# Modelling the demographic history of human North African genomes points to soft split divergence between populations

**DOI:** 10.1101/2023.11.07.565966

**Authors:** Jose M Serradell, Jose M Lorenzo-Salazar, Carlos Flores, Oscar Lao, David Comas

## Abstract

**Background:** North African human populations present a complex demographic scenario. The presence of an autochthonous genetic component and population substructure, plus extensive gene flow from the Middle East, Europe, and sub-Saharan Africa, have shaped the genetic composition of its people through time.

**Results:** We conducted a comprehensive analysis of 364 genomes to construct detailed demographic models for the North African region, encompassing its two primary ethnic groups, the Arab and Amazigh populations. This was achieved through the utilization of the Approximate Bayesian Computation with Deep Learning (ABC-DL) framework and a novel algorithm called Genetic Programming for Population Genetics (GP4PG). This innovative approach enabled us to effectively model intricate demographic scenarios, utilizing a subset of 16 whole-genomes at >30X coverage. The demographic model suggested by GP4PG exhibited a closer alignment with the observed data compared to the ABC-DL model. Both methods point to a back-to-Africa origin of North African individuals and a close relationship of North African with Eurasian populations. Results support different origins for Amazigh and Arab populations, with Amazigh populations originating back in Epipaleolithic times, as early as 22.3 Kya. GP4PG model supports Arabization as the main source of Middle Eastern ancestry in North Africa. The GP4PG model better explaining the observed data includes population substructure in surrounding populations (sub-Saharan Africa and Middle East) with continuous gene flow after the split between populations (migration decay). In contrast to what we observed in the ABC-DL, the best GP4PG model does not require pulses of admixture from surrounding populations into North Africa pointing to soft splits as drivers of divergence in North Africa.

**Conclusions:** We have built a demographic model on North Africa that points to a back-to-Africa expansion and a differential origin between Arab and Amazigh populations, emphasizing the complex demographic history at a population level.

## BACKGROUND

The North African region has a complex human demographic history with multiple migration events that have shaped the genetic makeup of its populations. Stone artifacts found in Algeria suggest that the first peopling of North Africa occurred around 2.4 million years ago [1]. However, the oldest human remains found in the region, at the Moroccan site of Jebel Irhoud, date back to 300,000 years ago [2]. Nonetheless, up to now there is no evidence that points to a continuity from these ancient humans to current North African people.

The oldest ancient DNA samples retrieved from North African individuals in the Taforalt site in Morocco date to Epipaleolithic times (15.1-13.9 Kya) [3]. The analysis of these Taforalt individuals shows a high affinity with Near Eastern Natufian populations. The presence of mitochondrial DNA haplogroups U6 and M1 in North Africans are consistent with a back-to-Africa event [3–5]. When compared with current inhabitants, the Taforalt ancestry component is present in all current North African populations following a West to East cline with highest frequencies observed on Amazigh individuals [6], pointing to a genetic continuity in the region at least since Epipaleolithic times.

Multiple migrations from surrounding regions have occurred in North Africa, leaving their genetic imprints on the local populations, which are characterized by an amalgam of ancestry components [7]. Most of these migrations originated in the Middle East, such as the Neolithic expansion associated with the spread of agriculture, which had a dramatic impact on the genetic makeup of North Africans, diluting the autochthonous Palaeolithic component in a similar demic process as in Europe [8]. In addition to the Neolithic migration event from the Middle East, recent data on ancient remains have shown that North Africa has also experienced gene flow from Europe in Neolithic times, as evidenced by the presence of Iberian genetic ancestry in Western North Africans in early Neolithic sites, suggesting a complex and heterogeneous gene flow in the region [9]. Furthermore, post-Neolithization gene flow has been observed in North Africa, resulting from events such as Arabization during the 7th to 11th centuries [10], the trans-Saharan slave trade from Roman times to the 19th century, and contacts with Mediterranean populations [7,11,12]. This complex pattern of migrations, with different temporal and geographical origins, has challenged the demographic reconstruction of North African population history.

Linguistics broadly classify the present populations of North Africa into two major groups: the Imazighen (Amazigh in singular), also known as the misnomer of Berbers, and the North African Arabs. After the Arab arrival to North Africa during the Arabization, most North African autochthonous groups adopted the Arab culture, and mixed with the immigrants [13]. However, a few groups retreated to remote places and maintained their customs, along with the Amazigh identity and language (i.e., Tamazight) [14,15]. Previous studies showcase large genetic heterogeneity within North African populations [6,7,15–18] with no clear differentiation between Arab and Amazigh populations as a whole. However, some Tamazight-speaking populations are genetic outliers with remarkable differences with their neighbouring Arab-speaking populations. This was attributed to isolation and drift, as well to differences in their demographic histories [6,18].

The genetic heterogeneity of North African populations plus the presence of an amalgam of genetic components, as a result of extensive gene flow coming from different areas and different time frames, have hampered the establishment of a demographic model for North African groups. To overcome this challenge, in this study we aim to reconstruct a demographic model of the Amazigh and Arab populations in North Africa that might be used as a neutral demographic model for future studies. We aim to tackle this issue by addressing three main questions. First, estimate the origin of North African groups and how they evolved. Secondly, address the number and amount of admixture events that have happened in the region. And finally, assess whether Imazighen and North African Arabs share the same demographic history. Overall, all these questions relate to the development of a demographic model that reproduces the rich and complex history of the region.

To answer these questions, we applied two different computational approaches. We first capitalized on the whole genome data of North Africa and used the Approximate Bayesian Computation framework coupled with Deep Learning (ABC-DL) [19]. Upon recognizing that the model identified by ABC-DL exhibited limited reproducibility with the observed data, we embarked on the development of a novel approach, Genetic Programming for Population Genetics (GP4PG), rooted in metaheuristics. GP4PG uses natural computing algorithms to infer the most optimum set of demographic events and associated parameters to explain the genetic variation observed in a given dataset.

## RESULTS

### Genetic Structure Analysis of North African samples

The Principal Component Analysis (PCA) performed on the whole genome dataset (see Methods) of North African individuals (N=30) and reference samples from Africa and Eurasia (N = 364), comprising 1.58 M SNPs, explains 8.64% of total variance on the first two principal components and recapitulates a similar population structure as previously described [6,7,12], with North African individuals clustered together in-between sub-Saharan African and Eurasian populations (Fig.1_a). A second PCA, focused on North African samples, shows in more detail the genetic structure of North Africa. East North Africa individuals cluster closer to Middle Easterns than West North Africans. Nonetheless, we observe a very heterogeneous pattern with the exception of the two North African Imazighen from Chenini (Tunisia) that cluster together isolated from the rest of North Africans (Fig. 1_b) (for full analysis see Additional file 1: Fig S1). The ADMIXTURE analysis (Fig. 1_c) identifies similar patterns to the PCA analysis. The lowest cross validation errors were found in the range between K=3 and K=9 (Additional file 1: Fig S2), with K=3, K=6, K=9 showing the least number of common modes among the different runs in pong [20]. At K=3 we observe the differentiation between sub-Saharan African (red), European (dark blue) and East Asian (light grey) components. All North African samples show similar ancestry patterns exposing an exclusive North Africa component at K = 9 (light blue) except for the Egyptian samples that show higher proportions of a purple component that is maximized in the Middle Eastern samples in K = 9 (Fig. 1_c). The rest of results at different K are in the Supplementary file (Additional file 1: Fig S3).

**Fig. 1:**
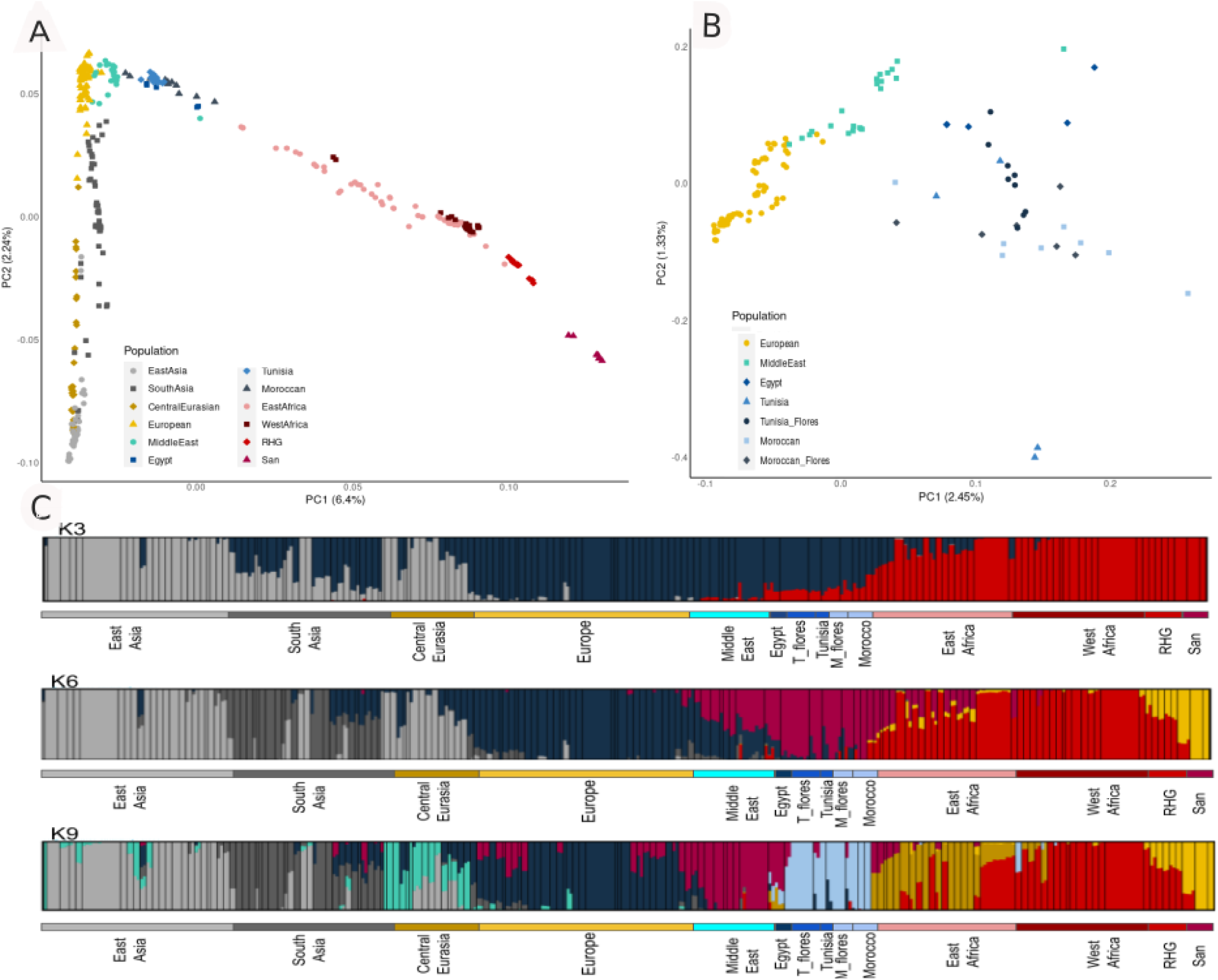
Genetic structure of North Africans and reference groups. **A**. PCA plot of all North African samples and the reference dataset. **B**. PCA plot focused on North African population substructure. **C**. Estimated ancestry proportions for the whole dataset in the three Ks most supported by pong [Behr2016].

### North African demographic model using an ABC-DL approach

Given these previous results showing the presence of population substructure in North Africa and a complex amalgam of genetic components, we conducted an Approximate Bayesian Computation (ABC) analysis coupled to a Deep Learning (DL) framework [19]. Our aim was to delve into the origins of contemporary North African populations and discern variations in the scenario based on the cultural backgrounds of these populations.

On one side, our dataset included North Africa Amazigh populations, considered the descendants of the autochthonous inhabitants of the region. On the other hand, we have also analysed North African Arab populations resulting from multiple cultural and migration events from the Middle East that vastly changed the genetic background of the populations from the region. We were also interested in measuring the genetic impact of migrations from neighbouring populations (Middle East, Europe, and sub-Saharan Africa) in both Amazigh and Arab groups.

We implemented seven demographic models where the North African groups diverge at different moments in time from the surrounding populations (Additional file 1: Fig S4): **Model A** considers North African groups as sister branches of West Africa. **Model B** shows North African populations splitting from the Middle East in a back-to-Africa movement. This is the model mostly supported in previous analyses by population structure and admixture-f3 approaches [6,7]. **Model C** implements a third variation of the possible origin of North Africa, placing it also in a back-to-Africa event prior to the split between Europeans and Middle Easterns. In **Model D** North African populations are split between Arab and Amazigh. Imazighen show a deeper origin splitting from the Eurasian branch, whereas Arab North Africans split more recently from the Middle Eastern branch. In **Model E** North African groups are separated, where Imazighen are a sister branch of West African populations while Arabs split from Middle Easterns. **Model F** represents the origin of Imazighen after the split of European populations but before the divergence between Middle Easterns and North African Arabs. Lastly, in **Model G** Imazighen appear as a sister branch of East African populations while Arabs split from Middle Easterns. Since all models comprised migration between North African populations and their surroundings, we did not include these parameters in the model ascertainment in order to improve the power of the DL prediction to discriminate between competing topologies.

After training the DL neural network for identification of the seven demographic models using as input the joint site frequency spectrum (jSFS) of each of the simulations to the observed data jSFS (see materials and methods), cross validation analyses using simulated data as observed showed that the ABC-DL can efficiently distinguish among the competing models. The minimum success rate for the discrimination is 67% and 76% for models **F** and **D** respectively, and the larger confusion is shown between these two models, which is expected as these are the two more similar topologies evaluated (Additional file 1: Table S1). The fact that **F** and **D** show such a level of misclassification is not surprising as both models show similar demographic topologies, only distinguishing on the branch where North African Imazighen diverge from the surrounding populations. In Model **D**, Imazighen have an origin that predates the European split while in Model **F** the Imazighen origin is slightly more recent, after Europeans have diverged from Middle Easterns. This similarity in the Models in addition with the fact that there is limited genetic differentiation between Middle Easterns, Europeans, and North Africans (Fig. 1), causes this misclassification rate in the ABC analysis.

When applied to the observed data, the posterior probability estimated by ABC-DL strongly supports model **D** (posterior probability of model given data =0.922) (Additional file 1: Table S2). Furthermore, only the models with Imazighen as a separate clade from Arabs and with a back-to-Africa movement show posterior probabilities greater than 0 (Additional file 1: Table S2). While both models are selected by the ABC, Model **D** represents the data 11.8 times better than Model **F** (Additional file 1: Table S3), presenting a robust result on the topology discrimination.

We further explored the **D** topology by adding migration pulses. We differentiate between migration and admixture pulses, with migrations referring to recent and continuous gene flow between two populations and admixture pulses defined as discrete and past gene flow from a target to a source population. We evaluated five different migration scenarios for Model **D** (Fig. 2): **Model D_1** represents Model **D** without migration or admixture events. **Model D_2** includes gene flow of both North African populations from/to surrounding populations. The model also includes several admixture pulses from Middle East to both Arab and Amazigh groups (which could mimic historic and pre-historic admixture events [13]) and admixture pulses to both North African populations from both East & West Africa representing that could represent the effect of trans-Saharan slave trade in the region [13]. **Model D_3** increases the complexity of **Model D_2** by including a “ghost” population from Eurasia. This population is described elsewhere [21] as a way of explaining basal substructure Out-of-Africa. We include two admixture pulses from this Basal Eurasian population, one to its sister branch and a second one to the European-Middle Eastern branch, representing a possible weaker influence of this Basal Eurasian on Amazigh respect to European, Middle East, and North African Arab populations. **Model D_4** is the most complex model since it includes a second “ghost” population in sub-Saharan Africa that represents the population substructure in Africa. This “ghost” population admixes with the “real” populations in Africa as described previously [22]. On top of that, this model also includes an admixture pulse from European populations to both North Africa [13] and one Middle Eastern admixture pulse to Imazighen. Finally, **Model D_5** reduces the complexity by excluding all admixture pulses that do not come from “ghost” populations. This model is a simplification of **Model D_4** to test if admixture pulses towards our target population are needed to explain the North Africa scenario.

**Fig. 2:**
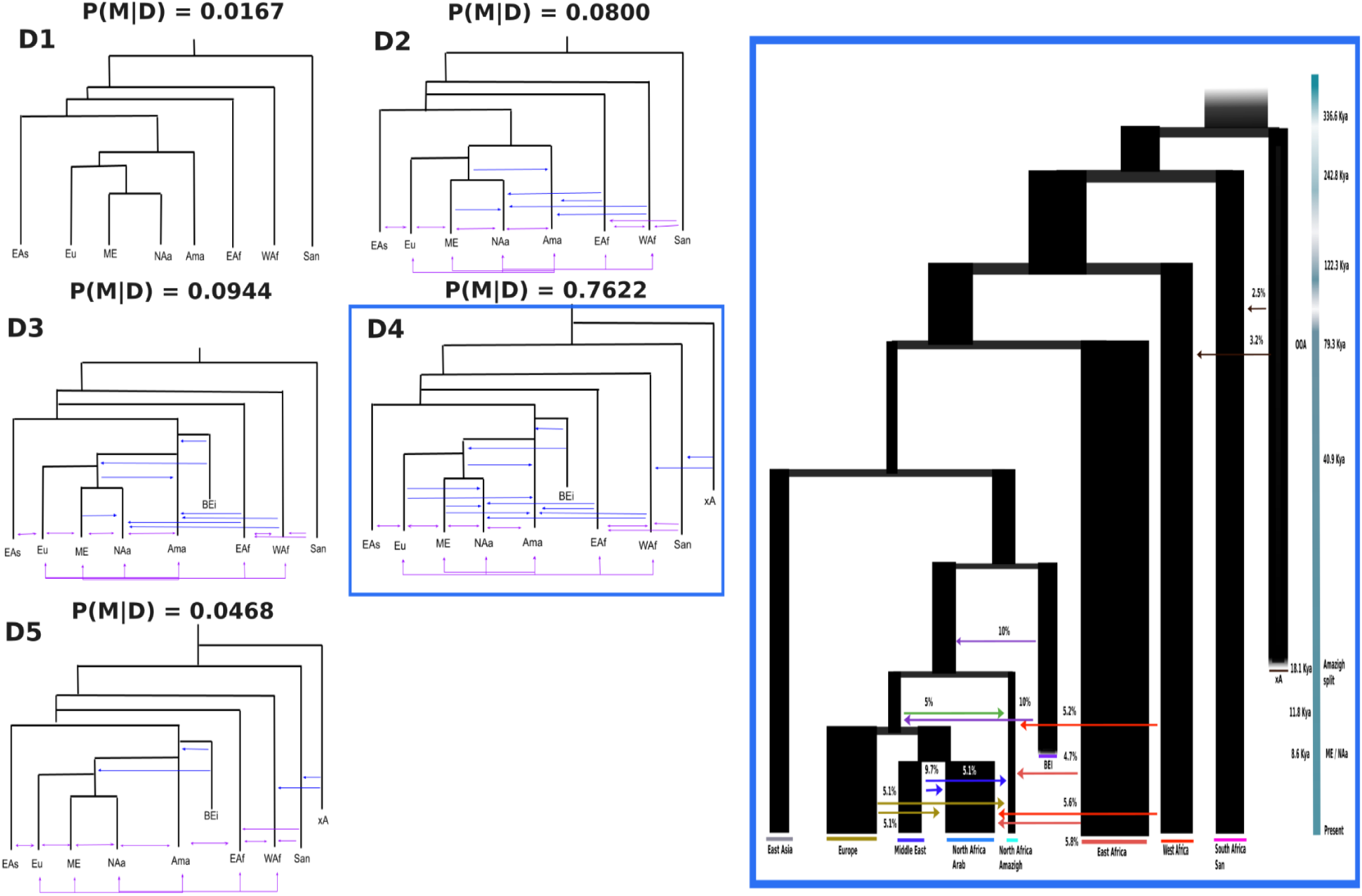
Tested demographic models including migration pulses. Left figures: variations of the selected topology on the first run of ABC-DL (Model D), included on the ABC-DL analyses considering North African Arab (NAa), North Africa Amazigh (Ama), Middle Eastern (ME), European (Eu), East Asian (EAs), East African (EAf), West African (WAf), and Ju/’hoansi (San) populations. Distinct levels of complexity are shown in each tested model, with purple lines representing recent migration and blue lines indicating admixture pulses from surrounding regions to both North African populations. The posterior probability obtained with our ABC-DL approach is shown on the top of each model. Right figure: fitted D_4 model with estimated parameters. Coloured lines represent admixture pulses from the specific population defined by the same color.

Cross validation analysis between these five models shows that ABC correctly identifies the model in 30-60% of the simulations (Additional file 1: Table S4). This limited sensitivity is compatible with the hypothesis that all models are remarkably similar between themselves with a lot of admixture events that may result in very similar statistics comparing one model to another.

When applied to the observed data, we obtained that **Model D_4** is the “best” model with 76.2% of accepted simulations (Additional file 1: Table S5) and a Bayes factor [23] of 8.07 to the second “best” model (Additional file 1: Table S6).

Next, we estimate the posterior probability for each of the 82 parameters form **Model D_4** by applying the ABC-DL approach [19,22]. For each parameter, we analysed to which extent the posterior distribution captures its real value. First, we computed the factor 2 statistic (Additional file 2: page 4: Table S1), defined as the number of times that the estimated mean is within the 50% to 200% of the true value of the parameter [23]. In most of the cases, the factor 2 analysis indicates high confidence in the estimation of the true value of each parameter. Time splits show significant better results than effective population size estimates, ranging from 98% (split of Europeans from the Middle East-North African Arab clade) to 100% in the West Africa split, compared to the 75% (effective population size of North Africa_ Middle East (NA_ME)) to 99% of West Africa effective population size (Additional file 2: page 4: Table S1). Regarding introgression parameters, introgression times show better factor 2 results (98% Basal Eurasian to European, Middle East, and North African populations) than the introgression amounts (77% NA_ME to Imazighen as the best value). In fact, the less accurate performance using the mean as proxy is for migration parameters. Factor 2 statistics show ∼ 70% chance of true value being in the 50% to 200% range of the estimated mean, which is similar to the value obtained when calculating the mean from random sampling of the prior distribution (Additional file 2: page 4: Table S1). Despite the weak performance in the estimation of migration parameters, the models with migration perform better in the ABC-DL than those without migration. These results indicate that the ABC-DL framework allows us to obtain a confident set of posteriors in most of the parameters, particularly for some of the more relevant ones since they show high factor 2 values and very significant differences between prior and posterior distributions (Additional file 2: page 4: Table S2). Of particular interest is the time of the split of Imazighen to the rest of European, Middle East, and North African populations. In 99.5% of cases, the mean posterior distribution of the time split of Imazighen to the rest of European, Middle East, and North African populations is within the factor 2. This suggests that the mean of the posterior distribution of this parameter can be considered a good proxy of the real value.

When the ABC-DL is applied to each parameter of the model (Fig. 2), we observed that the posterior distributions for a large number of parameters were significantly different from the prior distributions (Additional file 3: histograms; Additional file 2: page 4: Table S2) and that in most cases there is a correlation between predicted and simulated values in the ABC analysis (Additional file 4: Spearman correlation; Additional file 2: page 4: Table S3). According to the ABC-DL analysis, the North African Arab population diverged from Middle East common ancestor 8.6 Kya (97.5% credible interval (CI) ranging from 4.65 Kya to 15.40 Kya, assuming a generation time of 29 years per generation [24]). Imazighen split from the rest of Eurasian populations at 18.12 Kya (97.5% CI of 9.68 Kya to 27.33 Kya), whereas the Out-of-Africa event is estimated at 79.28 Kya (97.5% CI 51.11 Kya to 108.39 Kya). The effective population size indicates a reduction between the split with East Africa (Ne = 23,401 (97.5% CI: 4,465 to 39,557)) and the split with East Asia (Ne = 5,619 (97.5% CI: 1,292 to 9,713)), coinciding with the Out-of-Africa bottleneck. In some cases, the effective population size is slightly larger than the expected given our prior distributions (Additional file 3: histograms; Additional file 2: page 3). All parameters are estimated considering a mutation rate of 1.61e-8 ± 0.13e-8 mutations per bp [25].

The admixture estimates obtained from **Model D_4** show multiple migration pulses from Middle East, Europe, and sub-Saharan African populations towards both Amazigh and Arab populations in the last 200 generations. We observe that the amount of admixture from Middle East to North African Arab is larger than to Amazigh (9.7% [97.5% CI: 19.4% to 0.5%] vs 5.1% [97.5% CI: 9.8% to 0.3%]). Another interesting result is the 20% introgression from the “ghost” basal Eurasian population towards the MRCA of Middle East, European, and North African in at least two pulses of admixture. Other pulses of less intensity from East and West Africa to North Africa (∼ 5%) and from a “ghost” African population to San and West Africa are also observed (2.5% [97.5% CI: 4.9% to 0.2%] and 3.2% [97.5% CI: 4.9% to 0.4%], respectively).

Finally, we tested the robustness of the proposed model to generate datasets compatible with the observed genetic diversity in the data. We simulated 1000 datasets using the mean estimated at each parameter from **Model D_4** with *fastsimcoal2*. In order to quantify how similar each simulated dataset resembles the observed data, we compared the jSFS obtained for each simulation with the observed jSFS using a replication -unseen during the training of the DL-dataset by means of a PCA [26,27]. The results of the PCA indicate that the model does not correctly replicate the data since the observed jSFS falls as an outlier in the PCA of the simulations (Additional file 1: Fig. S5; Additional file 1: Fig. S6). This suggests that the current static models used might not properly capture the whole demographic complexity present in North Africa.

### Unravelling demographic history using Genetic Programming for Population Genetics (GP4PG)

To overcome the limitation of the ABC-DL approach in the reconstruction of North African population history, we have developed a novel approach to explore the demographic parameter and model topology space of a demographic model based on the paradigm of Genetic Programming. Genetic Programming is a meta-heuristic method inspired on evolution to generate formulae or programs coded as trees. Each node in the tree represents an operation or function, and the branches represent arguments or operands. An evolutionary approach Genetic Programming for Population Genetics (GP4PG) algorithm considers evolutionary events, such as changes in the effective population size or the increase or decrease of population substructure, as operations. GP4PG search the space of possible configurations of evolutionary events to define the demographic events that would generate genomic datasets similar to the observed ones (see material and methods). These demographic events include the ones already considered in ABC-DL, such as the presence of admixture, effective population sizes, or time of the demographic event. However, GP4PG allows modelling population substructure within each of the considered populations, or ecodemes [28,29], from the demographic model. In this framework, each ecodeme is formed by multiple topodemes that relate to each other following an isolation by distance pattern [30], where topodemes that are situated closer will migrate at higher rates than topodemes that are further apart (Additional file 1: Fig. S7). This allows generating reticulated and partially reticulated demographic models [31].

In addition, this new algorithm also takes into account population substructure when exploring different demographic scenarios. In our case, we considered eight ecodemes: San, West Africa, East Africa, Middle East, East Asia, Europe, North Africa Amazigh, and North Africa Arab, all with the same size and without distance between neighbouring ecodemes to simplify the models.

We apply the GP4PG algorithm with the six of the considered topologies used with the ABC-DL. We discarded **Model A** since it consistently showed the worst performance in multiple iterations of the ABC-DL reducing computing times and resource allocation. For the remaining six models, we used two different versions: a variant that includes migration between topodemes (populations) and a variant without including migrations resulting in 12 different topologies (Additional file 1: Fig. S8). Since GP4PG is a metaheuristic approach based on exploring graph topologies (see material and methods), it can be trapped into local optima. Therefore, we independently run 40 times the GP4PG algorithm, each for 200 iterations, retrieving the best demographic model from the end of the 200 iterations. After the 200 iterations the fitness error between the simulated dataset and the observed dataset at each run of GP4PG reaches a plateau (Additional file 1: Fig. S9). Interestingly, we observe differences in the final error of each run, ranging between 10 to 2, thus supporting that GP4PG tends to get trapped in local optima.

**Model D** is the one most supported by the GP4PG with 10 out of the 40 replications performed. The model that presents the minimum error is **Model D** (Model D_15 – Error = 2.56). Next, we assessed the performance of each model to predict the observed data. We selected the 10 models from the GP4PG runs with the least amount of fitness error. For each of them, we computed a thousand simulations. For each simulation we computed the 4-fold SFS of all the simulations. Finally, we compared them to a replication set of the observed data using PCA. We compared the results of the ABC-DL simulation, transforming the jSFS into a 4-fold SFS to evaluate if the performance of the GP4PG is better than the ABC-DL (Additional file 1: Fig. S9). Simulations from the ABC-DL **ModelD_4** fall further to the observed data than any of the GP4PG models in the PCA indicating that the models produced by the new methodology are consistently better at describing the observed data than the models produced by the ABC-DL (Additional file 1: Fig. S9).

After verifying that simulated data from models inferred by GP4PG fit the genetic diversity observed in the real data, we ascertained the model interpreted as the “best” after both ABC-DL and GP4PG (Fig. 3). This demographic model presents a topology that proposes different demographic histories between Amazigh and Arab populations, with Amazigh splitting from the MRCA around 22.3 thousand years ago. On the other hand, North African Arab individuals split from Middle Easterns around 1.6 Kya, which might be related to the Arabization process in North Africa. This result contrasts with the one obtained with the ABC-DL approach that estimated an older split of North African Arabs closer to Neolithic times. Another detail that we deduced from this demographic model of North Africa is that admixture pulses do not appear as drivers of genetic diversity in any of the best 10 models. Instead, most of the current genetic diversity can be explained by a combination of population substructure and migration decay between demes. The rest of the parameters of each model can be found in the supplementary material (Additional file 5: parameters for the best 10 models of the GP4PG).

**Fig. 3:**
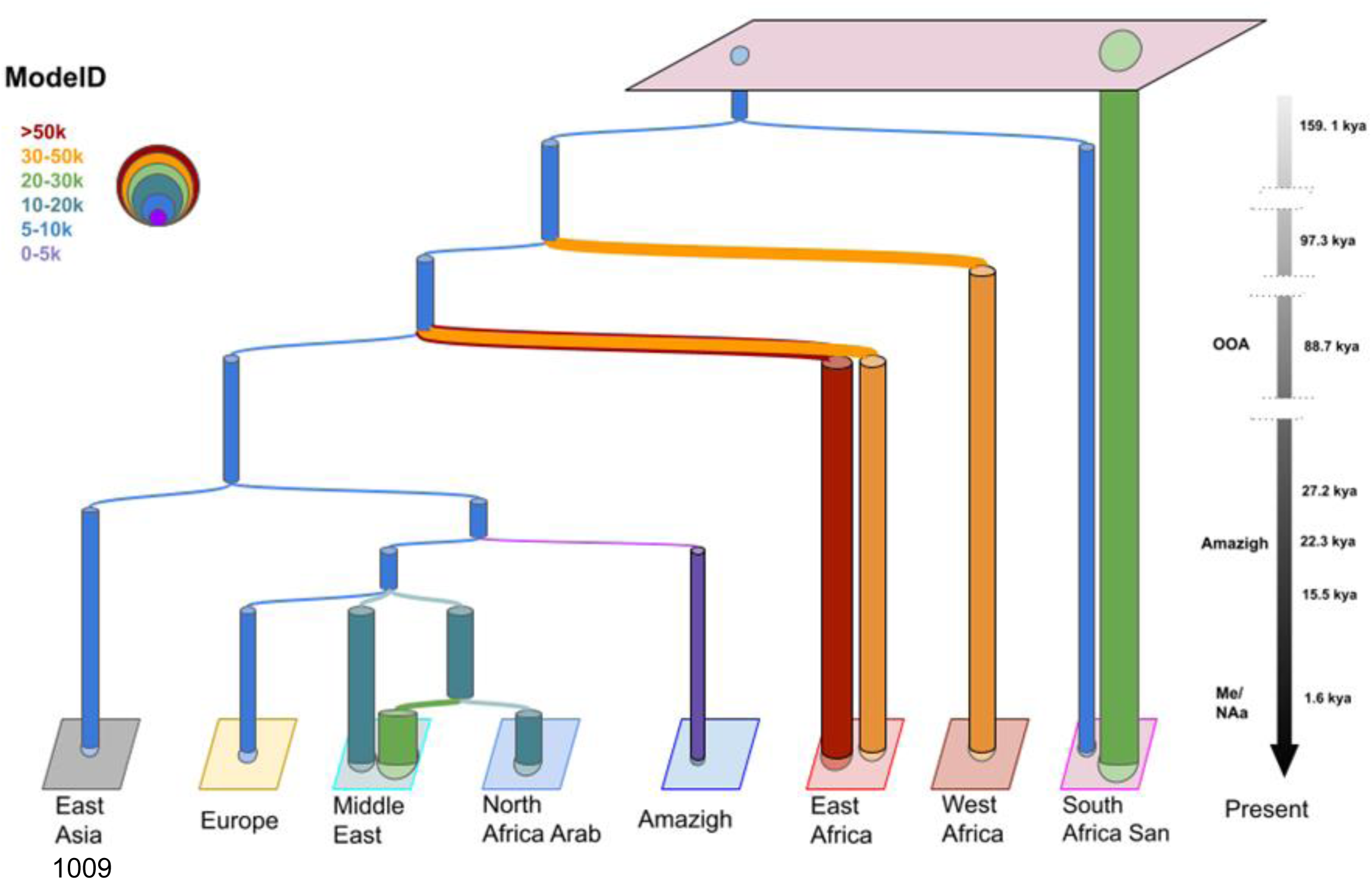
Fitted model obtained with GP4PG algorithm. Demographic model of North African populations with the least amount of error in the GP4PG algorithm (E = 2.56), iteration 15 out of 40. This model presents a different origin of North African Amazigh and Arab. Each color represents a different range of Ne also represented by the width of the columns. The model shows some population substructure, especially in sub-Saharan African populations and presents the ability to explain complex demographic models with population substructure and decaying migration after split patterns.

## DISCUSSION

North African populations are a demographic melting pot [7,13]. The presence of complex and recent demographic histories with multiple gene flow events from different sources difficult our ability to construct a feasible demographic model with current methodologies. The absence of model-specific statistics recapitulating particular events even for moderately complex evolutionary problems has fuelled the development of methods based on machine learning [32].

The application of the ABC-DL approach considering relatively complex demographic models supports that the Imazighen present a different demographic history compared to surrounding populations. Prior studies with autosomal markers on North Africa populations pointed to Amazigh populations presenting a unique genetic component that has been described as the autochthonous component for the region and that can be traced back to, at least, Epipaleolithic times [3,7]. This is supported by our ABC-DL model as we estimate a continuity of the Amazigh population up to 18.14 Kya (97.5% CI of 9.68 Kya to 27.33 Kya) falling in the range of the Epipaleolithic samples of Taforalt (15-13 Kya) [3]. The Arab population in North Africa, on the other hand, shows higher genetic affinity with Middle Eastern groups than to the ancient Taforalt samples. This affinity is described by an east to west cline on the amount of Middle East-like genetic ancestry across North Africa and has been hypothesized as a consequence of several migration movements from the Middle East to North Africa with the Neolithization and the Arab expansion being the more relevant ones [13]. The ABC-DL model supports the Neolithic expansion to North Africa as the main source of the Middle Eastern component. This is supported by divergence times between North African Arabs and Middle Easterns at 8.6 Kya (97.5% CI of 4.65 Kya to 15.40 Kya) which overlaps with previous studies on Neolithic expansion in North Africa [8]. We observe at least one pulse of admixture in late Neolithic from the Middle East to both North African populations around 5.6 - 5.9 thousand years ago (97.5% CI of 0.54 Kya to 13.94 Kya to Arab; 97.5% CI of 0.63 Kya to 14.03 Kya to Amazigh), highlighting the importance of Neolithic demic diffusion in North Africa [8,33]. Other admixture events from surrounding regions are also supported by our model. European admixture in Neolithic times is present, although always placed later than the Middle Eastern admixture (Additional file 2: page 4). This is observed both in both Amazigh and non-Amazigh populations in small proportions (∼5%) which is consistent with the hypothesis of European contact in western North Africa at least 5,000 years ago [8–10]. In addition to these gene flow events, it has been proposed the presence of a basal Eurasian population that could explain early diversity in West Eurasia, North Africa, and the Near East [21]. We observed that, when including this “ghost” basal Eurasian population in our models, the performance of the models increases, reinforcing the hypothesis of the basal Eurasian population proposed previously [21]. Focusing on the performance of the ABC-DL approach in North Africa, some of the main flaws are related to the estimation of effective population sizes. In some populations the posterior distribution of effective population size is larger than the initial prior distributions we provided. This might be an issue when estimating several parameters since effective population size and split times are highly correlated. Effective population size is dependent on heterozygosity [26]. As the heterozygosity increases, the effective population size or the split time of a population needs to increase in order to reach coalescence [26]. We observe that the heterozygosity in the North African Arab samples is higher compared to European and Middle Eastern populations (Additional file 1: Fig. S10) which can be a result of recent gene flow from sub-Saharan Africa. This increased heterozygosity hinders the estimation of split times and effect sizes as we may need bigger sample sizes or more complex scenarios to take this diversity into account.

Despite the reasonable performance of the ABC-DL approach, it has replicability issues. The accepted models cannot grasp all the diversity of the data and are biased by our preconceptions of the history of the populations. Moreover, a further drawback of the methodology is the black-box nature of DL approaches [34], which limits interpreting which parameters from the model should be modified in order to improve the performance of the inferred demography. To overcome these limitations, we have developed a new algorithm, GP4PG, inspired by the field of genetic programming and evolutionary algorithms [35–38], for automatizing the exploration of complex parameter-free demographic models. GP4PG performs, in terms of reproducing the observed genetic variation, better in the North Africa scenario than the best possible model obtained with ABC-DL algorithms by minimizing the difference between our dataset and the simulations obtained from the best model (Additional file 1: Fig. 9). It also reduces the amount of bias that can be imputed due to the building of each of the competing models, given that the only inputs to the GP4PG algorithm are the topologies for each of the models.

The GP4PG analysis, like the ABC-DL, supports that Amazigh and Arab populations in North Africa have different demographic histories despite both originated as back-to-Africa movements from Eurasian populations. Amazigh groups appeared before the European, Middle Eastern, and North Africa Arab branch, splitting from a common ancestor around 22.3 Kya (Fig. 3). This date precedes the oldest genomic data available in the region [3] for at least seven thousand years, pointing to a genetic continuity even before than expected, and which is closer to the earliest known appearance of Iberomaurisian culture in Northwest Africa (25.85 to 25.27 cal. Kyra) at Tamar Hat [39]. On the other hand, North African Arab populations appear to have split from Middle Easterns 1.6 Kya, placing this population closer to an Arabization replacement on North Africa than to a Neolithization demic diffusion. This result contrasts with the ABC-DL results that show a deeper impact of the Neolithization process in the North African Arab - Middle Eastern split. The difference in the divergence time can be attributed to the effect of migration. From a coalescent perspective, migration between two populations generates a distinction between the coalescence time between the lineages and the split time between the populations. In models where migration is included as a parameter, the split time between North African Arab and Middle East populations is closer to the one observed in the ABC-DL analysis (ModelD_7 at 6.41 Kya; ModelD_26 at 13.39 Kya; Additional file 5) implying that the inclusion of migration increases the time of coalescence between the populations.

The absence of admixture events in the best model of the GP4PG algorithm can be attributed to the fact that our algorithm is weighted towards modelling population splits by a soft process in which migration decays forward in time between related populations. Therefore, whenever a split between two populations occurs, migration between the newly formed topodemes continues for generations, decaying as the demes grow further apart until they achieve a constant migration rate. Our selected model presents population substructure in current sub-Saharan African populations that extends to ancient times (Fig. 3). These observations support the results obtained by Ragsdale et al [31], where reticulated population substructure tens of thousands years ago could explain some of the genetic diversity previously attributed to archaic introgression [19,22]. Although the most robust model supports this idea of soft splits between populations, we do not rule out the possibility of admixture pulses as we observe in the ABC-DL analysis because we hypothesize that the effect of continuous migration after split could be mixed up with the effect of admixture pulses.

Since our analysis includes a limited number of samples (*n =* 2 per population), the study lacks some power to confidently corroborate some of these results, especially for the sub-Saharan population. Despite this, the results indicate that the models always perform better when including some amount of population substructure. On top of that, we must be aware that the selected Imazighen individuals are part of a very isolated population, the Chenini Amazigh. This population is an outlier in North Africa (Fig.1_b) due to isolation. This characteristic makes it useful as a proxy of the North Africa autochthonous component given that the amount of sub-Saharan and Middle Eastern genetic components is lower than in the rest of North African Imazighen groups [40]. Nonetheless, taking into account the heterogeneity within Imazighen groups due to different amounts of genetic components coming from neighbouring populations, using other Imazighen groups could lead to slight differences in some of the studied parameters.

Most of the selected topologies in the 40 runs of the GP4PG are either Model D (25%), Model C (17.5%), or Model F (20%). These three models present very similar topologies, with slight variations, mainly in setting the Amazigh origin. The current implementation of the GP4PG algorithm has enough statistical power to discriminate between competing models but falls short to detect fine scale migration and admixture events. This is due to the multimodal nature of the SFS that can lead to similar genomic patterns [26,41] with different demographic models. Implementation of haplotype-based summary statistics to the GP4PG algorithm in the future could solve some of these issues in very complex demographic scenarios such as that of North Africa.

## CONCLUSION

In sum, we have built a robust model of the demographic scenario of the North African populations. By implementing an ABC-DL algorithm and a novel GP4PG algorithm based on metaheuristics, we have defined a clear topology that proposes a back-to-Africa origin and differentiates the Amazigh-speaking population from the Arab-speaking groups in the origin and settlement in northern Africa. Our data point to a complex scenario where population substructure and admixture events had a significant impact on the genetic structure of current North African populations.

## MATERIAL & METHODS

### Databases

To analyse the population structure and demographic history of North Africa a dataset was compiled consisting of 32 whole genomes sequenced on deep coverage from North Africa. This dataset includes fifteen newly published samples from Morocco (*n* = 6) and Tunisia (*n =* 9) *(Ref : EGAXXXX),* Imazighen (*n* =4) and non-Imazighen (*n* = 6) individuals from Serra-Vidal 2019 [6], Egyptian samples (*n* = 3) from Pagani et al. 2015 [42] and Saharawi (*n = 2*) and Mozabite (*n* = 2) North Africa individuals extracted from the Simons Genome Diversity Project (SGDP) [43]. These North African samples were merged with a panel of world-wide populations from the SGDP (*n =* 295) [43], the 1000 Genomes Project (*n* = 38) [44] and high coverage Qatari individuals (*n =* 9) from Fakhro et al [45]. The final dataset used for the population structure analysis consists of 374 whole genome high coverage individuals. Furthermore, to perform the demographic inference analysis, a subset of individuals from each of the proxy groups for every population was analysed (Additional file 1: Table S7). Two North African Arab speaking population (Tunisian Arab), two from a Tamazight speaking group in North Africa (Tunisian Chenini), two Middle Eastern representatives from Qatar, two Northern European from Utah (CEU), two East Asian from Han (CHB), two West African from Yoruba population in Ibadan (YRI), two East African from Luhya in Kenya (LWK) and two South African San from Ju/’hoansi North in Namibia (JHN). The final dataset for the demographic inference analysis comprised 16 individuals with an individual whole-genome coverage of >30X.

### Read Mapping and Variant Calling

Single-nucleotide polymorphism (SNP) genotype variation of each sample was obtained by the following procedure. Read quality assessment of the fastq files was performed with fastQC (https://www.bioinformatics.babraham.ac.uk/projects/fastqc/). Reads were mapped to the hg19 reference genome using the Burrows-Wheeler Aligner (BWA-MEM v0.7.13) [46]. Reads were then sorted using SAMtools v1.2 [47] and duplicates were removed using MarkDuplicates from Picard (https://broadinstitute.github.io/picard/). Indels were realigned and quality scores were recalibrated using the Genome Analysis Toolkit (GATK 4.1.8.1) [48]. Variant calling was done using the HaplotypeCaller and merging of each sample into a multisample VCF was done using the GATK GenomicDB-GenotypeGVCFs functions [48].

We obtained the structure dataset by merging all samples, keeping the common SNPs across all samples, and applying SNP (geno = 0.1) and individual missingness (mind = 0.1) in PLINK v1.9 [49–51]. We then excluded the SNPs with MAF < 0.01 and removed the individuals with cryptic relatedness using KING [52] based on a cutoff of 0.325. The final dataset comprised 9.68 M SNPs on 365 individuals from 150 different populations.

### Data filtering

For demographic modelling, the dataset was further filtered to obtain a confident set of variants by implementing the following criteria: (i) a minimum of 5 reads mapped for each locus, (ii) a quality score threshold for the alternative allele of the variant, with a minimum score of 20 in the QUAL field of the VCF file, (iii) a PASS in the genotyping quality, (iv) exclusion of regions covered by structural variants [22] using TandemRepeatMarker repeats of length greater than 80 bp (UCSC browser) and 1000 Genomes Project copy number variants

(https://www.ncbi.nlm.nih.gov/dbvar/studies/studyvariants_for_estd199.csv), (v) exclusion of regions adjacent to indels with a 6bp flanking region, and (vi) exclusion of multiallelic variants. Based on these filtering steps, we obtained a 1.962.660.202 bp-long callable genome containing a high-confidence set of 16.4 M SNPs for downstream analyses. The ancestral state of each variant in these genomes was set to the chimpanzee reference genome (panTro4 genome assembly from http://hgdownload.cse.ucsc.edu/goldenPath/hg19/vsPanTro4/reciprocalBest) to avoid any discrepancy between African and non-African populations as detailed elsewhere [19,22].

### Structure analysis

#### Principal component analysis

Principal component analysis was performed using flashPCA [53], pruning the data for linkage disequilibrium between the markers using PLINK v1.90 [49–51] based on an r^2^ threshold of 0.4 in every continuous window of 200 SNPs with a step of 25 SNPs (i.e., – indep-pairwise 200 25 0.4).

#### ADMIXTURE analysis

ADMIXTUREv1.3 [54] was applied on the whole structure dataset, which was previously pruned for linkage disequilibrium between markers (–indep-pairwise 200 25 0.4). ADMIXTURE-ready dataset had 1.58 M SNPs on 365 samples. ADMIXTURE in unsupervised mode was run assuming several ancestral clusters ranging from K = 2 to K = 12 with 10 independent runs for each K using different randomly generated seeds for each run. The cross-validation error was assessed for each run, with K = 3 to K = 9 giving the minimum error. We then ran pong [20] in the greedy mode in order to identify common modes among different runs for each K to align clusters across different values of K.

### Demographic Model

To decipher the complex demographic scenario of North Africa, we used an Approximate Bayesian Computation with Deep Learning as explained more in detail in Mondal et al. [19]. The ABC-DL implementation is a three-step analysis. First, we generated thousands of simulations with fastSimcoal2 [55,56] for each of the competing models using the joint multidimensional site frequency spectrum among populations (jSFS) as a raw summary statistic. This statistic contains the information to run most of the frequency-based statistical analyses used in population genetics which are informative for detecting most of the demographic parameters considered in our demographic models (more in [19]. Second, we trained a DL to predict, from the jSFS, the most informative summary statistic (SS-DL) of the considered parameter or set of models. A potential limitation of this approach is the fact that the DL is trained with simple data and compared to the real model generated by the observed data, possibly overfitting our models. To avoid biases in the DL prediction, we injected jSFS noise in each simulation from the real data (see [19]). Finally, we performed a classical ABC approach using the SS-DL in a new set of simulated datasets.

The callable fraction of the genome we used for the demographic modelling is a modification of that defined by Pouyet [57]. We further cleaned the original callable genome, defined by Pouyet, to identify neutral regions by masking genomic regions containing Ensembl genes within a 20 kb range and masking CpG islands as defined elsewhere [19]. After that, we excluded all regions failing to reach a SNP density over 90% on 10 Kb windows with a sliding step of 2500 bp. to obtain 53.7 Mb of callable genome that was used in the next steps.

Ten Deep Learning (DL) networks were created, with four hidden layers each, and trained with 20,000 simulations each. An additional set of 180,000 simulations per model was generated, injecting noise from the observed jSFS of an individual of each population (BTUN01, TUN01, NA18559, NA12878, SR098230, NA19037, NA19207, HGDP01032), and the probability of each model was predicted using the 10 DL networks. The results were averaged to get the summary statistic (SS-DL) for the Approximate Bayesian Computation (ABC) analysis, carried out using the “abc” package in R [58,59]. We have assumed a mutation rate of 1.61e-8 ± 0.13e-8 [25] and a generation time of 29 years [24].

The ABC process can be divided into two steps. First, we checked how well the “abc” was able to distinguish between competing models. To do so, we applied the cv4postpr function on the “abc” package that performs a cross validation analysis on the ability of distinguishing between models [58]. This was done using 50 simulations per model and the same ABC parameters we used when analysing the observed data. The cv4postpr runs the ABC using simulated data as observed data and counting the number of times that the model with the highest posterior probability was, in fact, the model that generated the simulated data. Once the “abc” was able to distinguish between the different competing models, we began the discrimination of the “best” model. To do so we applied the “postpr” function of the “abc” package [58], keeping the 1000 best simulations (out of a total of 180,000 simulations per model) under a “mnlogistic” option for model comparison (tol = 0.001). We applied this procedure twice, one for the discrimination between the 7 competing models and another one to select the best variant for the topology selected before.

The DL process and ABC calculation were repeated for the parameter estimation of each parameter in the “best” model to get the posterior range for all demographic parameters. Mean, median, mode, 95% Credible Interval (CI) and 95% Highest Density Interval (HDI) [60] were calculated for each parameter. Finally, Spearman correlation, factor2 [23], and Kullback-Leiber distance [61] were used to assess the quality of the posterior predictions.

### GP4PG

Model comparison by ABC requires defining which are the models to consider. These models are, by definition, simplifications of reality. However, basic assumptions about the demographic events, and particularly population substructure, can significantly bias the model ascertainment [31]. Previous studies have dealt with the presence of hidden population substructure by modelling “ghost” populations [19,21,22] or by generating ancient weak structured stems that interact forming the current populations [31]. To bypass this issue, we have developed the Genetic Programming for Population Genetics (GP4PG), which is based on using genetic programming (GP), a branch of natural computing.

Natural computing refers to meta-heuristic algorithms inspired by nature to solve –by means of optimizing an error function– complex problems that are otherwise intractable. The underlying rationale of natural computing is that strategies used by nature to solve natural problems can be applied by reverse engineering to human-based-problems. Within the context of natural computing, Darwinian evolution inspires a broad family of algorithms called Evolutionary Algorithms that mimic the process of how evolution works to adapt an organism to its environment. Within the evolutionary algorithm family, Genetic algorithms have been already used in population genomics for demographic parameter definition [62]. However, GP is better suited for generating formulae and population relationships as the algorithm codes solutions in the form of a graphical (tree) structure whose nodes or edges represent parameters [37,38]. GP is an automated invention machine, which routinely delivers high-return human competitive machine intelligence, duplicating the functionality of previously patented inventions, infringed a previously issued patent, or created a patentable new invention [63]. The basic workflow of a GP embraces the basics of the biological evolution of a population, including selection, recombination, and mutation. Within this machine learning framework, a proposed solution, coded in the form of a graph, is considered as a biological being, which is subject to selection according to how good is for predicting the parameter of interest –how good is the summary statistic to distinguish models, or to predict a parameter of a given model. A set of solutions define a population that evolves over generations, exchanging information through recombination and exploring the surrounding space through mutation, to minimize an error function. Within the context of demographic modelling, the nodes refer to possible demographic events – and the tree depicts a demographic model (Additional file 1: Fig. S11; Additional file 1: Table S8).

The proposed GP4PG framework considers the particular features of demographic modelling and applies some modifications to the classical GP algorithm to account for them. GP4PG organizes populations in fundamental homogeneous groups called “demes” that are nested within “topodemes" and “ecodemes” [29]. Gilmour and Gregor defined these concepts from an ecological perspective. Ecodeme refers to those topodemes sharing a given habitat, while topodeme is used to group the individuals that are from the same locality. Each ecodeme can have one or more topodemes depending on the heterogeneity of the population. In a population genetics and demographics scenario, we used topodeme as a unit of population substructure to resemble the different population nuclei that we could observe in a population and that suggest the internal diversity that be observed in a given region (“ecodeme”) [28].

The GP4PG algorithm models the migration rate of each ecodeme within itself and with adjacent ecodemes following an isolation by distance approach [30]. Thus, populations situated further apart would have less chance to exchange migrants than those that are closer together. For the sake of simplicity, we considered that populations all occupy a same size space and that the distance between adjacent populations was zero, meaning that geographical barriers, such as the Sahara Desert or the Mediterranean Sea, were not considered to order the populations in the space. All models included a migration decay function. Following the split between two ecodemes, these demes continue to exchange migrants, but the quantity is gradually reduced over time until it reaches the migration rate defined between the two populations or they stop sharing migrants if the models do not have migration. This modelling approach accounts for the fact that the migration rate between two populations is not constant from the outset, but rather fluctuates and diminishes exponentially with time as they become more isolated and distinct.

First of all, the nodes used in GP4PG represent demographic events rather than operations to be added to a formula. Each demographic event requires a time when it occurs, and such time determines the relationship with its preceding nodes in backward. The GP4PG will also choose from the several possible demographic events (admixture, addition and reduction of demes or changes in the migration rates) and different combinations to produce the simulations that are going to be tested against the observed data. All the simulations were ranked by means of a standardized error comparing the jSFS (4-wise SFS) of each of the simulations to the observed data jSFS. The best simulations will, following the concepts of GP, produce more offspring than the worst simulations, that will suffer modifications using mutations and recombination with other models, similar to an evolutionary process, optimizing the models after each generation. In this way, at each iteration, we’ll obtain better models to explain the observed data.

### Exploring the space of possibilities within the GP4PG framework

In canonical GP, exploration of the space of solutions is mainly accomplished by generating solutions from the most successful ones by means of recombination: exchanging sub-trees at a given node in both parents. Thus, the offspring is a combination of both trees using a subtree-crossover operator [36]. In the GP framework, modifications of the parental structure -i.e., mutation-are less frequent [36]. However, classical recombination approaches applied to demographic models could easily produce non-compatible solutions, where the root node of the replaced subtree from one parent could have older times than its preceding node. This does not occur if only the mutation process is applied, as in this case changing the time can be constrained to be between the ranges of the previous and next demographic event. There are different evolutionary strategies and evolutionary strategies-like algorithms whose main exploration force is a type of mutation. We adapted the invasive weed optimization algorithm (IWO) [64,65] to be used for GP-tasks. IWO emulates the process of colonizing new environments by invasive plants. In IWO, each solution present in the population reproduces proportional to its fitness. First, a finite number of seeds (demographic models) is produced. Reproduction of each solution depends on a seed offspring function [Eq.1] that lets those solutions that have better fitness reproduce more than those that are further from the optimum. This reproduction technique allows the chance to survive and reproduce for unfeasible solutions, hence not discarding possible useful information carried by low fitness individuals. Once the new solutions are proposed proportional to how good each parental solution is, the new solutions are ranked from the best to the worse and a new population of the original size is generated by disregarding worse performing solutions.

In our case, we rank all the possible solutions according to how close the simulated dataset produces summary statistics (SS) close to the summary statistics observed in the real data (Eq. 1):

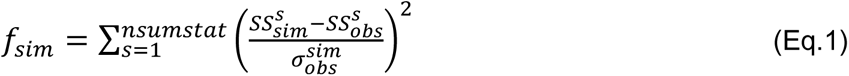

Where each element s comes from the jSFS of each simulated model computed among all possible combinations of four populations (4jSFS) against the 4jSFS of the observed data. The standard error of each element *s* is obtained by Monte Carlo resampling with replacement from the considered genomic fragments 1,000 datasets of the same genomic size as in the training dataset and computing for each dataset the 4jSFS.

By using 4-population-fold jSFS instead of the full multidimensional SFS (mSFS) among all populations, the total number of summary statistics is reduced to Eq. 2 instead of Eq. 3. This avoids the exponential explosive nature of the full multidimensional site frequency spectrum and the associated curse of dimensionality [66], reducing the number of mSFS combinations with value 0, while allowing to recapitulate the demographic relationships among populations [67]:

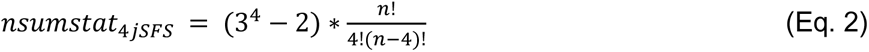

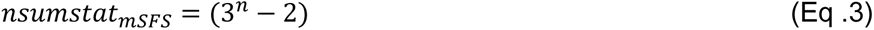

### Evolving the population of answers

The best performing solution in our population (i.e., the one whose demographic model produces 4jSFS close to the one in the observed data) generated eight new solutions, each of them showing in probability differences on the time of the events, the events and particular parameters related to the events. The worst performing solution in our population produced two offspring. Other solutions reproduced in proportion S (Eq.4). In our analysis we have determined a *S*_*min*_ of 2 and a *S*_*max*_ of 8, this allows the preservation of low fitness solutions that could potentially give higher fitness offspring that otherwise with a more restrictive *S*_*min*_ would be lost:

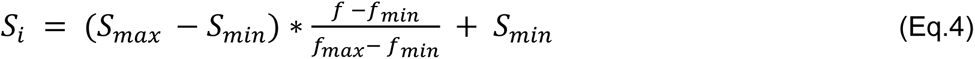

### Model comparison with GP4PG

We tested six competing topologies in the GP4PG algorithm (Fig. 4), which are the same topologies tested for ABC-DL except for **Model A**. On top of that, we constructed two models for each topology, one considering migration between “ecodemes” and the other without it. For the GP4PG algorithm, in order to speed up the algorithm, we have used half the masking regions filtered from Pouyet [57], and a second set of data to validate the analysis. The GP4PG algorithm gives us the model that presents the least amount of error to the data for each iteration. In our case we have performed 40 iterations with 200 generations for each iteration. The 10 best solutions were then compared to the observed data by performing a PCA of the 4jSFS of a thousand simulations of each model against the 40 4jSFS of the replication dataset to take into account the deviation in the SFS due to the random selection of masked regions.

**Fig. 4:**
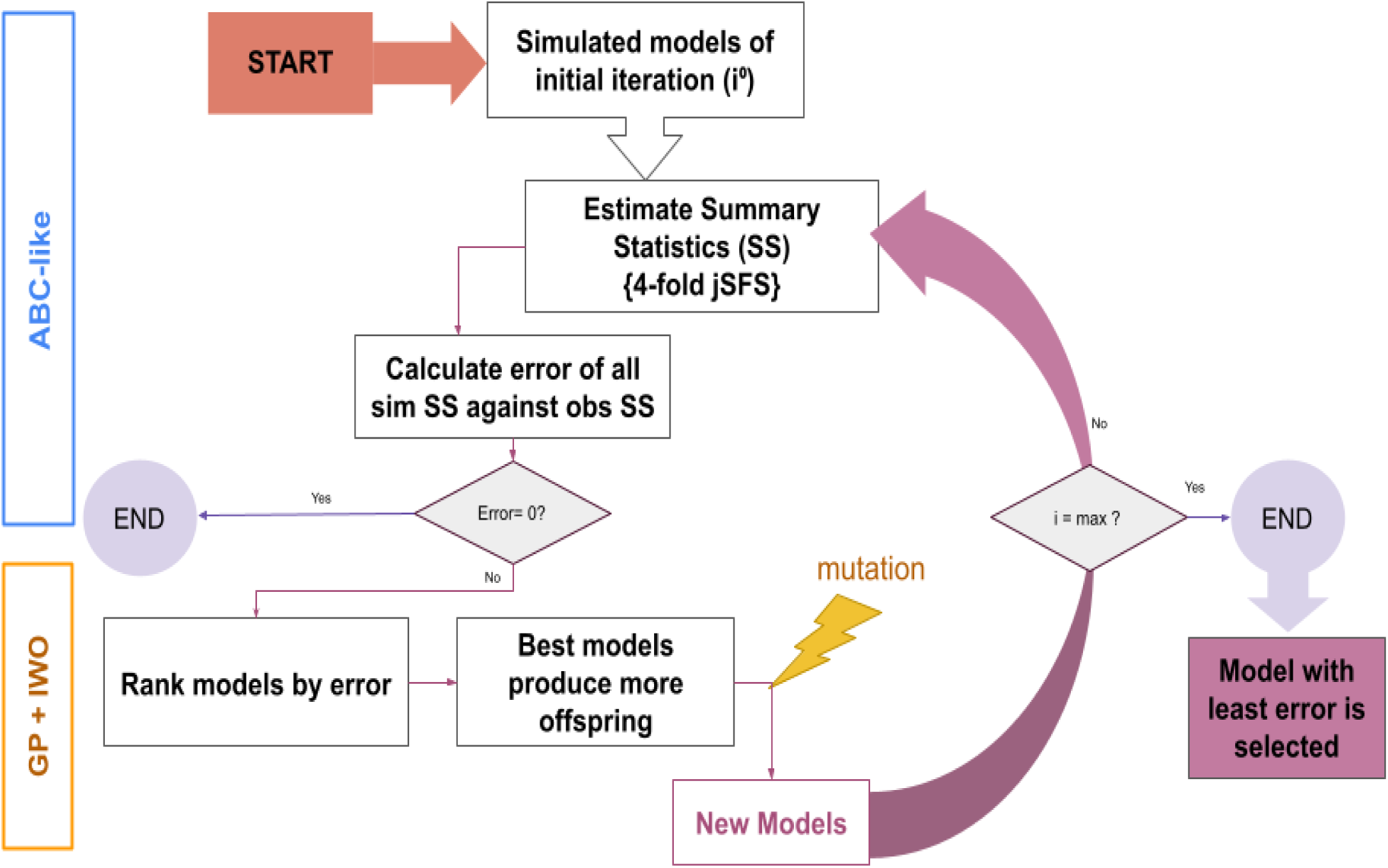
Structure of the GP4PG algorithm. From an initial set of models, we compute a summary statistic (4jSFS) for each simulation and compare it to the observed data by the means of a fitness error function. Then, the errors are ranked. We produce the offspring (2nd generation) following an invasive weed optimization algorithm modifying each child model (mutation). We repeat this procedure until the error is 0 or it reaches a plateau.

## Supporting information

Supplementary File Description

## DECLARATIONS

### Ethics approval and consent to participate

The present project has the corresponding IRB approval (*Comitè d’Ètica d’Investigació-Parc de Salut Mar* 2019/8900/I, Barcelona, Spain).

### Consent for publication

The authors declare no competing interests and agree with the publication of the results.

### Availability of data and materials

#### Funding

This work was supported by the Spanish Ministry of Science and Innovation (grant numbers PID2019-106485GB-I00 and RTC-2017–6471-1 AEI/FEDER, UE), Fundación CajaCanarias and Fundación Bancaria “La Caixa” (2018PATRI20), and “Unidad María de Maeztu” (CEX2018-000792-M) funded by the MCIN and the AEI (DOI:10.13039/501100011033). J.M.S was supported with a Formació de Personal Investigador fellowship from Generalitat de Catalunya (FI_B100135).

#### Authors’ contributions

J.M.S., O.L., and D.C. conceived the work. J.M.S. and O.L. performed the computational analyses, generated the figures, and wrote the manuscript. D.C. contributed to analysis and/or interpretation of results. All authors approved the final manuscript.

## Acknowledgements

We would like to thank the Scientific Computing Core Facility at the UPF (https://www.upf.edu/web/sct-sit) for their technical help and support.

