## Supplementary File 3 for "Modelling the demographic history of human North African genomes points to soft split divergence between populations"

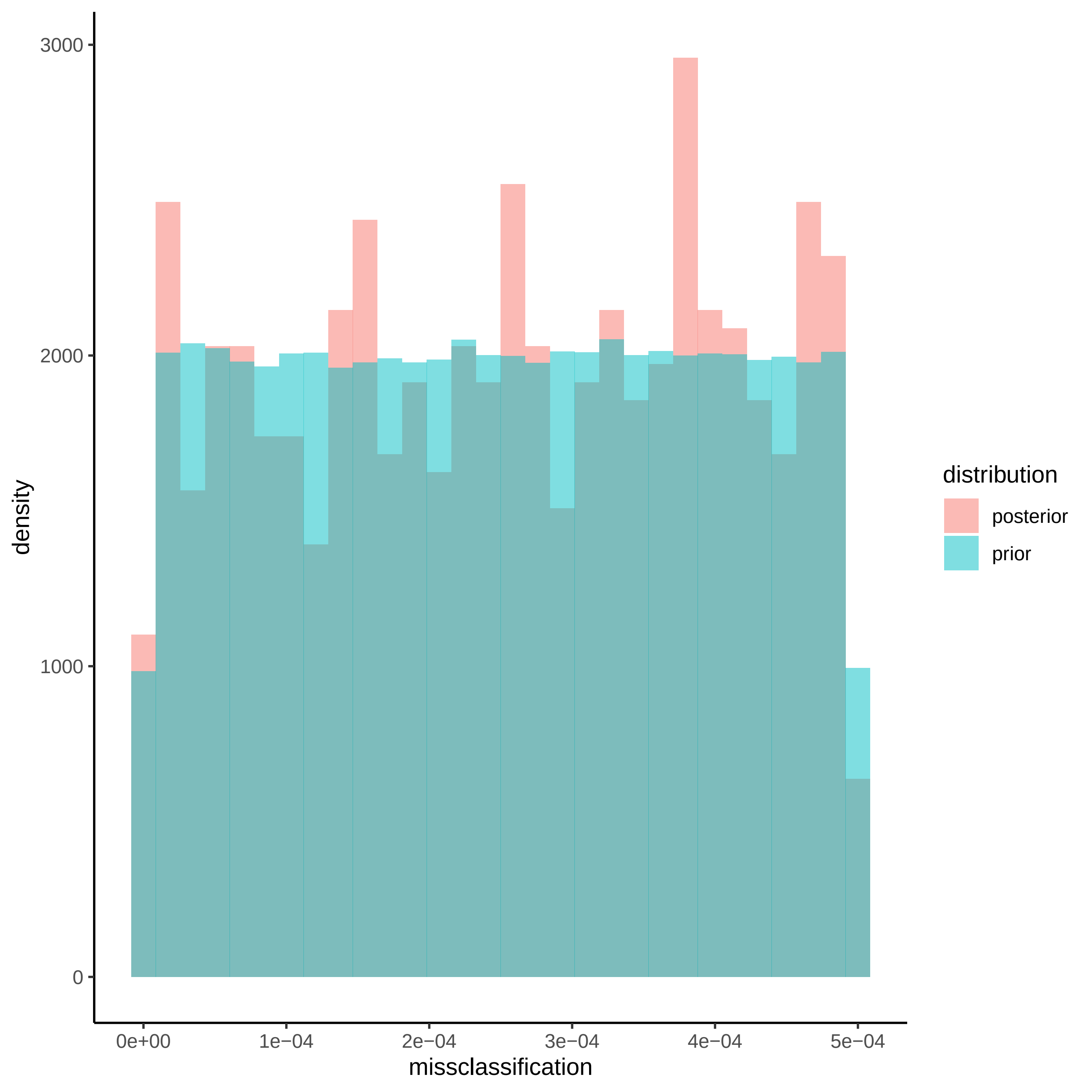

3000
2000
1000
0
density
0e+00
1e−04
2e−04
3e−04
4e−04
5e−04
missclassification
distribution
posterior
prior

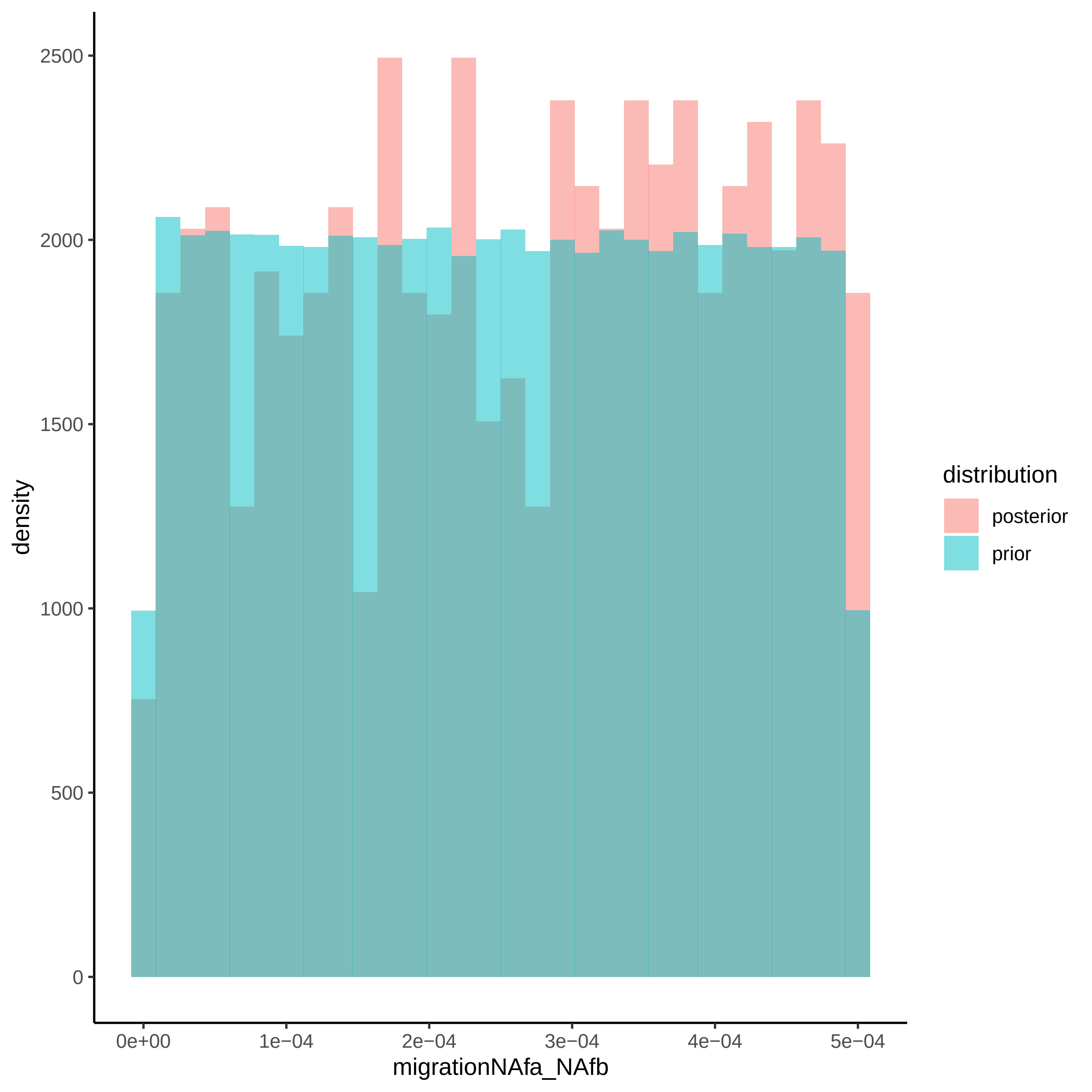

2500
2000
1500
1000
500
0
density
0e+00
1e−04
2e−04
3e−04
4e−04
5e−04
migrationNAfa_NAfb
distribution
posterior
prior

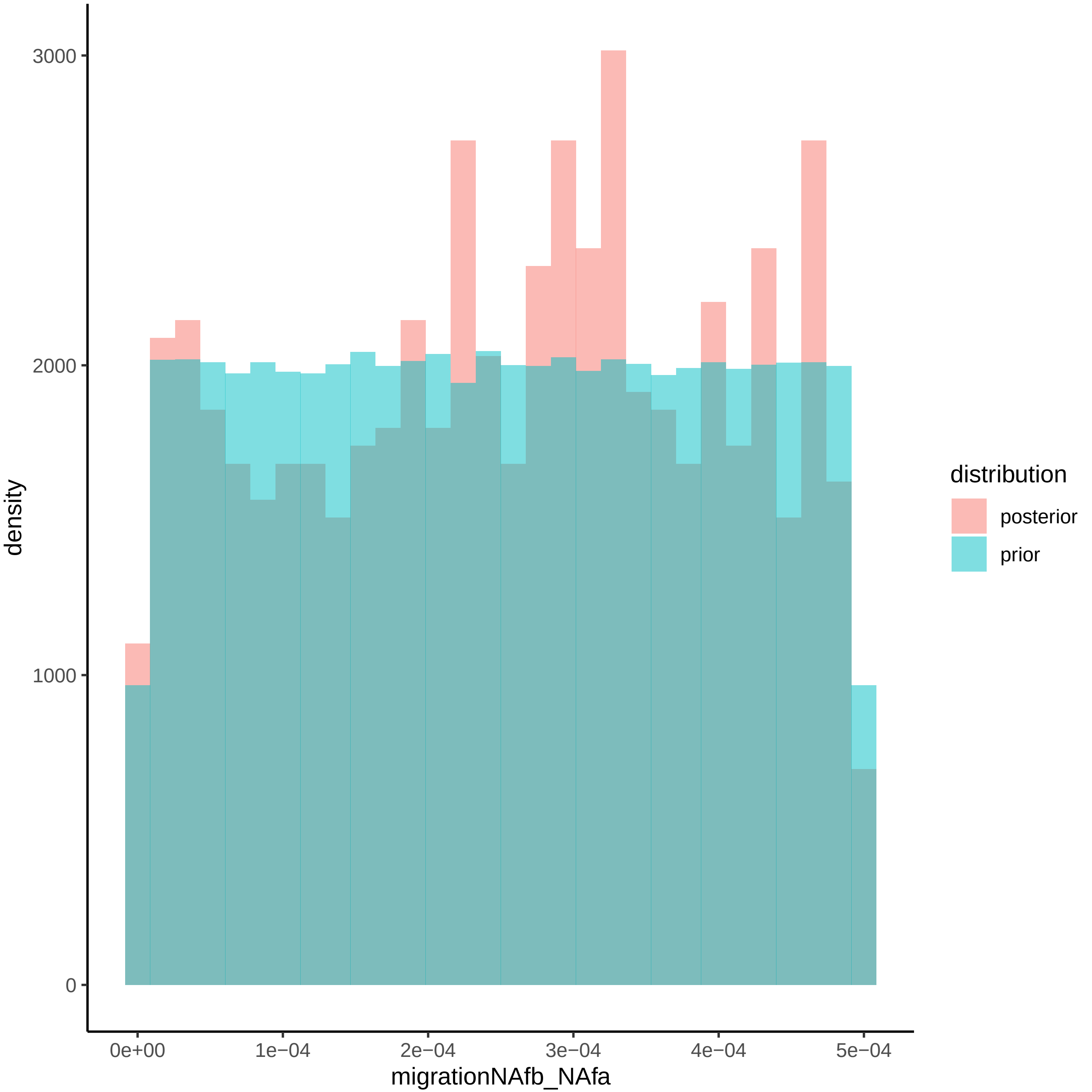

density
3000
2000
1000
0
0e+00
1e−04
2e−04
3e−04
4e−04
5e−04
migrationNAfb_NAfa
distribution
posterior
prior

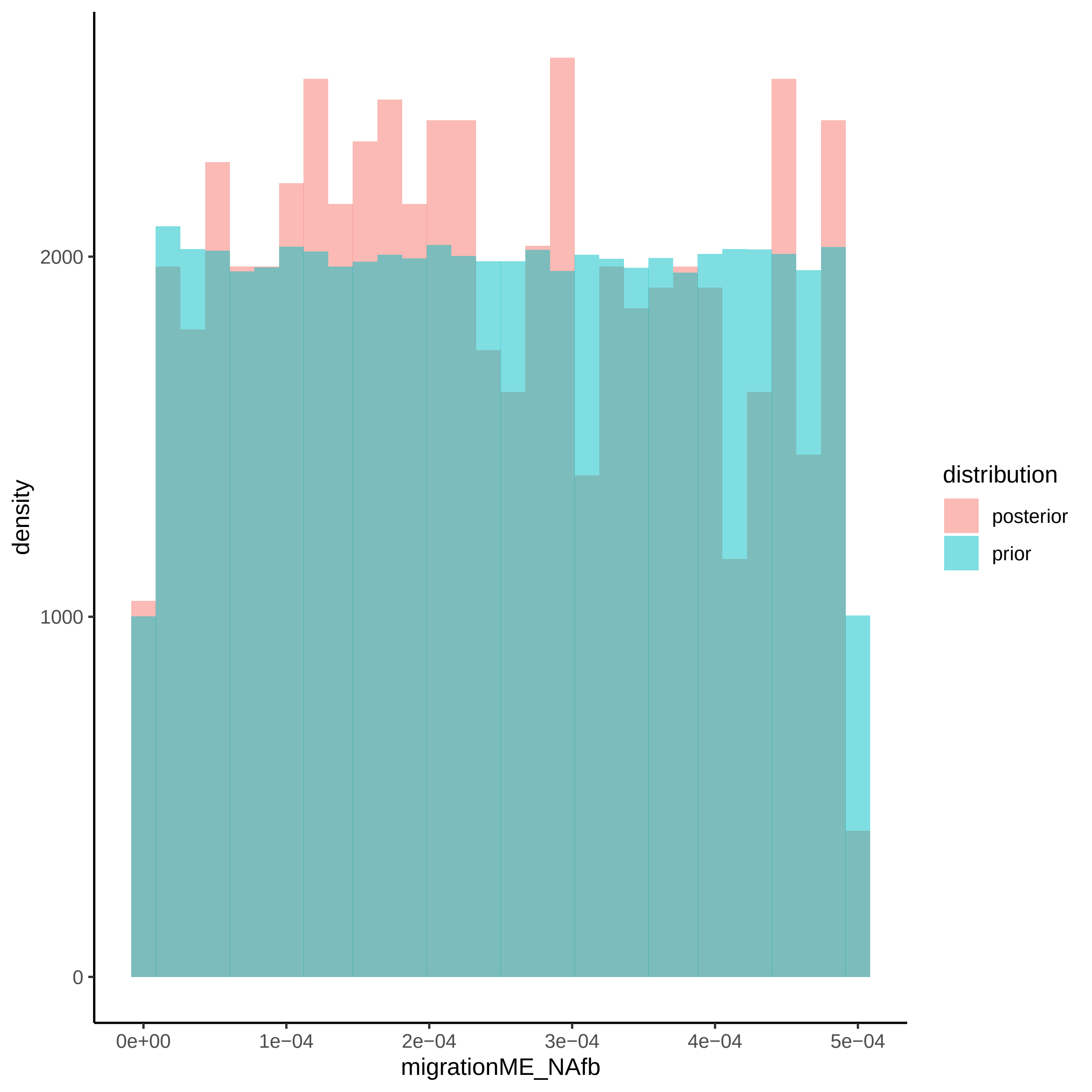

density
2000
1000
0
0e+00
1e−04
2e−04
3e−04
4e−04
5e−04
migrationME_NAfb
distribution
posterior
prior

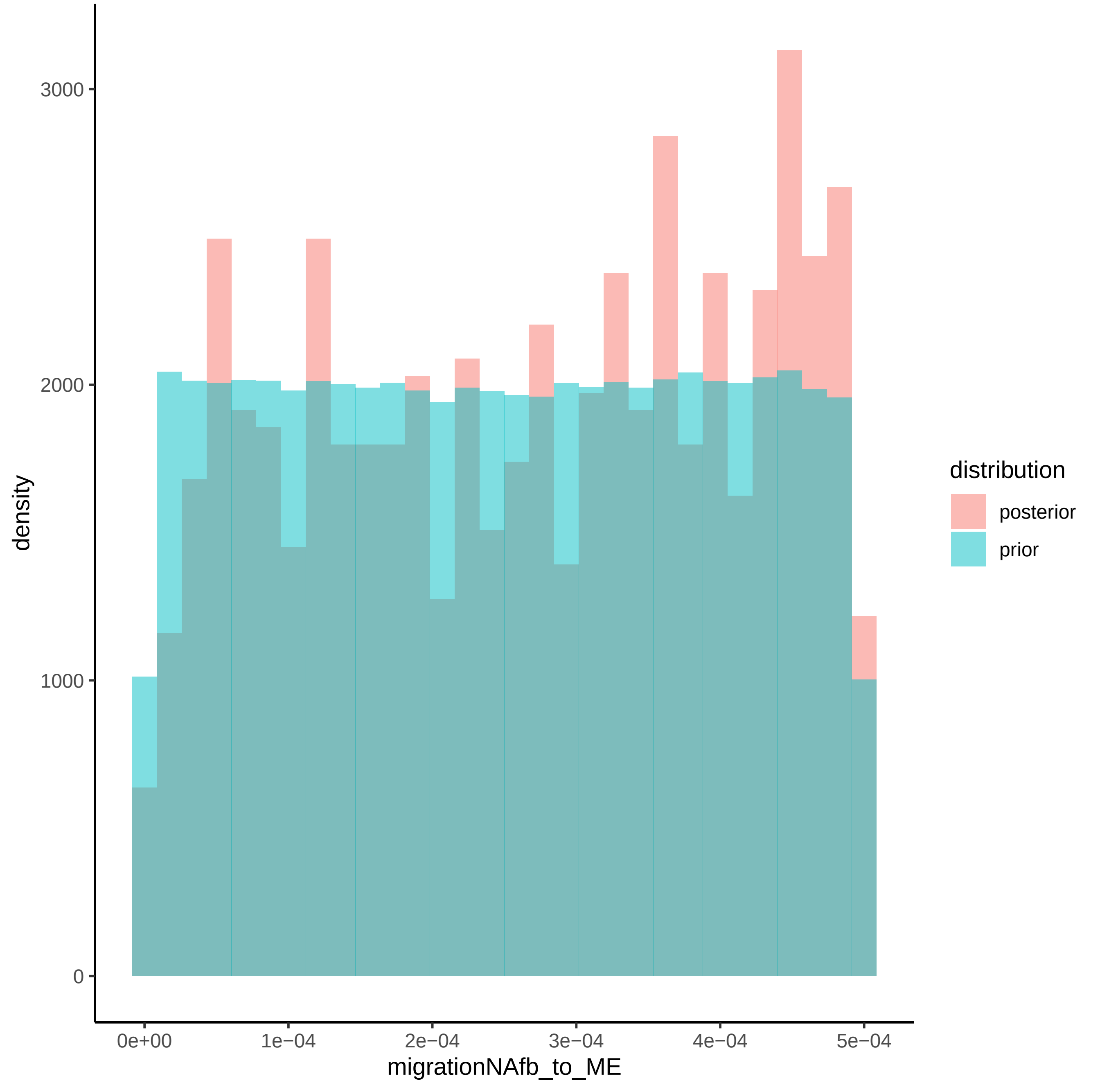

density
0
1000
2000
3000
0e+00
1e−04
2e−04
3e−04
4e−04
5e−04
migrationNAfb_to_ME
distribution
posterior
prior

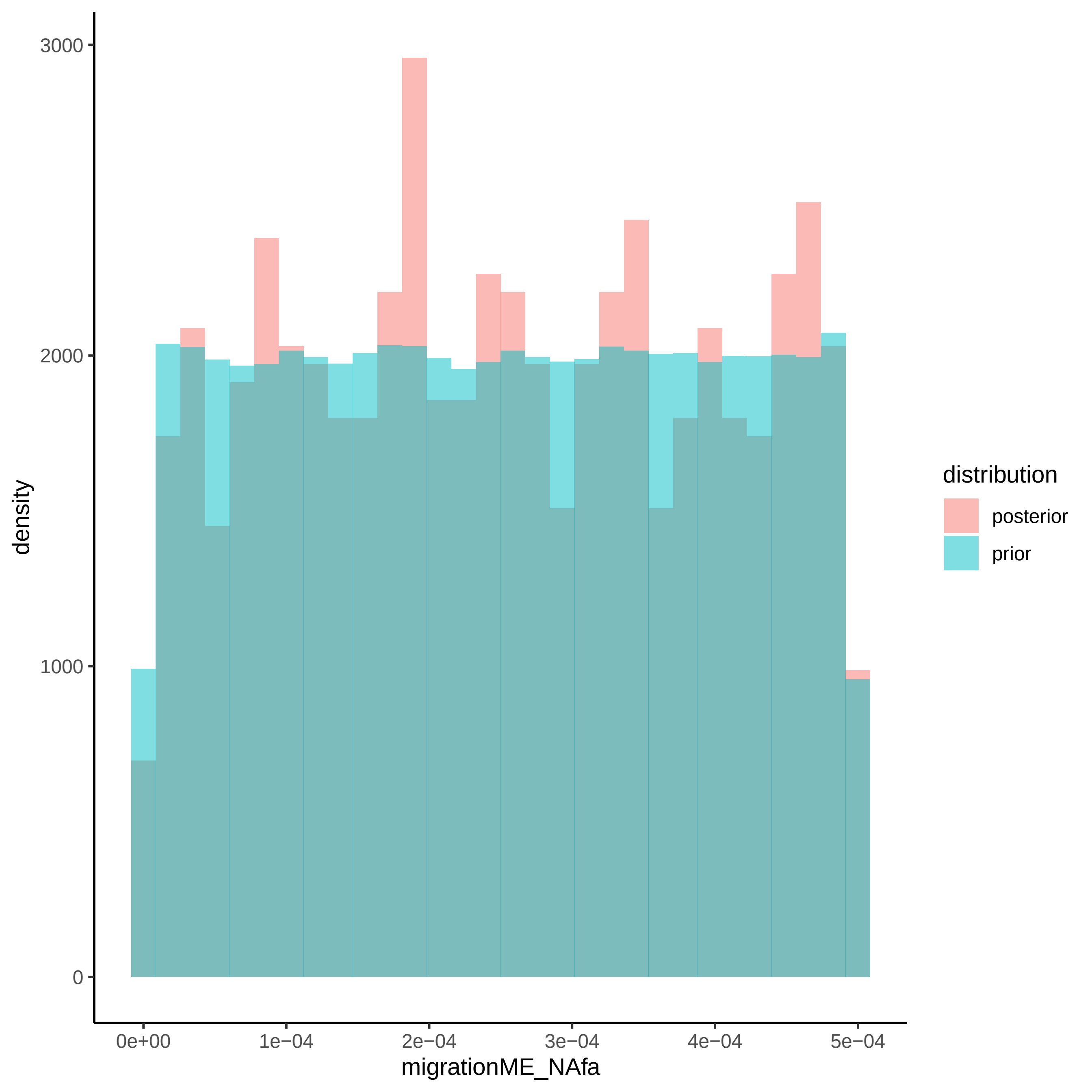

3000
2000
1000
0
density
0e+00
1e−04
2e−04
3e−04
4e−04
5e−04
migrationME_NAfa
distribution
posterior
prior

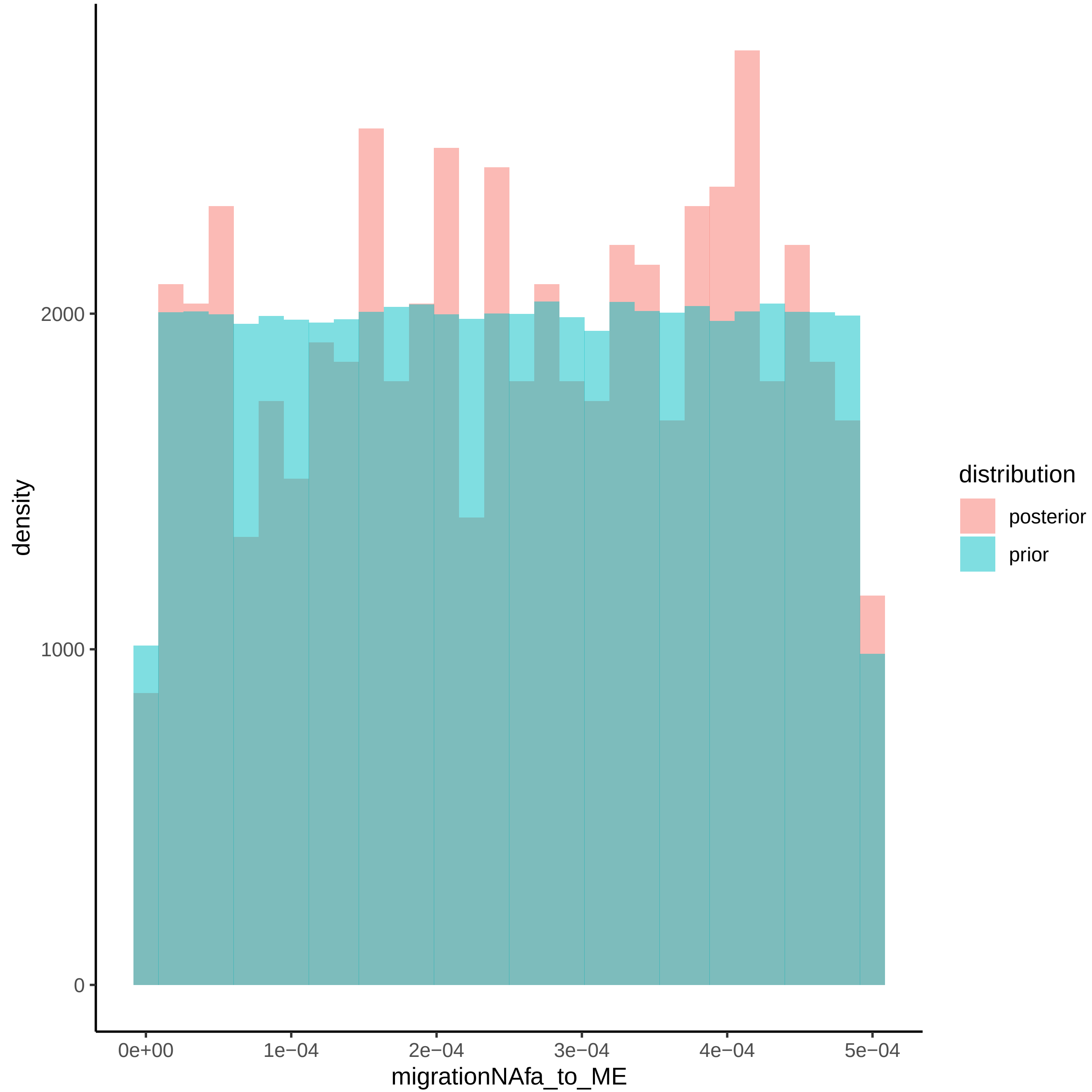

density
2000
1000
0
0e+00
1e−04
2e−04
3e−04
4e−04
5e−04
migrationNAfa_to_ME
distribution
posterior
prior

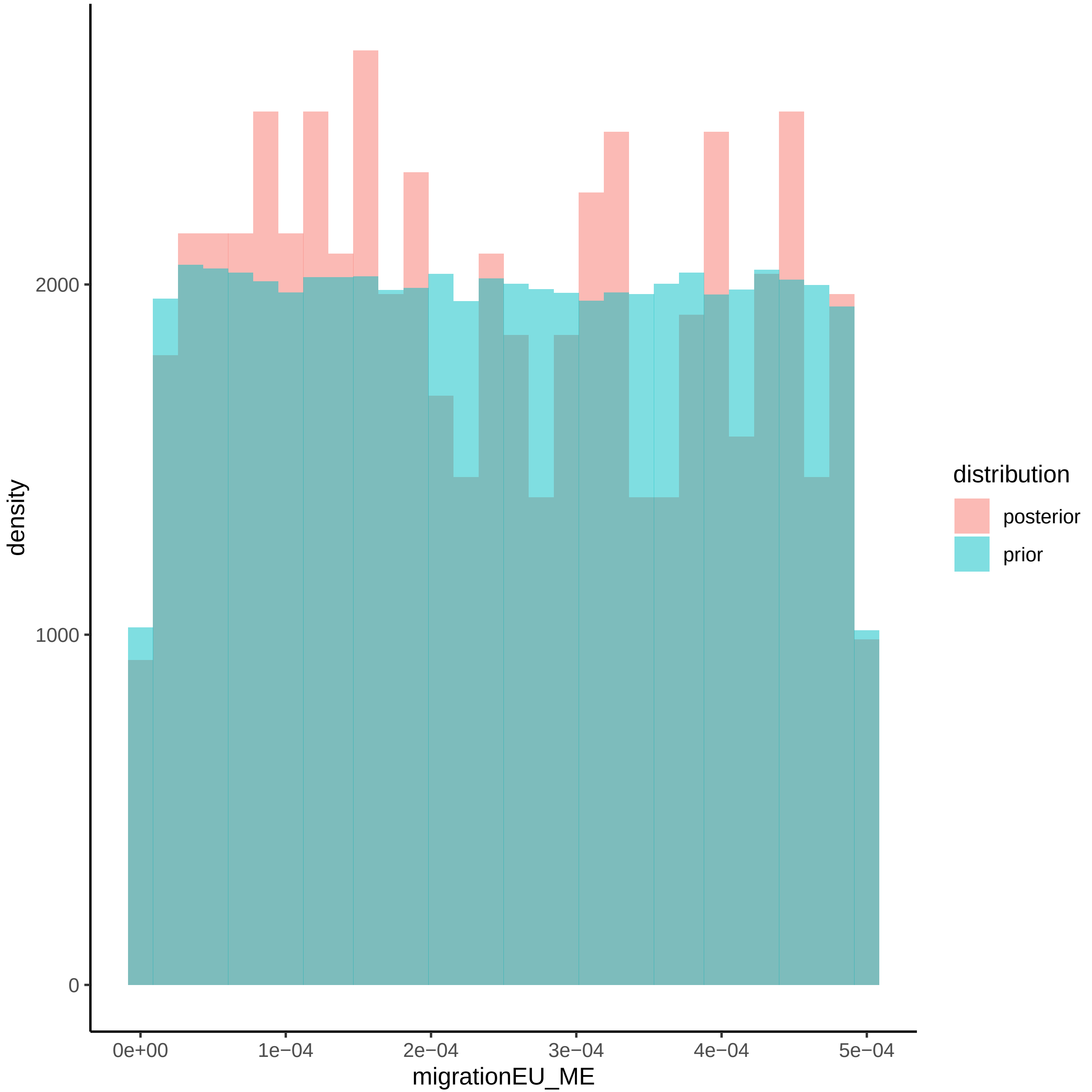

density
2000
1000
0
0e+00
1e−04
2e−04
3e−04
4e−04
5e−04
migrationEU_ME
distribution
posterior
prior

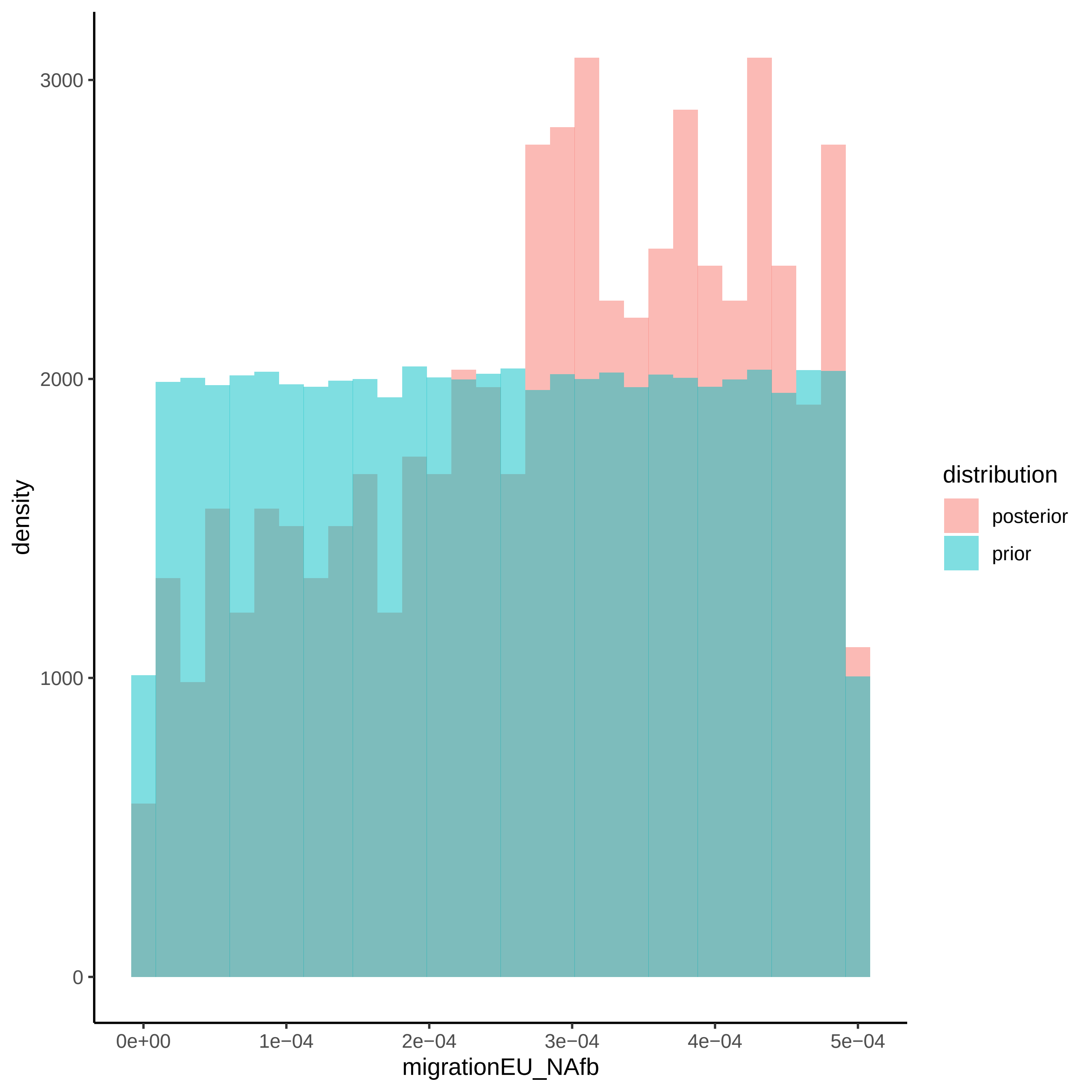
3000
2000
1000
0
density
0e+00
1e−04
2e−04
3e−04
4e−04
5e−04
migrationEU_NAfb
distribution
posterior
prior

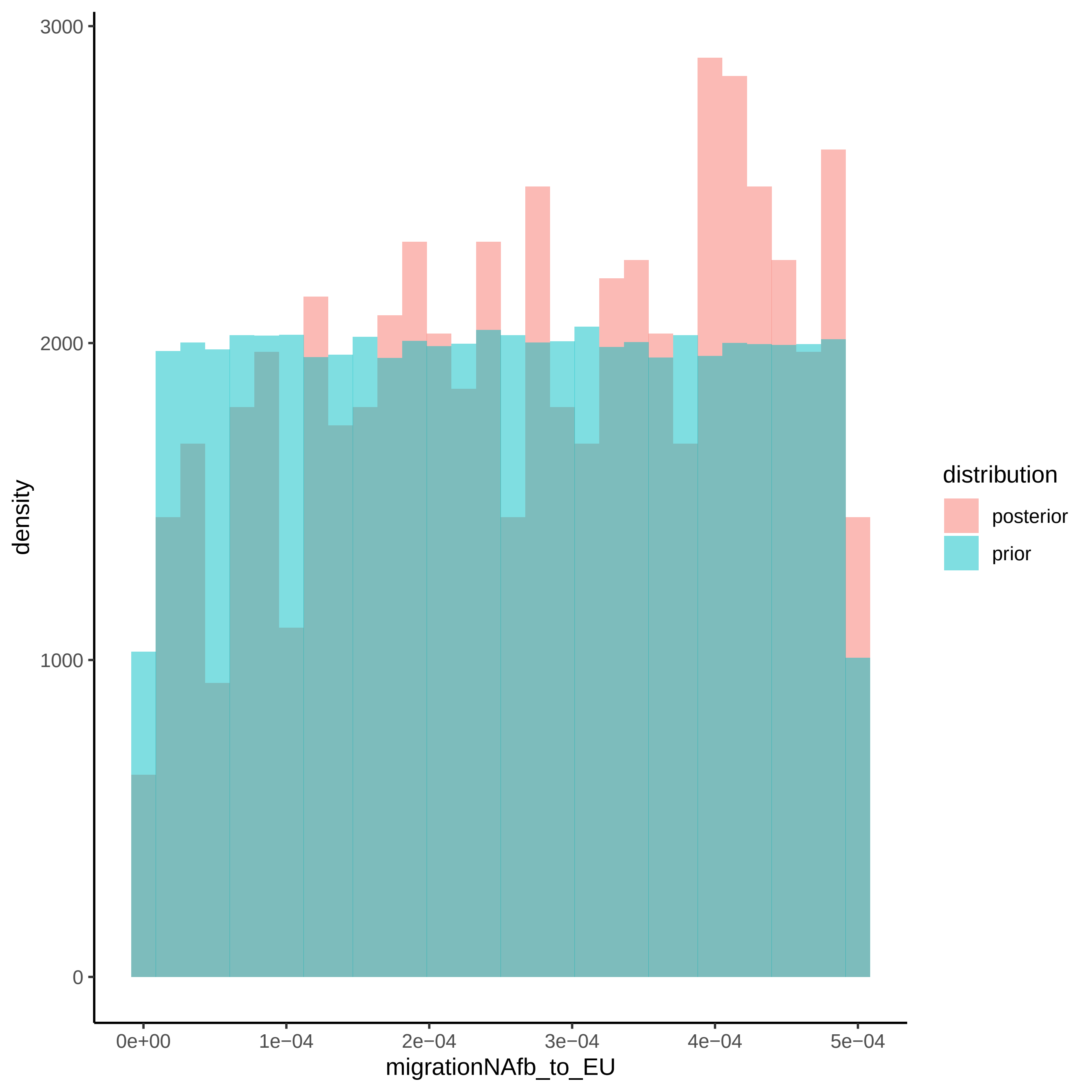

3000
2000
1000
0
density
0e+00
1e−04
2e−04
3e−04
4e−04
5e−04
migrationNAfb_to_EU
distribution
posterior
prior

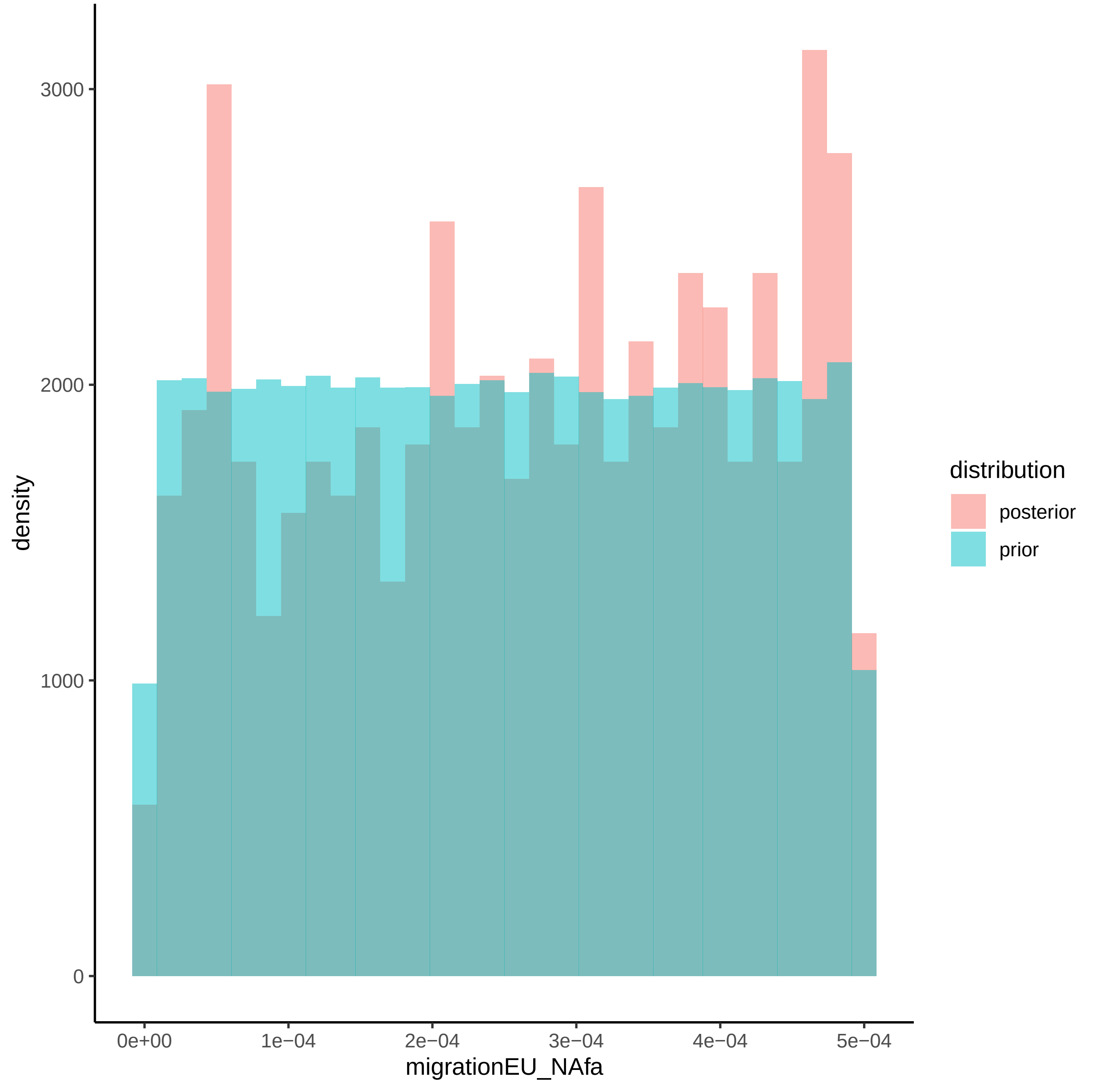

3000
2000
1000
0
density
0e+00
1e−04
2e−04
3e−04
4e−04
5e−04
migrationEU_NAfa
distribution
posterior
prior

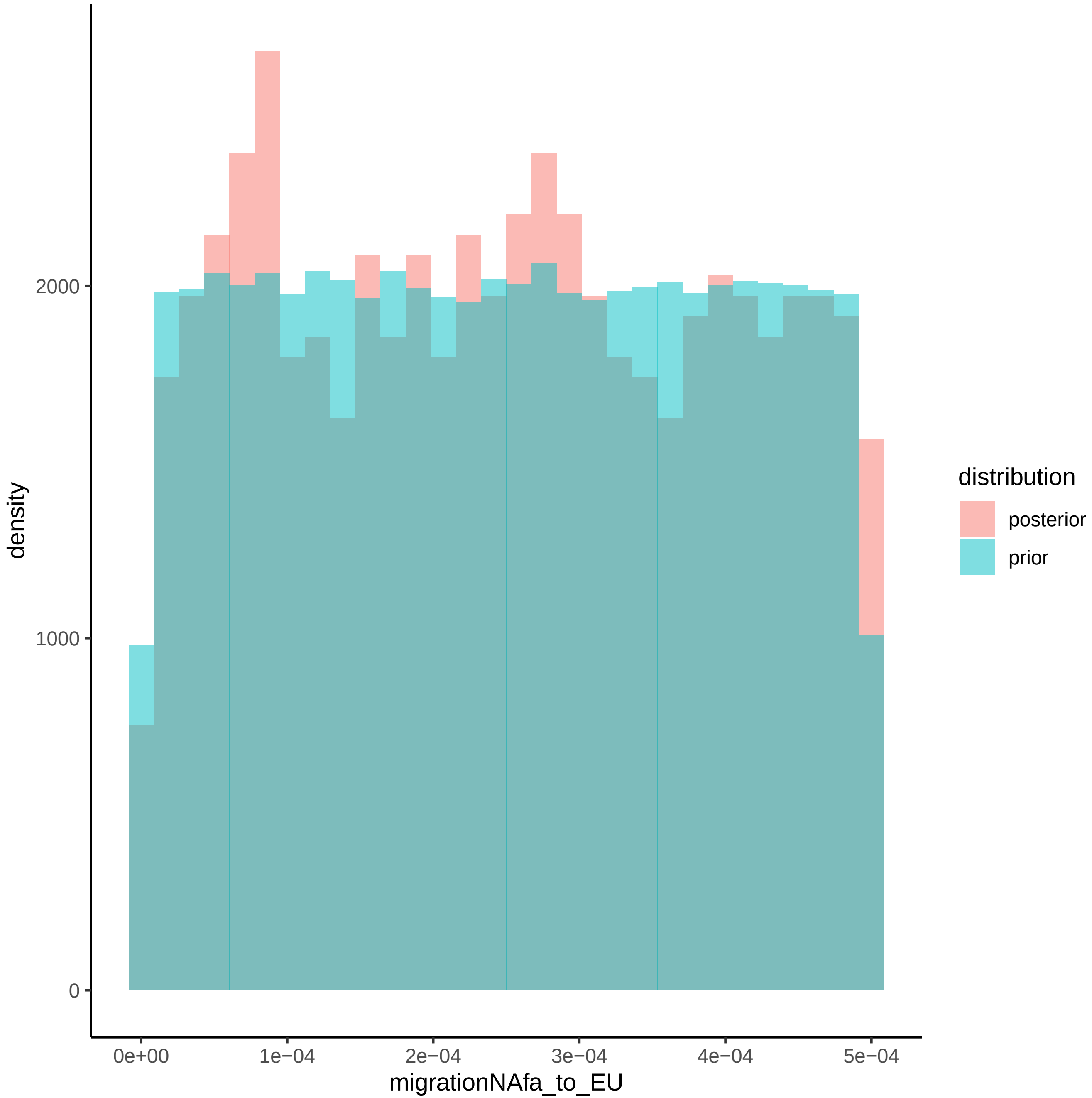
density
2000
1000
0
0e+00
1e−04
2e−04
3e−04
4e−04
5e−04
migrationNAfa_to_EU
distribution
posterior
prior

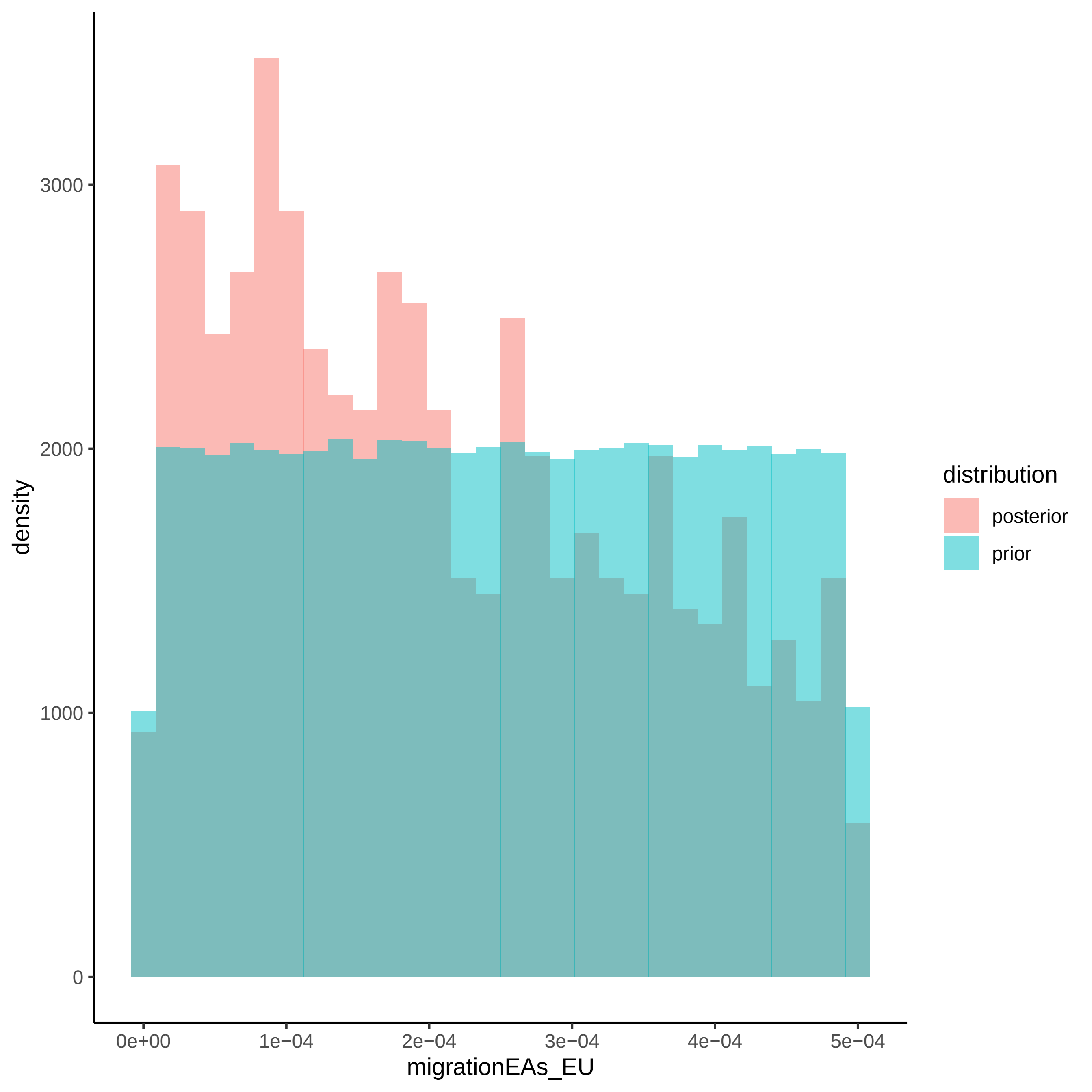
density
3000
2000
1000
0
0e+00
1e−04
2e−04
3e−04
4e−04
5e−04
migrationEAs_EU
distribution
posterior
prior

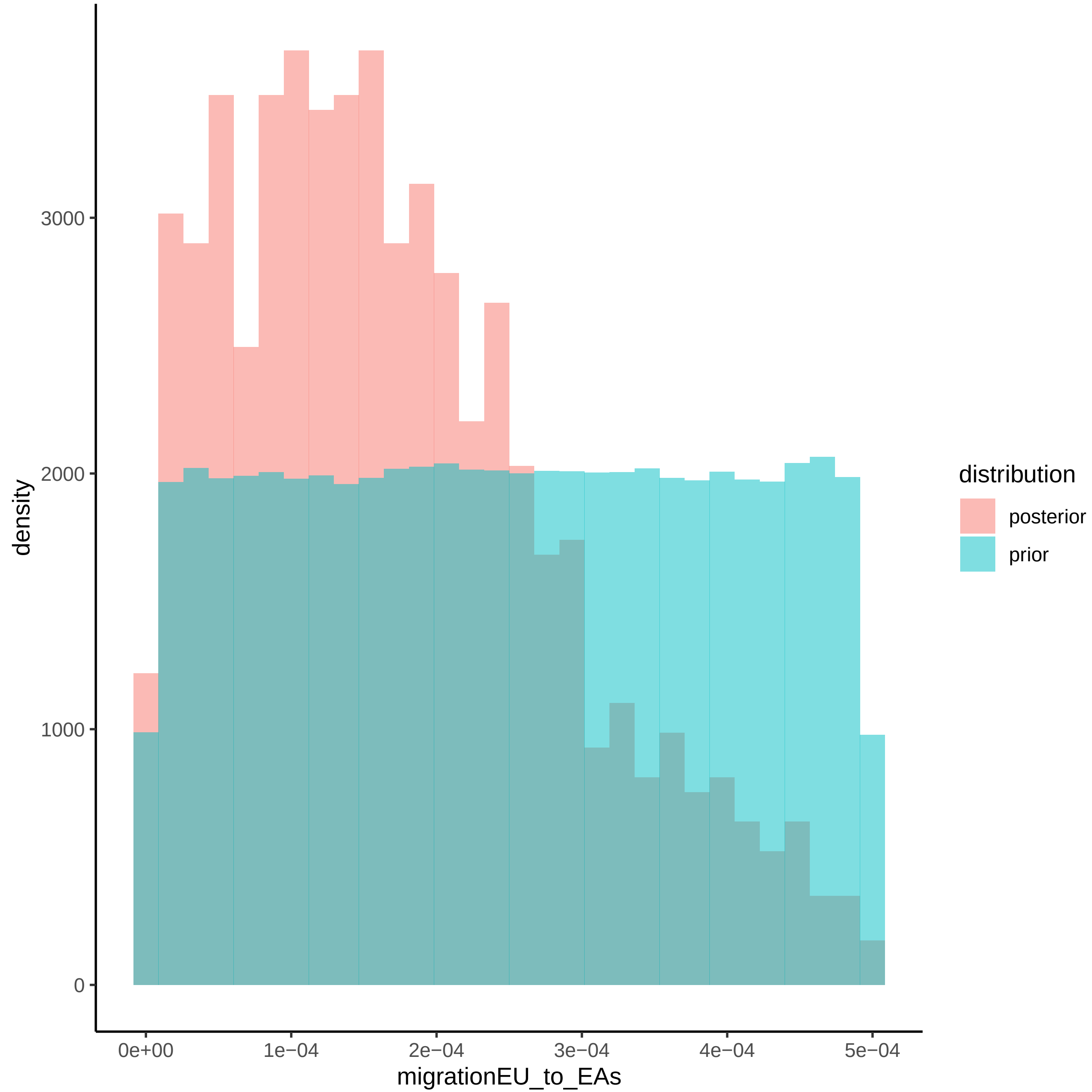
density
3000
2000
1000
0
0e+00
1e−04
2e−04
3e−04
4e−04
5e−04
migrationEU_to_EAs
distribution
posterior
prior

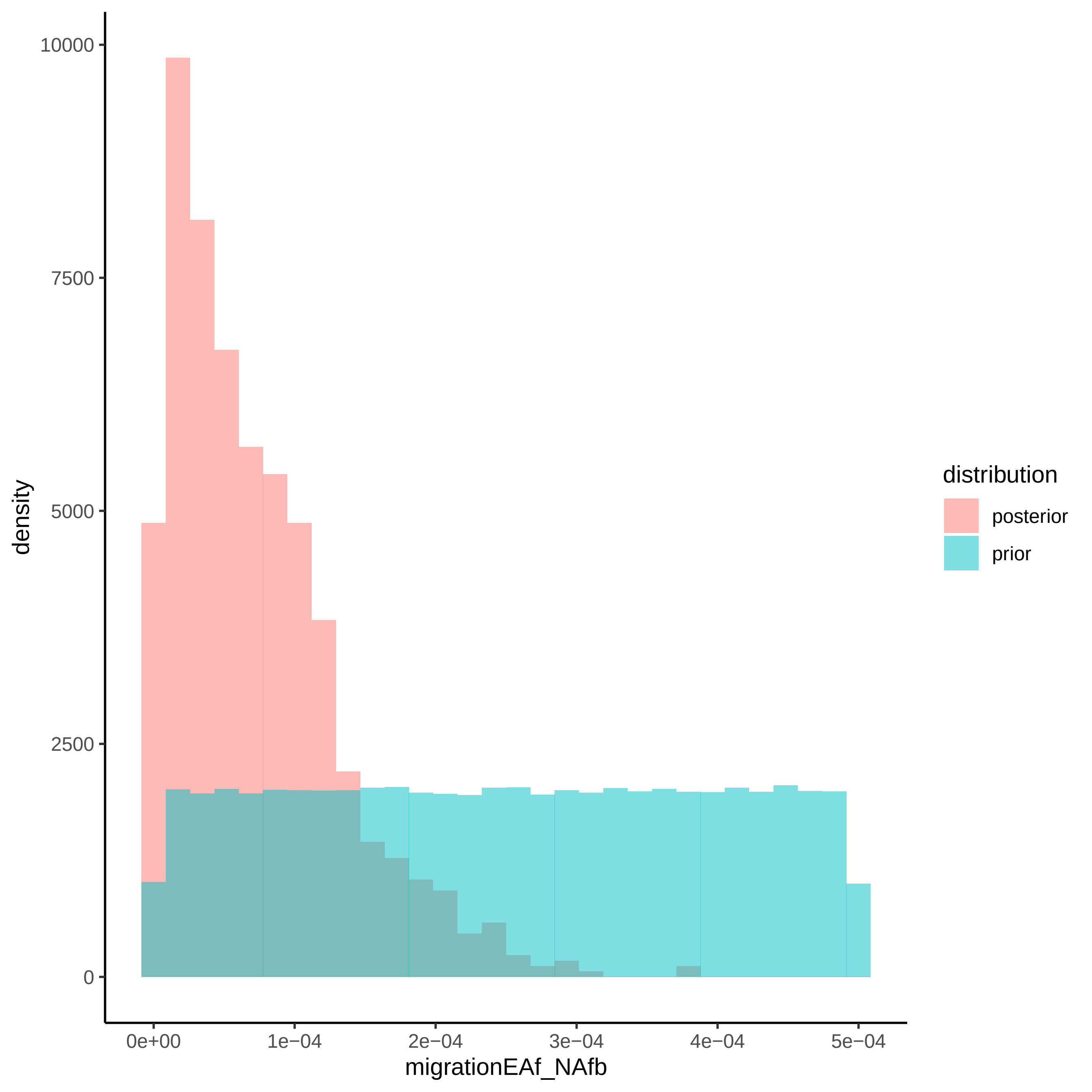

density
10000
7500
5000
2500
0
0e+00
1e−04
2e−04
3e−04
4e−04
5e−04
migrationEAf_NAfb
distribution
posterior
prior

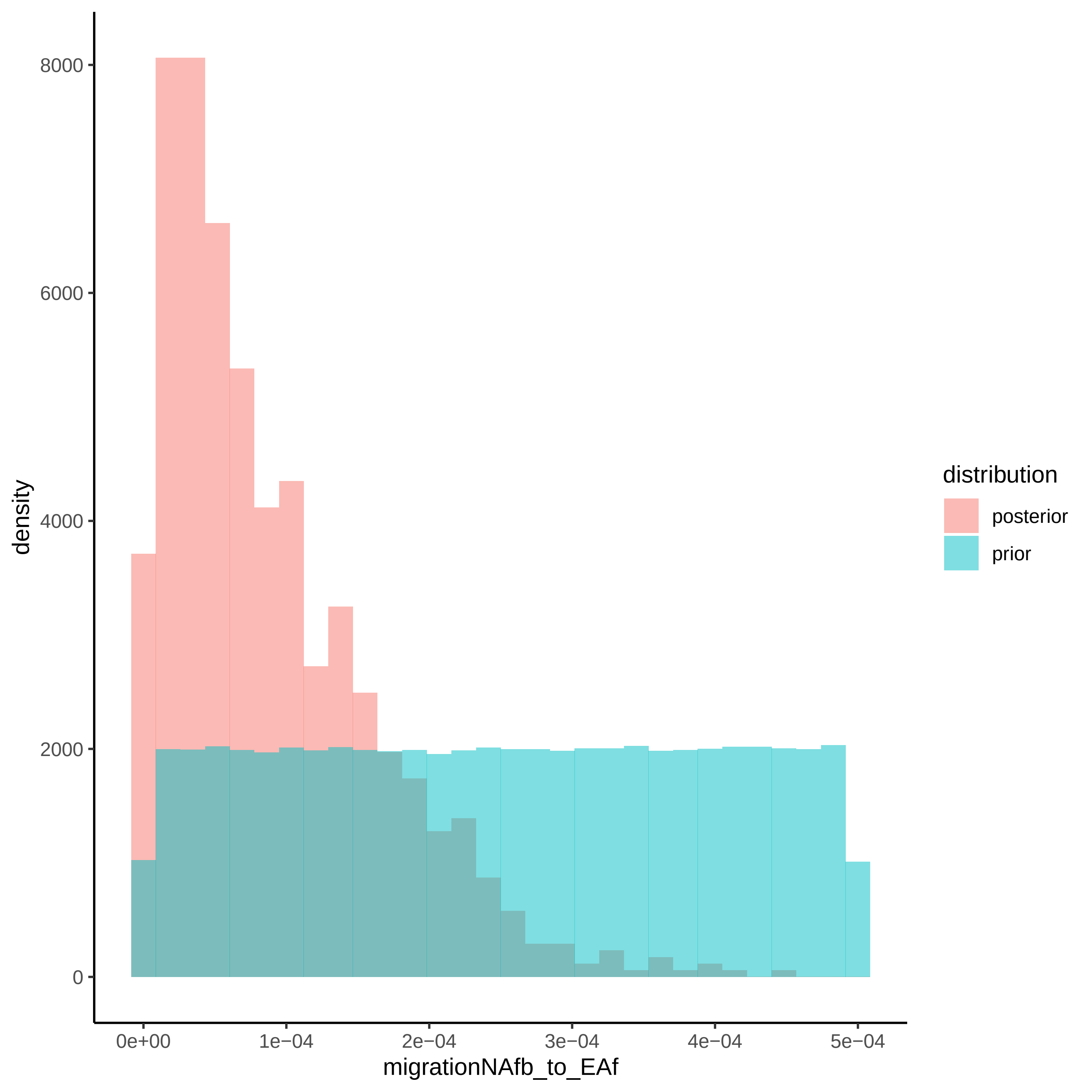

density
8000
6000
4000
2000
0
0e+00
1e−04
2e−04
3e−04
4e−04
5e−04
migrationNAfb_to_EAf
distribution
posterior
prior

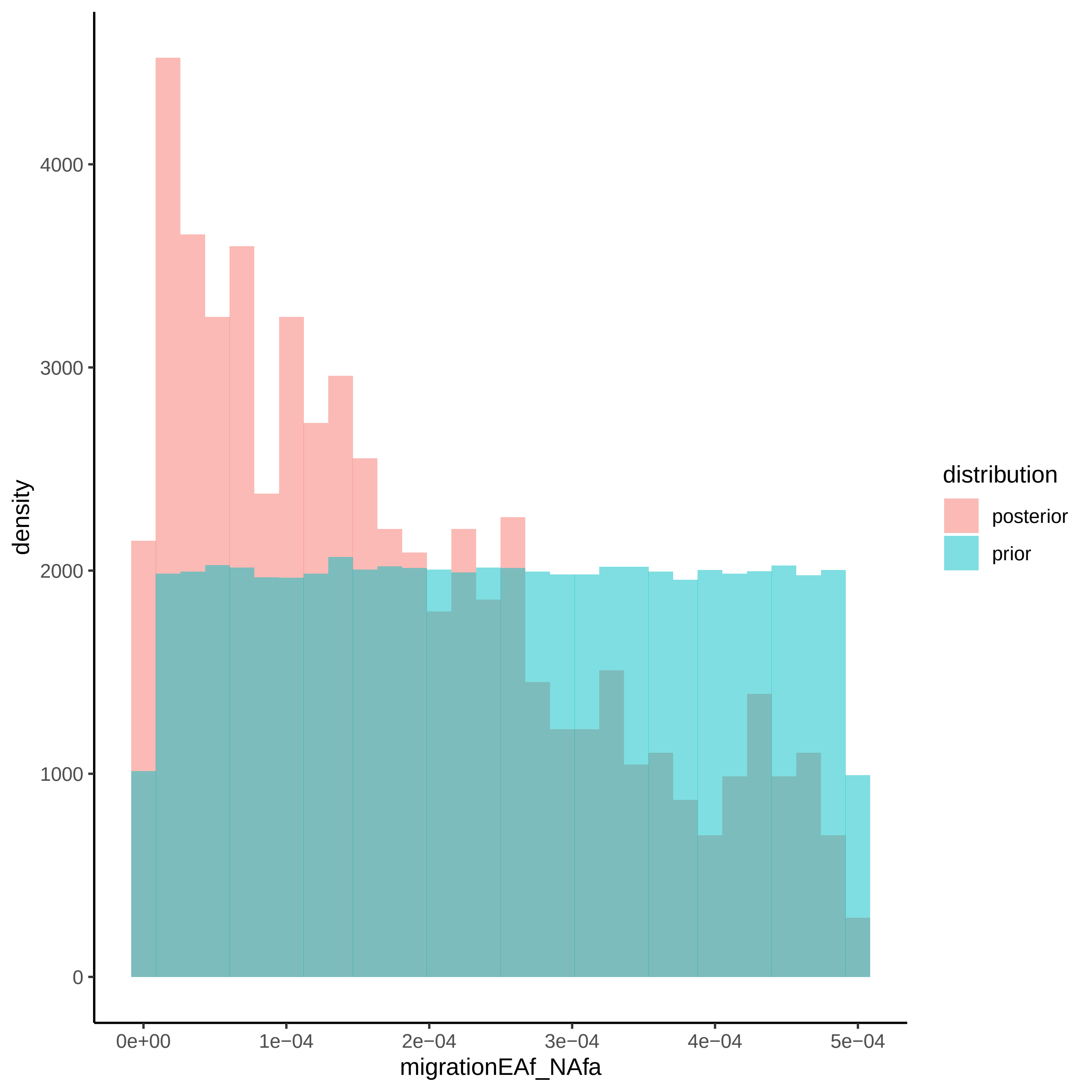

density
4000
3000
2000
1000
0
0e+00
1e−04
2e−04
3e−04
4e−04
5e−04
migrationEAf_NAfa
distribution
posterior
prior

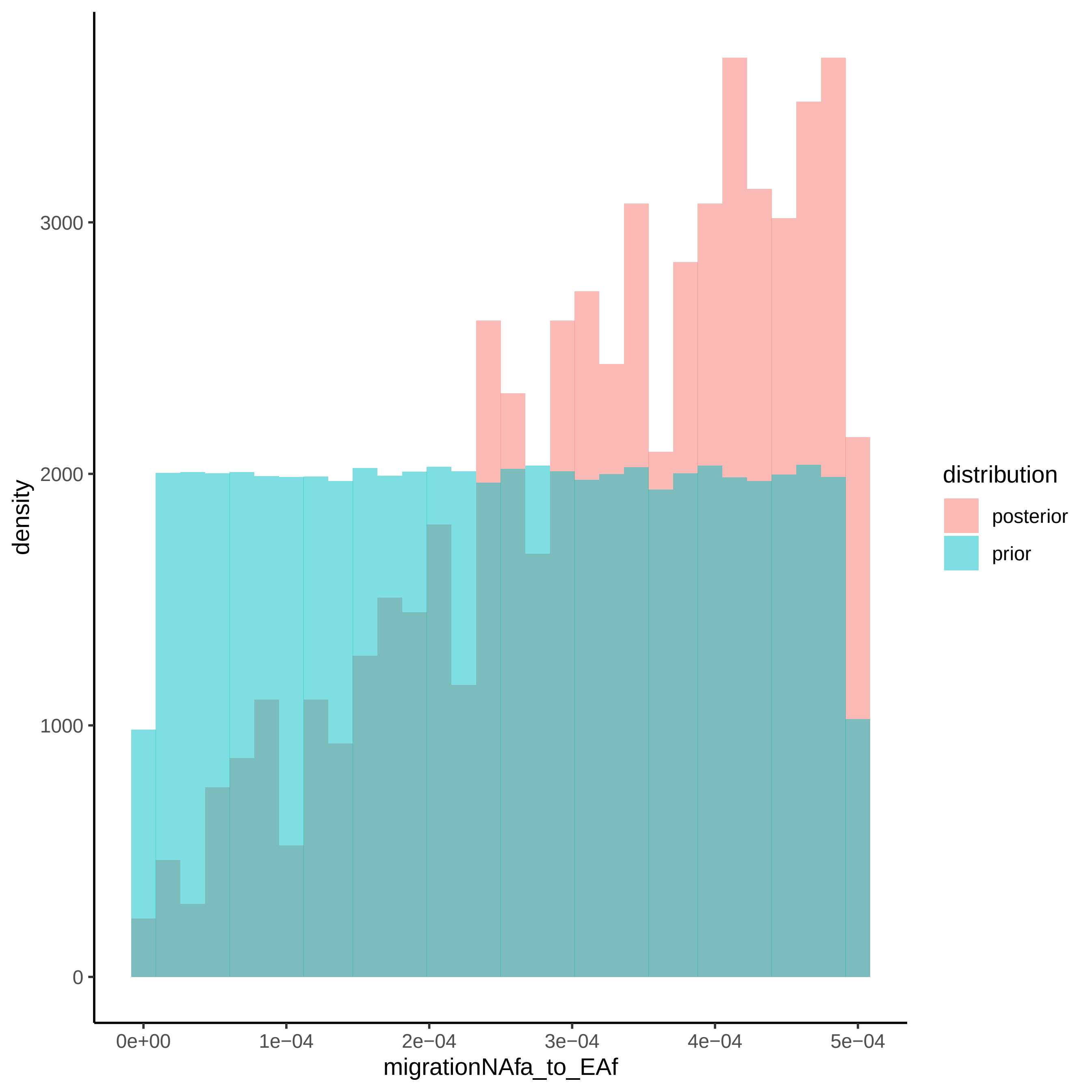

density
3000
2000
1000
0
0e+00
1e−04
2e−04
3e−04
4e−04
5e−04
migrationNAfa_to_EAf
distribution
posterior
prior

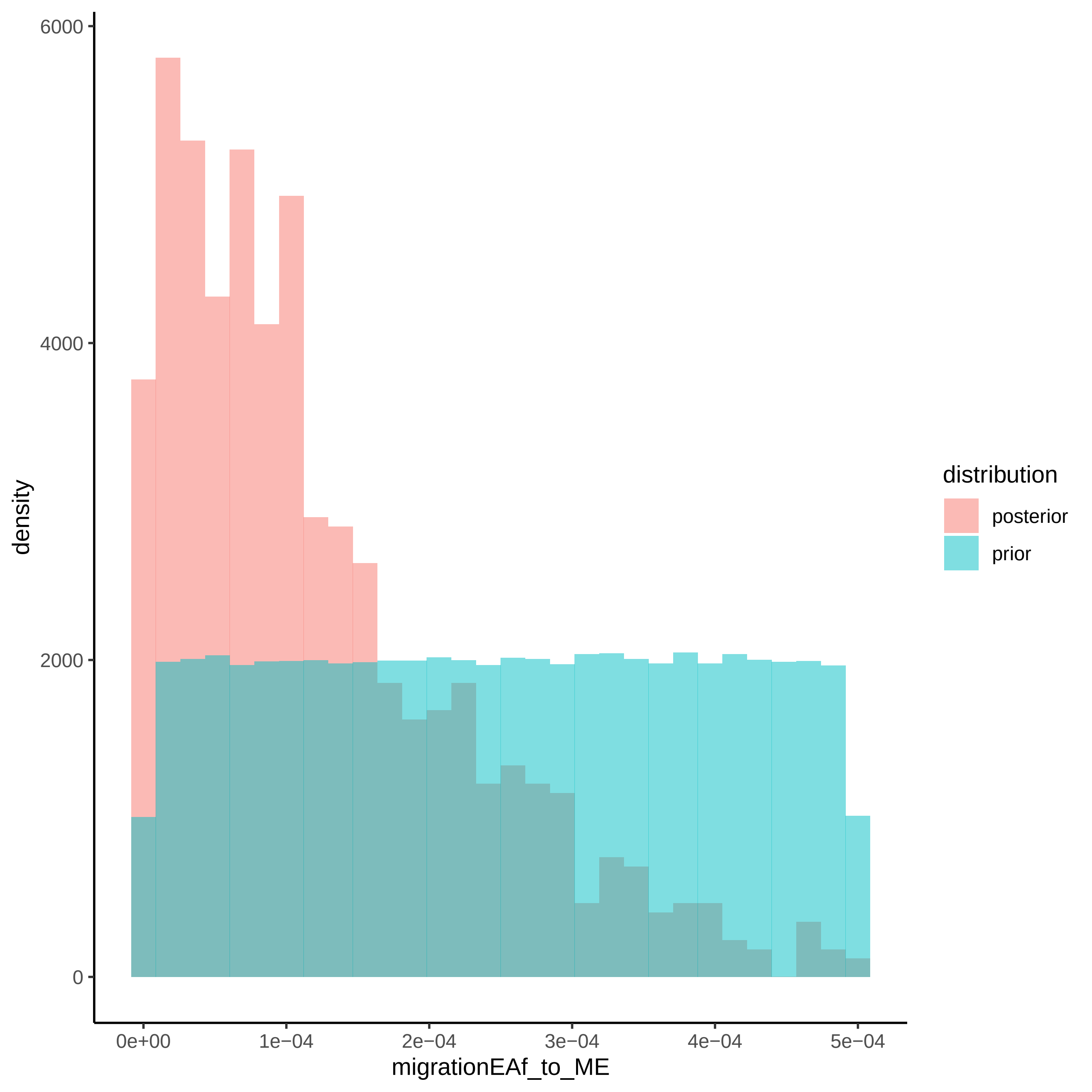
6000
4000
2000
0
density
0e+00
1e−04
2e−04
3e−04
4e−04
5e−04
migrationEAf_to_ME
distribution
posterior
prior

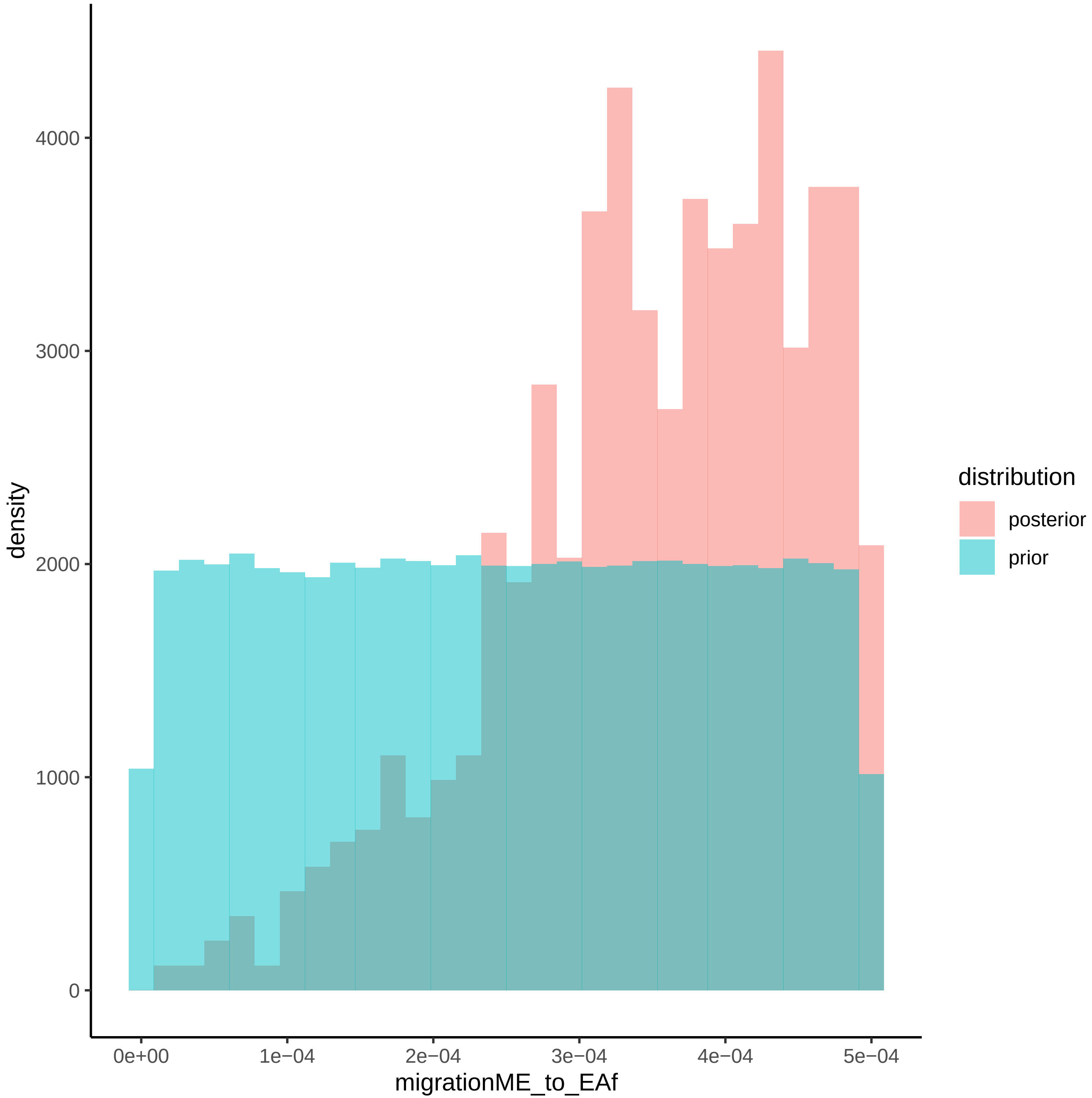

density
4000
3000
2000
1000
0
0e+00
1e−04
2e−04
3e−04
4e−04
5e−04
migrationME_to_EAf
distribution
posterior
prior

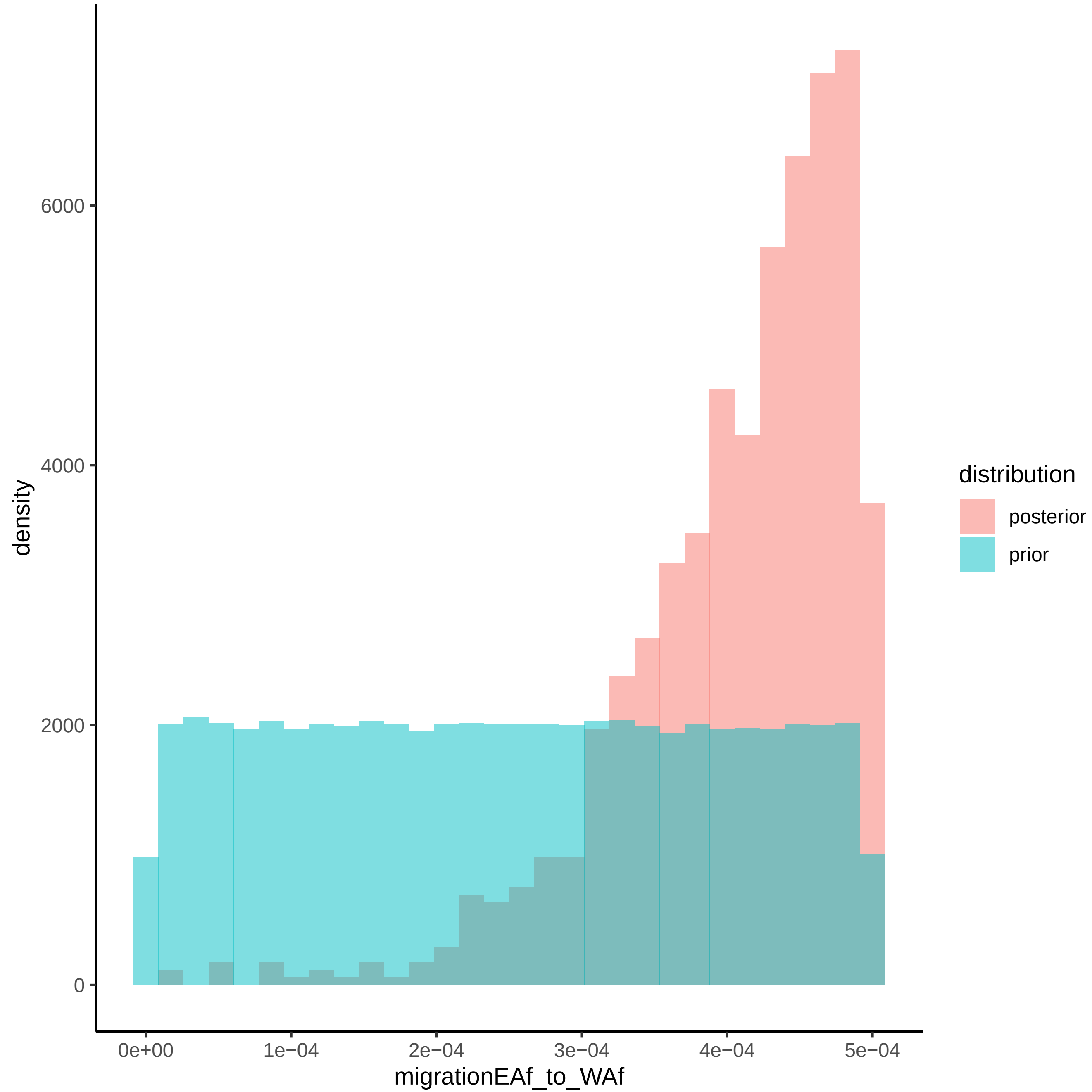

density
6000
4000
2000
0
0e+00
1e−04
2e−04
3e−04
4e−04
5e−04
migrationEAf_to_WAf
distribution
posterior
prior

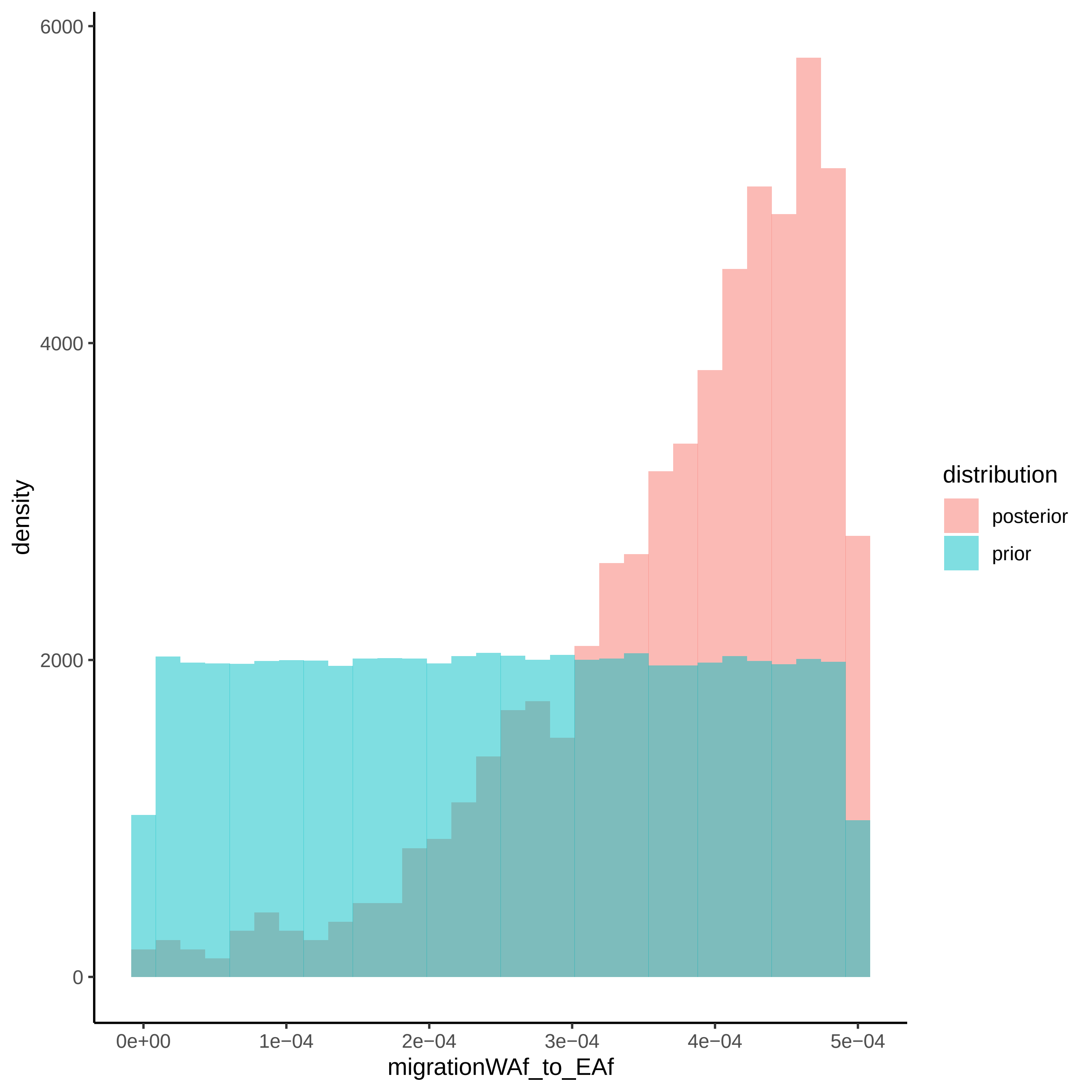

density
6000
4000
2000
0
0e+00
1e−04
2e−04
3e−04
4e−04
5e−04
migrationWAf_to_EAf
distribution
posterior
prior

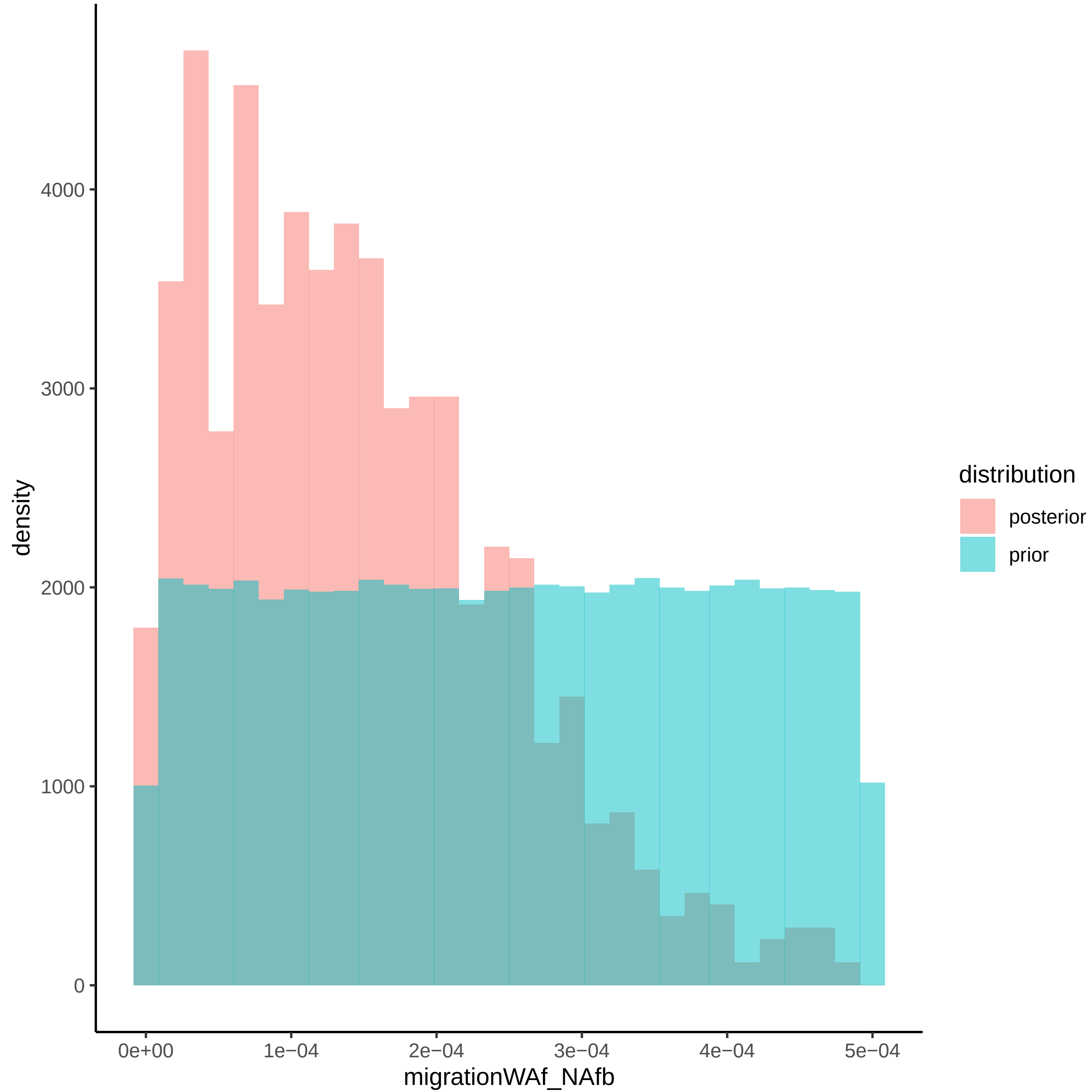

density
4000
3000
2000
1000
0
0e+00
1e−04
2e−04
3e−04
4e−04
5e−04
migrationWAf_NAfb
distribution
posterior
prior

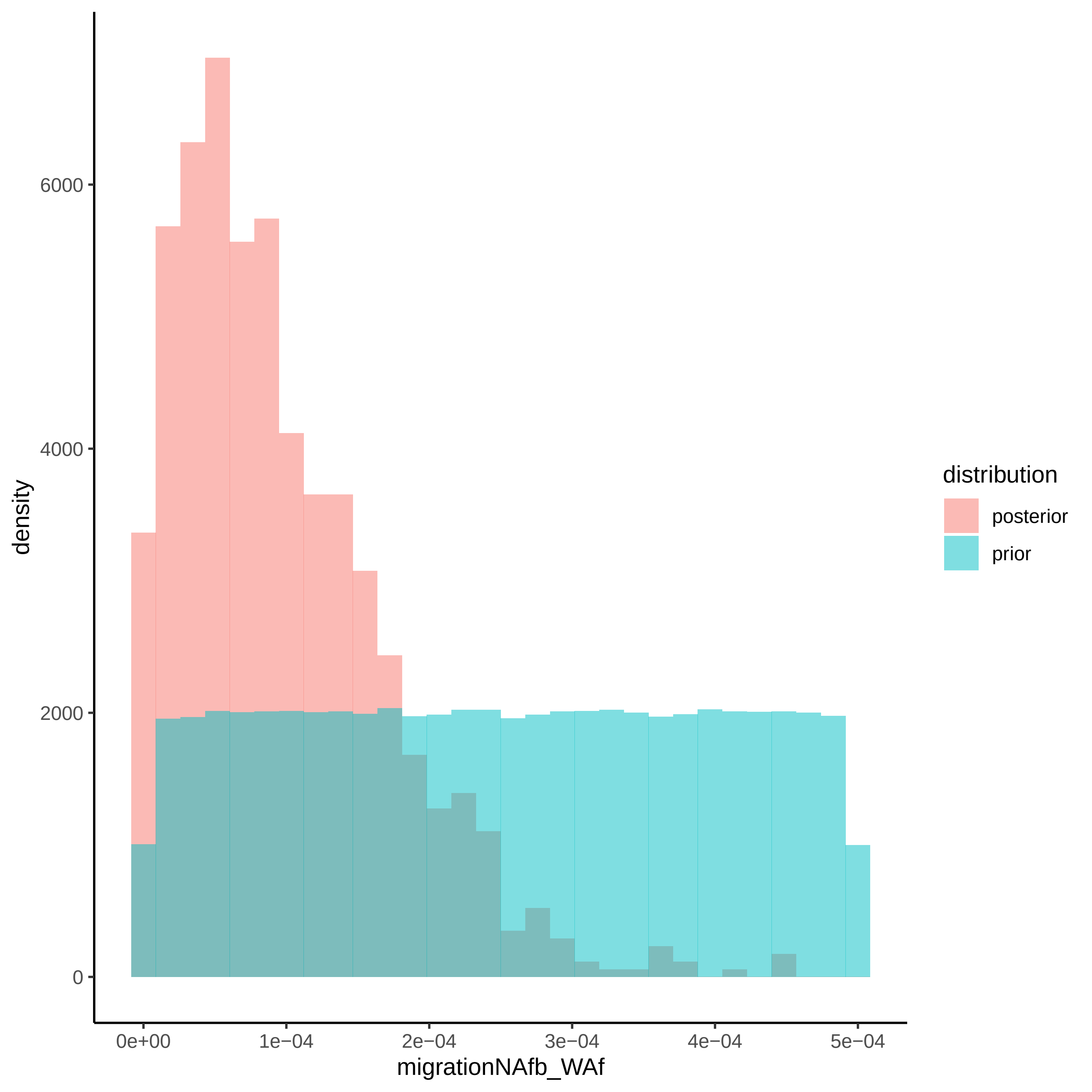

density
6000
4000
2000
0
0e+00
1e−04
2e−04
3e−04
4e−04
5e−04
migrationNAfb_WAf
distribution
posterior
prior

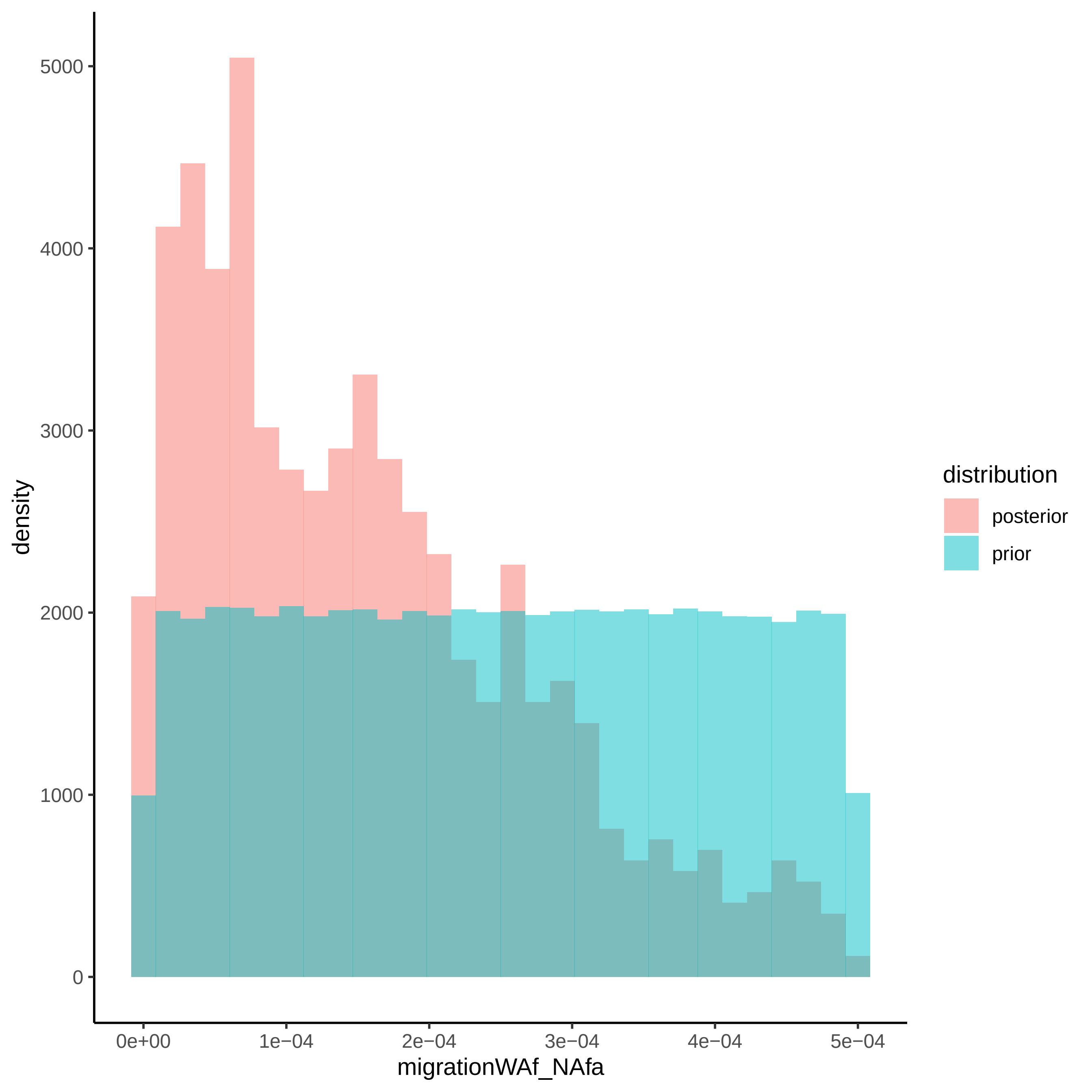
density
migrationWAf_NAfa
distribution
posterior
prior
0
1000
2000
3000
4000
5000
0e+00
1e−04
2e−04
3e−04
4e−04
5e−04

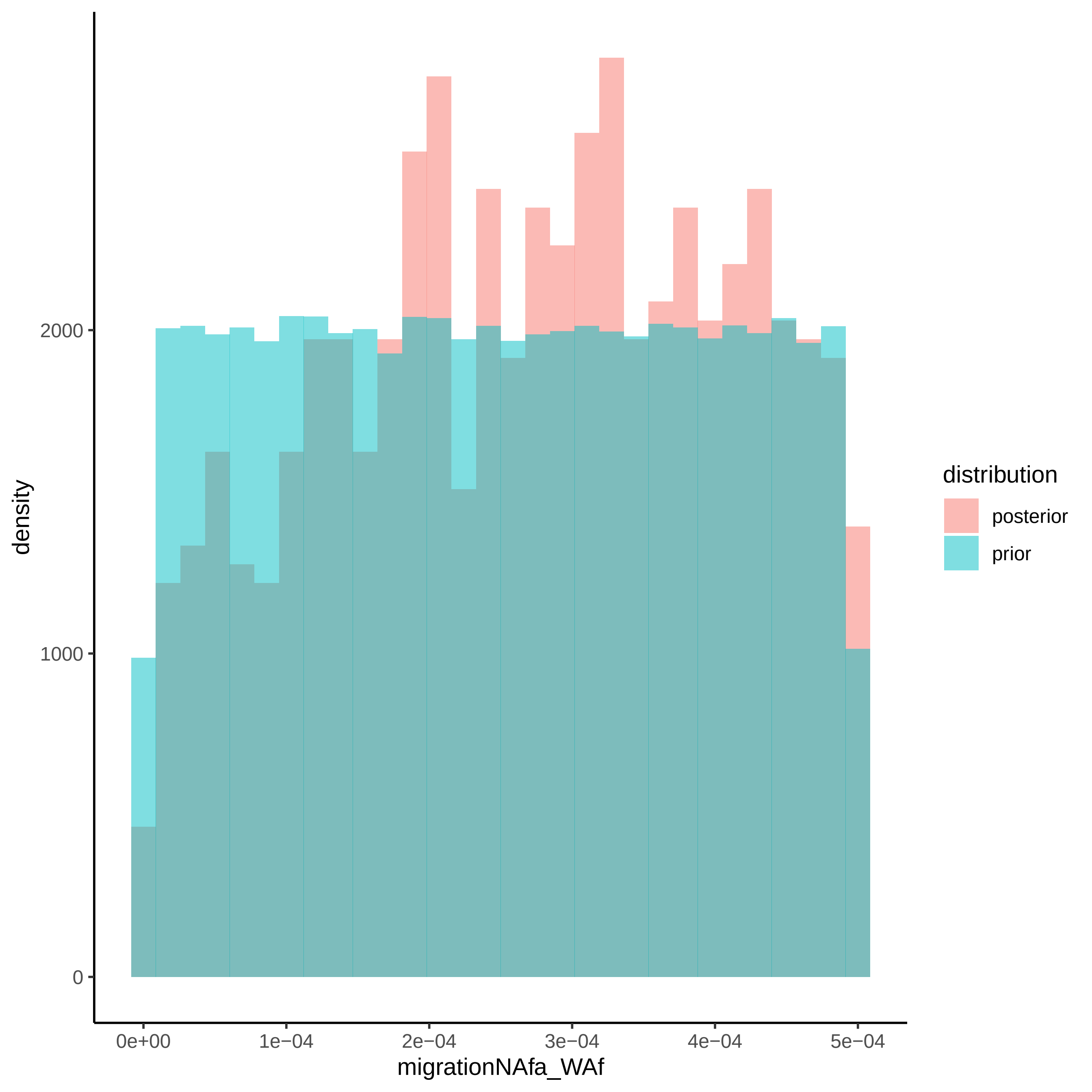

density
2000
1000
0
0e+00
1e−04
2e−04
3e−04
4e−04
5e−04
migrationNAfa_WAf
distribution
posterior
prior

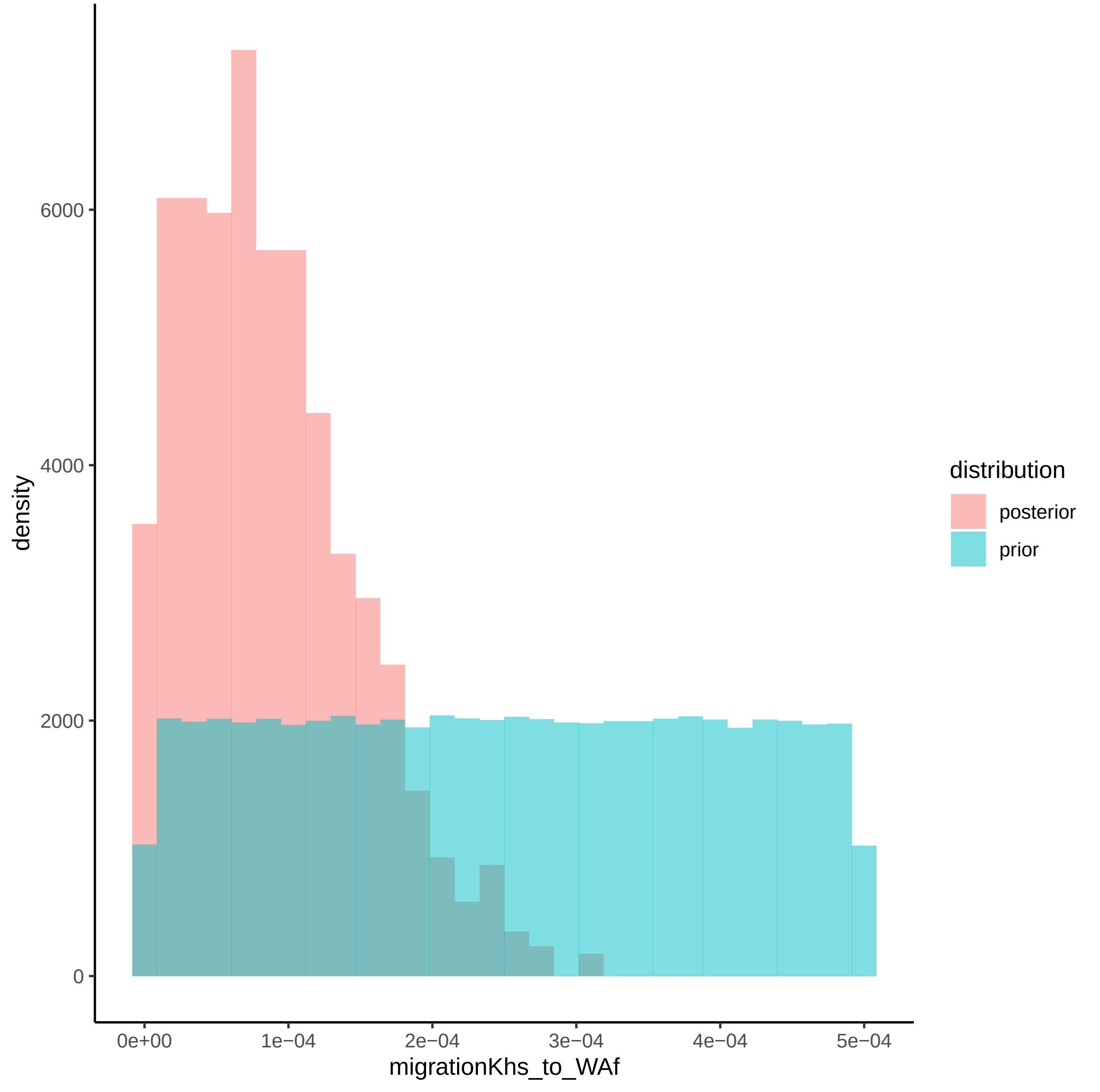

density
6000
4000
2000
0
0e+00
1e−04
2e−04
3e−04
4e−04
5e−04
migrationKhs_to_WAf
distribution
posterior
prior

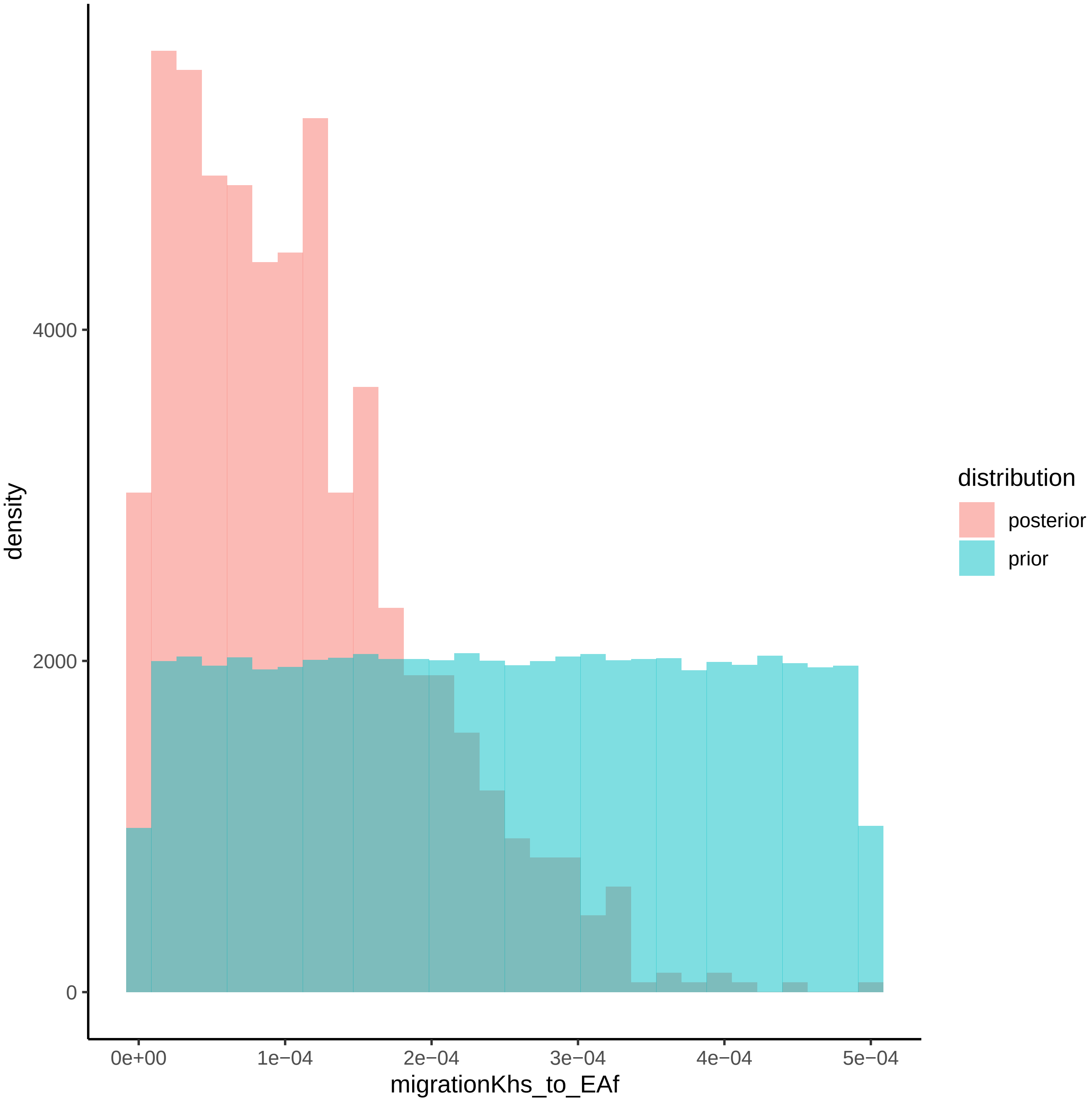
density
4000
2000
0
0e+00
1e−04
2e−04
3e−04
4e−04
5e−04
migrationKhs_to_EAf
distribution
posterior
prior

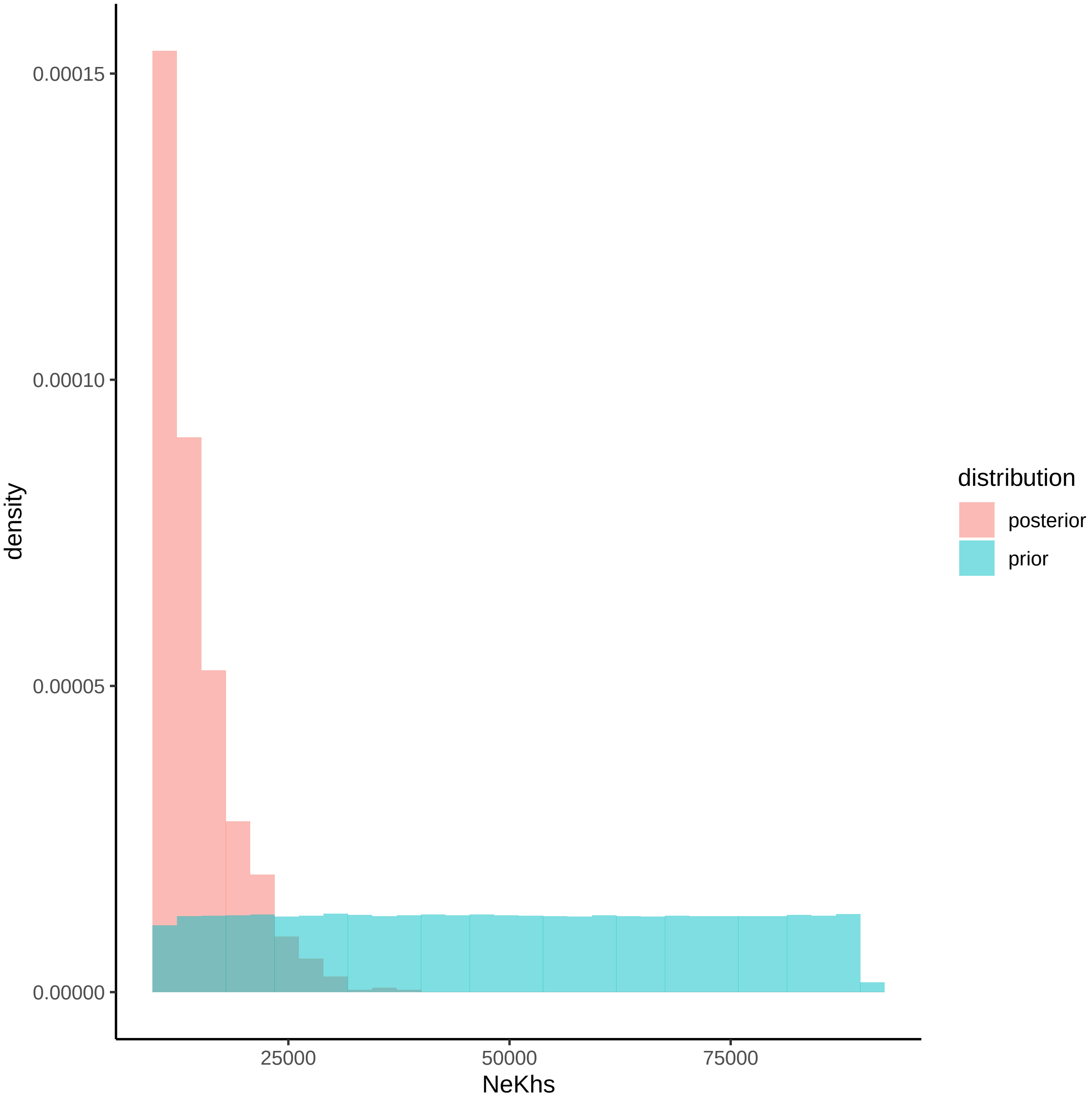
0.00015
0.00010
0.00005
0.00000
density
25000
50000
75000
NeKhs
distribution
posterior
prior

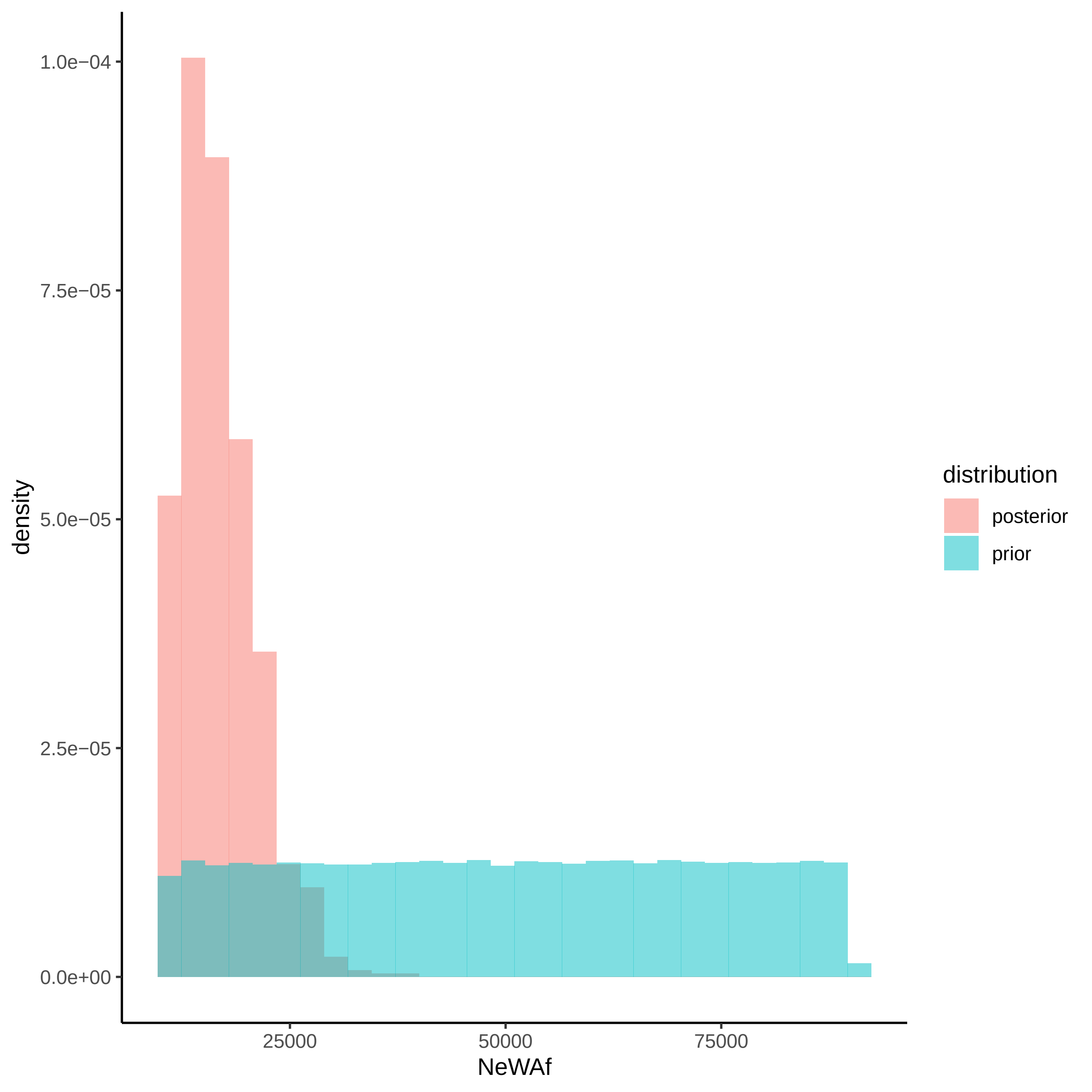

density
1.0e−04
7.5e−05
5.0e−05
2.5e−05
0.0e+00
25000
50000
75000
NeWAf
distribution
posterior
prior

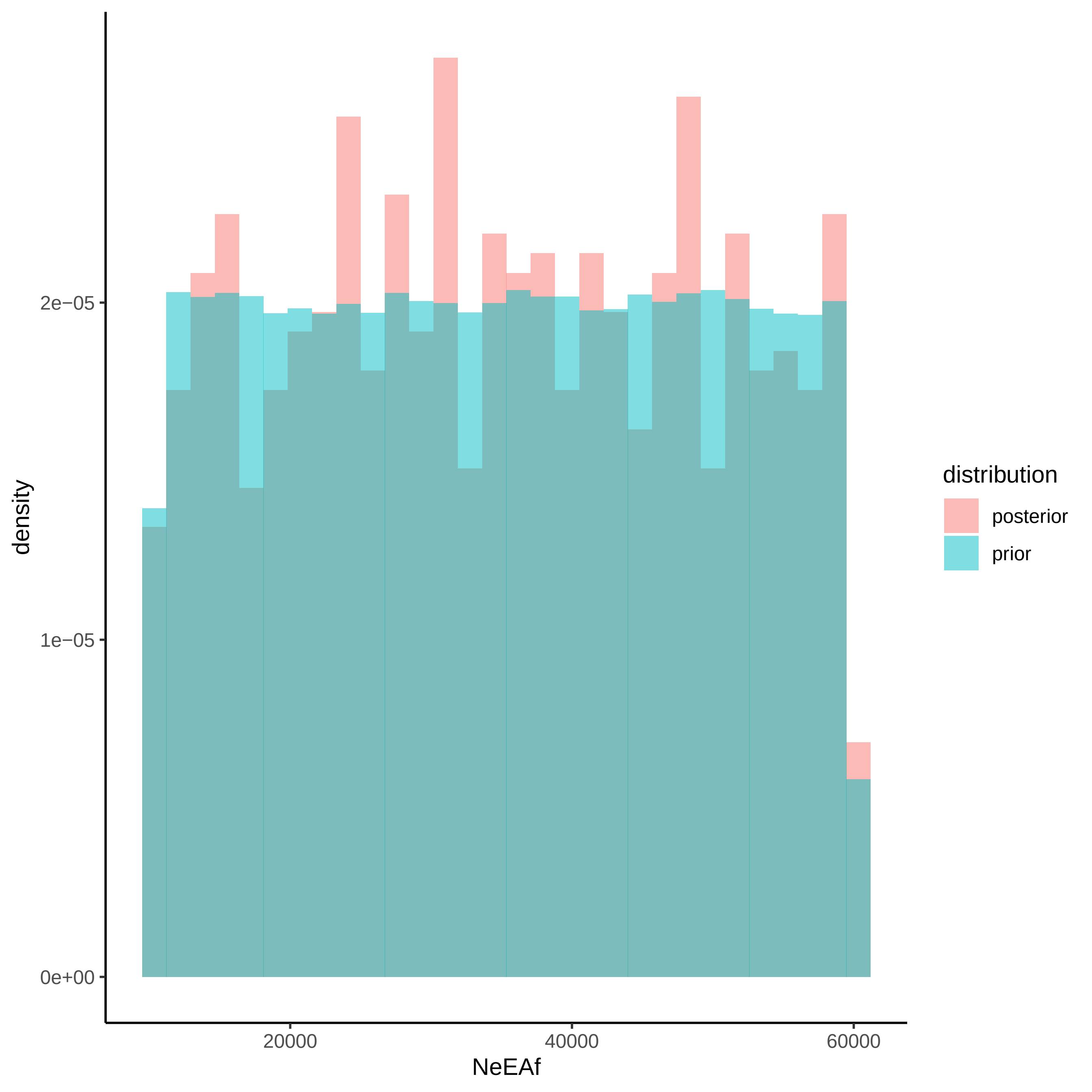

density
2e−05
1e−05
0e+00
20000
40000
60000
NeEAf
distribution
posterior
prior

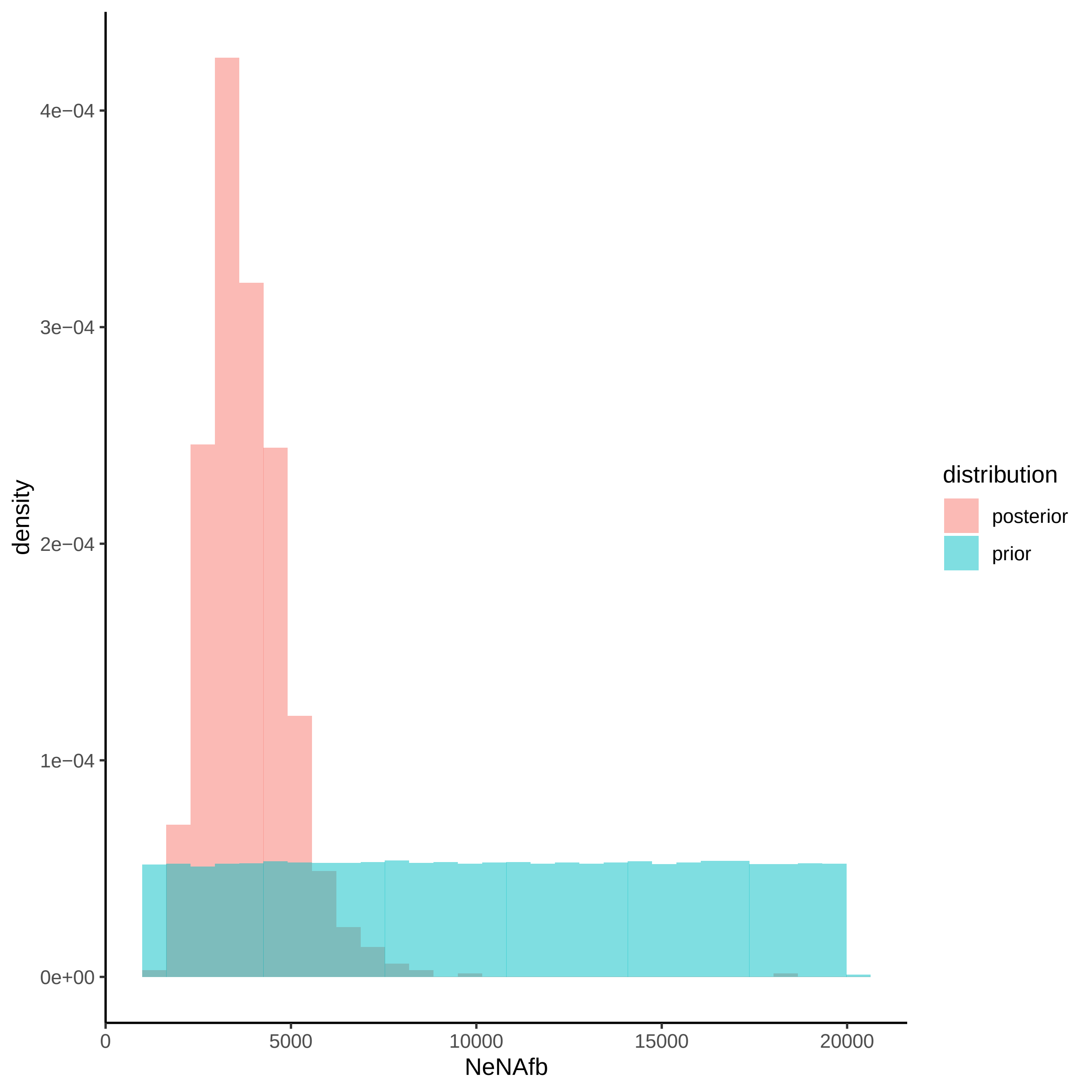

density
4e−04
3e−04
2e−04
1e−04
0e+00
0
5000
10000
15000
20000
NeNAfb
distribution
posterior
prior

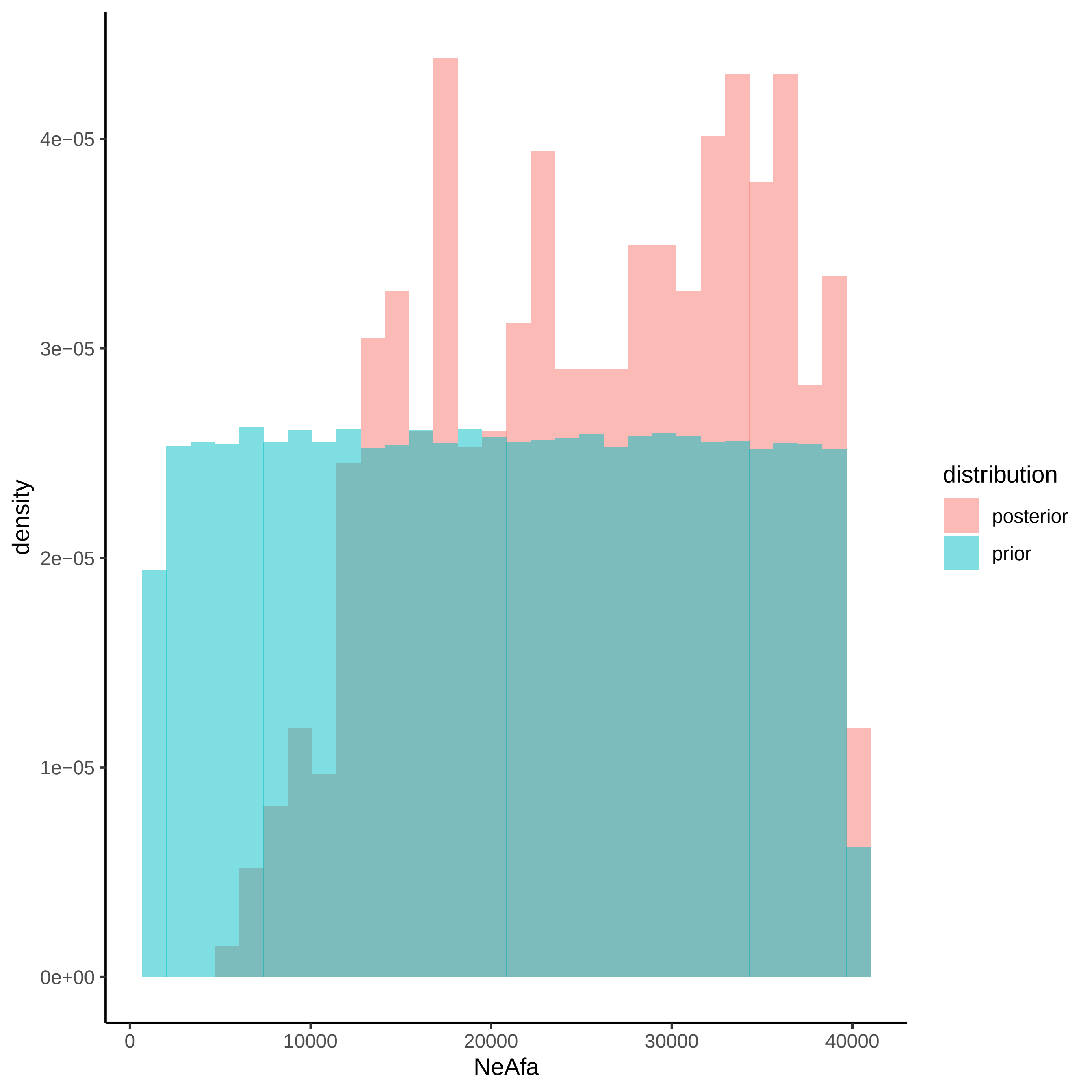

density
4e−05
3e−05
2e−05
1e−05
0e+00
0
10000
20000
30000
40000
NeAfa
distribution
posterior
prior

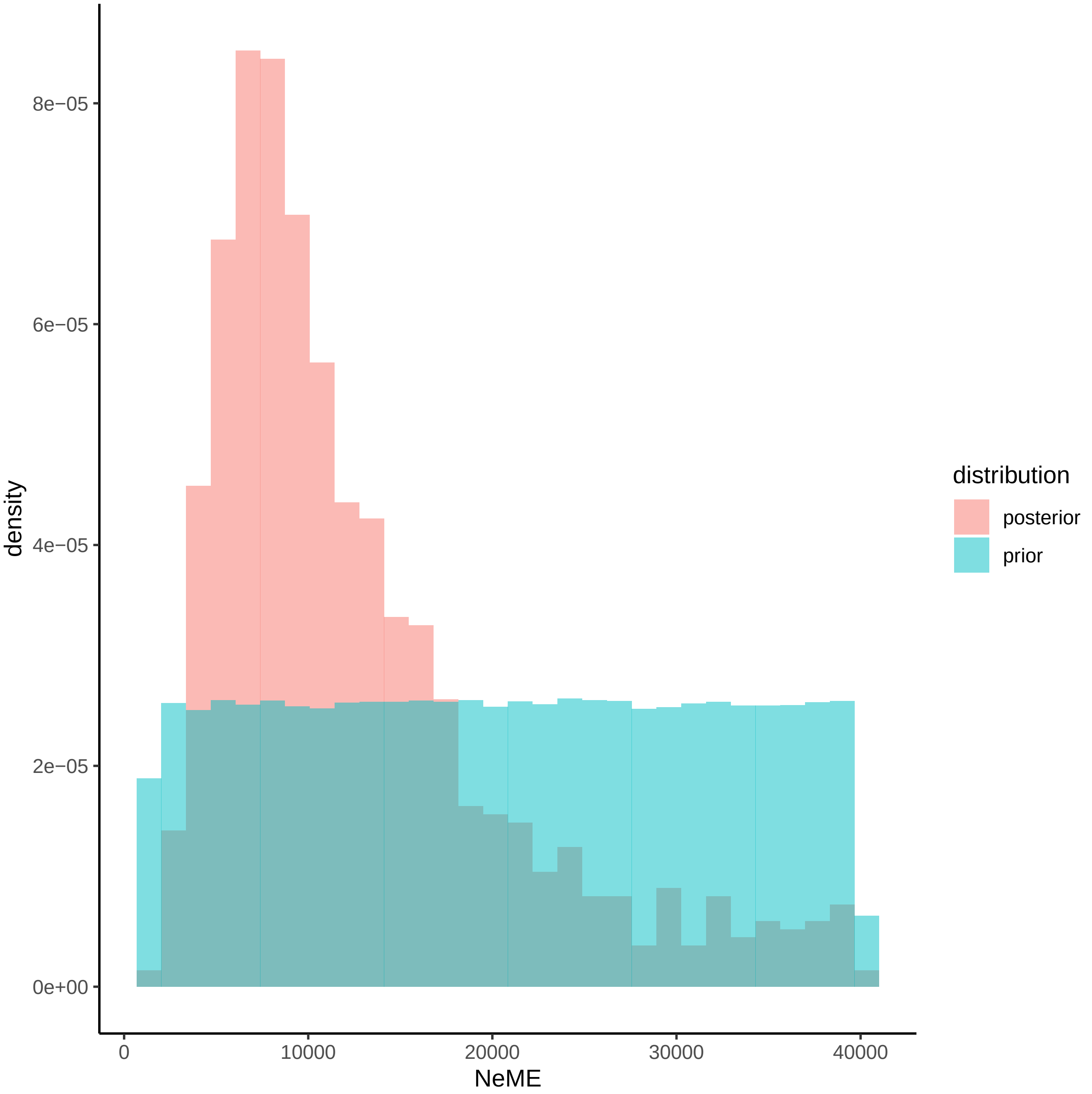

8e−05
6e−05
4e−05
2e−05
0e+00
density
0
10000
20000
30000
40000
NeME
distribution
posterior
prior

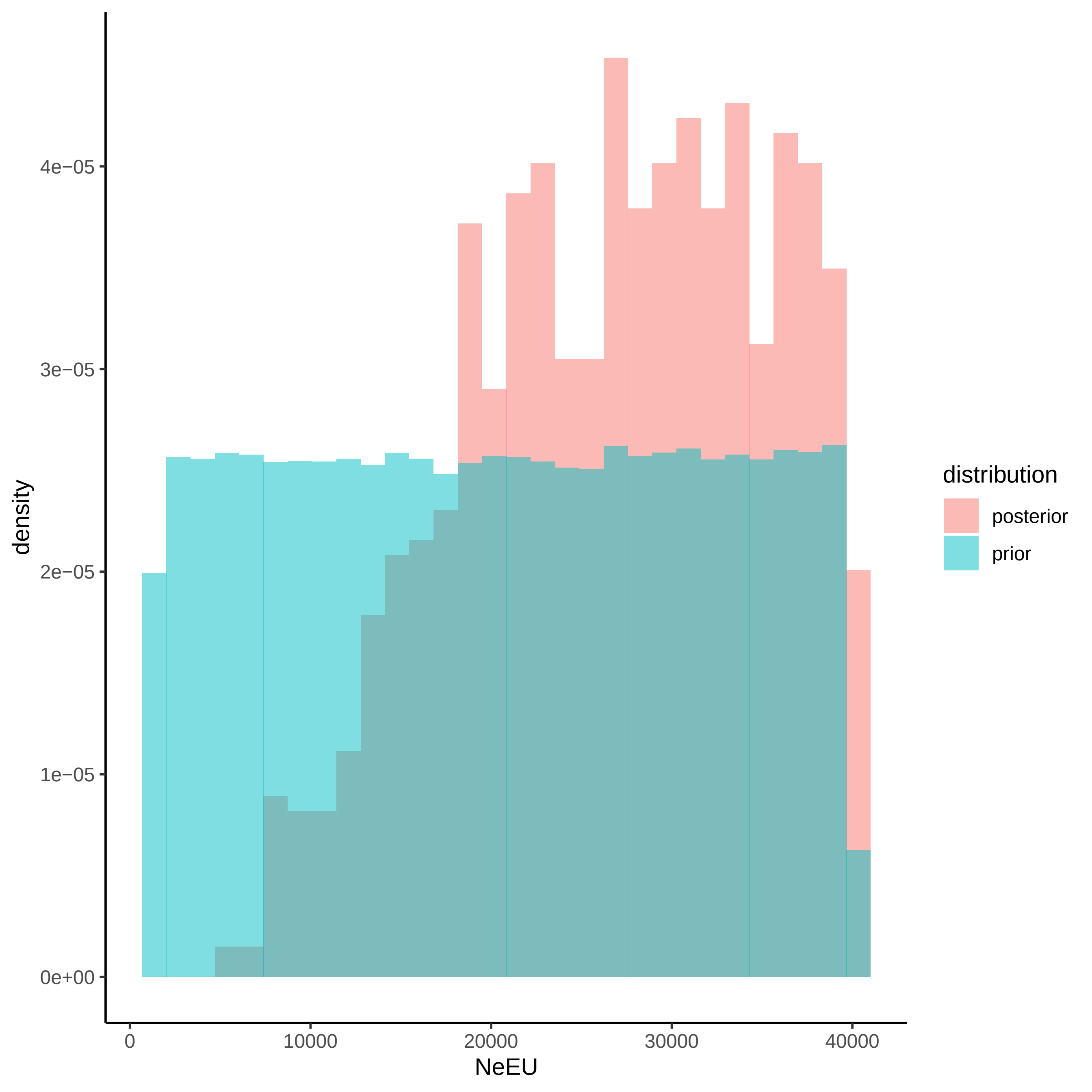
density
4e−05
3e−05
2e−05
1e−05
0e+00
0
10000
20000
30000
40000
NeEU
distribution
posterior
prior

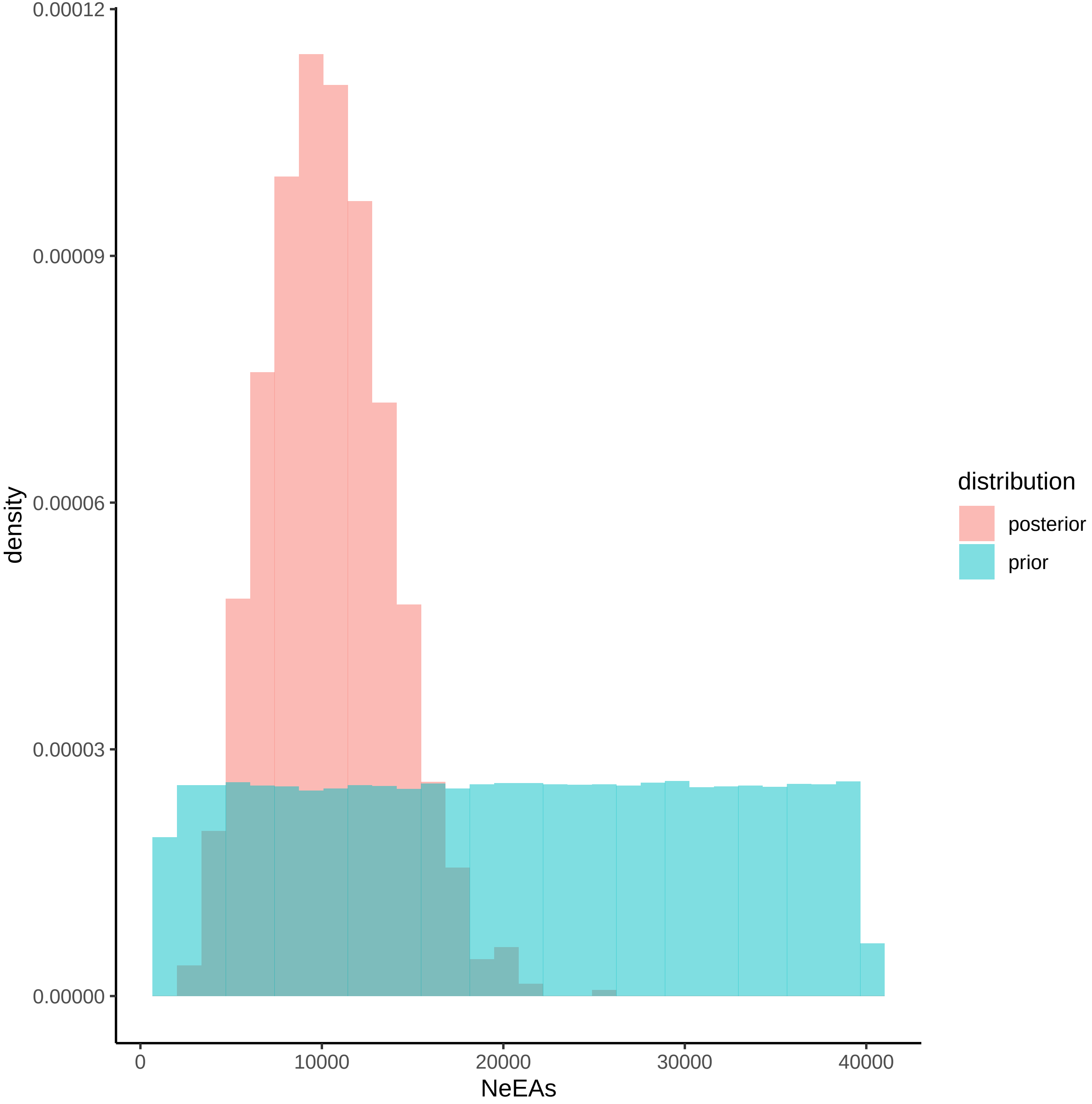

0.00012
0.00009
0.00006
0.00003
0.00000
density
0
10000
20000
30000
40000
NeEAs
distribution
posterior
prior

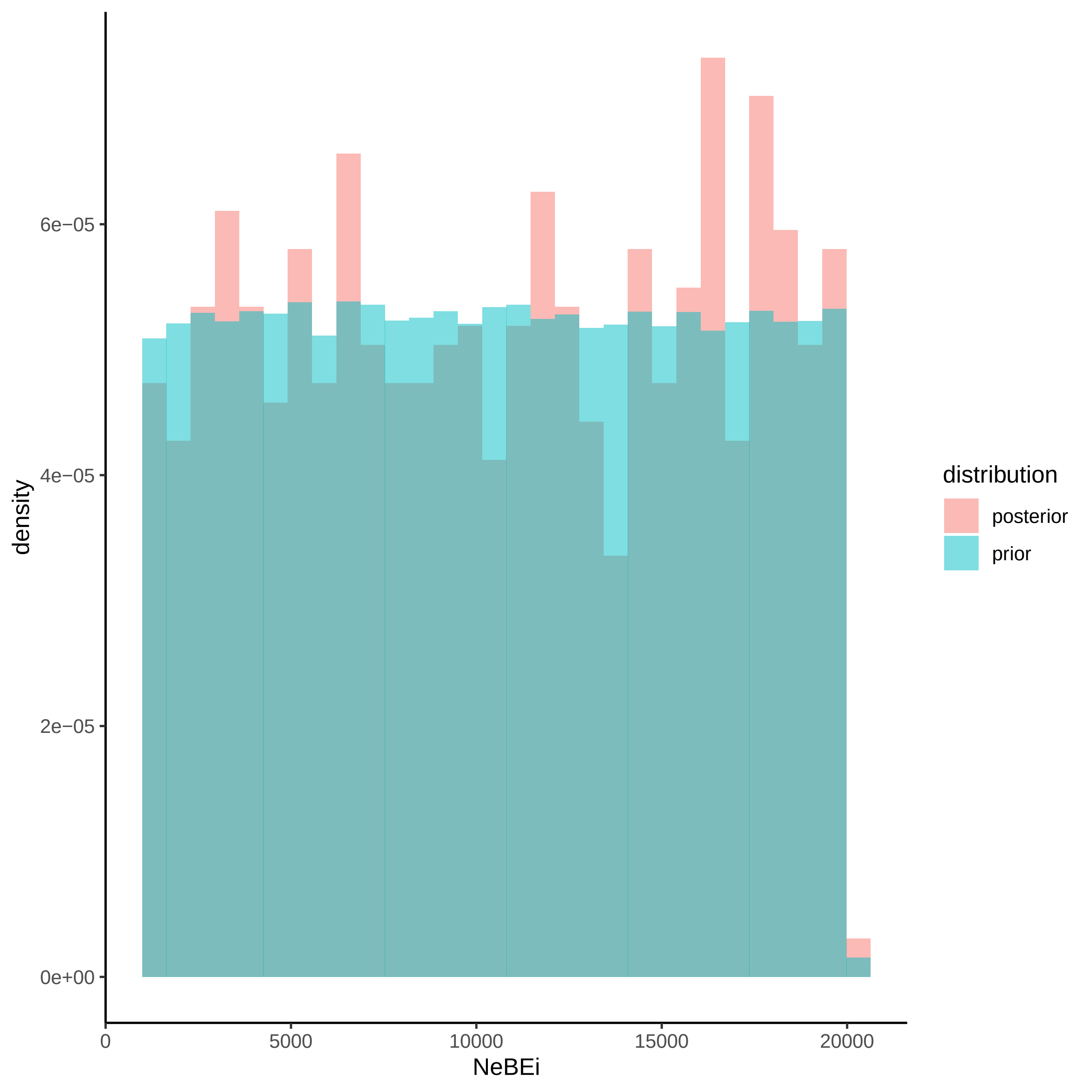
density
6e−05
4e−05
2e−05
0e+00
0
5000
10000
15000
20000
NeBEi
distribution
posterior
prior

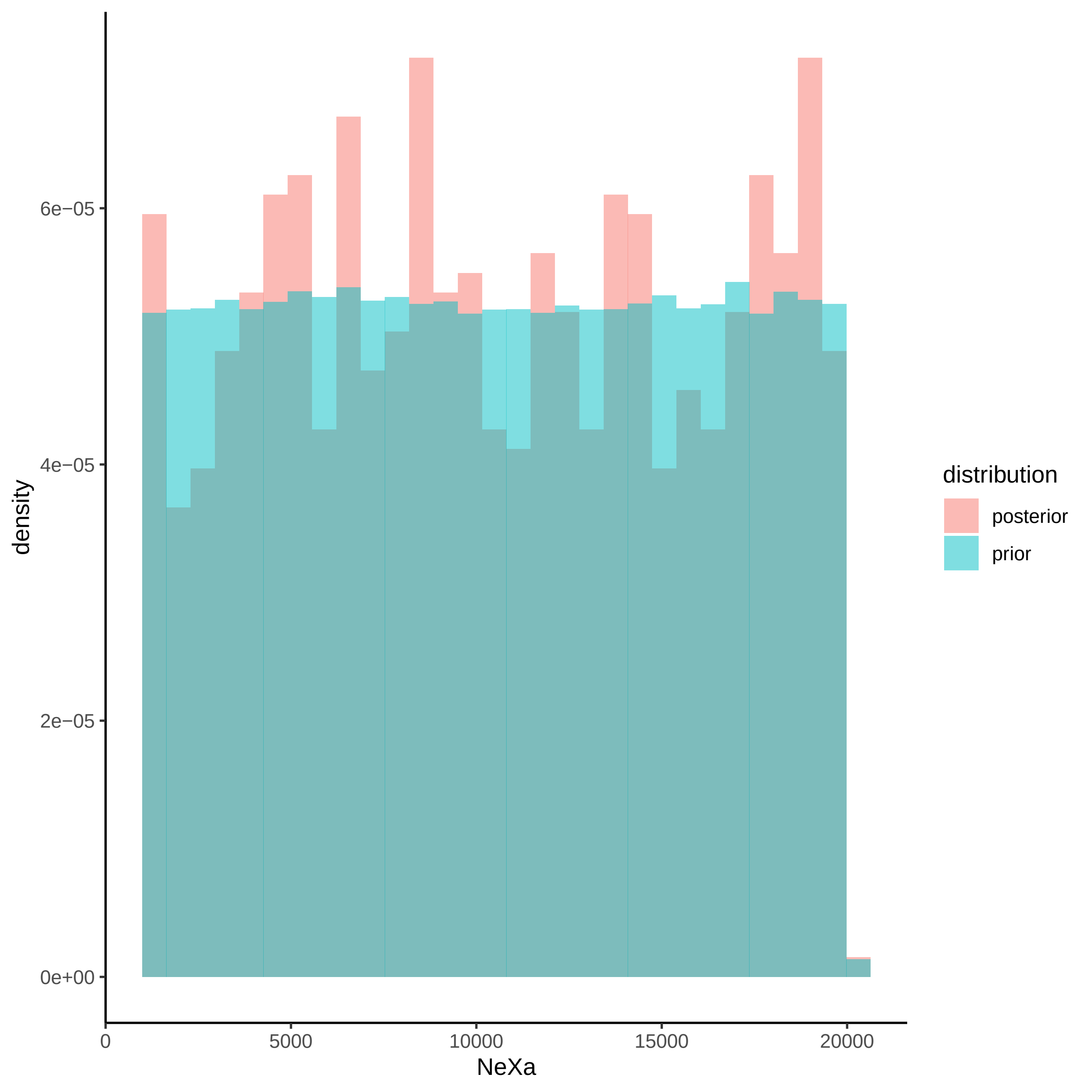

6e−05
4e−05
2e−05
0e+00
density
0
5000
10000
15000
20000
NeXa
distribution
posterior
prior

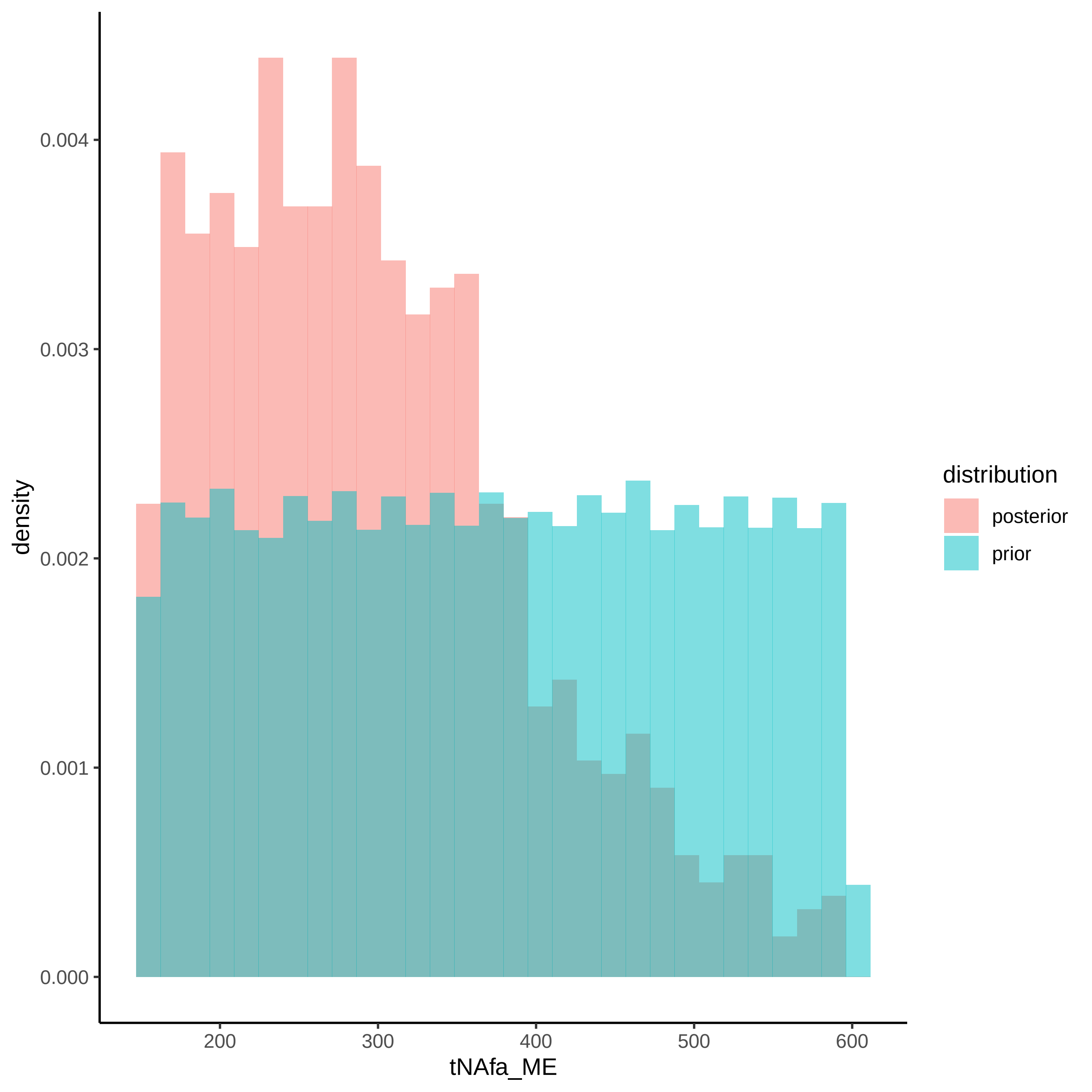

density
0.000
0.001
0.002
0.003
0.004
200
300
400
500
600
tNAfa_ME
distribution
posterior
prior

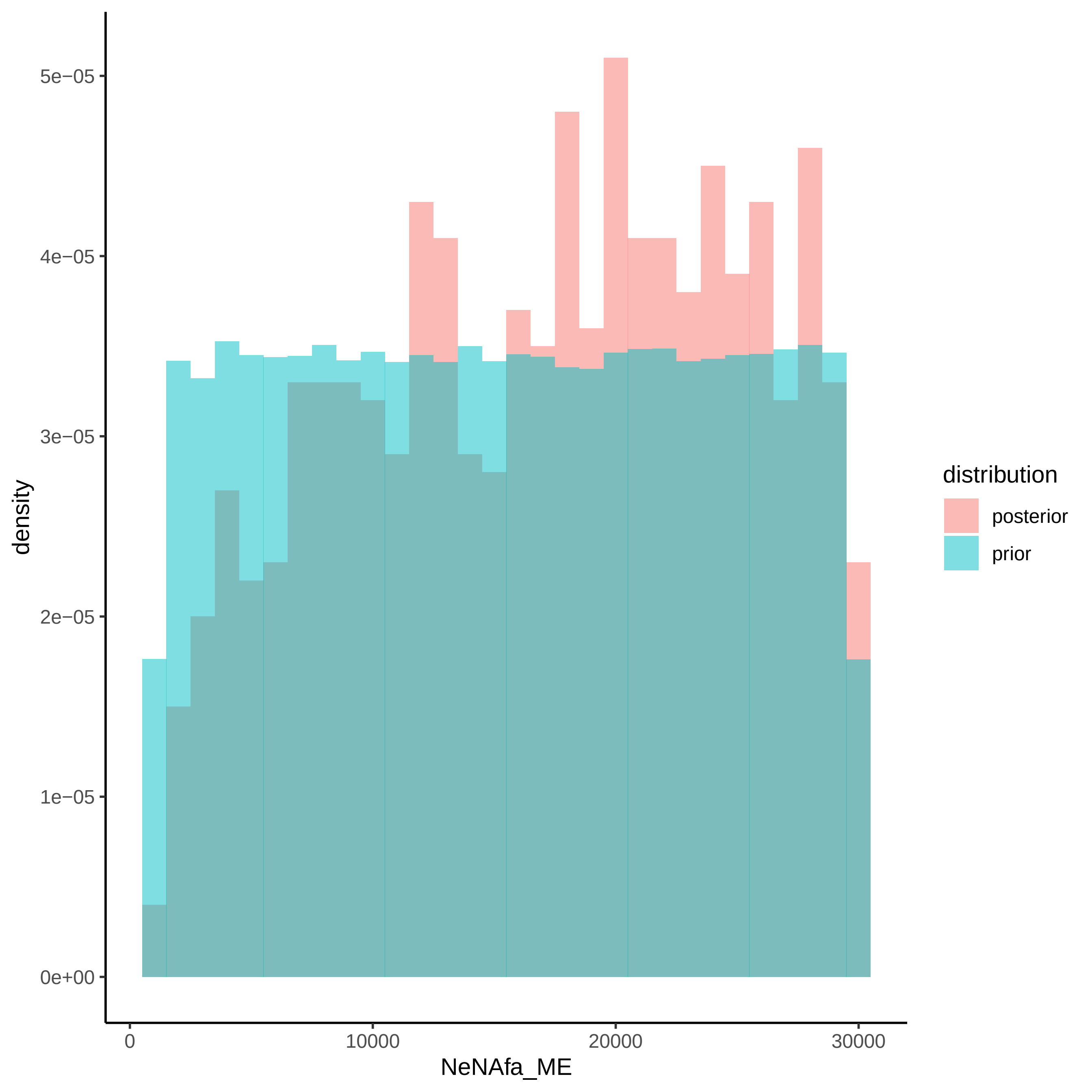

density
5e−05
4e−05
3e−05
2e−05
1e−05
0e+00
0
10000
20000
30000
NeNAfa_ME
distribution
posterior
prior

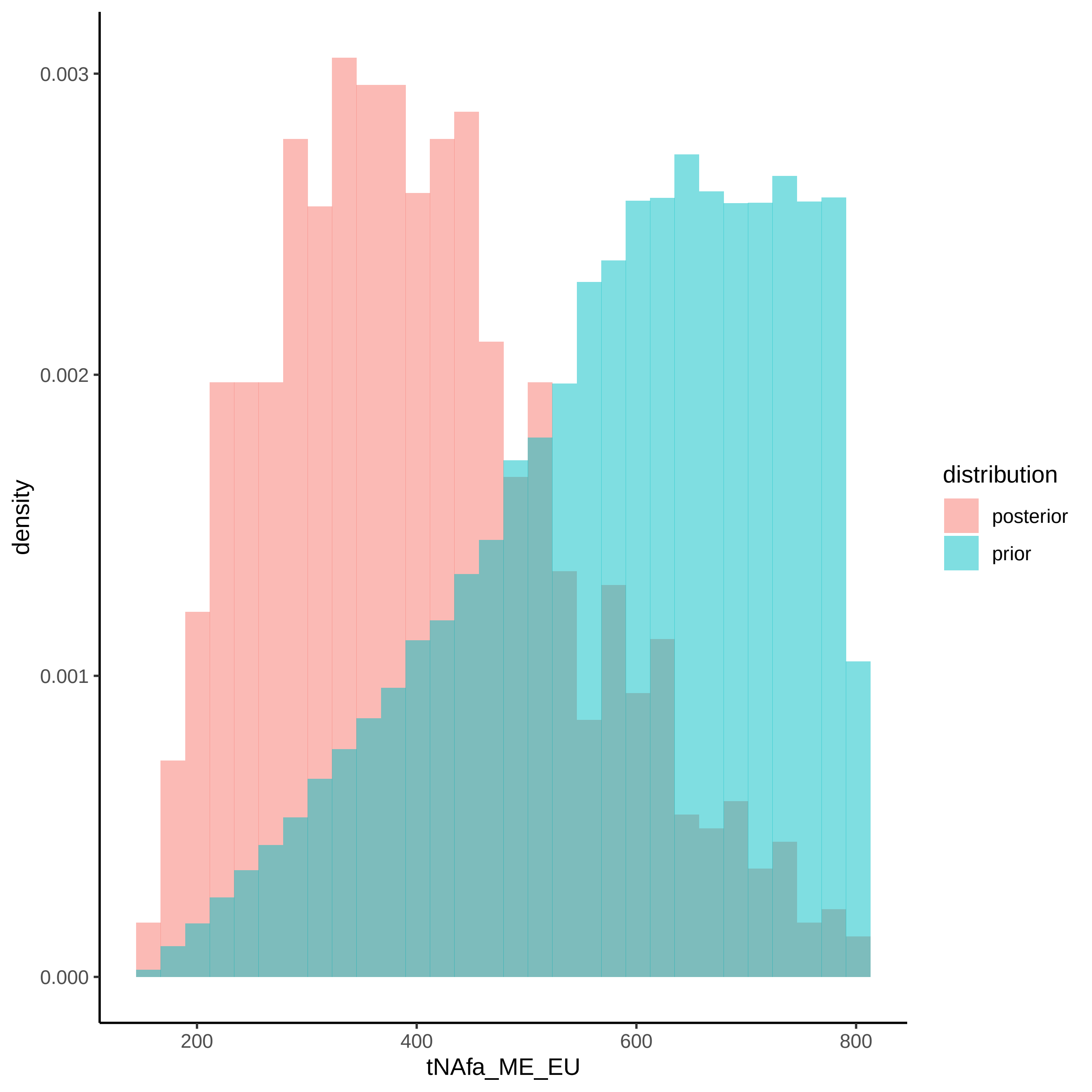

0.003
0.002
0.001
0.000
density
200
400
600
800
tNAfa_ME_EU
distribution
posterior
prior

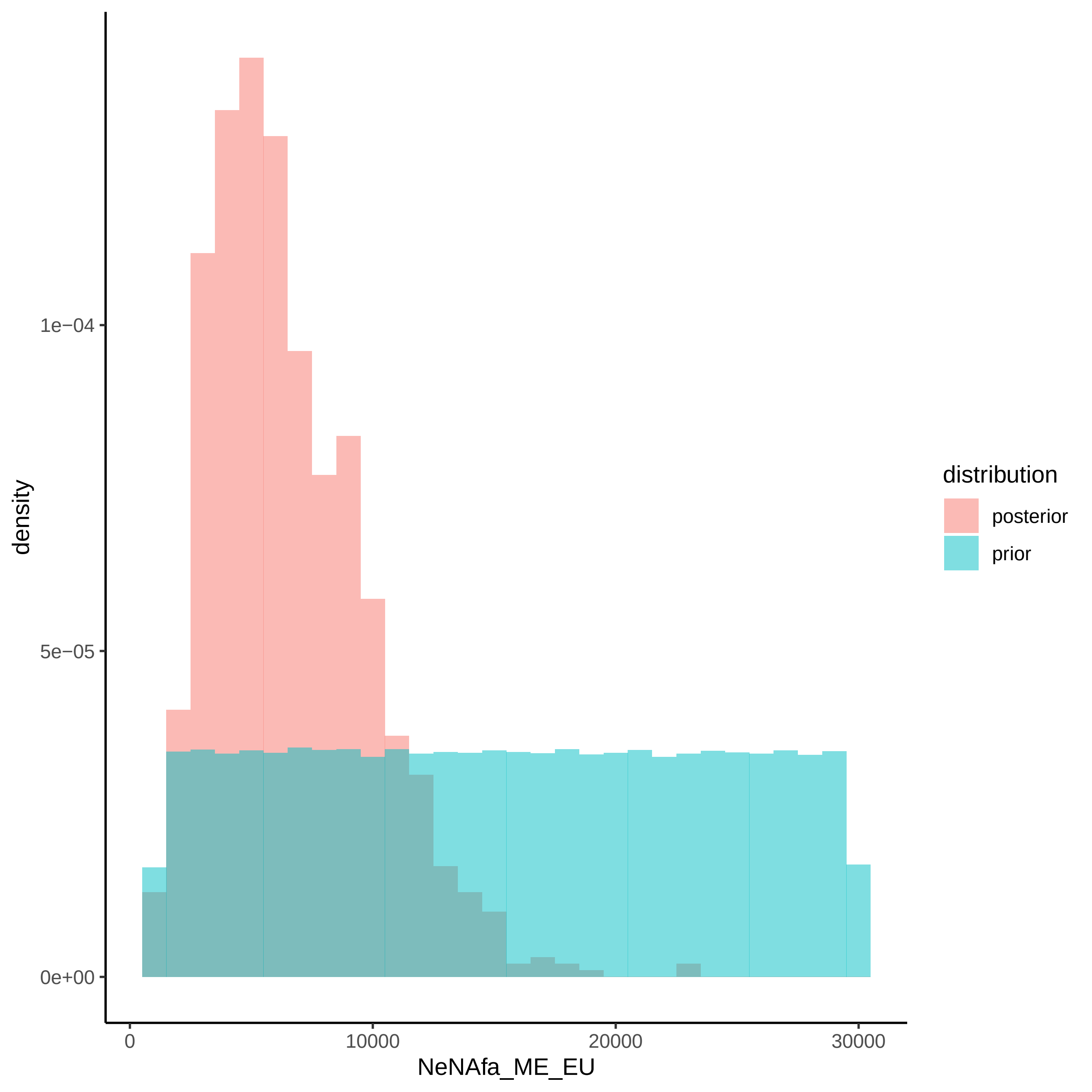

density
1e−04
5e−05
0e+00
0
10000
20000
30000
NeNAfa_ME_EU
distribution
posterior
prior

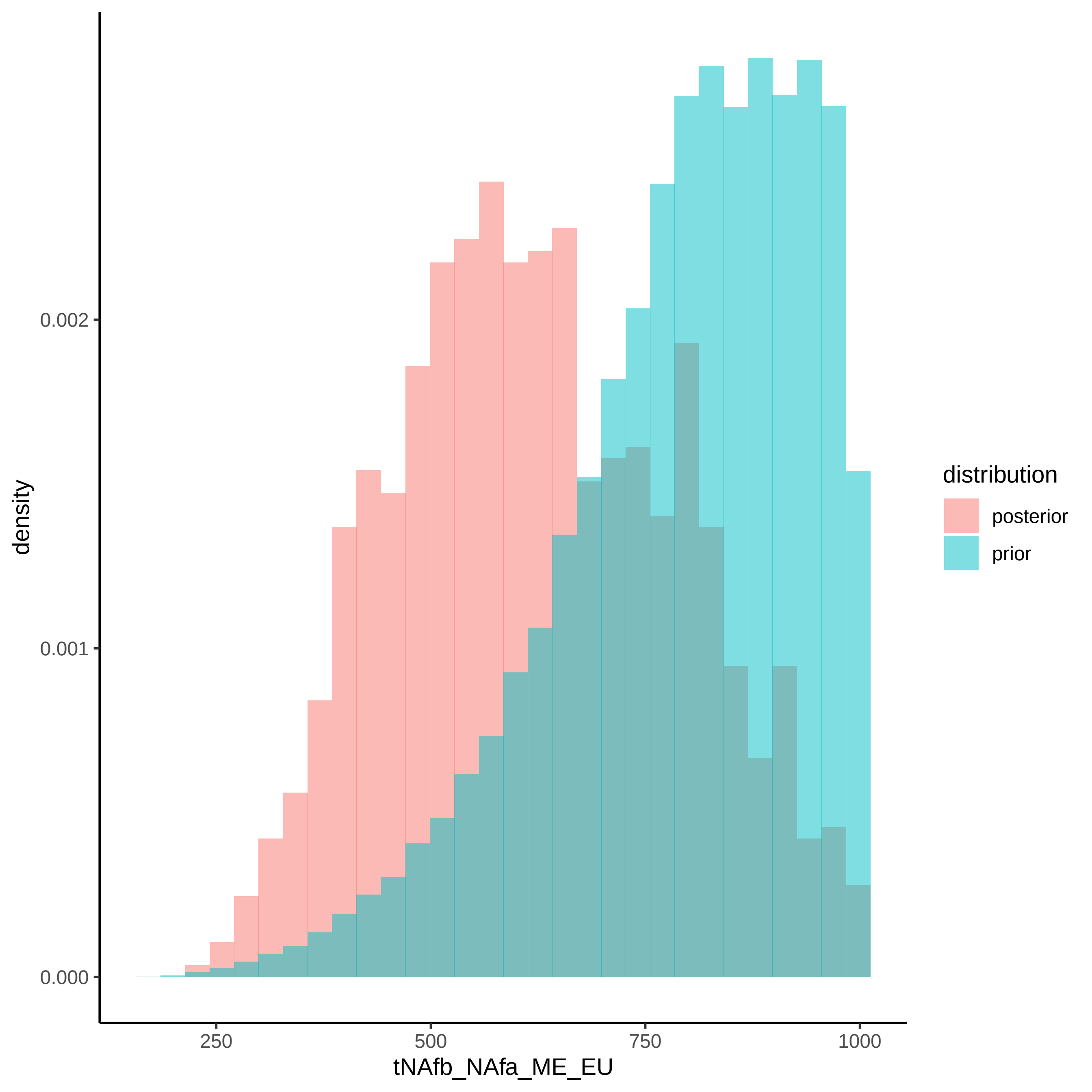

density
0.000
0.001
0.002
250
500
750
1000
tNAfb_NAfa_ME_EU
distribution
posterior
prior

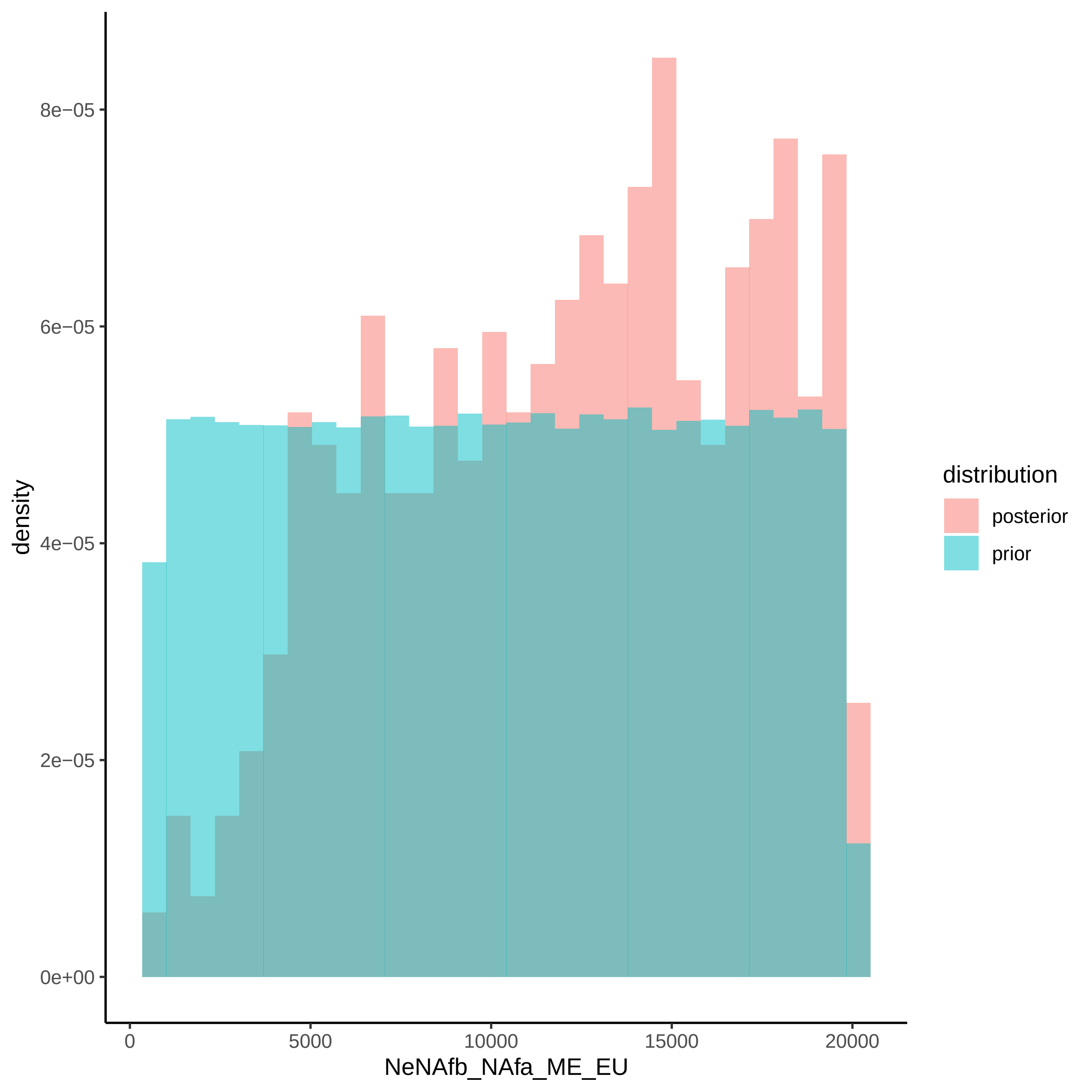

density
8e−05
6e−05
4e−05
2e−05
0e+00
0
5000
10000
15000
20000
NeNAfb_NAfa_ME_EU
distribution
posterior
prior

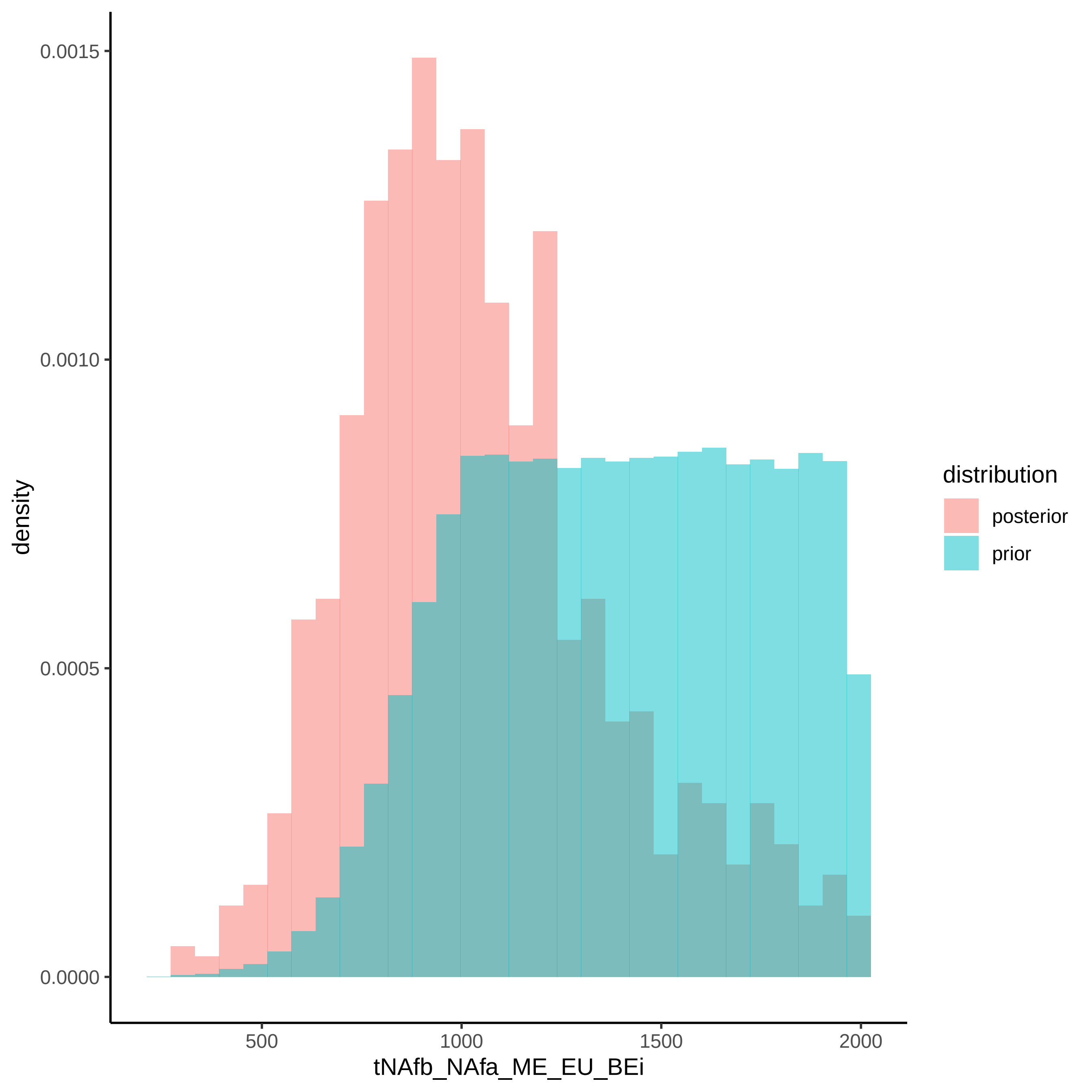
0.0015
0.0010
0.0005
0.0000
density
500
1000
1500
2000
tNAfb_NAfa_ME_EU_BEi
distribution
posterior
prior

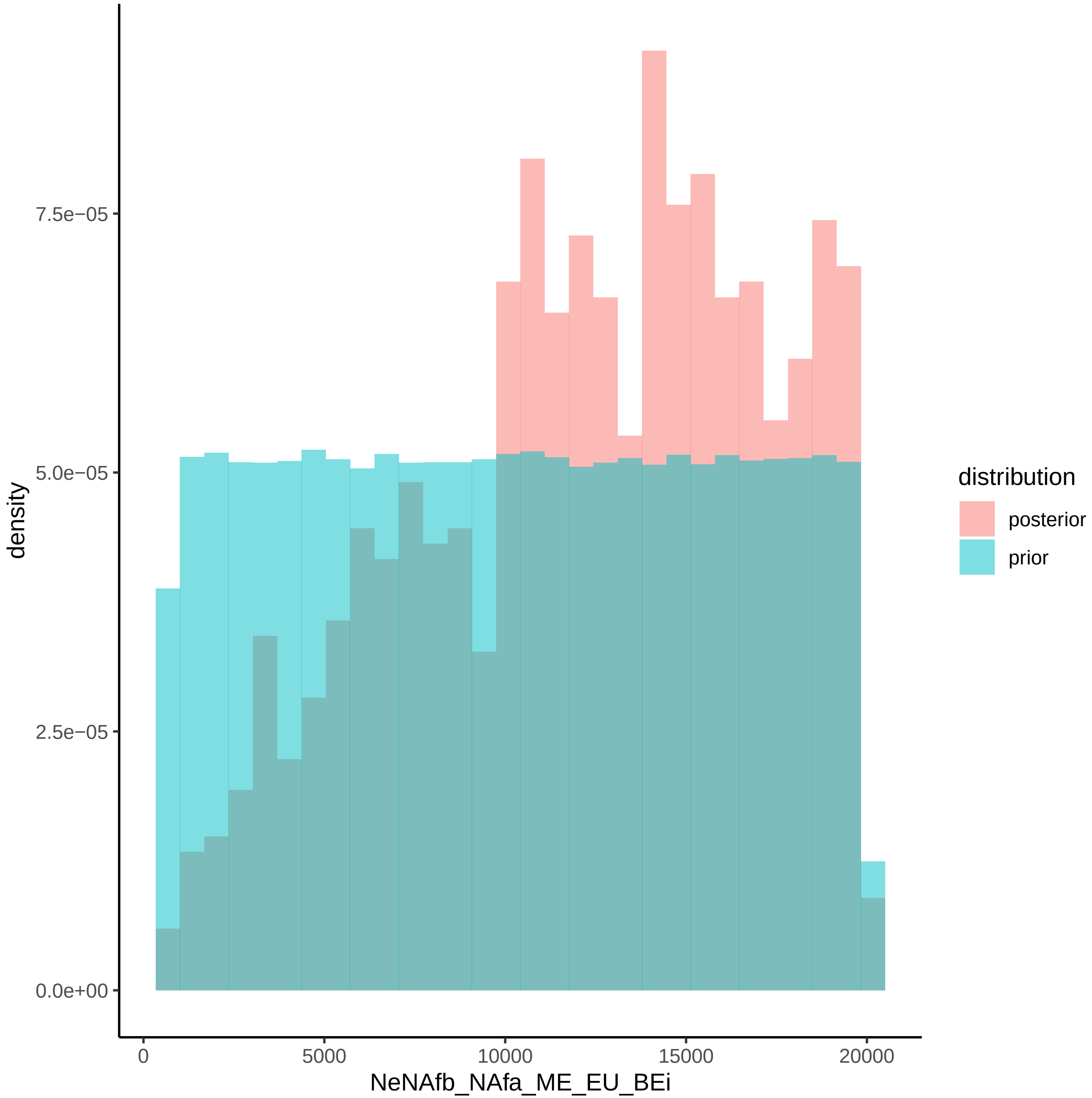

density
7.5e−05
5.0e−05
2.5e−05
0.0e+00
0
5000
10000
15000
20000
NeNAfb_NAfa_ME_EU_BEi
distribution
posterior
prior

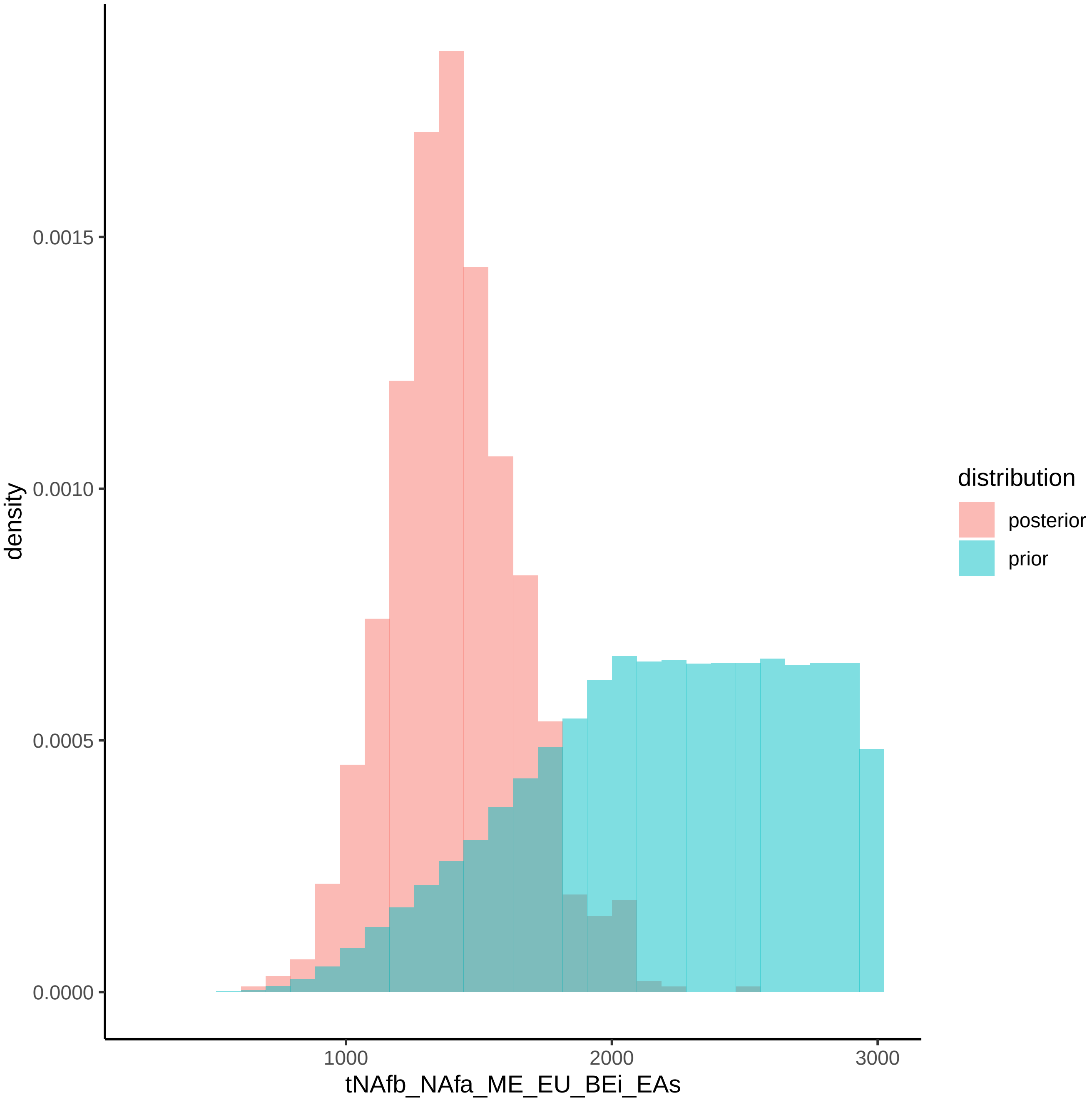

density
0.0015
0.0010
0.0005
0.0000
1000
2000
3000
tNAfb_NAfa_ME_EU_BEi_EAs
distribution
posterior
prior

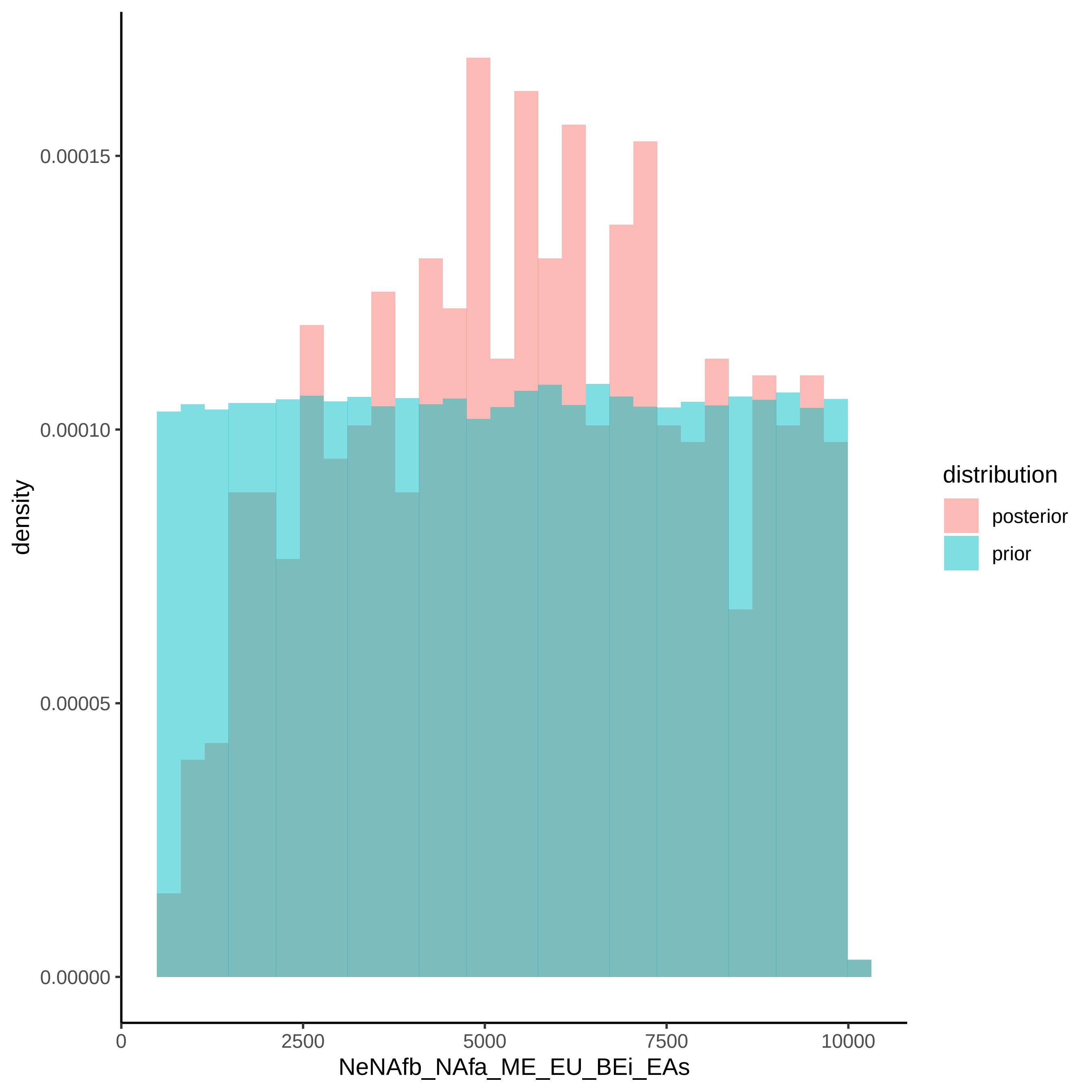

density
0.00015
0.00010
0.00005
0.00000
0
2500
5000
7500
10000
NeNAfb_NAfa_ME_EU_BEi_EAs
distribution
posterior
prior

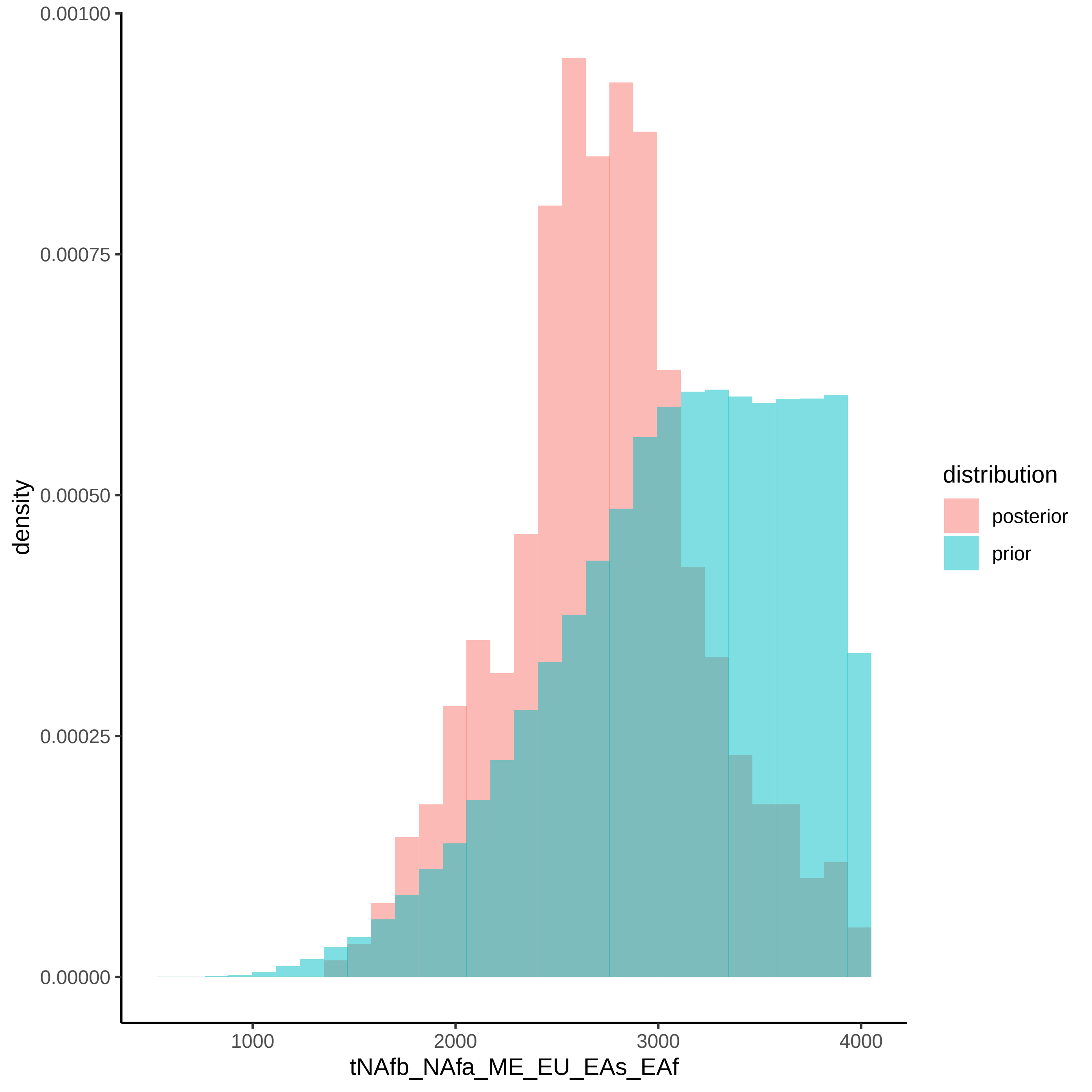
0.00100
0.00075
0.00050
0.00025
0.00000
density
1000
2000
3000
4000
tNAfb_NAfa_ME_EU_EAs_EAf
distribution
posterior
prior

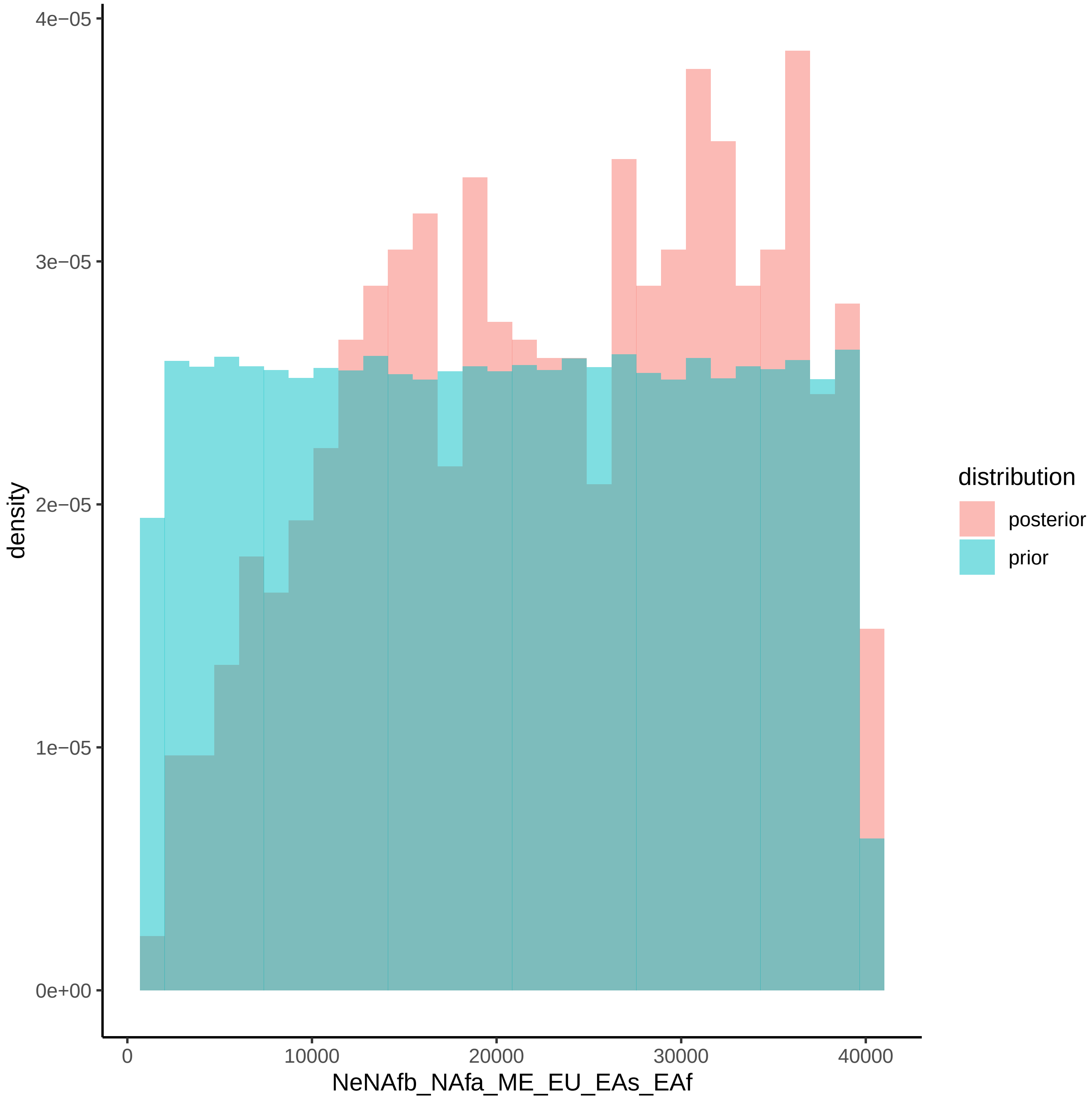
4e−05
3e−05
2e−05
1e−05
0e+00
density
0
10000
20000
30000
40000
NeNAfb_NAfa_ME_EU_EAs_EAf
distribution
posterior
prior

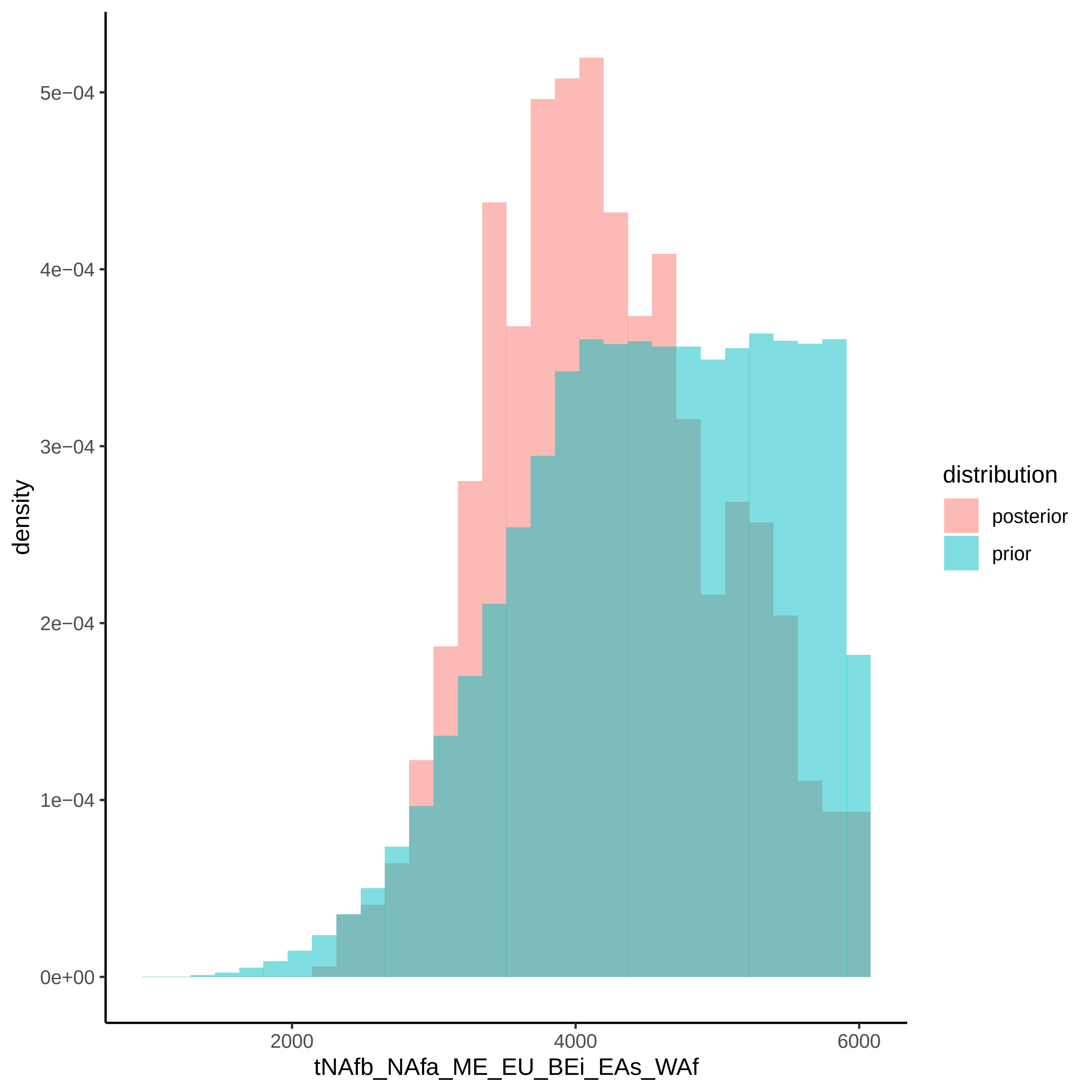

density
5e−04
4e−04
3e−04
2e−04
1e−04
0e+00
2000
4000
6000
tNAfb_NAfa_ME_EU_BEi_EAs_WAf
distribution
posterior
prior

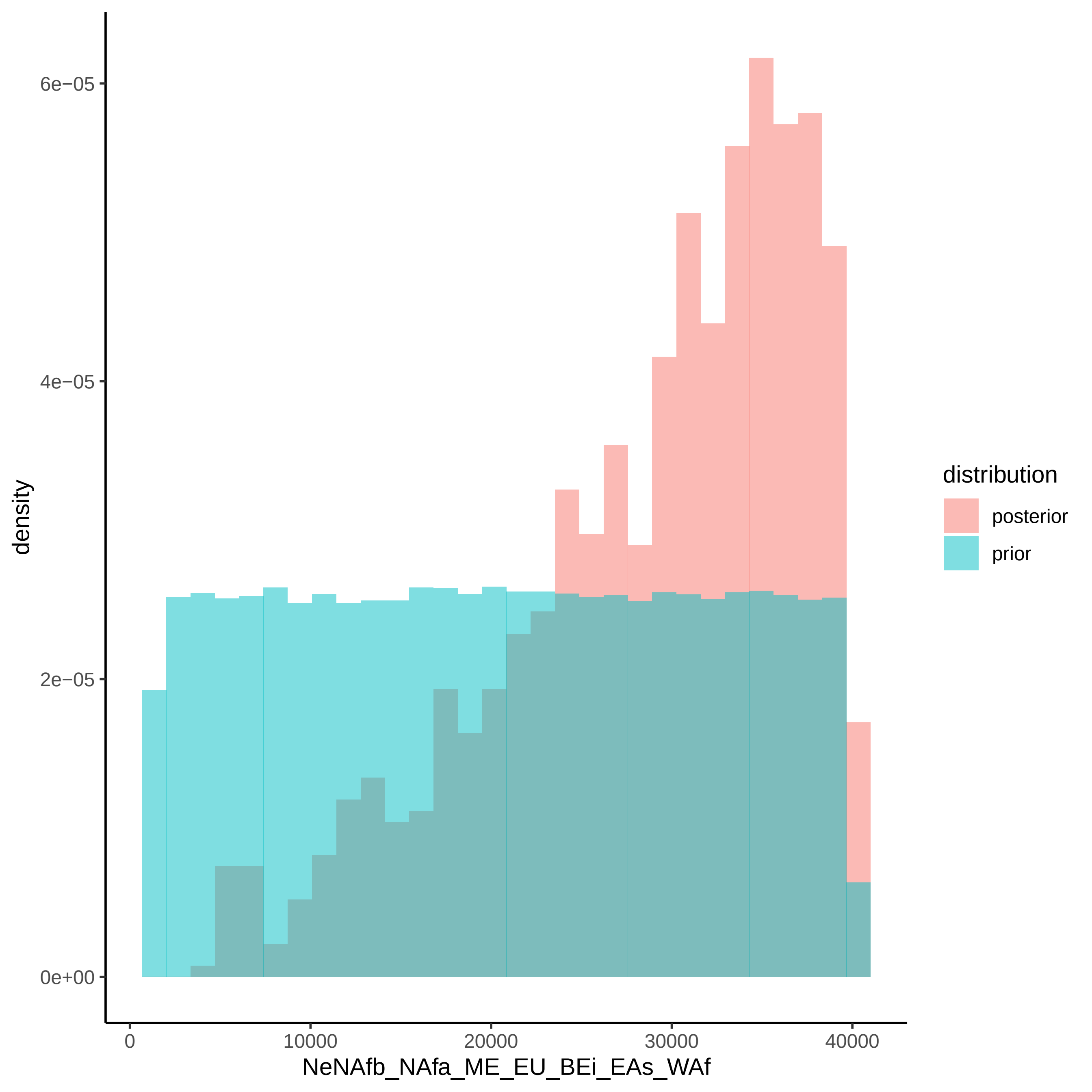

density
6e−05
4e−05
2e−05
0e+00
0
10000
20000
30000
40000
NeNAfb_NAfa_ME_EU_BEi_EAs_WAf
distribution
posterior
prior

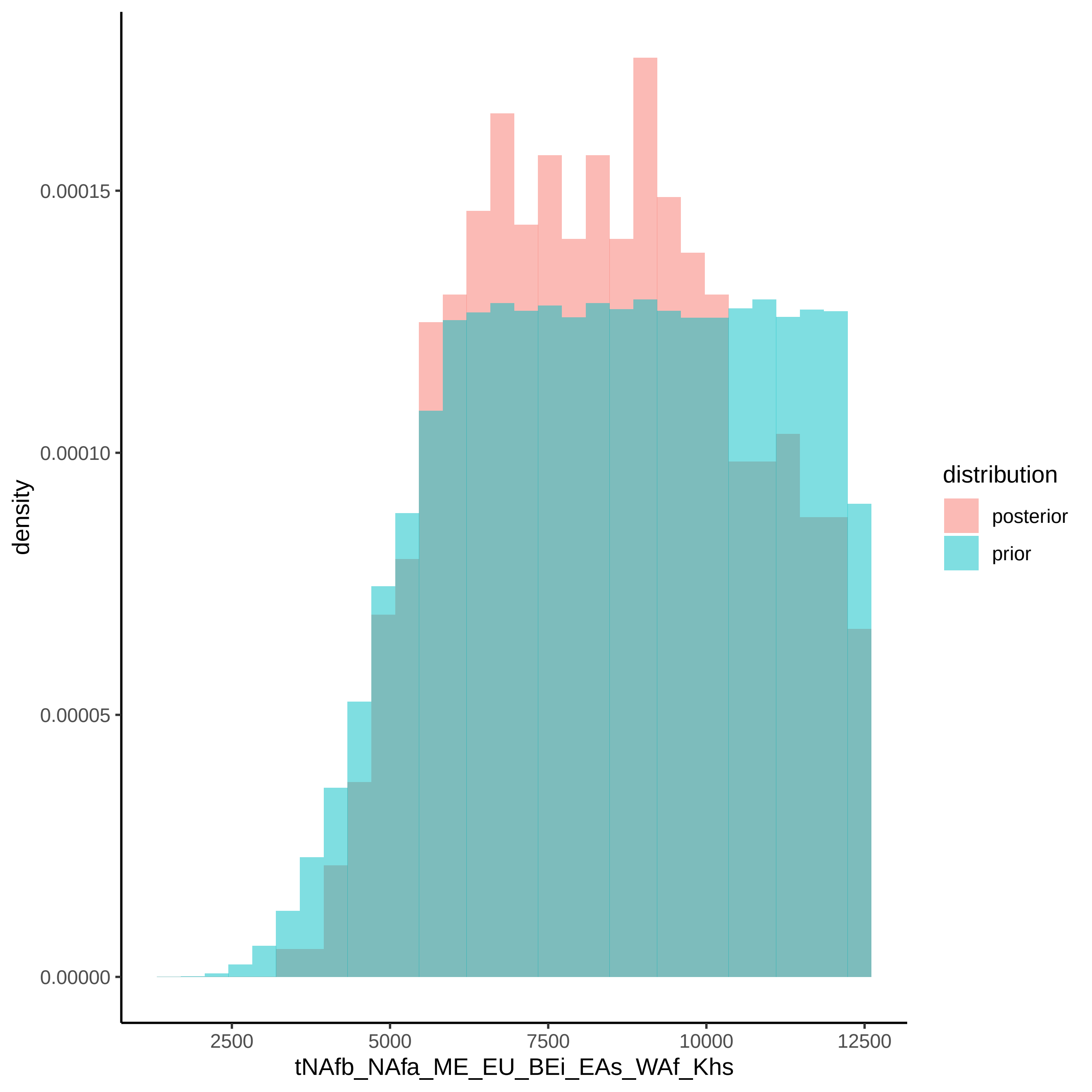

density
0.00015
0.00010
0.00005
0.00000
2500
5000
7500
10000
12500
tNAfb_NAfa_ME_EU_BEi_EAs_WAf_Khs
distribution
posterior
prior

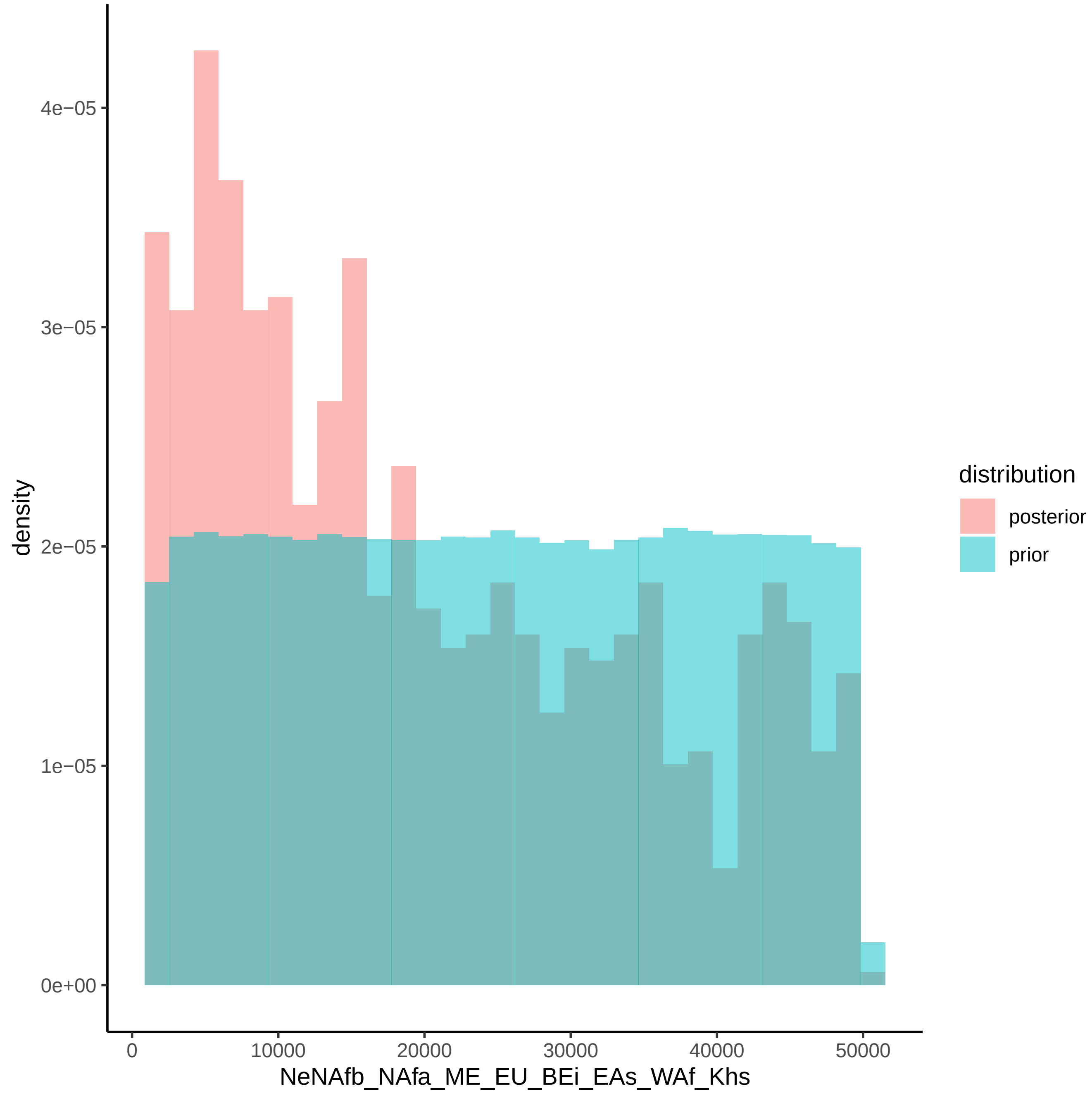

4e−05
3e−05
2e−05
1e−05
0e+00
density
0
10000
20000
30000
40000
50000
NeNAfb_NAfa_ME_EU_BEi_EAs_WAf_Khs
distribution
posterior
prior

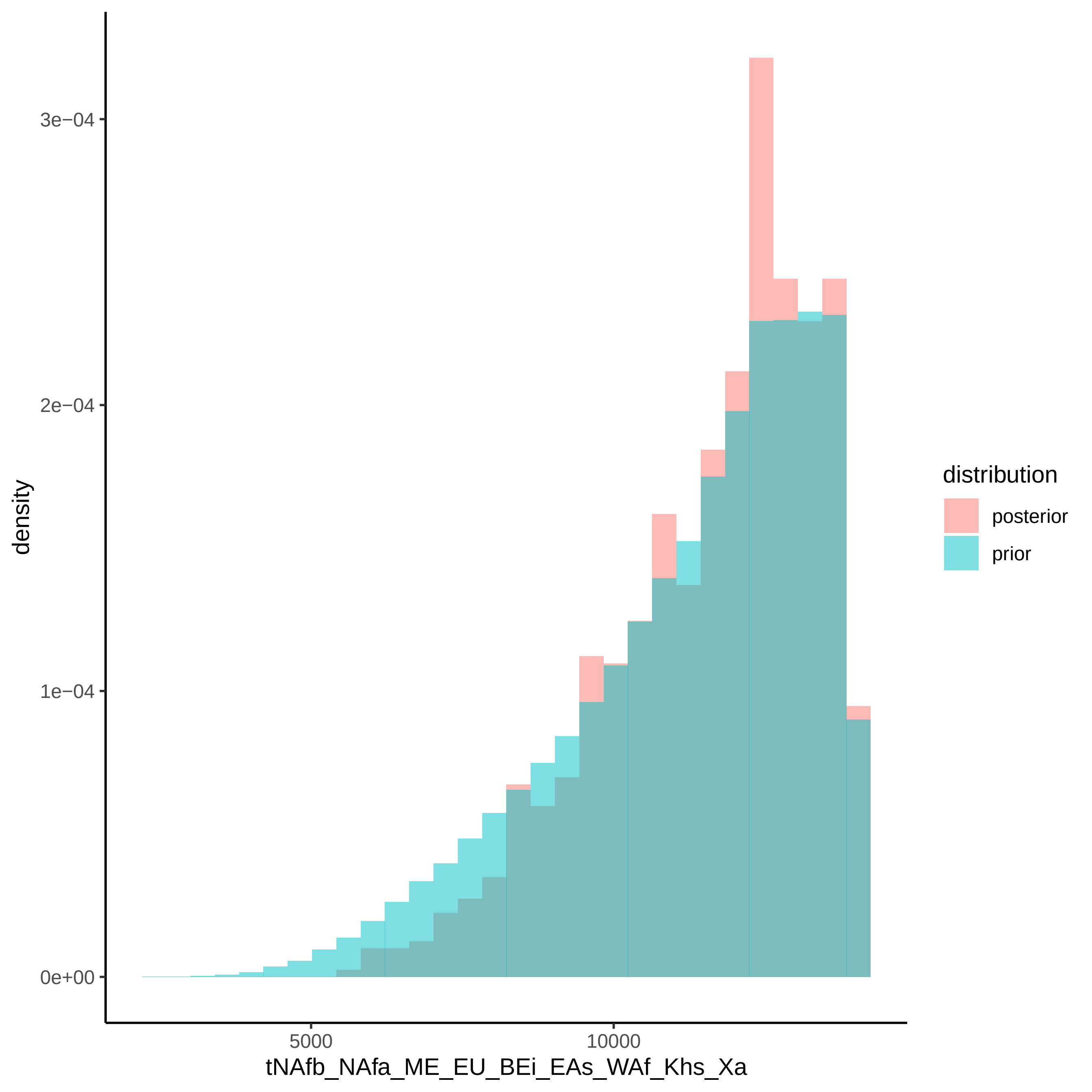
3e−04
2e−04
1e−04
0e+00
density
5000
10000
tNAfb_NAfa_ME_EU_BEi_EAs_WAf_Khs_Xa
distribution
posterior
prior

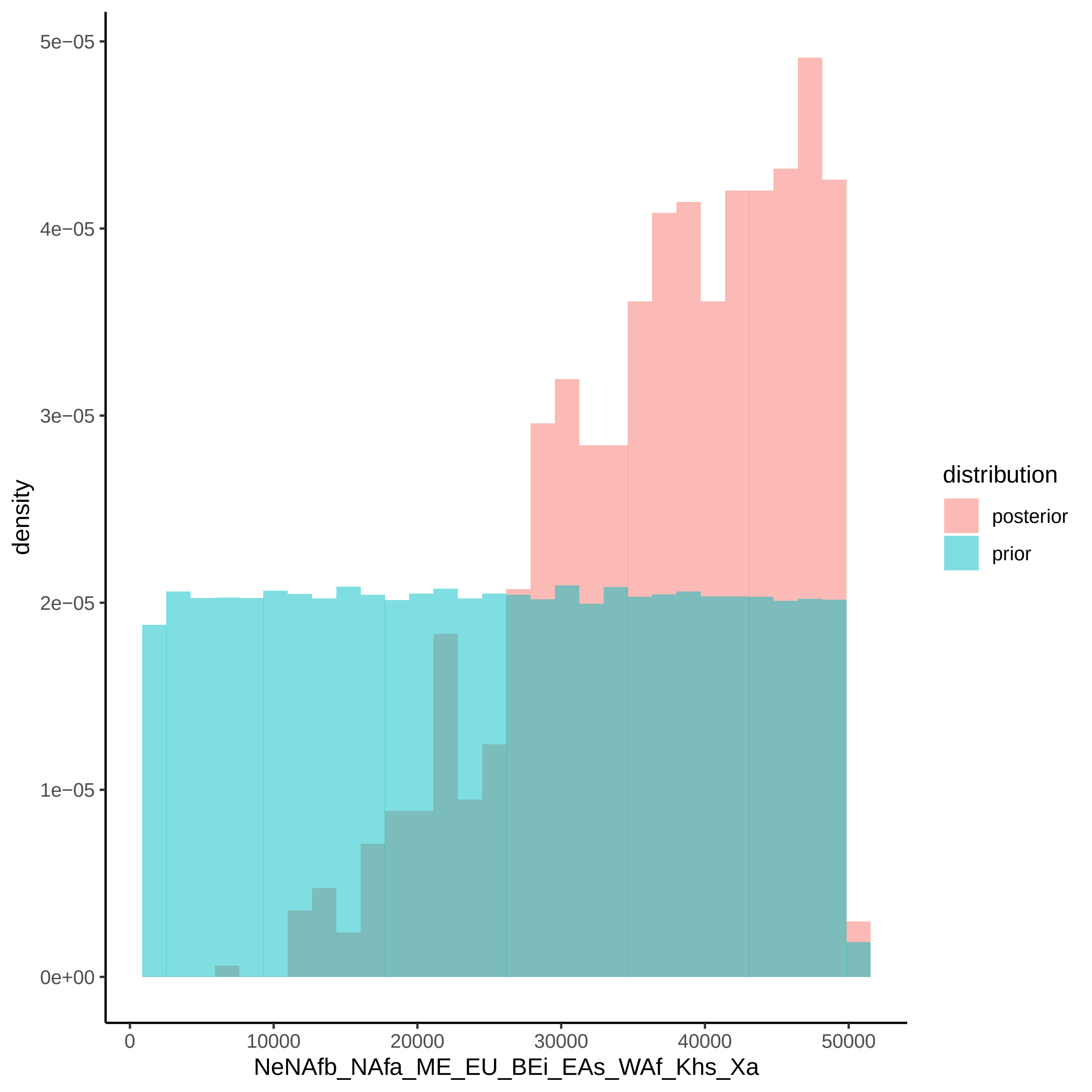

5e−05
4e−05
3e−05
2e−05
1e−05
0e+00
density
0
10000
20000
30000
40000
50000
NeNAfb_NAfa_ME_EU_BEi_EAs_WAf_Khs_Xa
distribution
posterior
prior

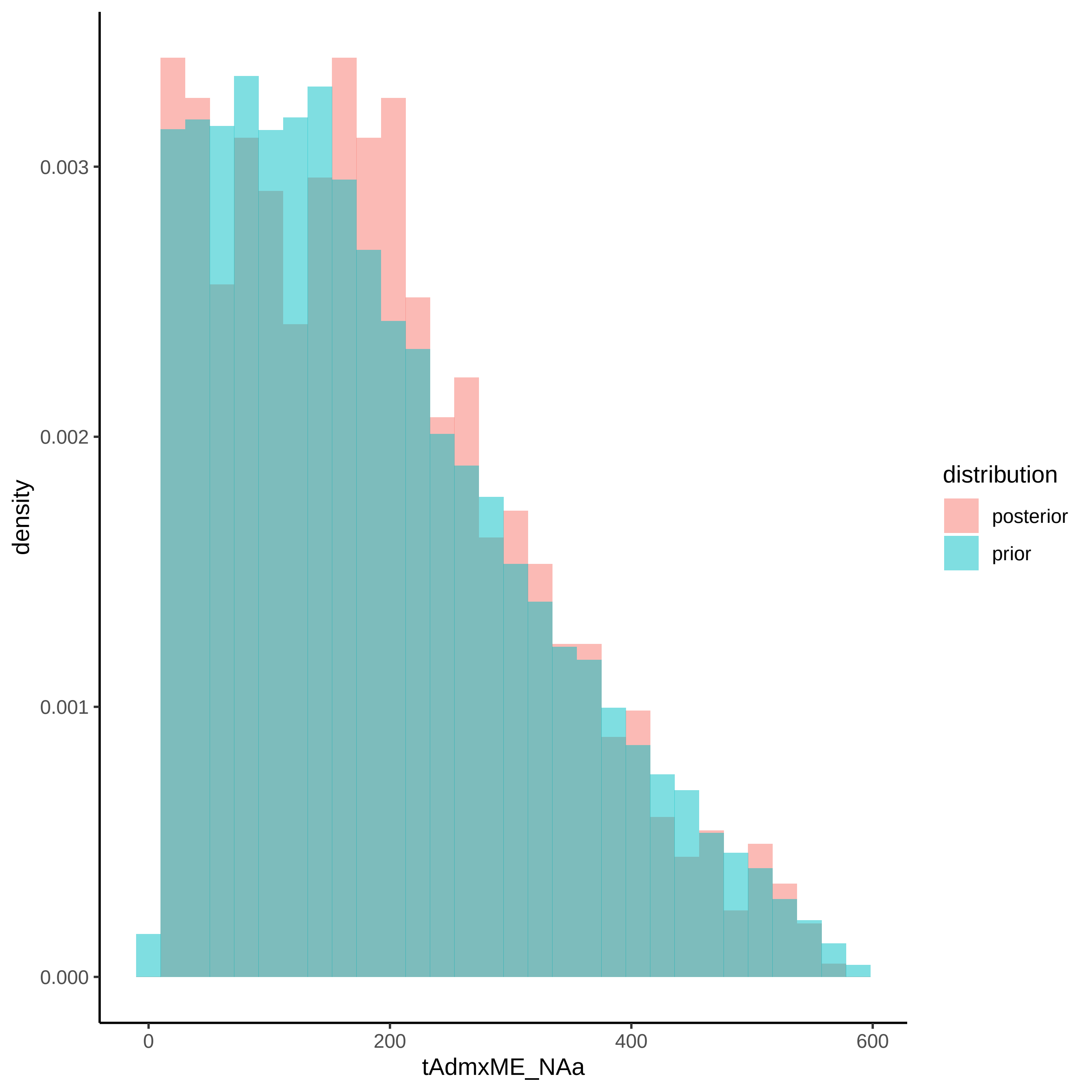
0.003
0.002
0.001
0.000
density
0
200
400
600
tAdmxME_NAa
distribution
posterior
prior

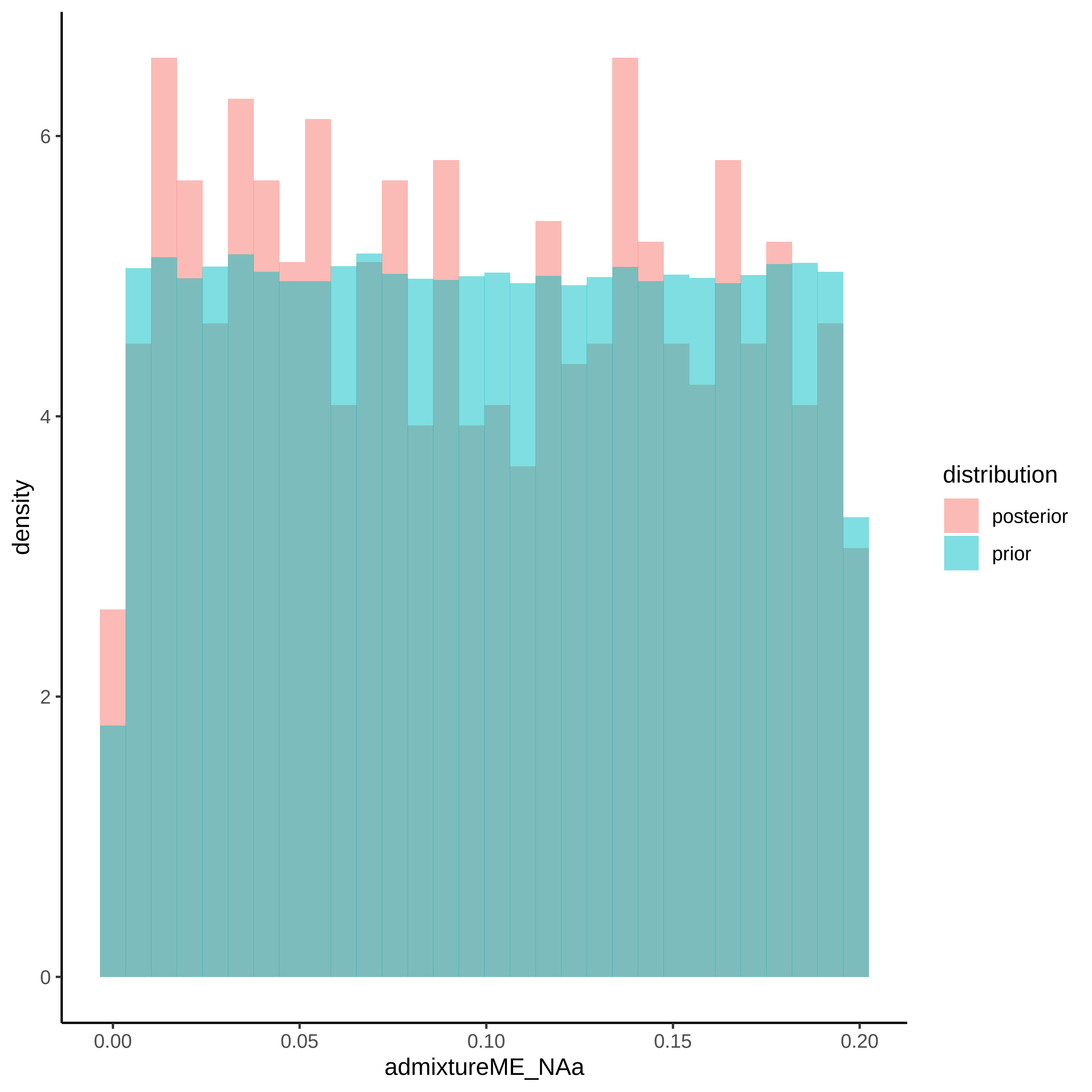

density
admixtureME_NAa
distribution
posterior
prior
0
2
4
6
0.00
0.05
0.10
0.15
0.20

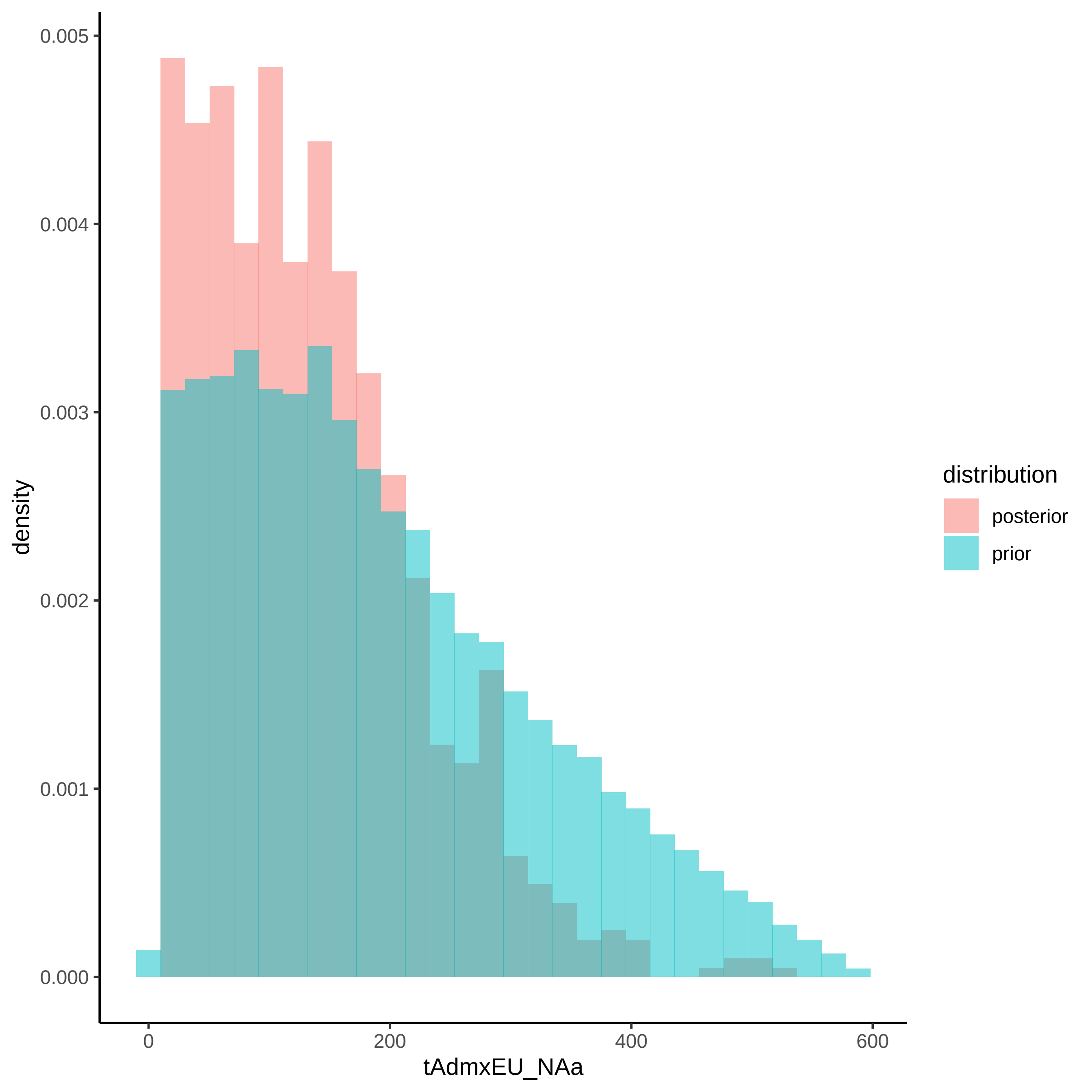

0.005
0.004
0.003
0.002
0.001
0.000
density
0
200
400
600
tAdmxEU_NAa
distribution
posterior
prior

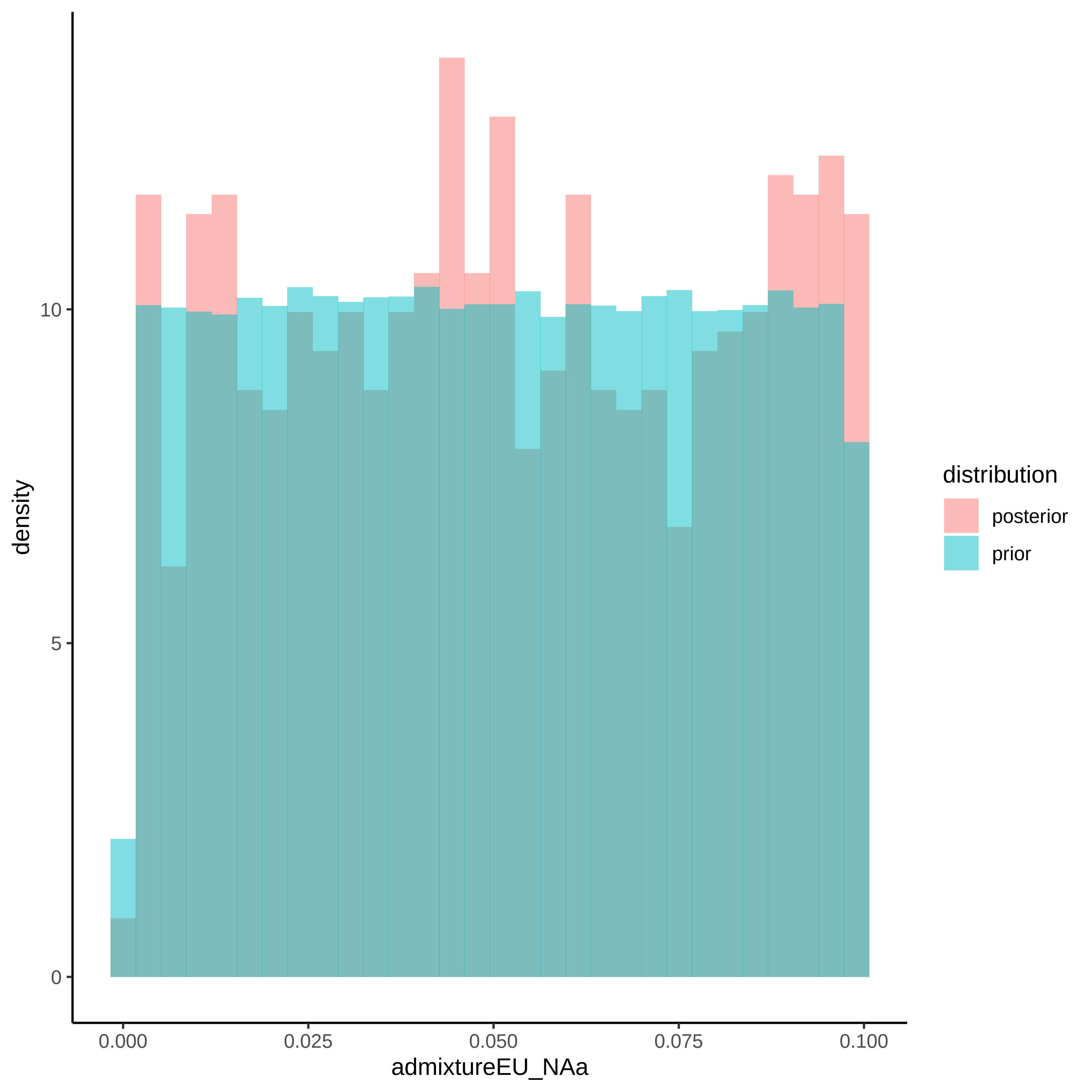
density
10
5
0
0.000
0.025
0.050
0.075
0.100
admixtureEU_NAa
distribution
posterior
prior

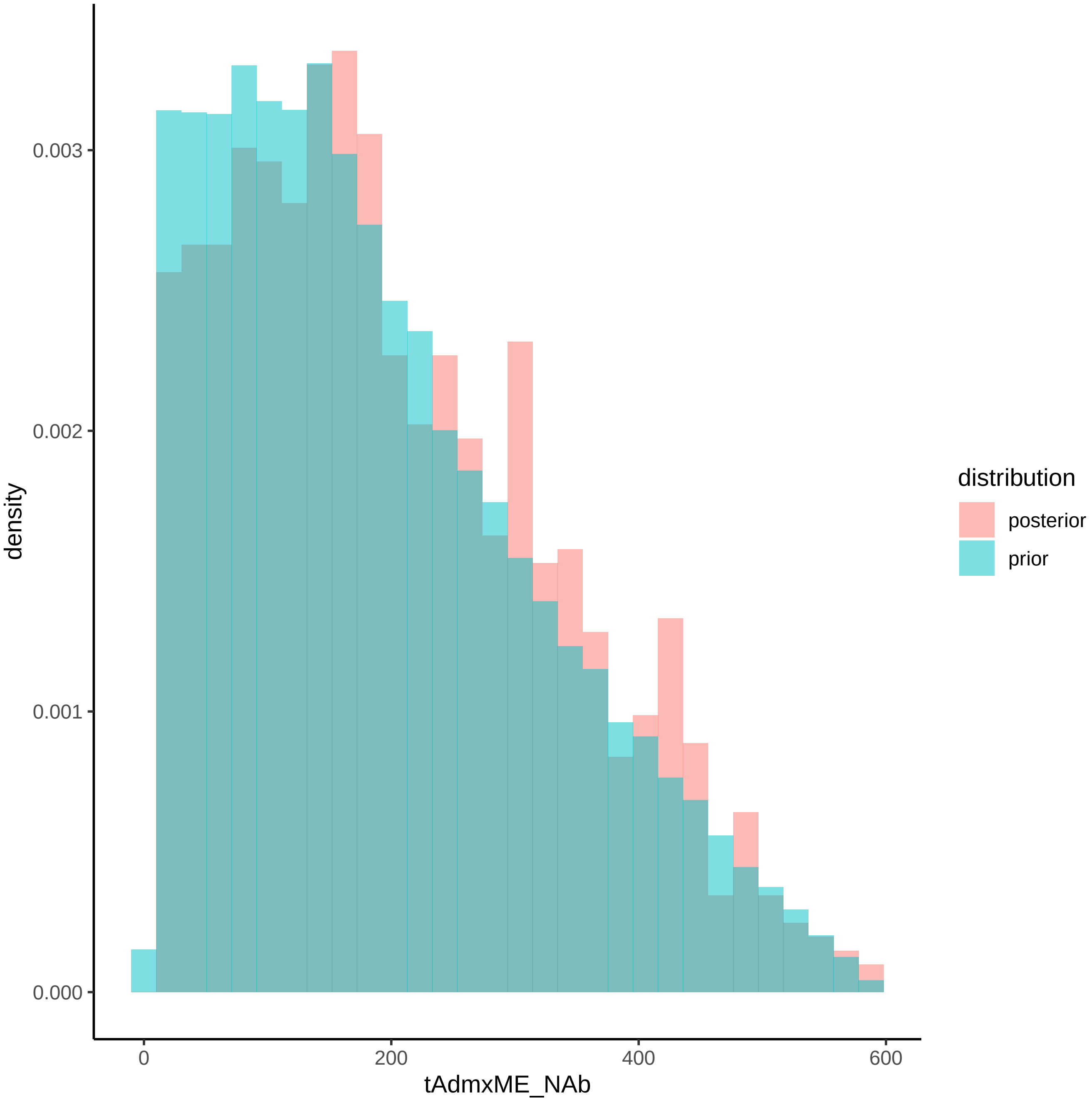

density
0.000
0.001
0.002
0.003
0
200
400
600
tAdmxME_NAb
distribution
posterior
prior

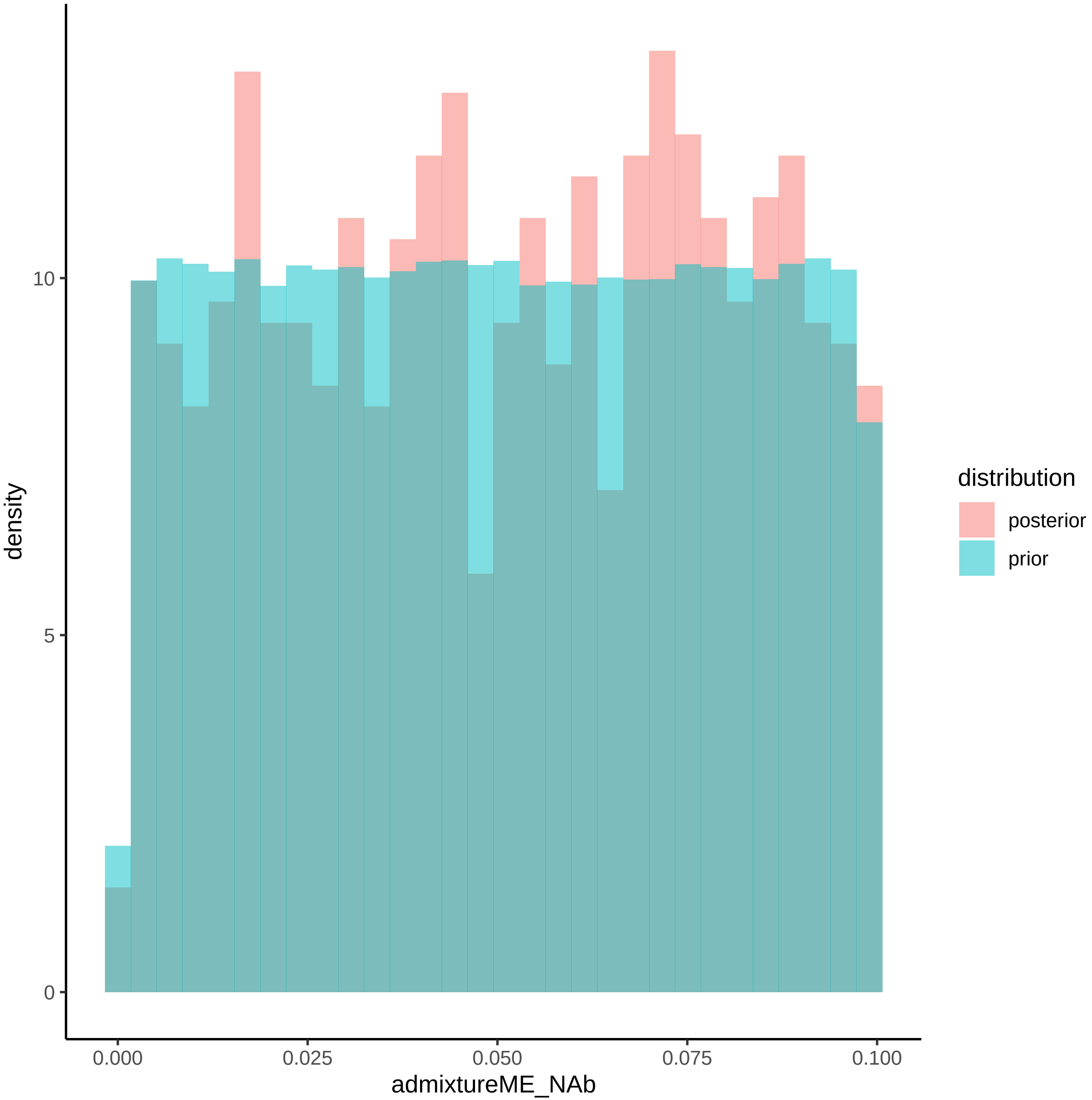

density
10
5
0
0.000
0.025
0.050
0.075
0.100
admixtureME_NAb
distribution
posterior
prior

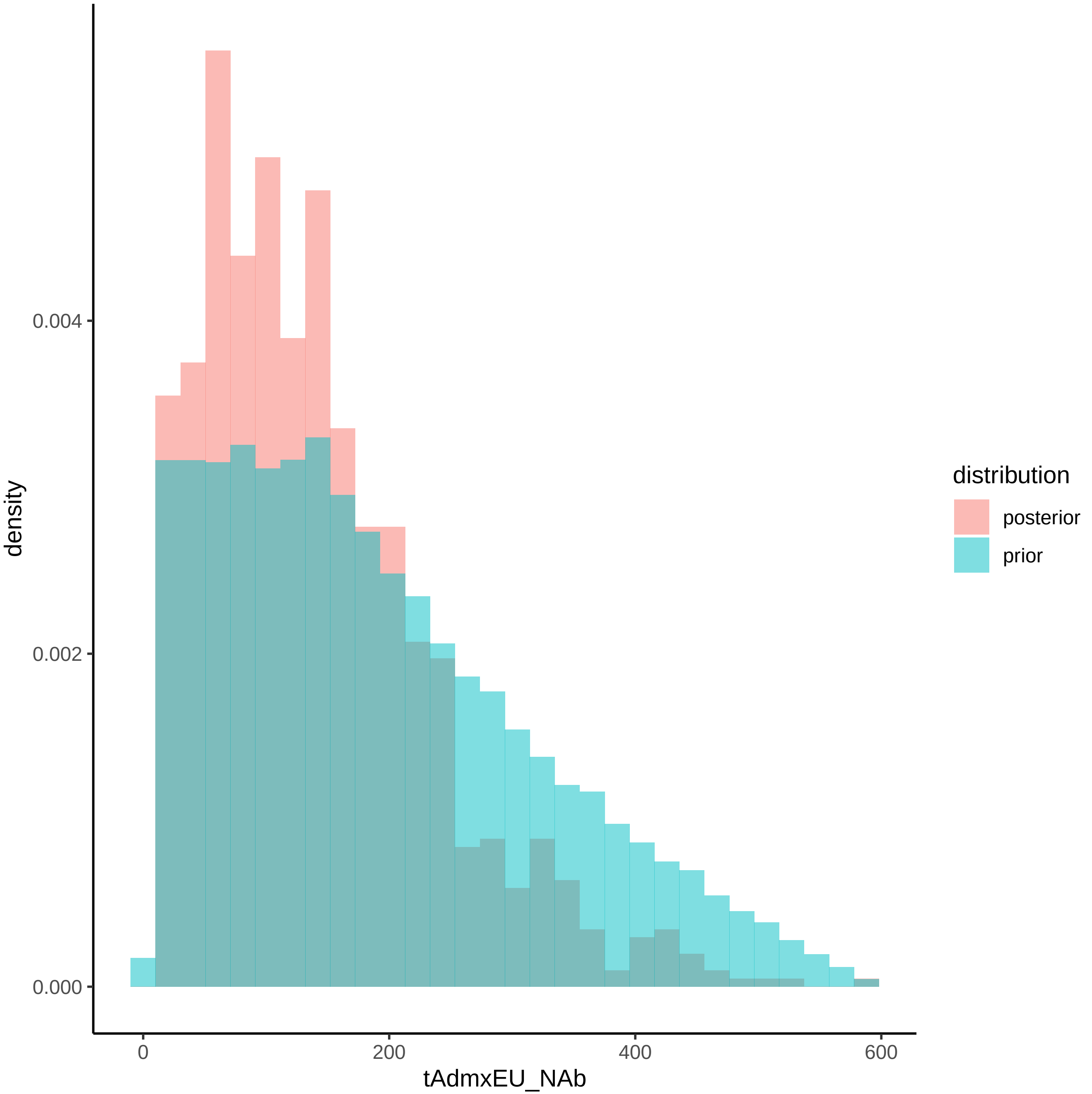

0.004
0.002
0.000
density
0
200
400
600
tAdmxEU_NAb
distribution
posterior
prior

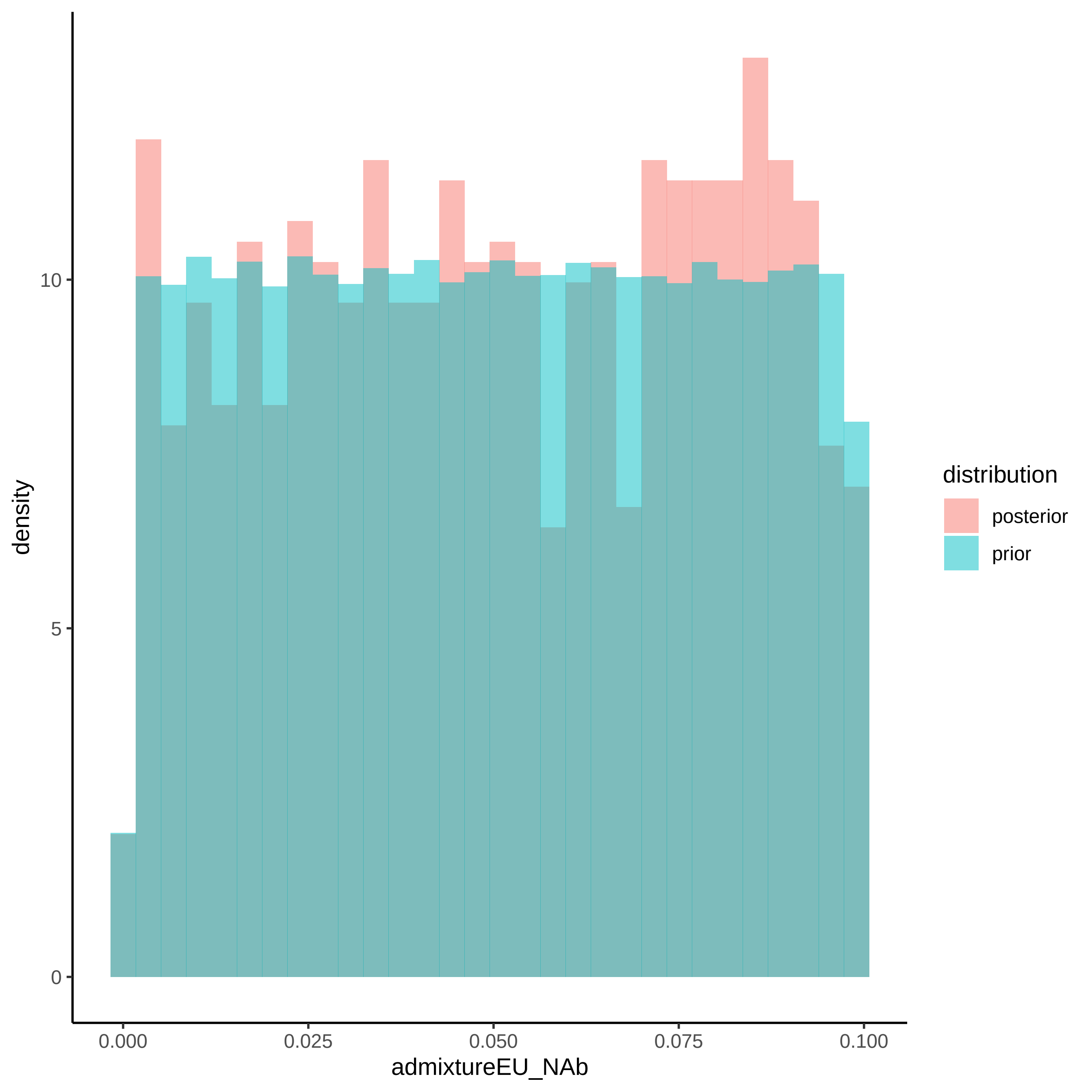

density
10
5
0
0.000
0.025
0.050
0.075
0.100
admixtureEU_NAb
distribution
posterior
prior

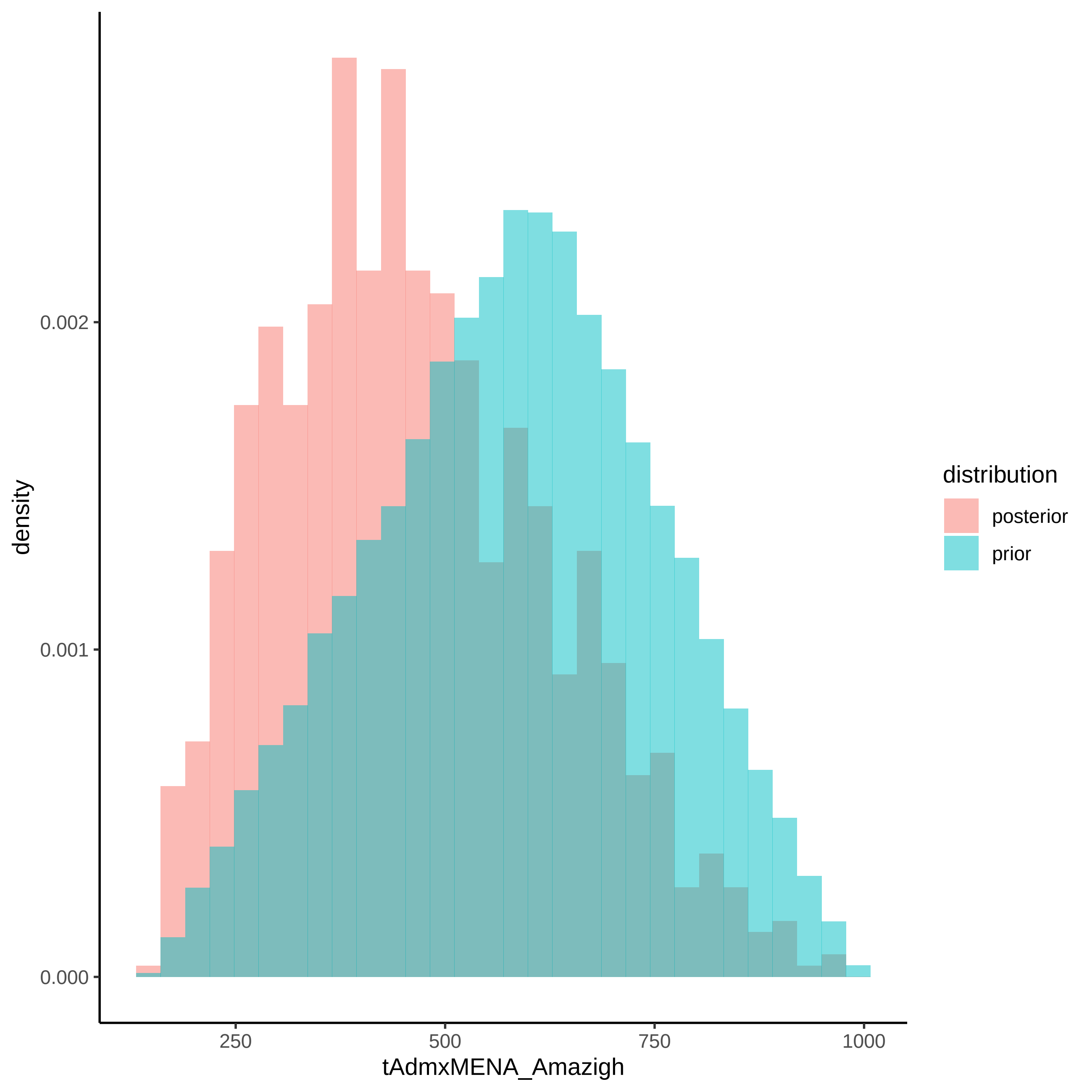

density
0.000
0.001
0.002
250
500
750
1000
tAdmxMENA_Amazigh
distribution
posterior
prior

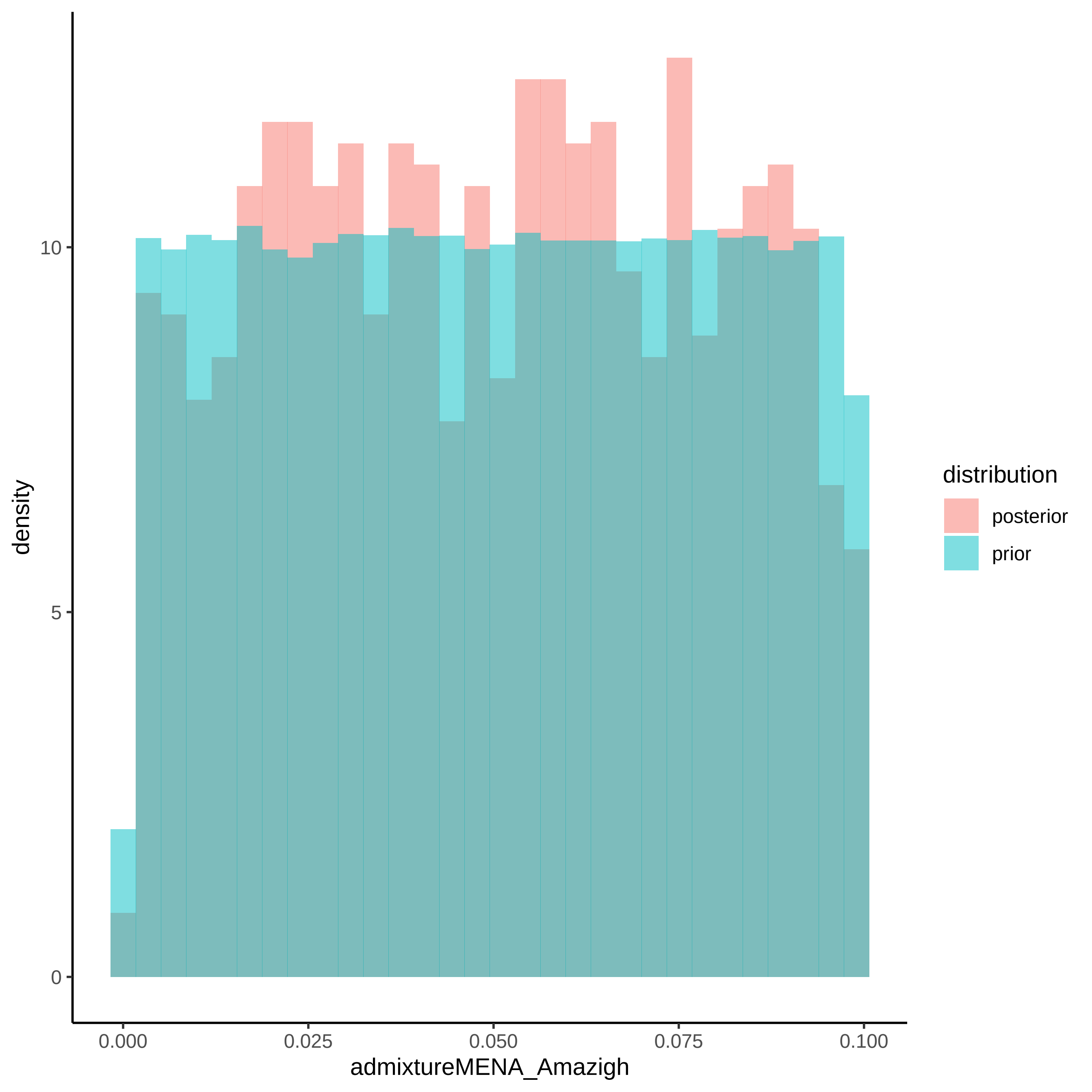
density
10
5
0
0.000
0.025
0.050
0.075
0.100
admixtureMENA_Amazigh
distribution
posterior
prior

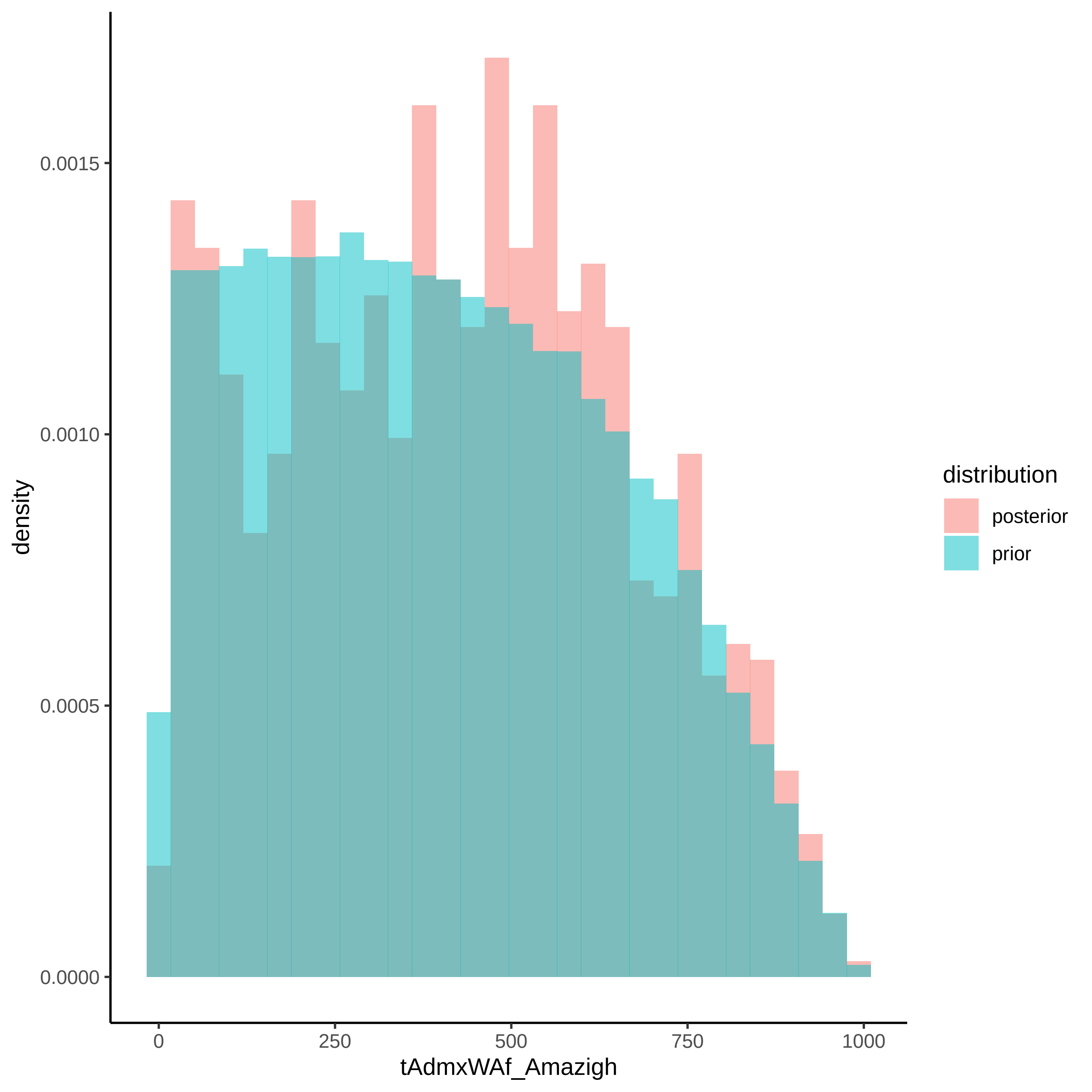

density
0.0015
0.0010
0.0005
0.0000
0
250
500
750
1000
tAdmxWAf_Amazigh
distribution
posterior
prior

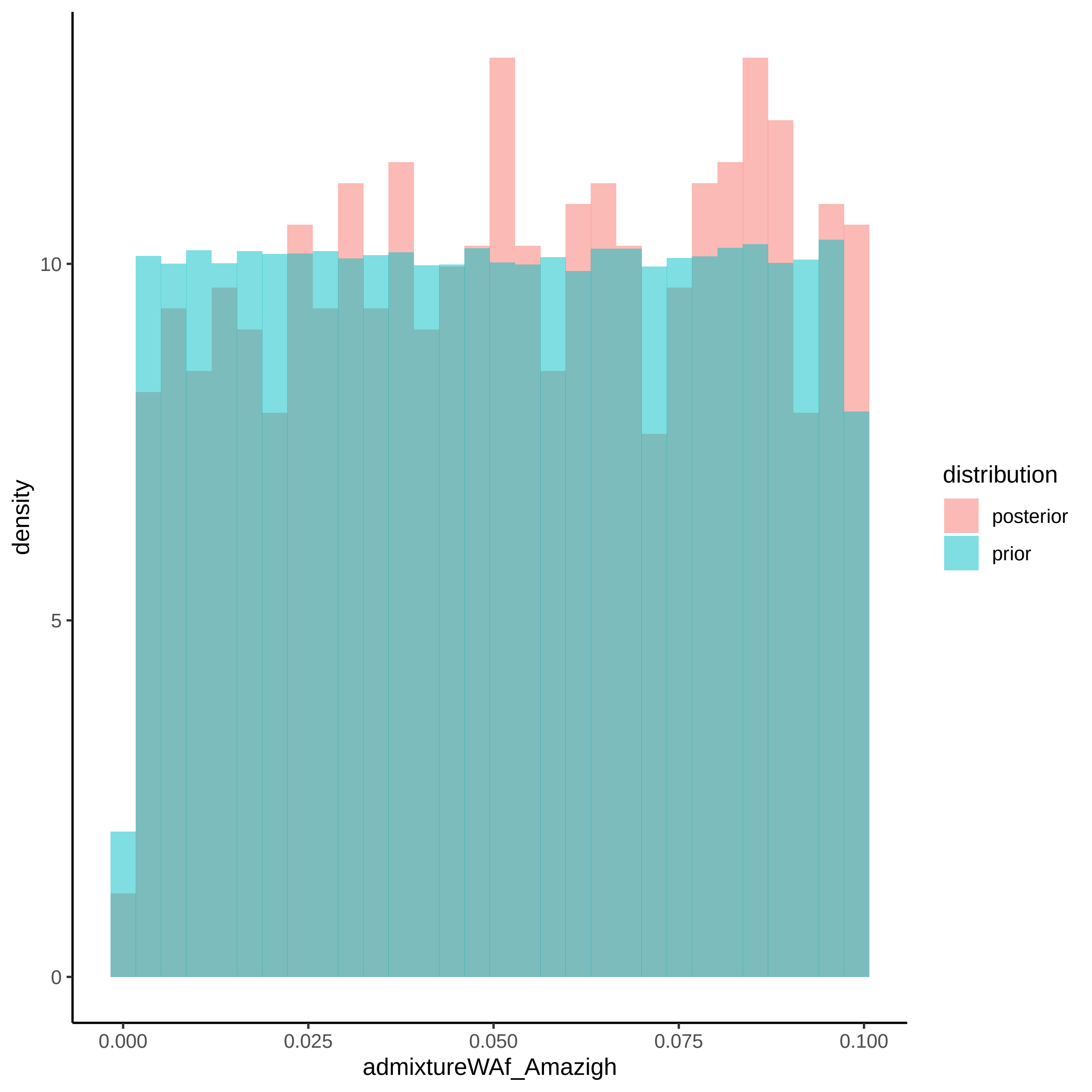

density
10
5
0
0.000
0.025
0.050
0.075
0.100
admixtureWAf_Amazigh
distribution
posterior
prior

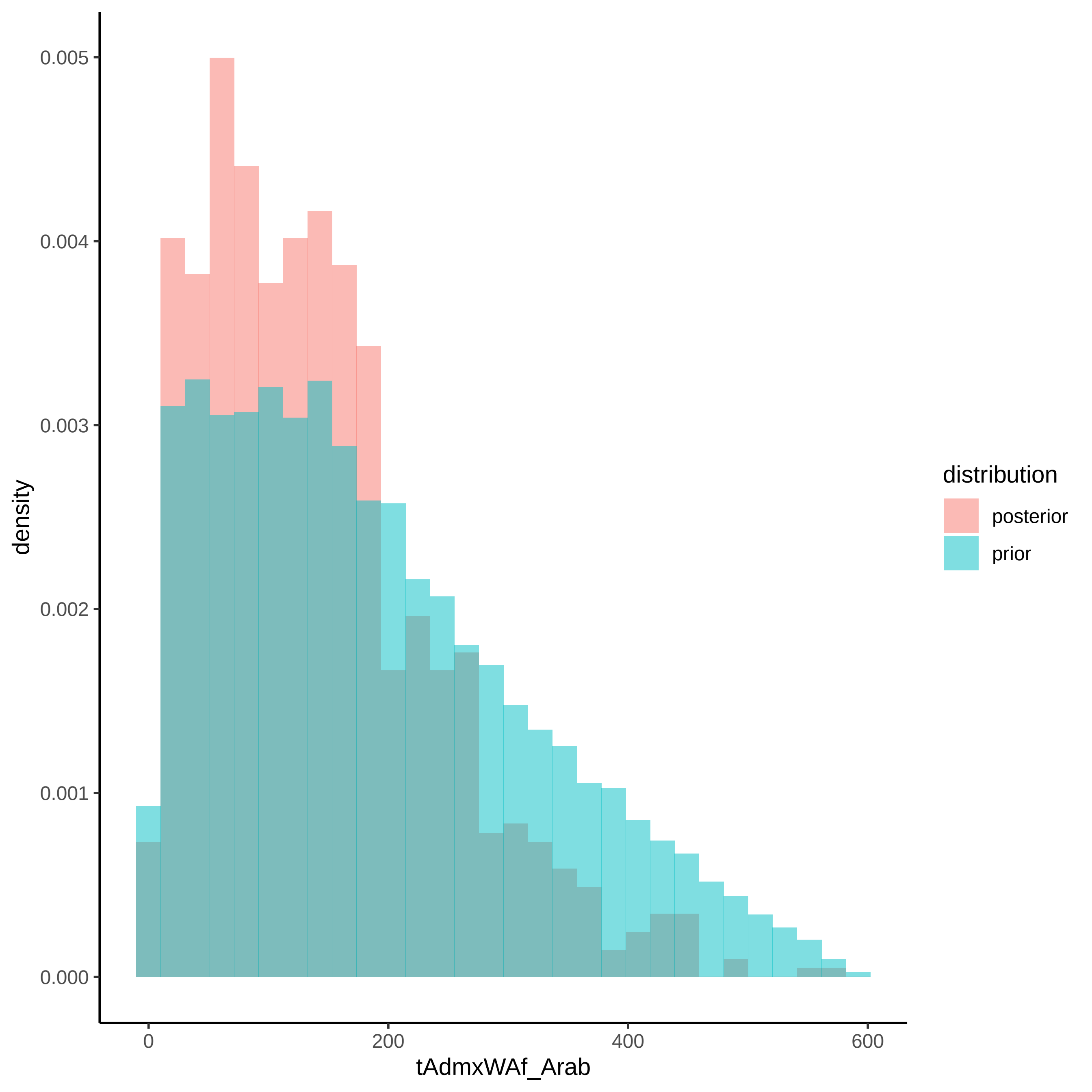

density
0.000
0.001
0.002
0.003
0.004
0.005
0
200
400
600
tAdmxWAf_Arab
distribution
posterior
prior

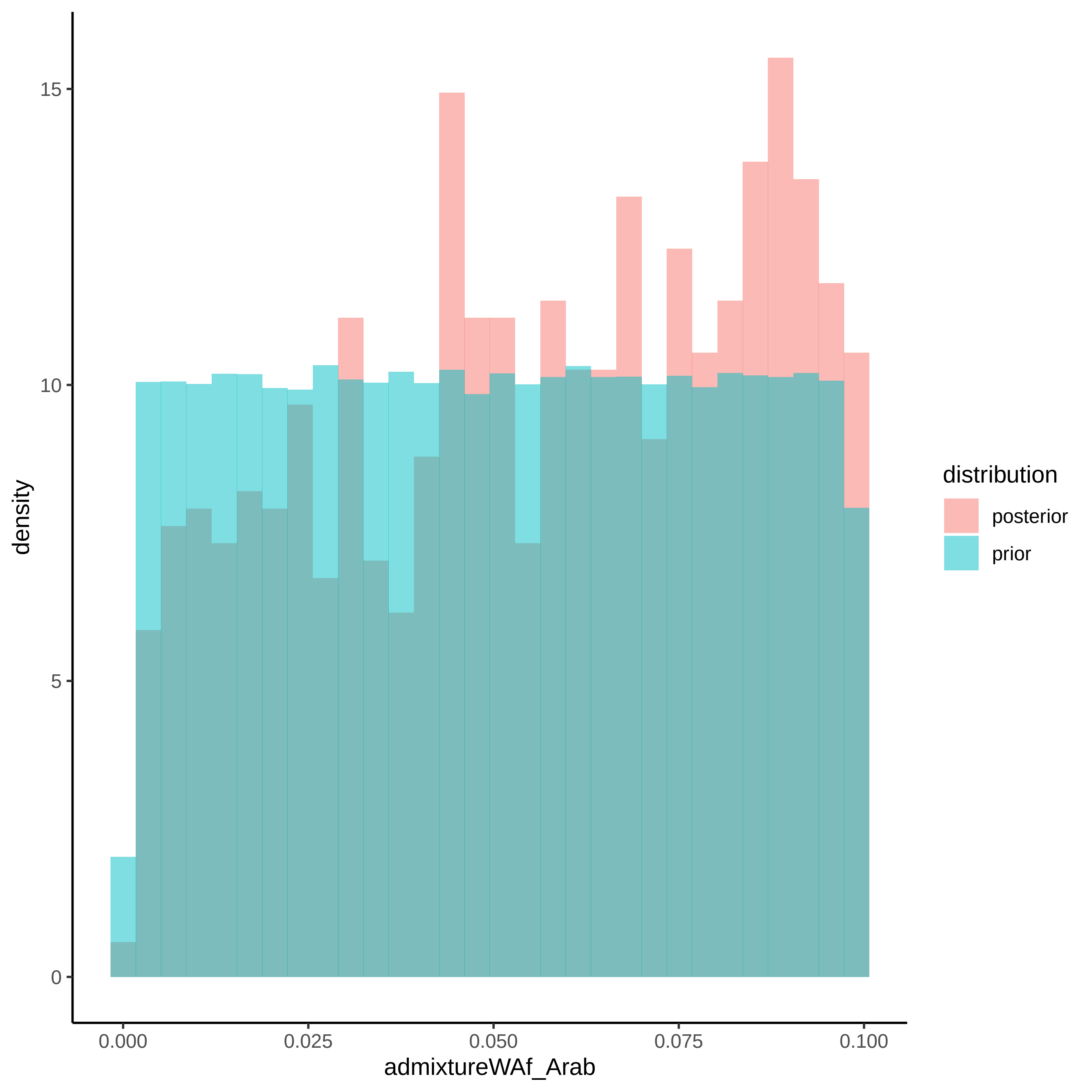

15
10
5
0
density
0.000
0.025
0.050
0.075
0.100
admixtureWAf_Arab
distribution
posterior
prior

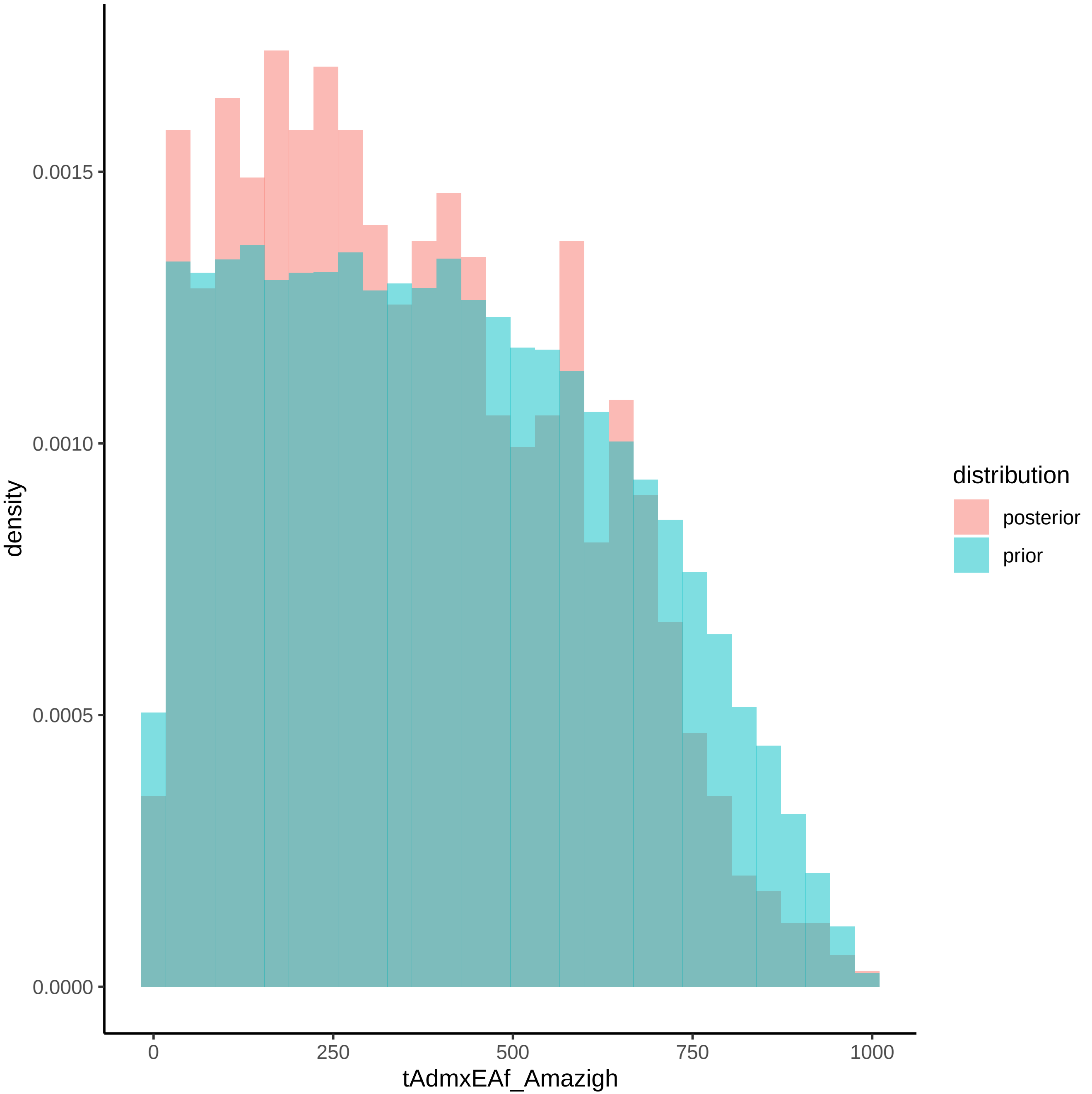

density
0.0000
0.0005
0.0010
0.0015
0
250
500
750
1000
tAdmxEAf_Amazigh
distribution
posterior
prior

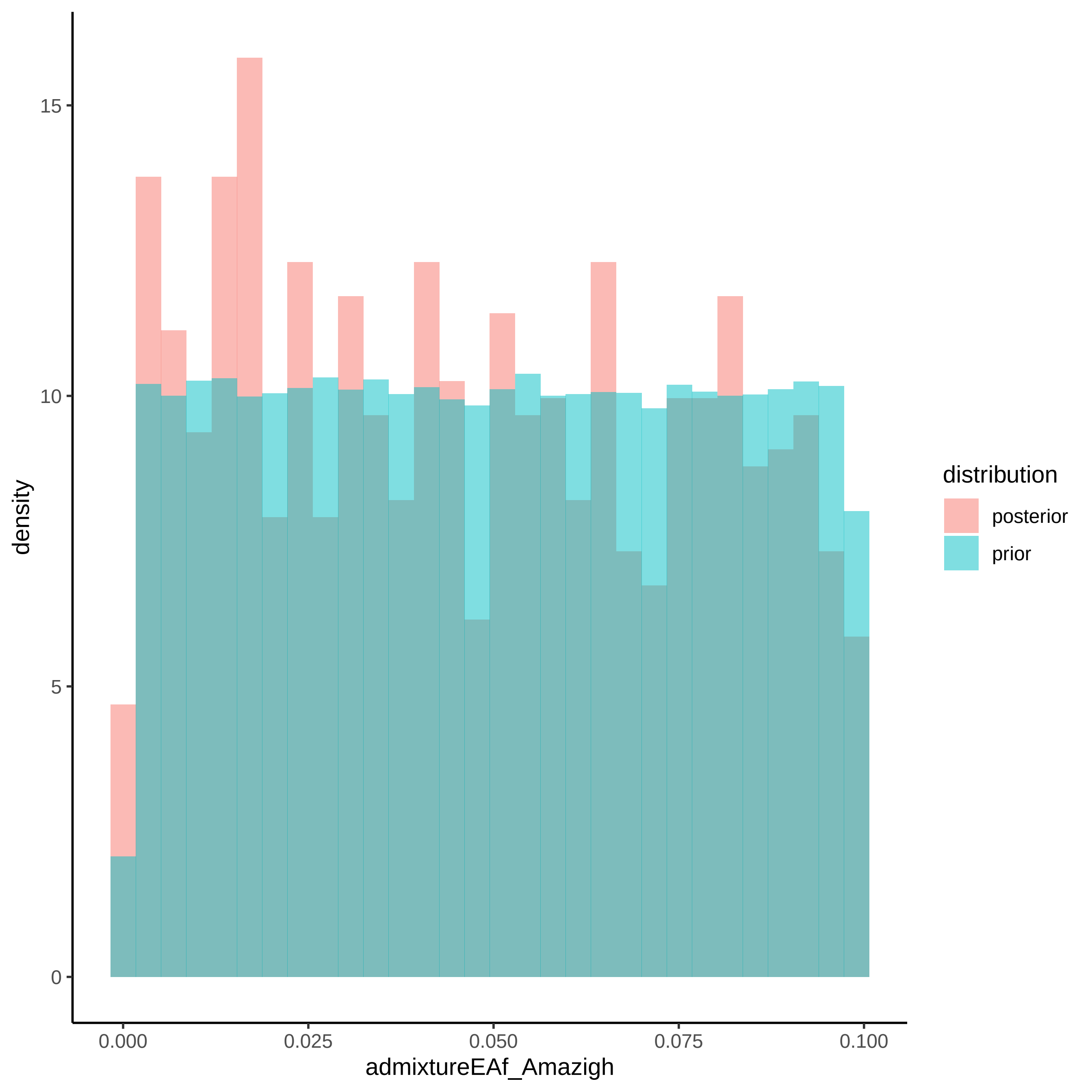

density
15
10
5
0
0.000
0.025
0.050
0.075
0.100
admixtureEAf_Amazigh
distribution
posterior
prior

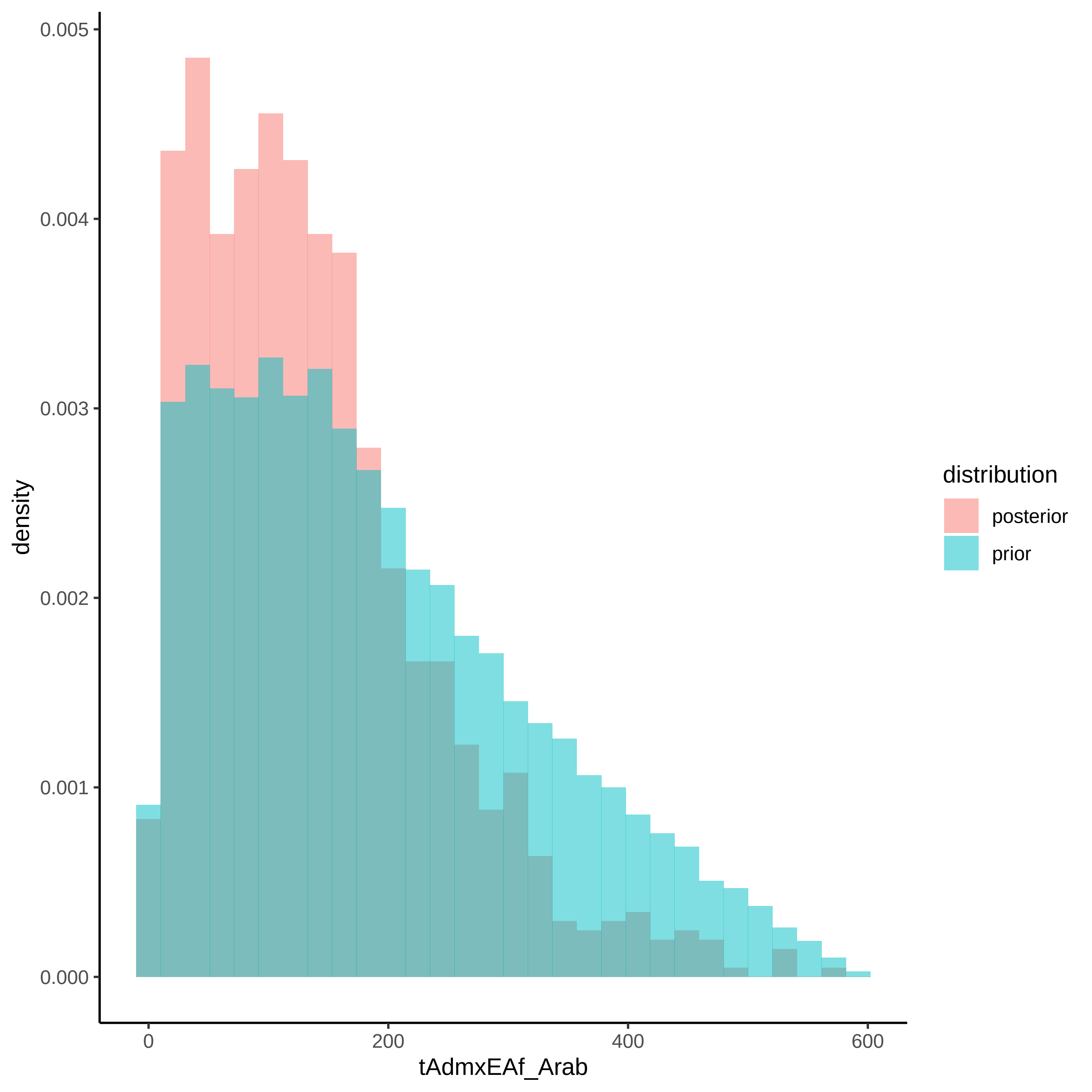

density
0.005
0.004
0.003
0.002
0.001
0.000
0
200
400
600
tAdmxEAf_Arab
distribution
posterior
prior

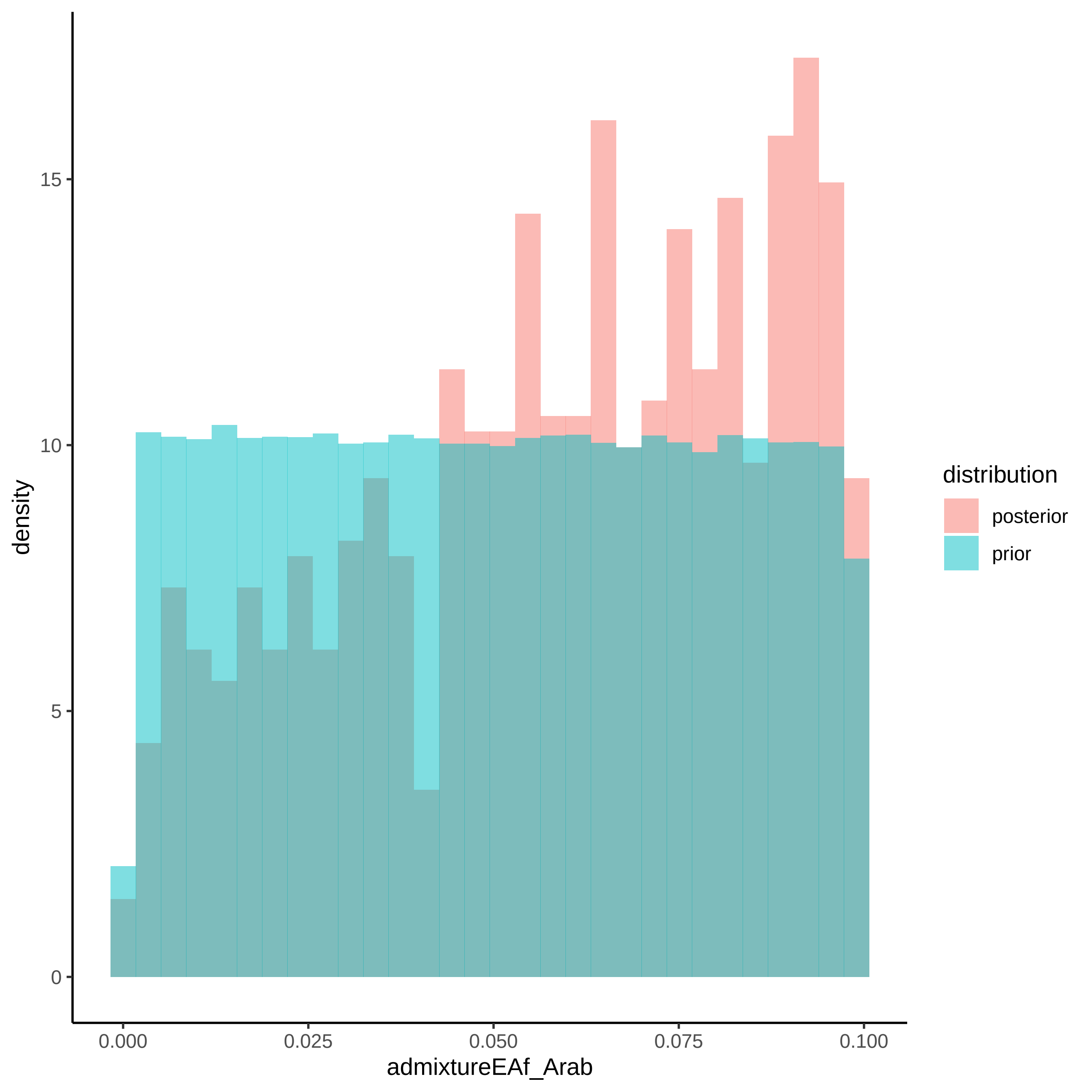

density
15
10
5
0
0.000
0.025
0.050
0.075
0.100
admixtureEAf_Arab
distribution
posterior
prior

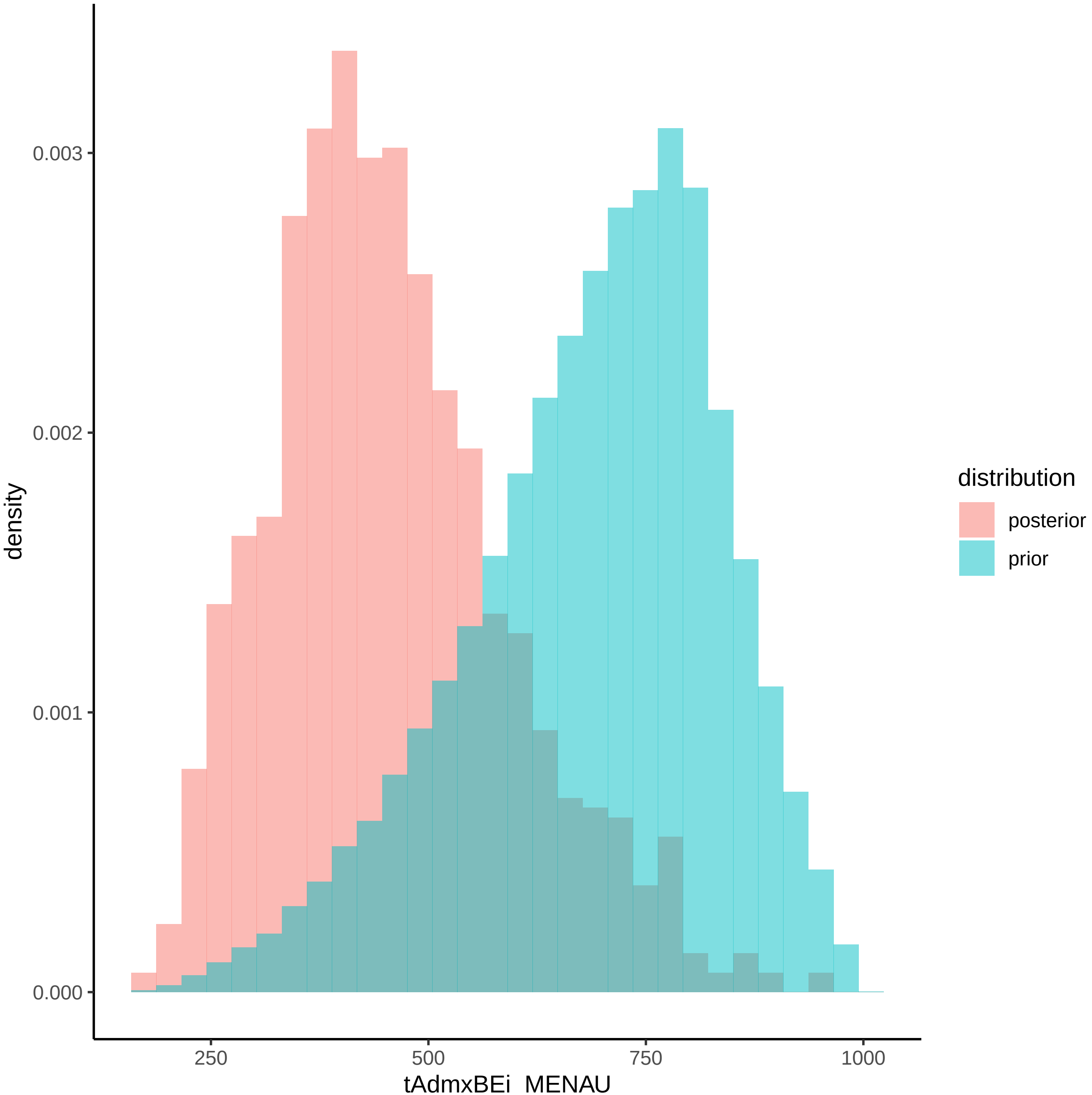
density
0.003
0.002
0.001
0.000
250
500
750
1000
tAdmxBEi_MENAU
distribution
posterior
prior

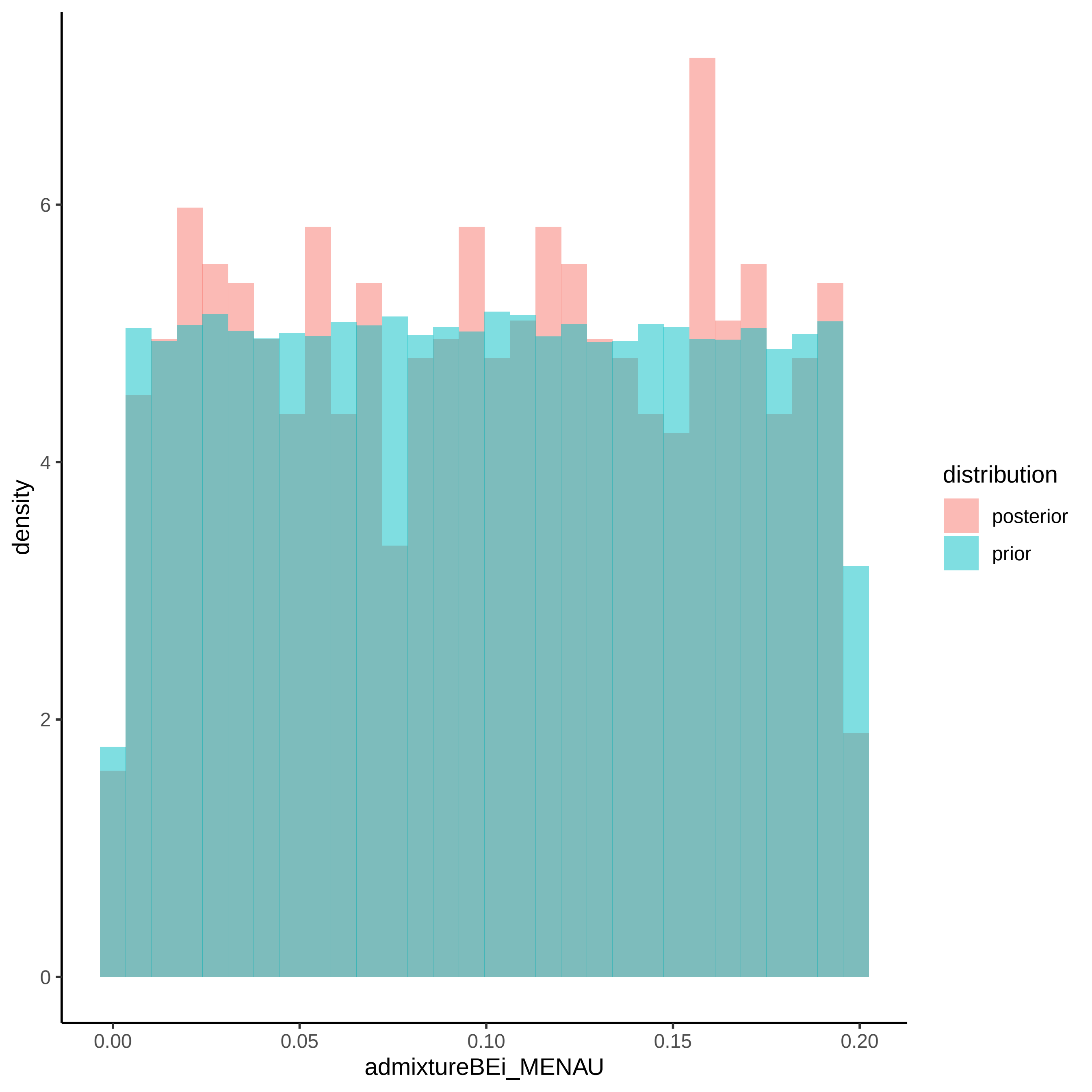

density
6
4
2
0
0.00
0.05
0.10
0.15
0.20
admixtureBEi_MENAU
distribution
posterior
prior

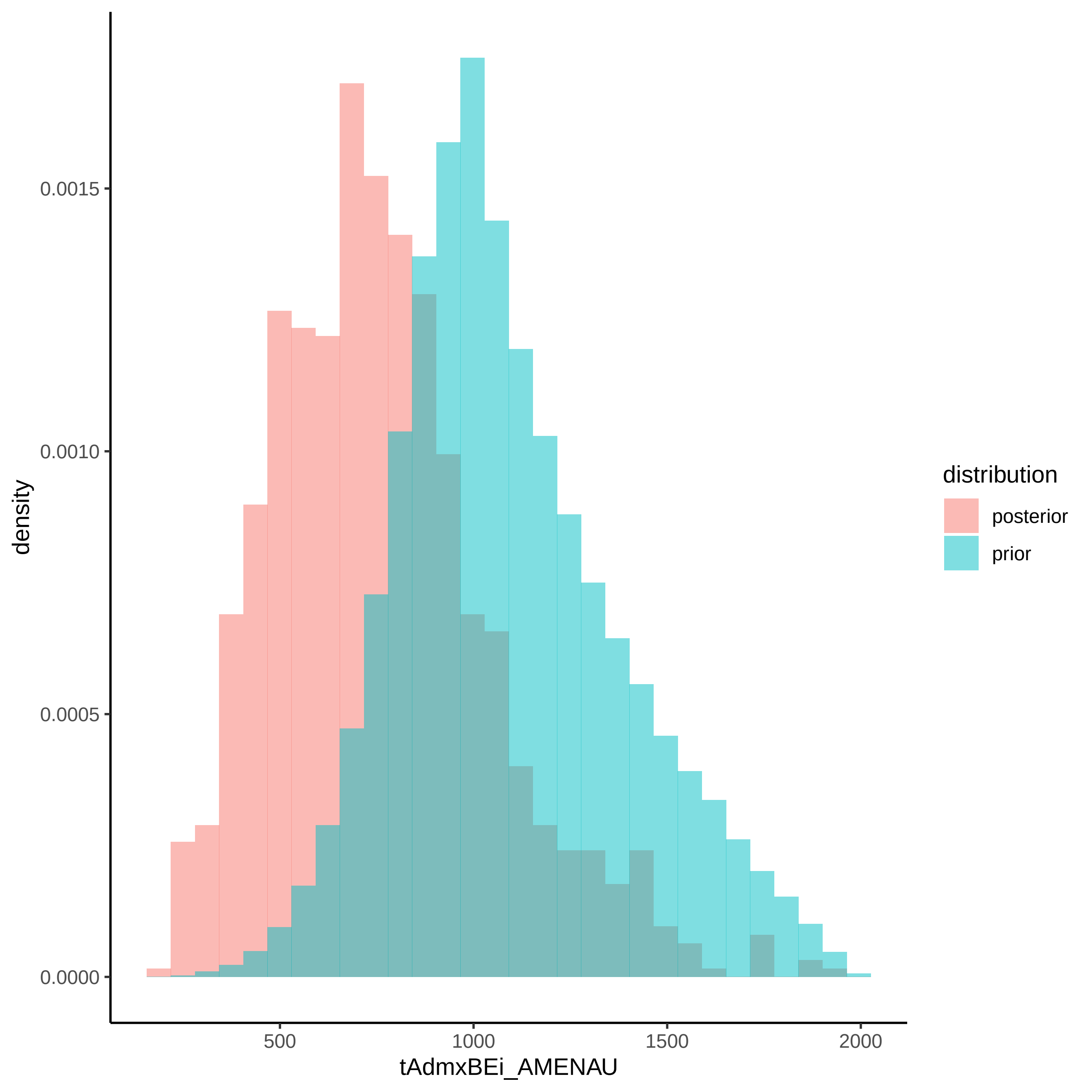

density
0.0015
0.0010
0.0005
0.0000
500
1000
1500
2000
tAdmxBEi_AMENAU
distribution
posterior
prior

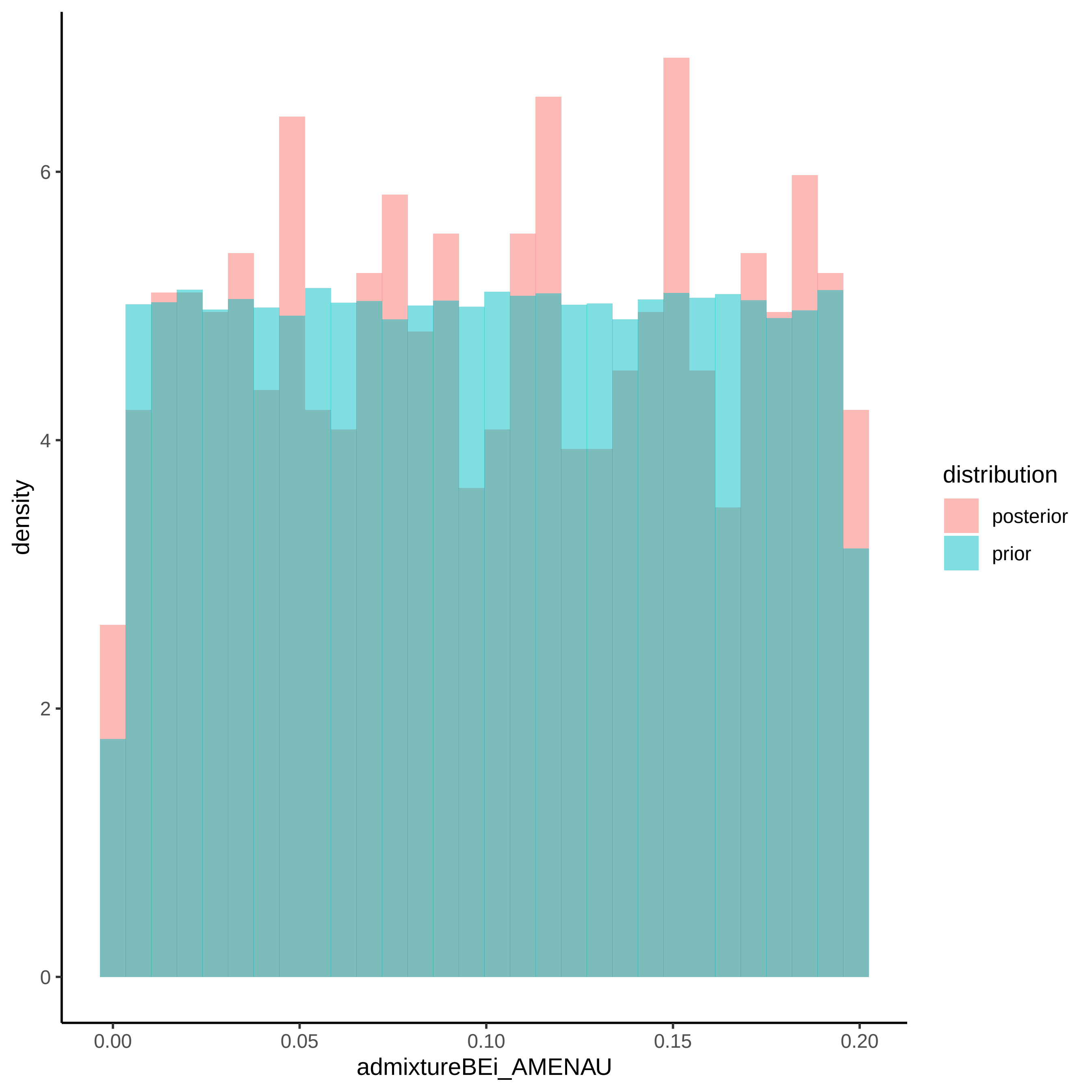
density
admixtureBEi_AMENAU
distribution
posterior
prior
0
2
4
6
0.00
0.05
0.10
0.15
0.20

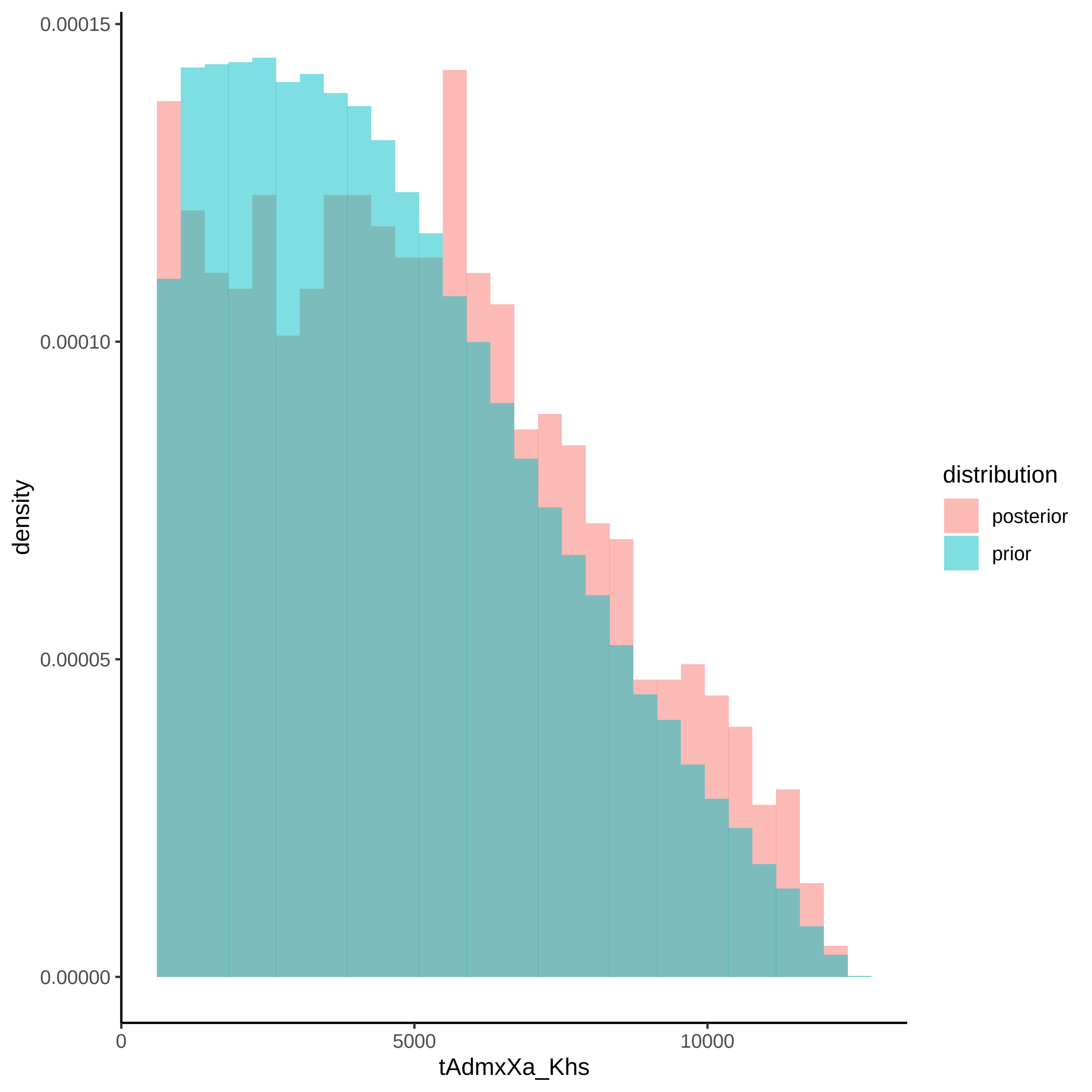

0.00015
0.00010
0.00005
0.00000
density
0
5000
10000
tAdmxXa_Khs
distribution
posterior
prior

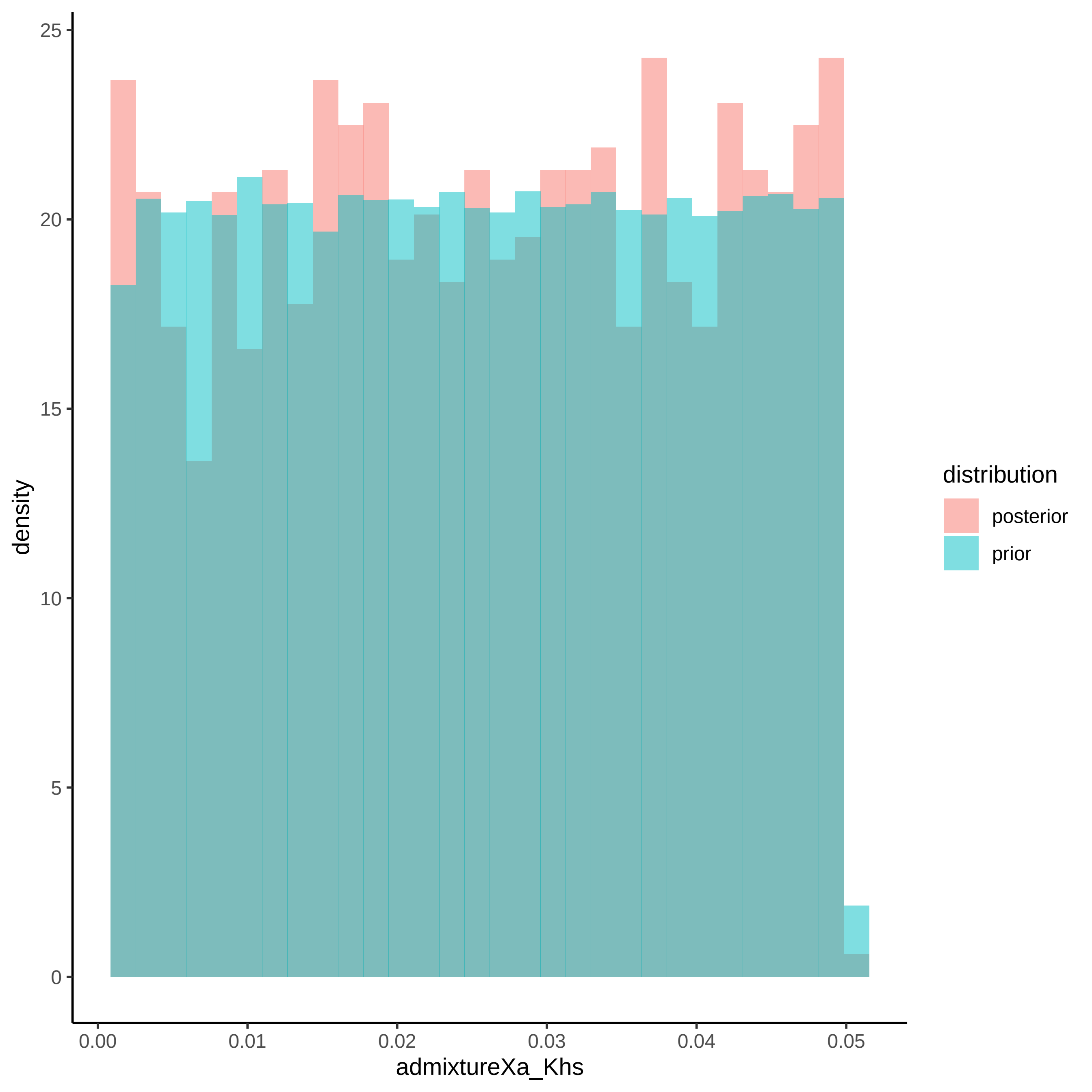

density
admixtureXa_Khs
0
5
10
15
20
25
0.00
0.01
0.02
0.03
0.04
0.05
distribution
posterior
prior

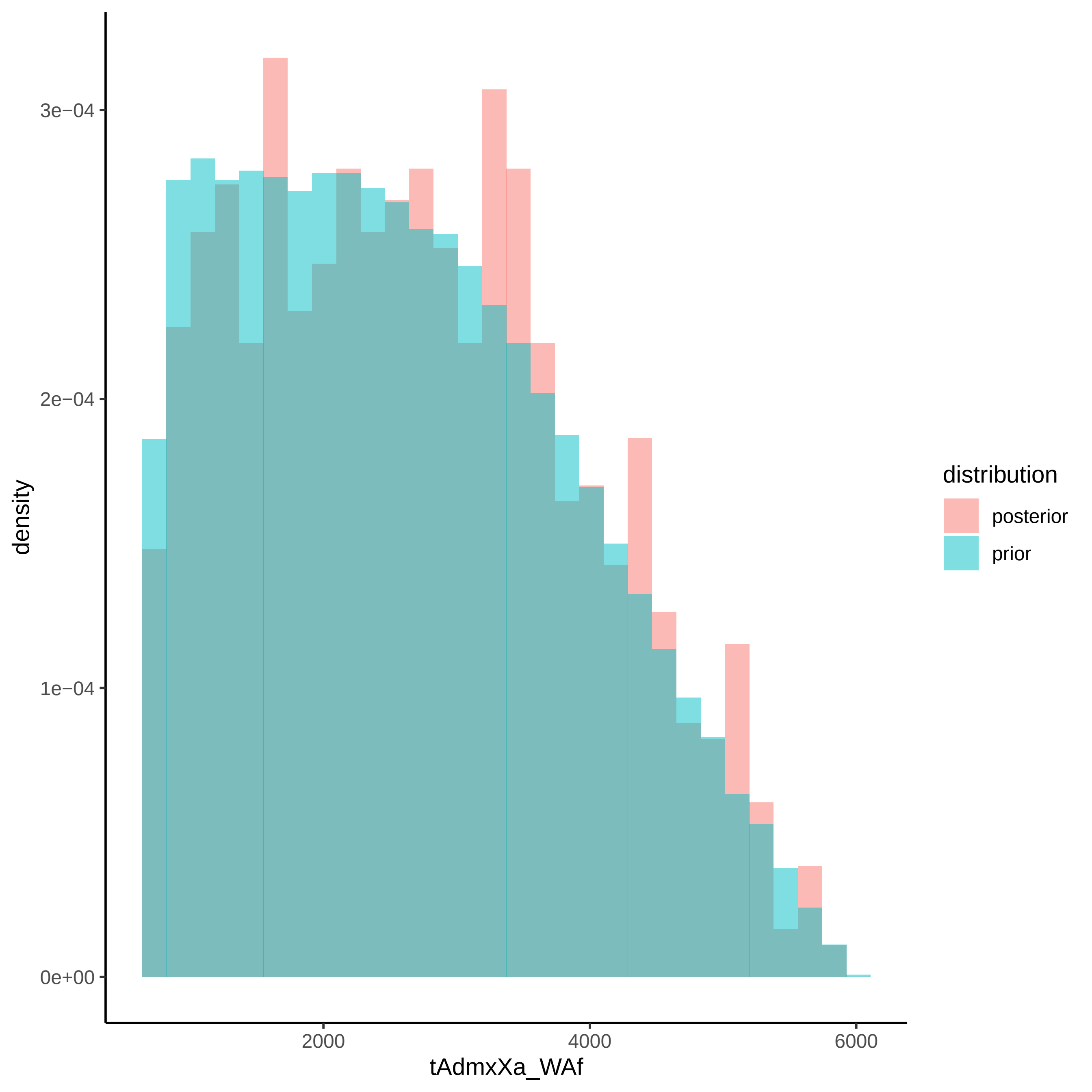

3e−04
2e−04
1e−04
0e+00
density
2000
4000
6000
tAdmxXa_WAf
distribution
posterior
prior

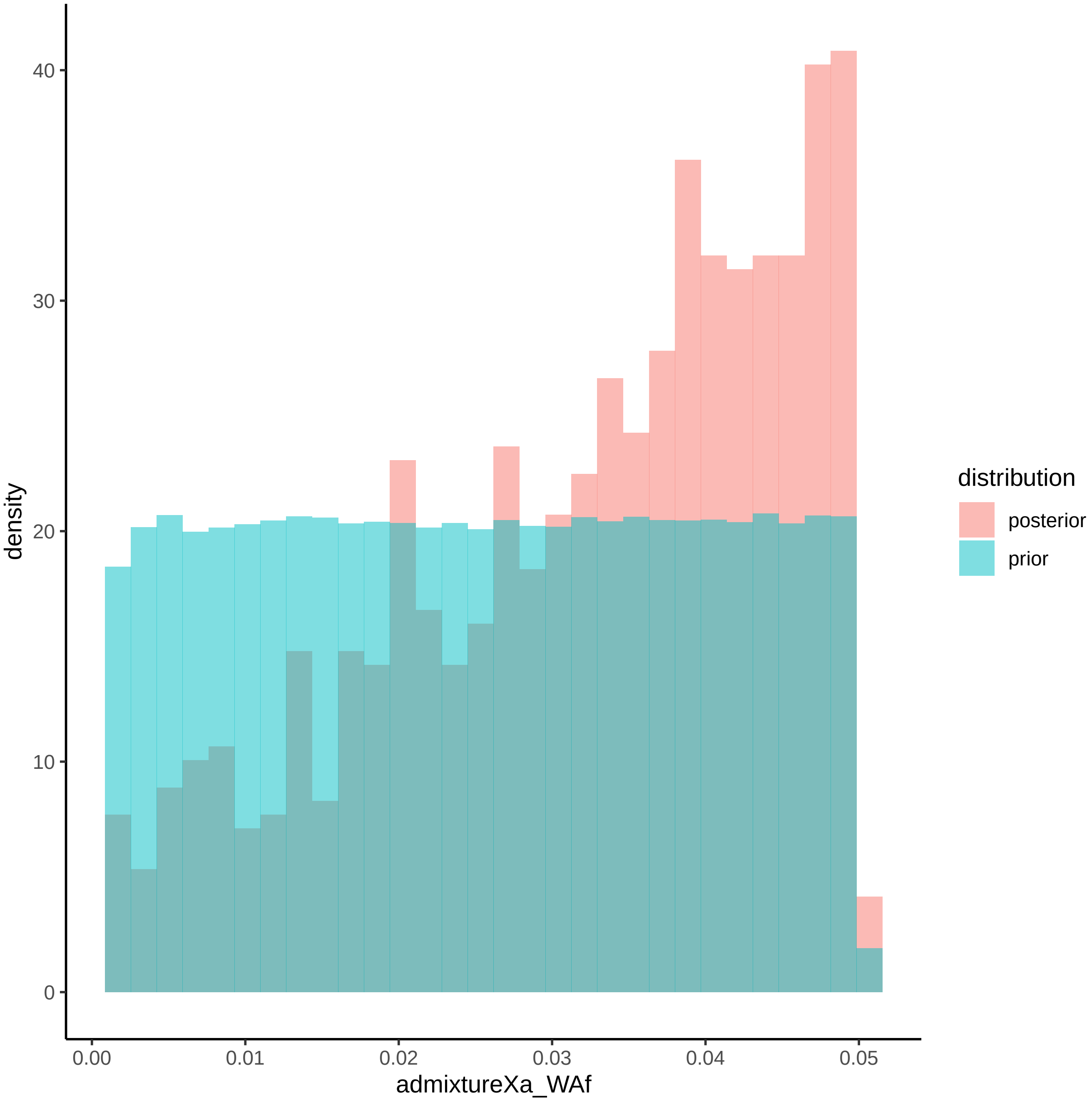

density
40
30
20
10
0
0.00
0.01
0.02
0.03
0.04
0.05
admixtureXa_WAf
distribution
posterior
prior
