## Supplementary File 4 for "Modelling the demographic history of human North African genomes points to soft split divergence between populations"

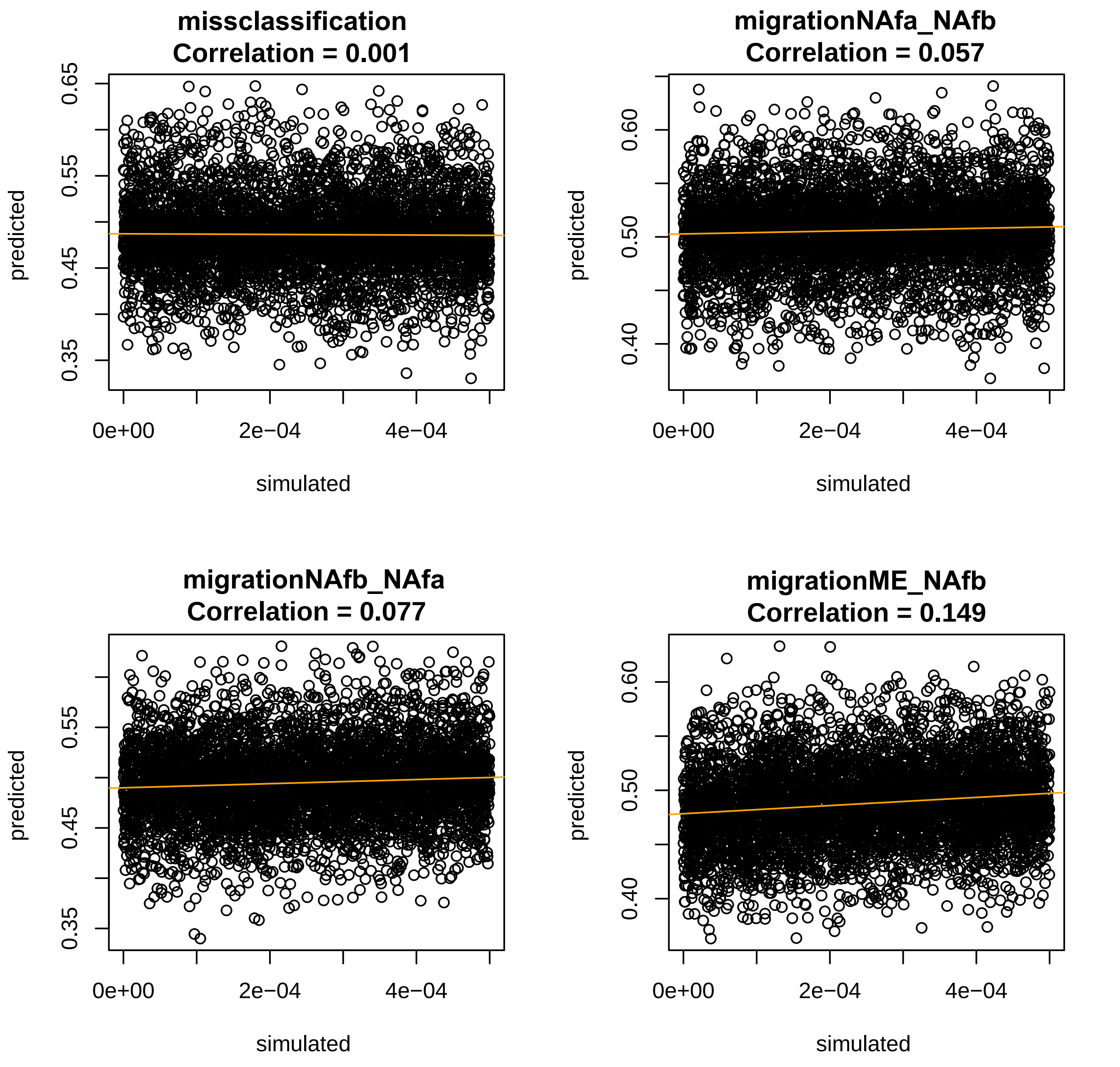
missclassification
Correlation = 0.001
migrationNAfa_NAfb
Correlation = 0.057
migrationNAfb_NAfa
Correlation = 0.077
migrationME_NAfb
Correlation = 0.149
predicted
simulated
0.65
0.55
0.45
0.35
0.60
0.50
0.40
0e+00
2e−04
4e−04

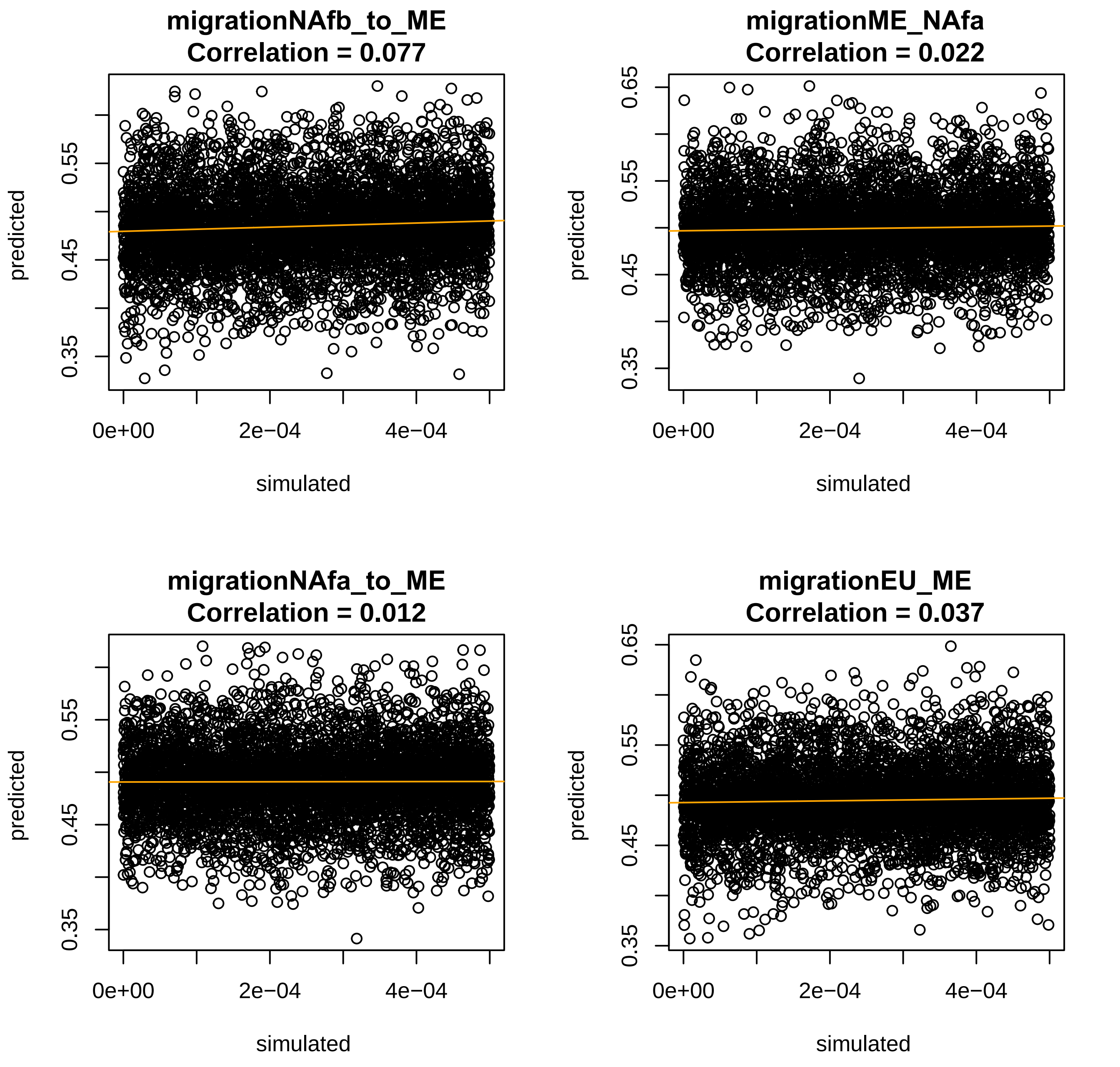
migrationNAfb_to_ME
Correlation = 0.077
predicted
simulated
0.35
0.45
0.55
0e+00
2e−04
4e−04
migrationME_NAfa
Correlation = 0.022
predicted
simulated
0.35
0.45
0.55
0.65
0e+00
2e−04
4e−04
migrationNAfa_to_ME
Correlation = 0.012
predicted
simulated
0.35
0.45
0.55
0e+00
2e−04
4e−04
migrationEU_ME
Correlation = 0.037
predicted
simulated
0.35
0.45
0.55
0.65
0e+00
2e−04
4e−04

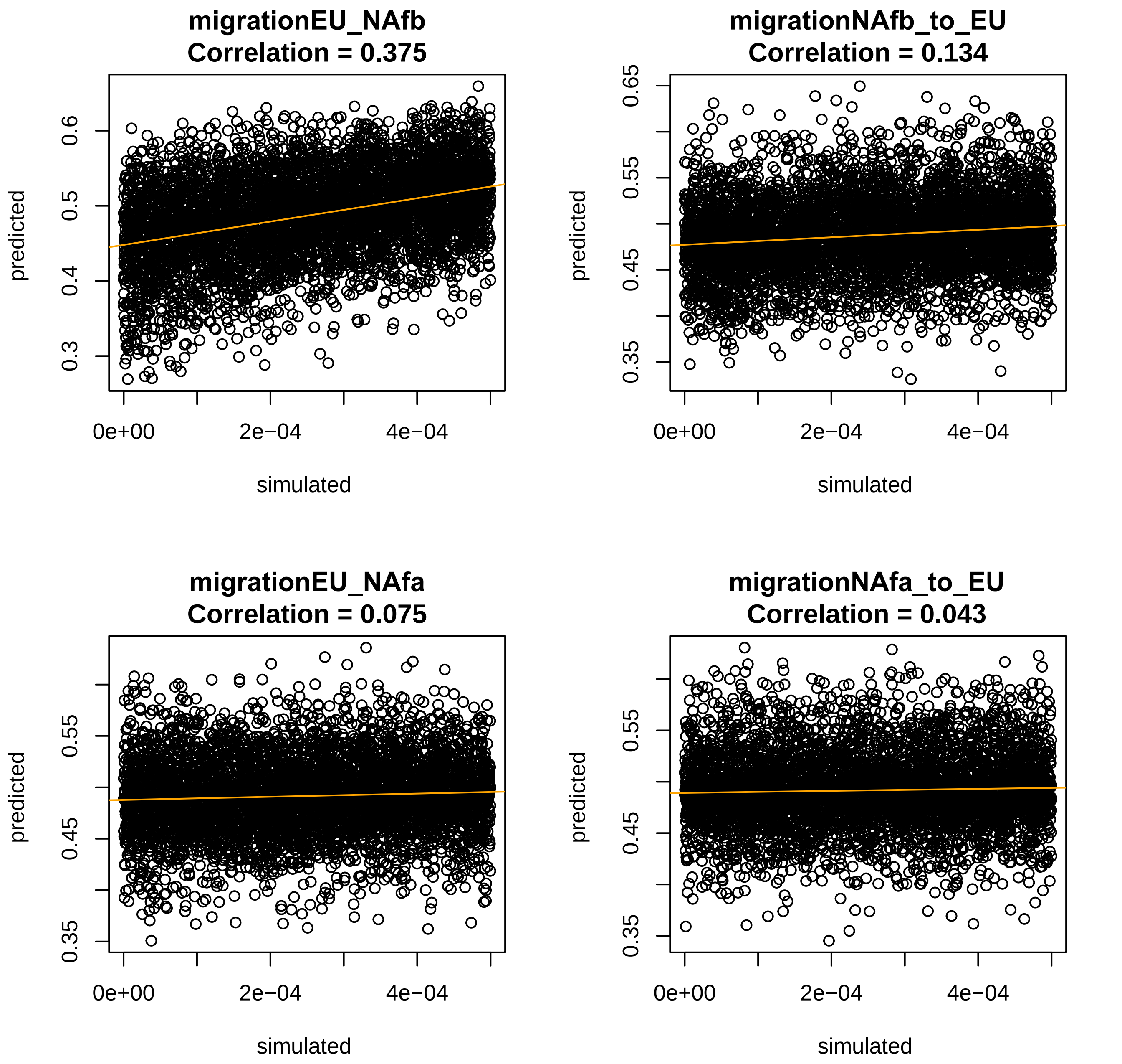

migrationEU_NAfb
Correlation = 0.375
migrationNAfb_to_EU
Correlation = 0.134
migrationEU_NAfa
Correlation = 0.075
migrationNAfa_to_EU
Correlation = 0.043
predicted
simulated
0e+00
2e−04
4e−04
0.3
0.4
0.5
0.6
0.35
0.45
0.55
0.65

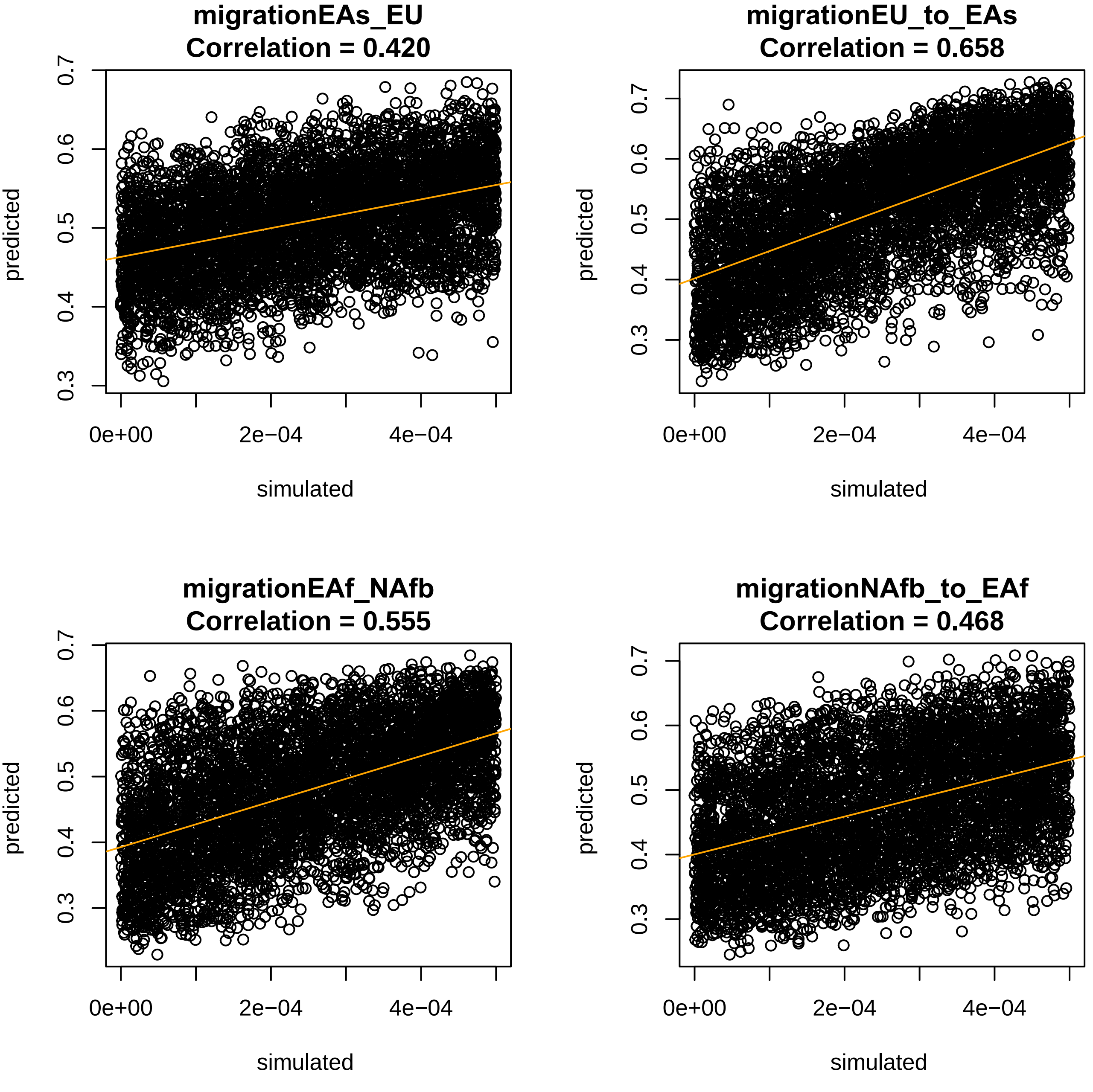
migrationEAs_EU
Correlation = 0.420
migrationEU_to_EAs
Correlation = 0.658
migrationEAf_NAfb
Correlation = 0.555
migrationNAfb_to_EAf
Correlation = 0.468
predicted
simulated
0.3
0.4
0.5
0.6
0.7
0e+00
2e−04
4e−04

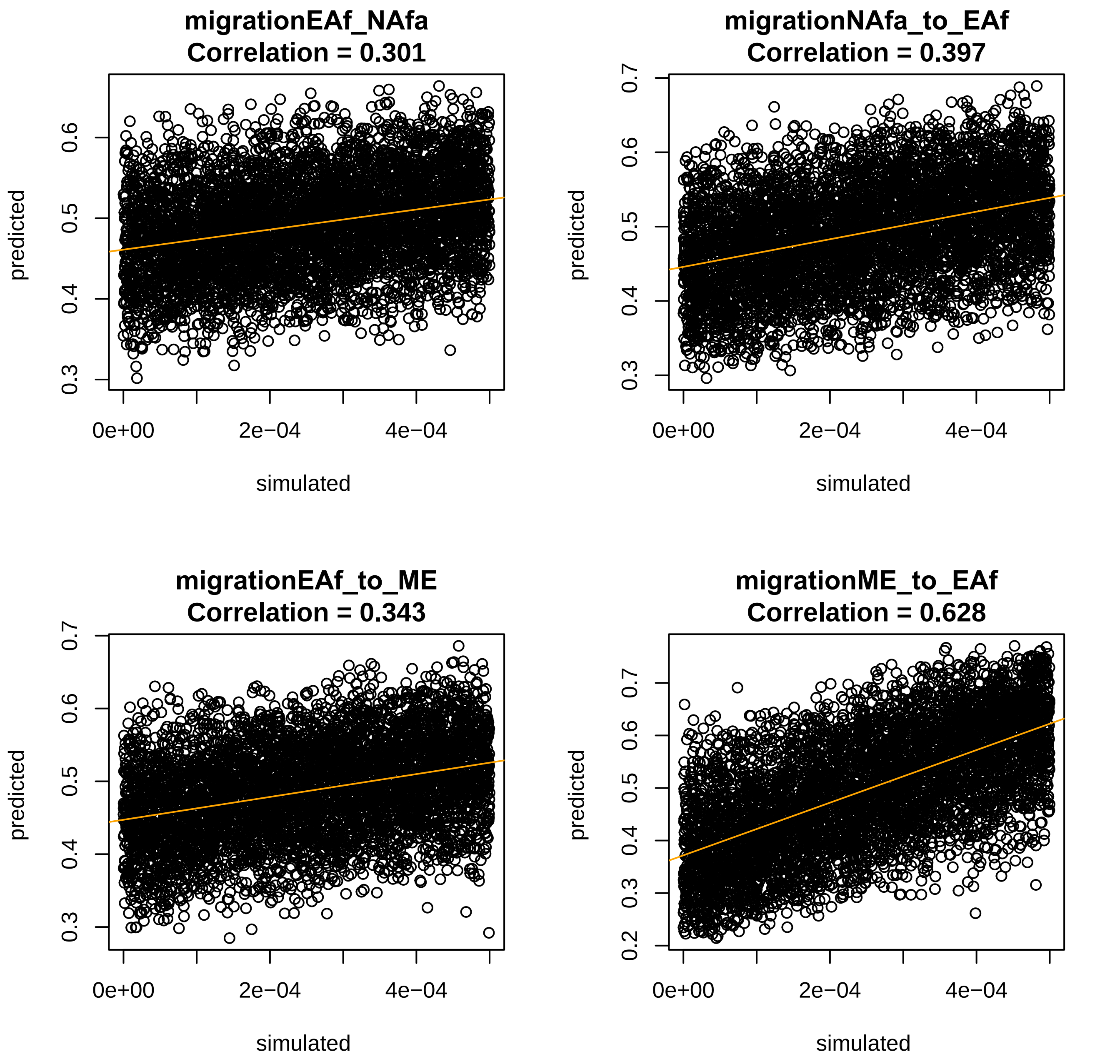
migrationEAf_NAfa
Correlation = 0.301
migrationNAfa_to_EAf
Correlation = 0.397
migrationEAf_to_ME
Correlation = 0.343
migrationME_to_EAf
Correlation = 0.628
predicted
simulated
0e+00
2e−04
4e−04
0.2
0.3
0.4
0.5
0.6
0.7

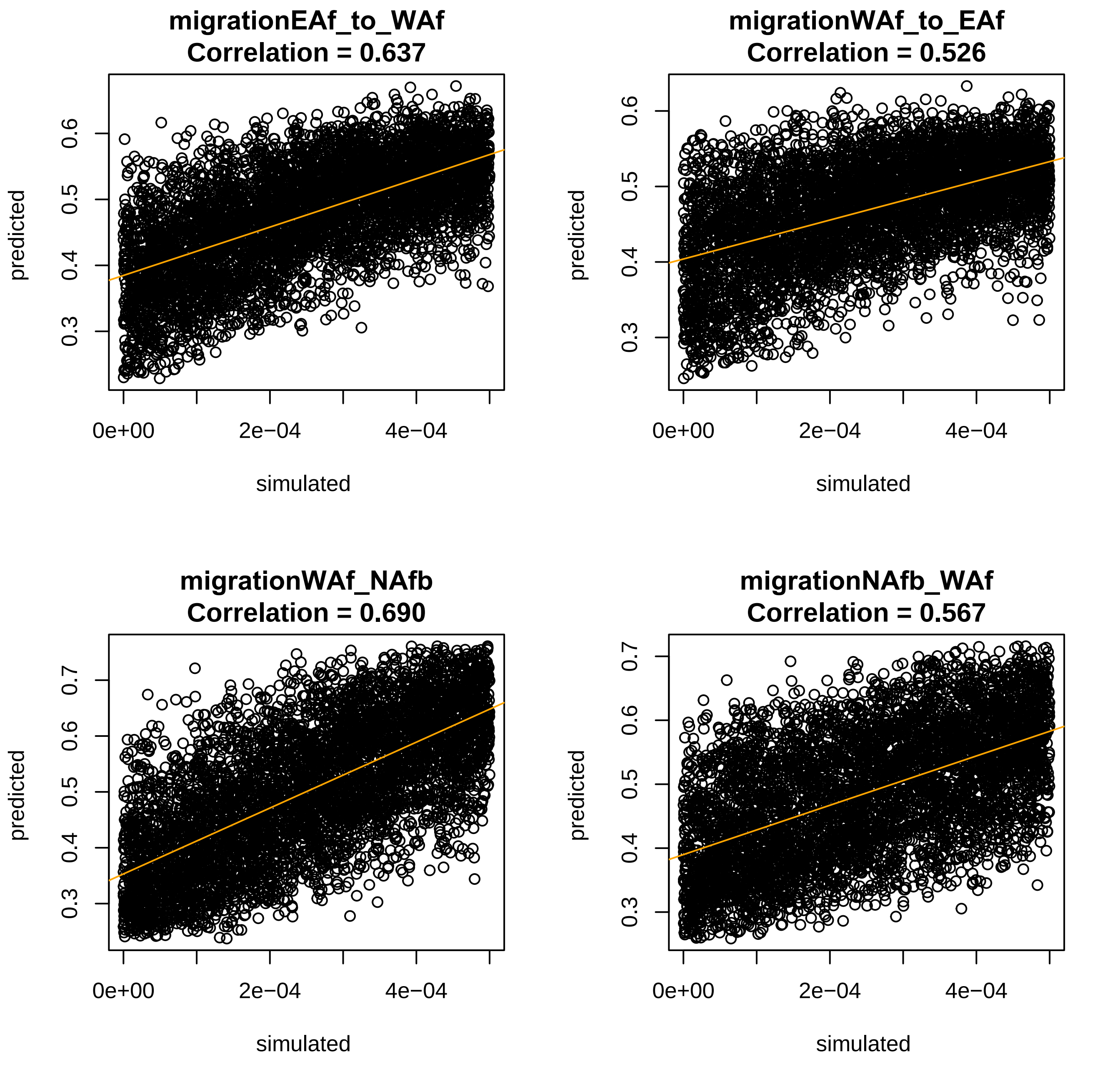
migrationEAf_to_WAf
Correlation = 0.637
migrationWAf_to_EAf
Correlation = 0.526
migrationWAf_NAfb
Correlation = 0.690
migrationNAfb_WAf
Correlation = 0.567
predicted
simulated
0e+00
2e−04
4e−04
0.3
0.4
0.5
0.6
0.7

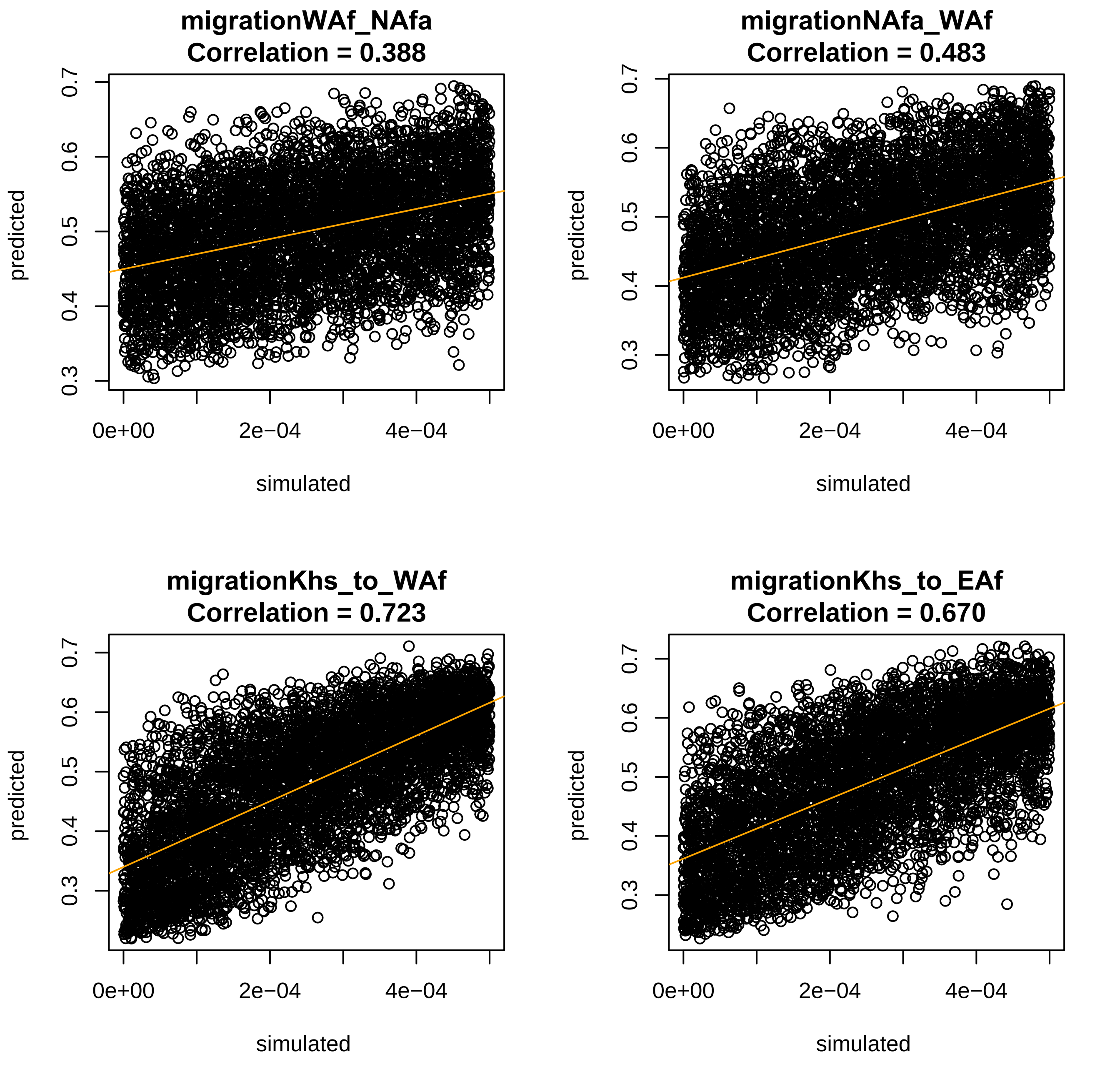
migrationWAf_NAfa
Correlation = 0.388
migrationNAfa_WAf
Correlation = 0.483
migrationKhs_to_WAf
Correlation = 0.723
migrationKhs_to_EAf
Correlation = 0.670
predicted
simulated
0.3
0.4
0.5
0.6
0.7
0e+00
2e−04
4e−04

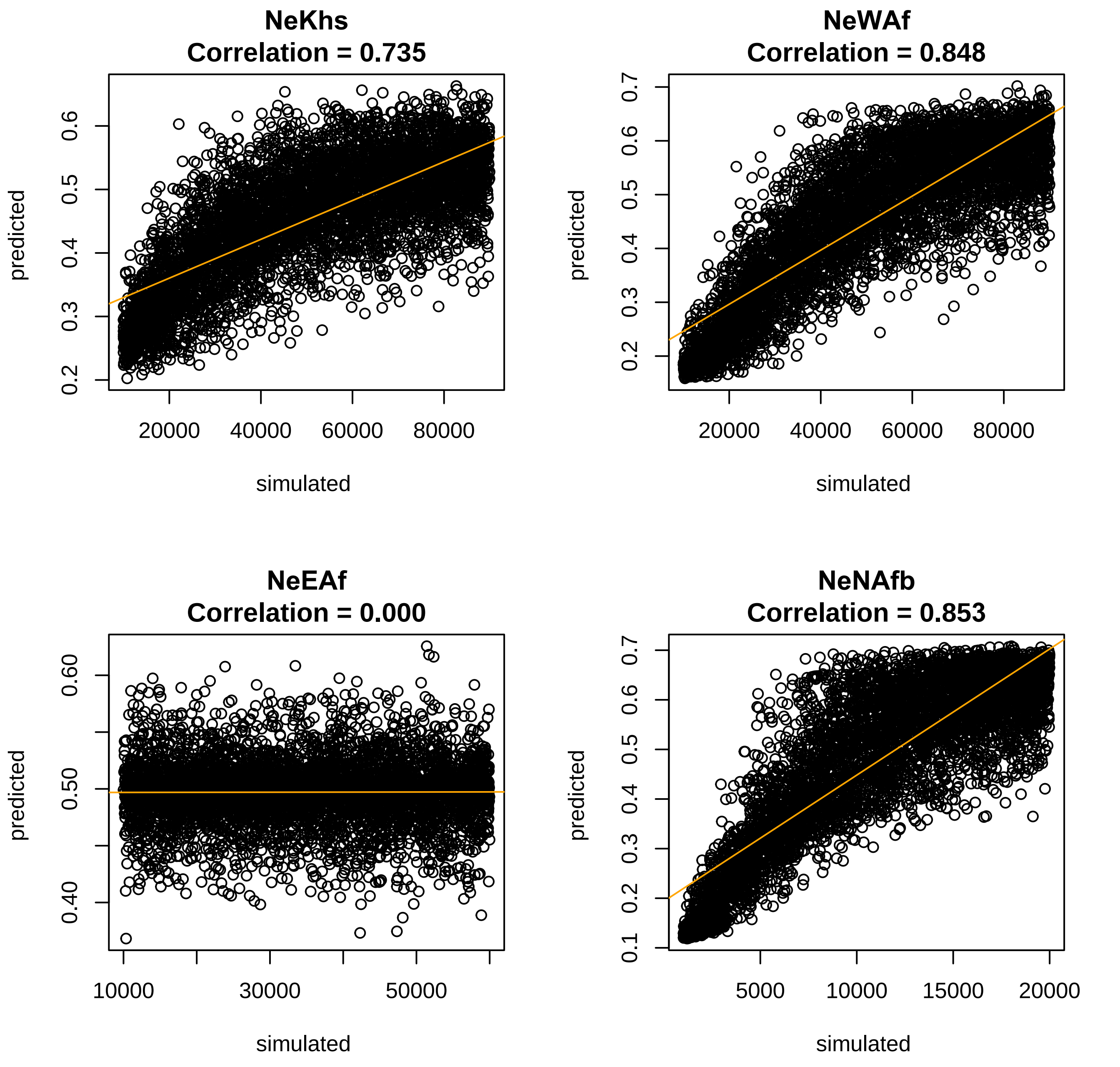
NeKhs
Correlation = 0.735
NeWAf
Correlation = 0.848
NeEAf
Correlation = 0.000
NeNAfb
Correlation = 0.853
predicted
simulated
0.2
0.3
0.4
0.5
0.6
0.7
0.1
0.40
0.50
0.60
20000
40000
60000
80000
10000
30000
50000
5000
15000

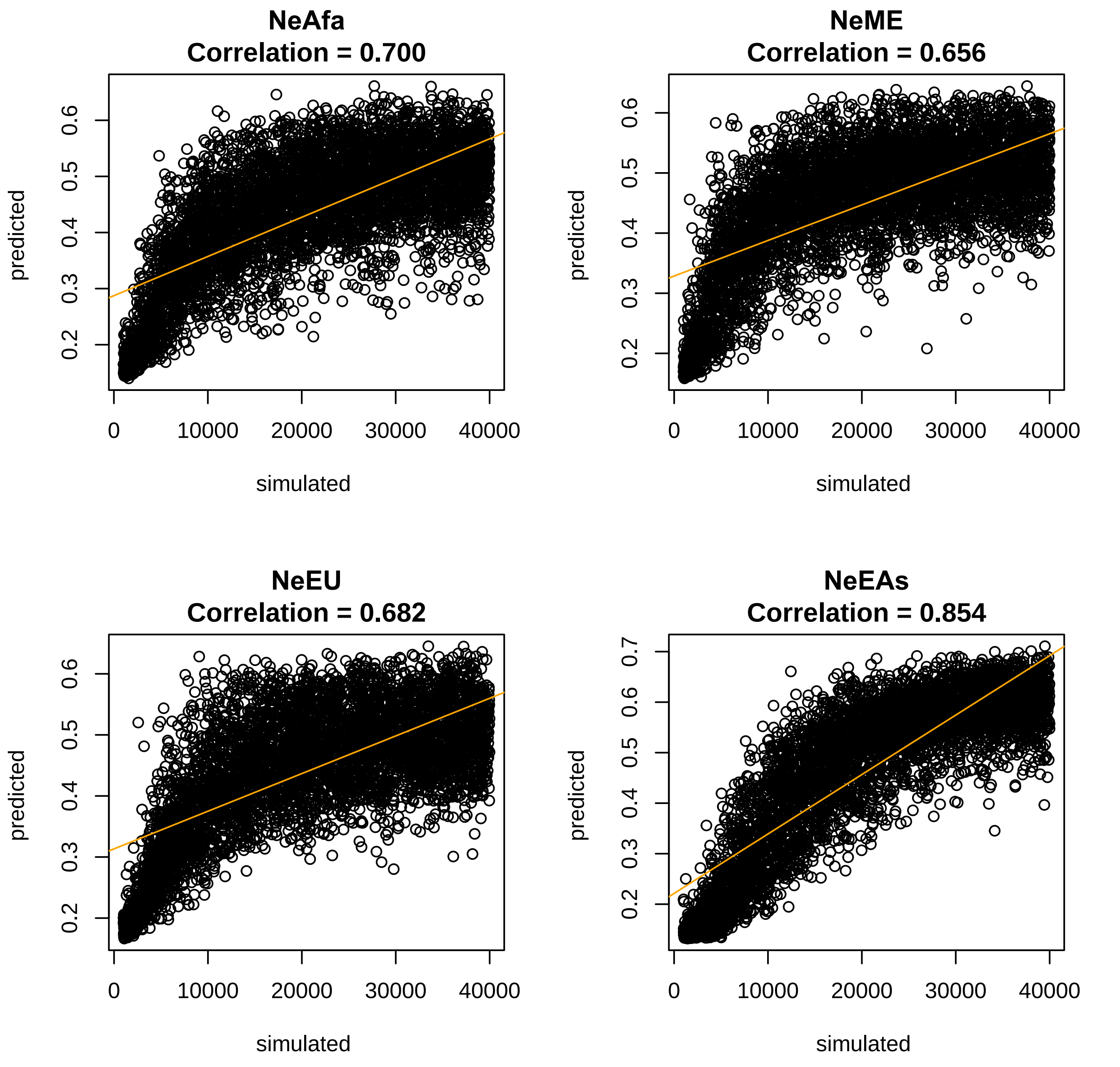
NeAfa
Correlation = 0.700
NeME
Correlation = 0.656
NeEU
Correlation = 0.682
NeEAs
Correlation = 0.854
predicted
simulated
0
10000
20000
30000
40000
0.2
0.3
0.4
0.5
0.6
0.7

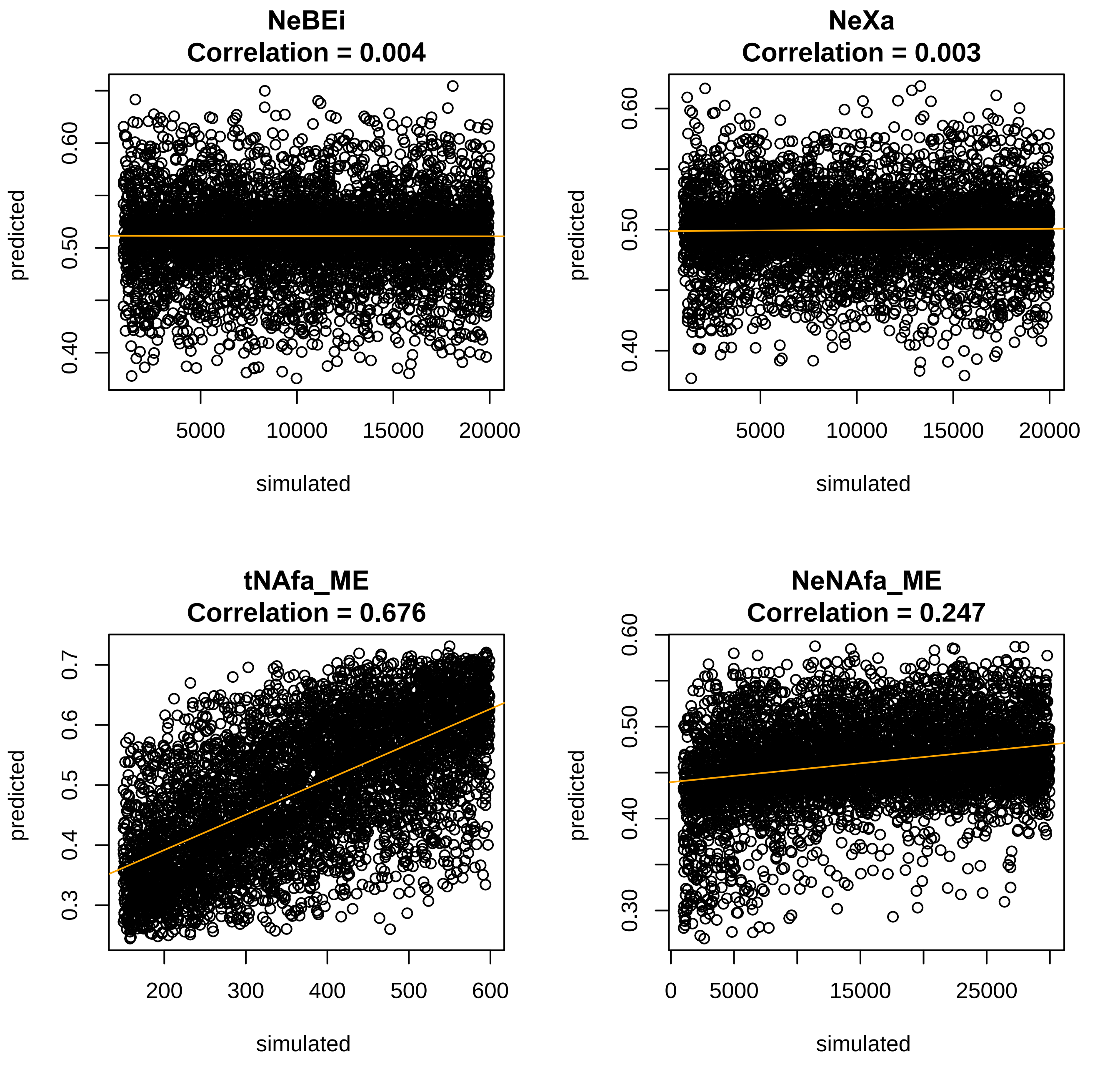
NeBEi
Correlation = 0.004
NeXa
Correlation = 0.003
tNAfa_ME
Correlation = 0.676
NeNAfa_ME
Correlation = 0.247
predicted
simulated
0.40
0.50
0.60
0.3
0.4
0.5
0.6
0.7
0.30
5000
10000
15000
20000
25000
0
200
300
400
500
600

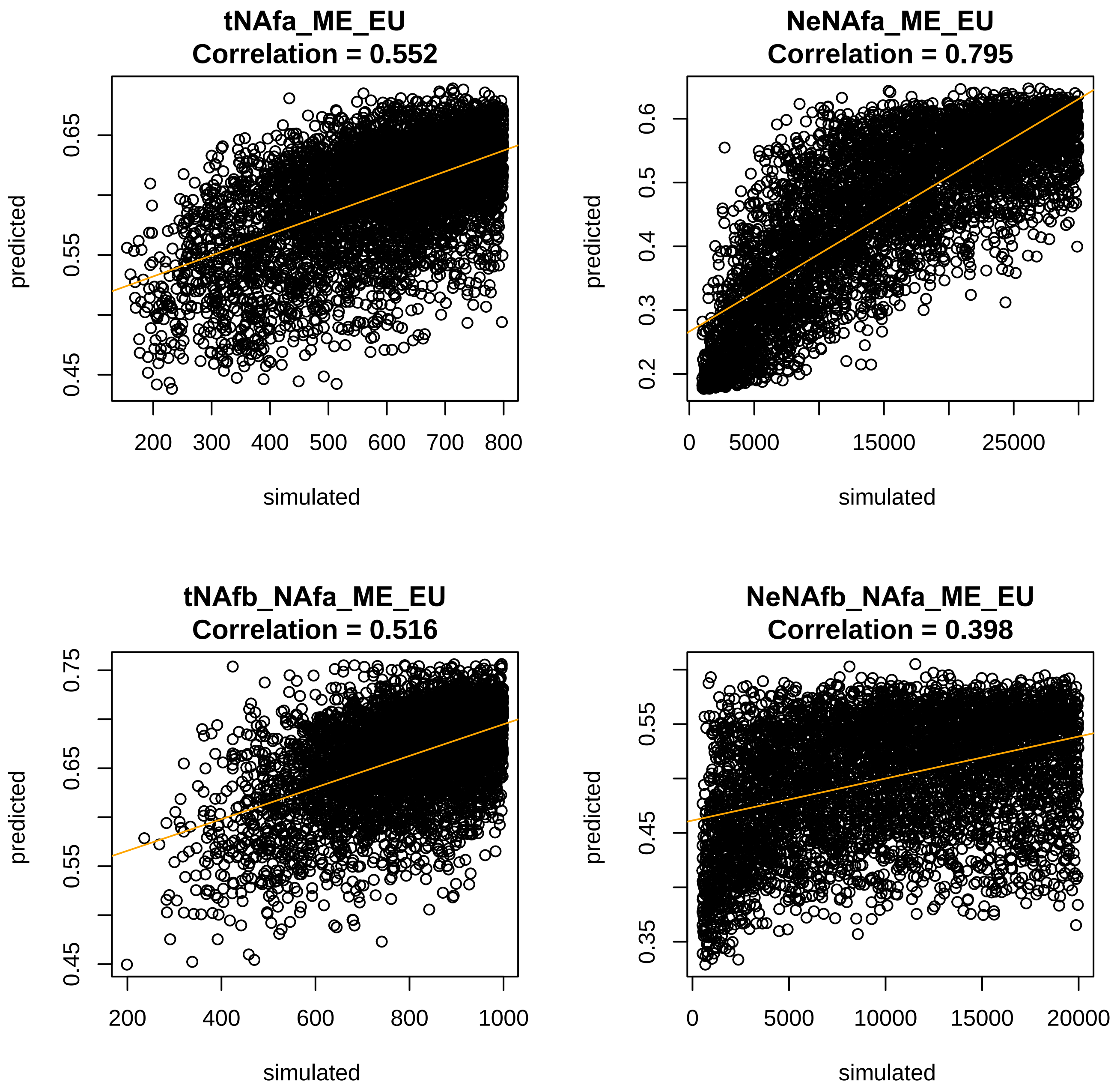
tNAfa_ME_EU
Correlation = 0.552
predicted
0.45
0.55
0.65
200 300 400 500 600 700 800
simulated
NeNAfa_ME_EU
Correlation = 0.795
predicted
0.2 0.3 0.4 0.5 0.6
0 5000 15000 25000
simulated
tNAfb_NAfa_ME_EU
Correlation = 0.516
predicted
0.45 0.55 0.65 0.75
200 400 600 800 1000
simulated
NeNAfb_NAfa_ME_EU
Correlation = 0.398
predicted
0.35 0.45 0.55
0 5000 10000 15000 20000
simulated

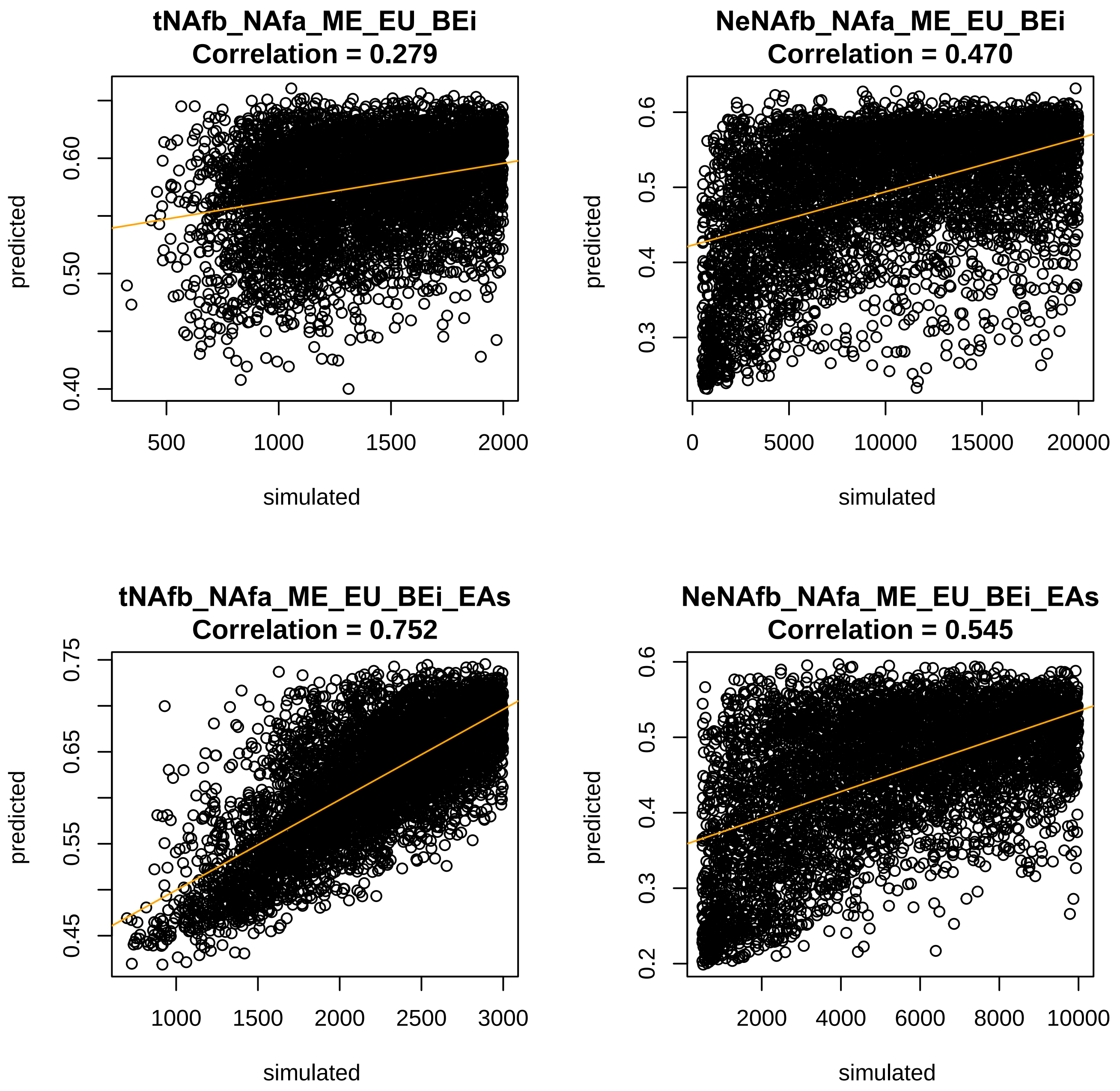
tNAfb_NAfa_ME_EU_BEi
Correlation = 0.279
predicted
simulated
0.40
0.50
0.60
500
1000
1500
2000
NeNAfb_NAfa_ME_EU_BEi
Correlation = 0.470
predicted
simulated
0.3
0.4
0.5
0.6
0
5000
10000
15000
20000
tNAfb_NAfa_ME_EU_BEi_EAs
Correlation = 0.752
predicted
simulated
0.45
0.55
0.65
0.75
1000
1500
2000
2500
3000
NeNAfb_NAfa_ME_EU_BEi_EAs
Correlation = 0.545
predicted
simulated
0.2
0.3
0.4
0.5
0.6
2000
4000
6000
8000
10000

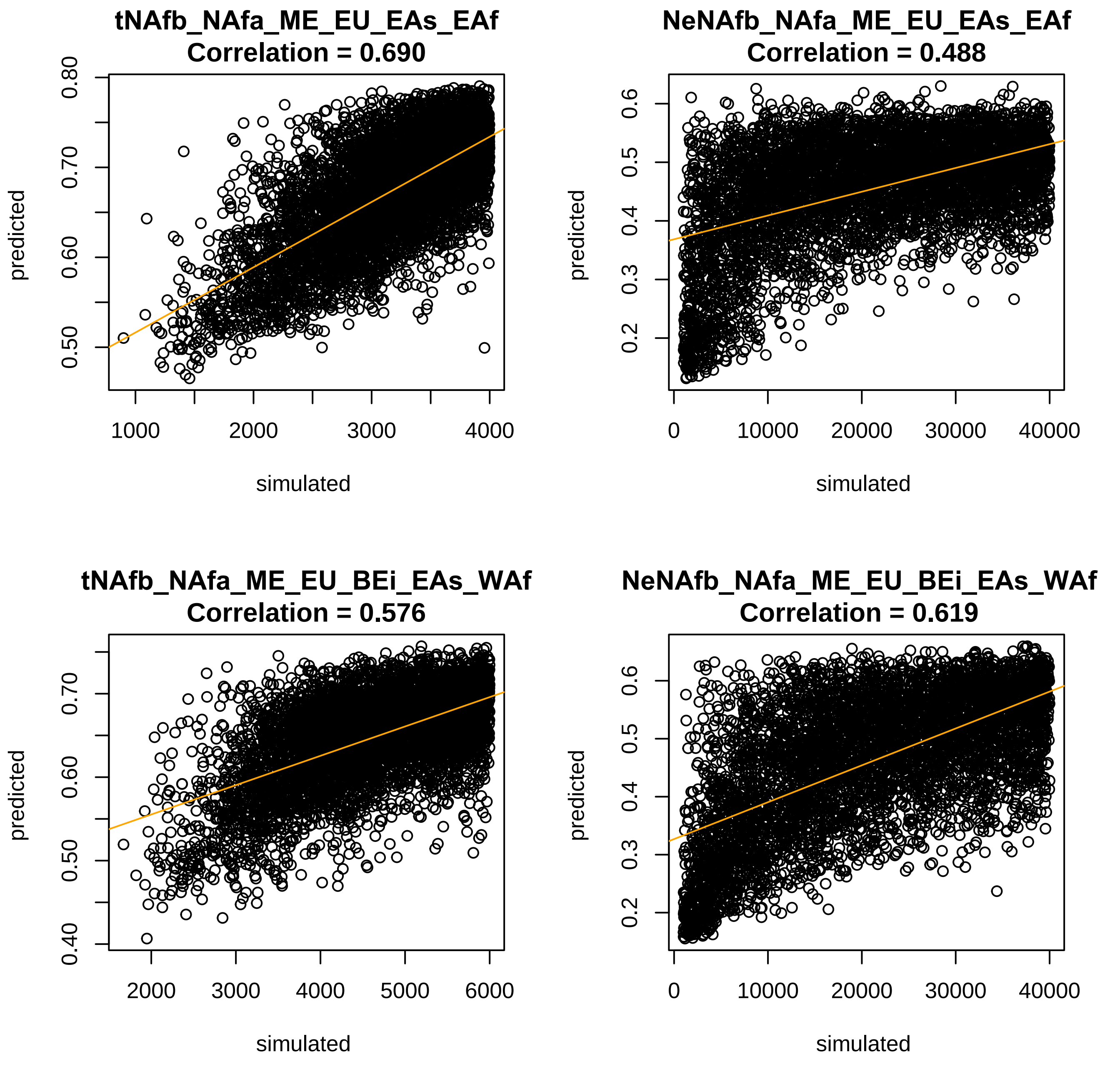

tNAfb_NAfa_ME_EU_EAs_EAf
Correlation = 0.690
NeNAfb_NAfa_ME_EU_EAs_EAf
Correlation = 0.488
tNAfb_NAfa_ME_EU_BEi_EAs_WAf
Correlation = 0.576
NeNAfb_NAfa_ME_EU_BEi_EAs_WAf
Correlation = 0.619
predicted
simulated

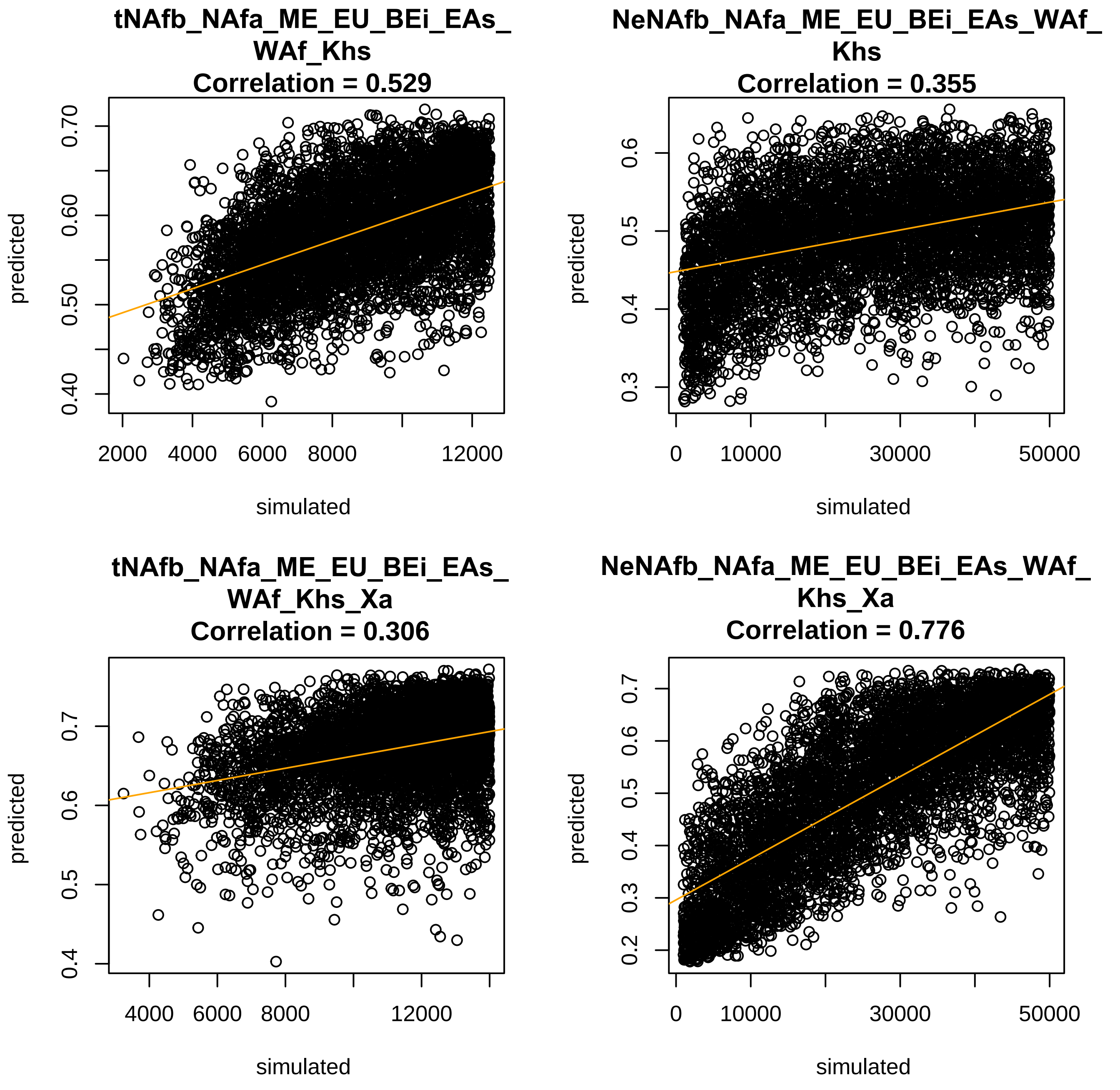
tNAfb_NAfa_ME_EU_BEi_EAs_WAf_Khs
Correlation = 0.529
predicted
0.40
0.50
0.60
0.70
2000
4000
6000
8000
12000
simulated
NeNAfb_NAfa_ME_EU_BEi_EAs_WAf_Khs
Correlation = 0.355
predicted
0.3
0.4
0.5
0.6
0
10000
30000
50000
simulated
tNAfb_NAfa_ME_EU_BEi_EAs_WAf_Khs_Xa
Correlation = 0.306
predicted
0.4
0.5
0.6
0.7
4000
6000
8000
12000
simulated
NeNAfb_NAfa_ME_EU_BEi_EAs_WAf_Khs_Xa
Correlation = 0.776
predicted
0.2
0.3
0.4
0.5
0.6
0.7
0
10000
30000
50000
simulated

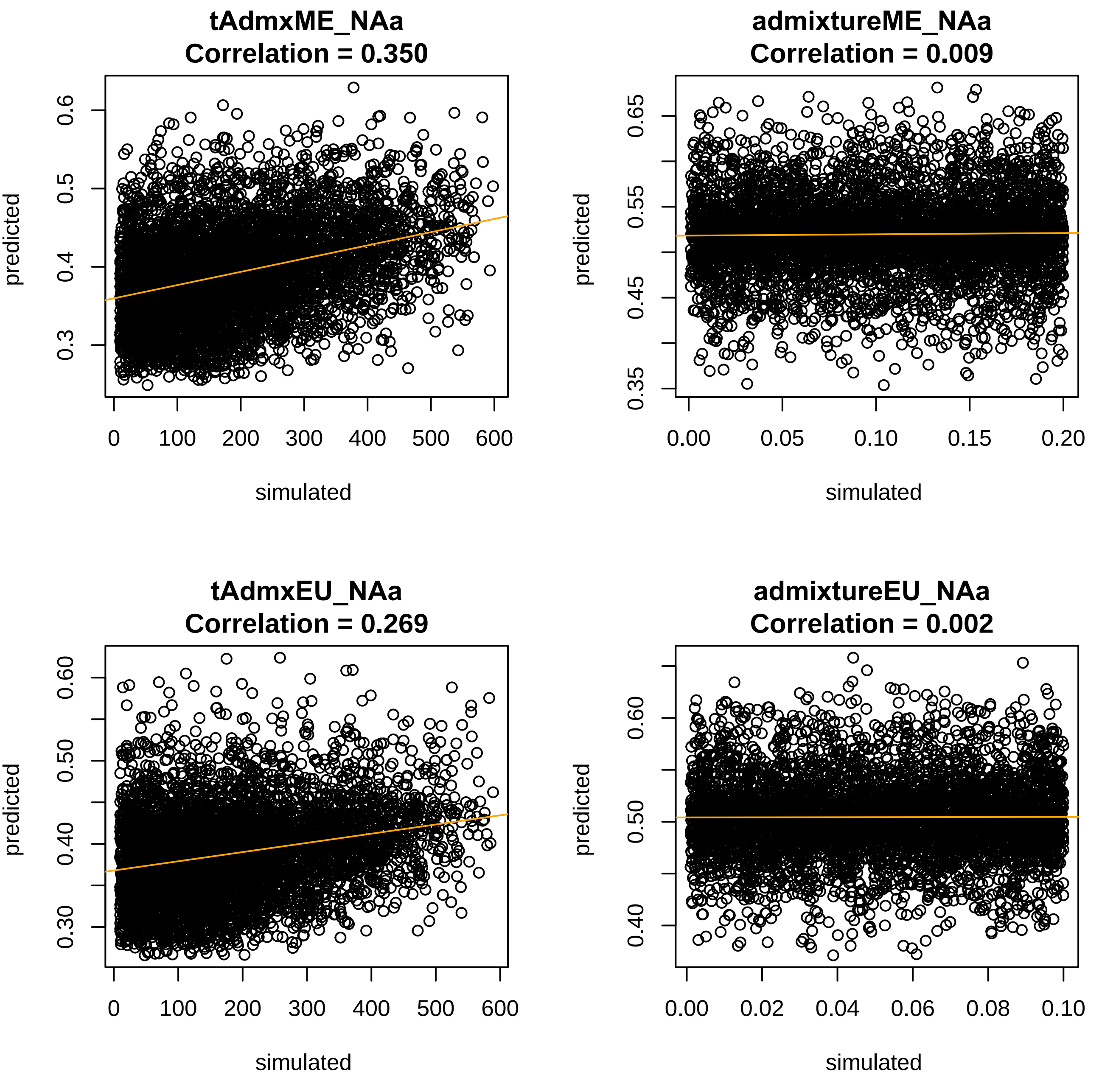
tAdmxME_NAa
Correlation = 0.350
predicted
simulated
admixtureME_NAa
Correlation = 0.009
predicted
simulated
tAdmxEU_NAa
Correlation = 0.269
predicted
simulated
admixtureEU_NAa
Correlation = 0.002
predicted
simulated

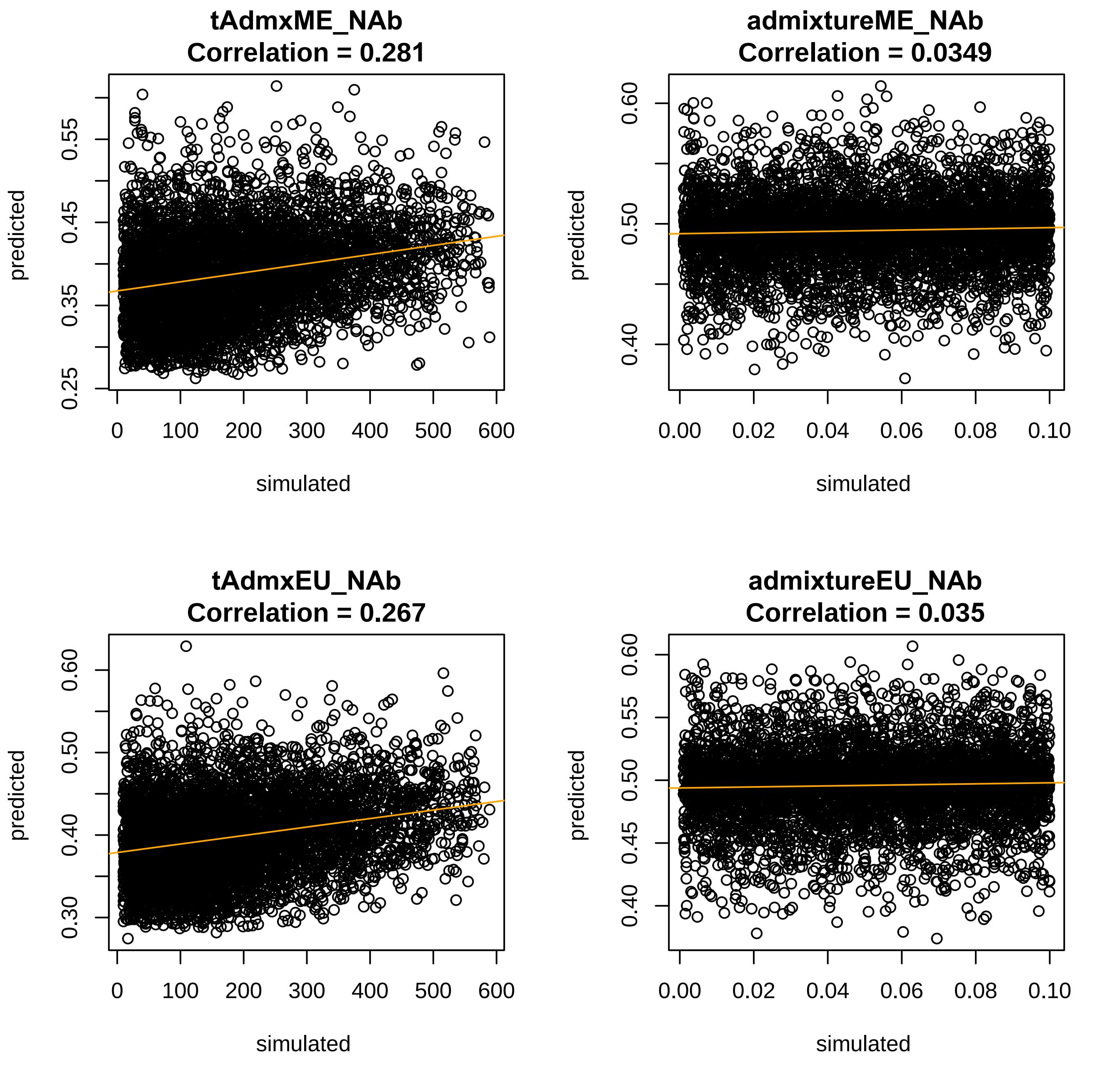
tAdmxME_NAb
Correlation = 0.281
admixtureME_NAb
Correlation = 0.0349
tAdmxEU_NAb
Correlation = 0.267
admixtureEU_NAb
Correlation = 0.035
predicted
simulated
0.25
0.35
0.45
0.55
0.30
0.40
0.50
0.60
0.45
0.55
0
100
200
300
400
500
600
0.00
0.02
0.04
0.06
0.08
0.10

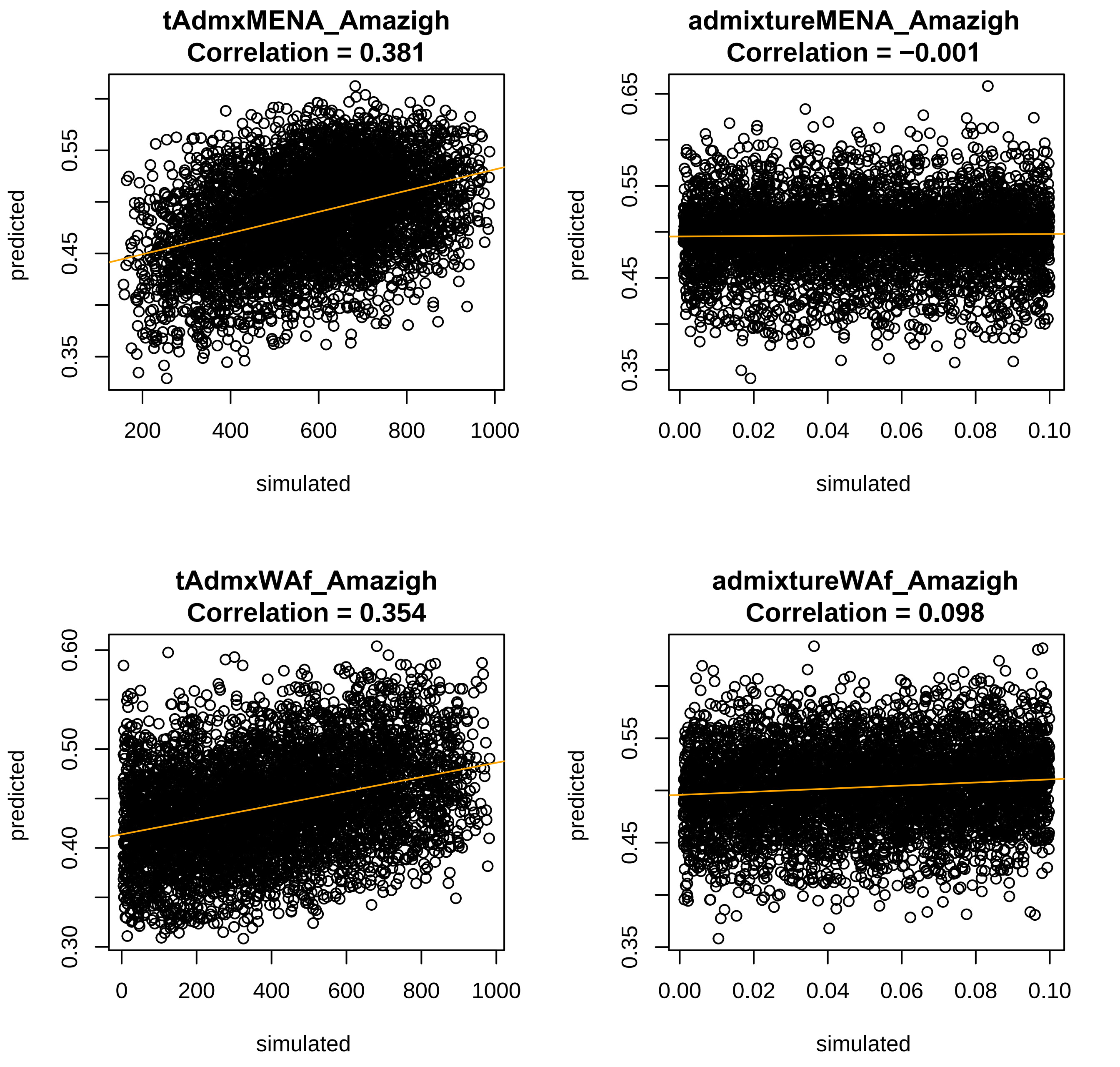
tAdmxMENA_Amazigh
Correlation = 0.381
admixtureMENA_Amazigh
Correlation = −0.001
tAdmxWAf_Amazigh
Correlation = 0.354
admixtureWAf_Amazigh
Correlation = 0.098
predicted
simulated

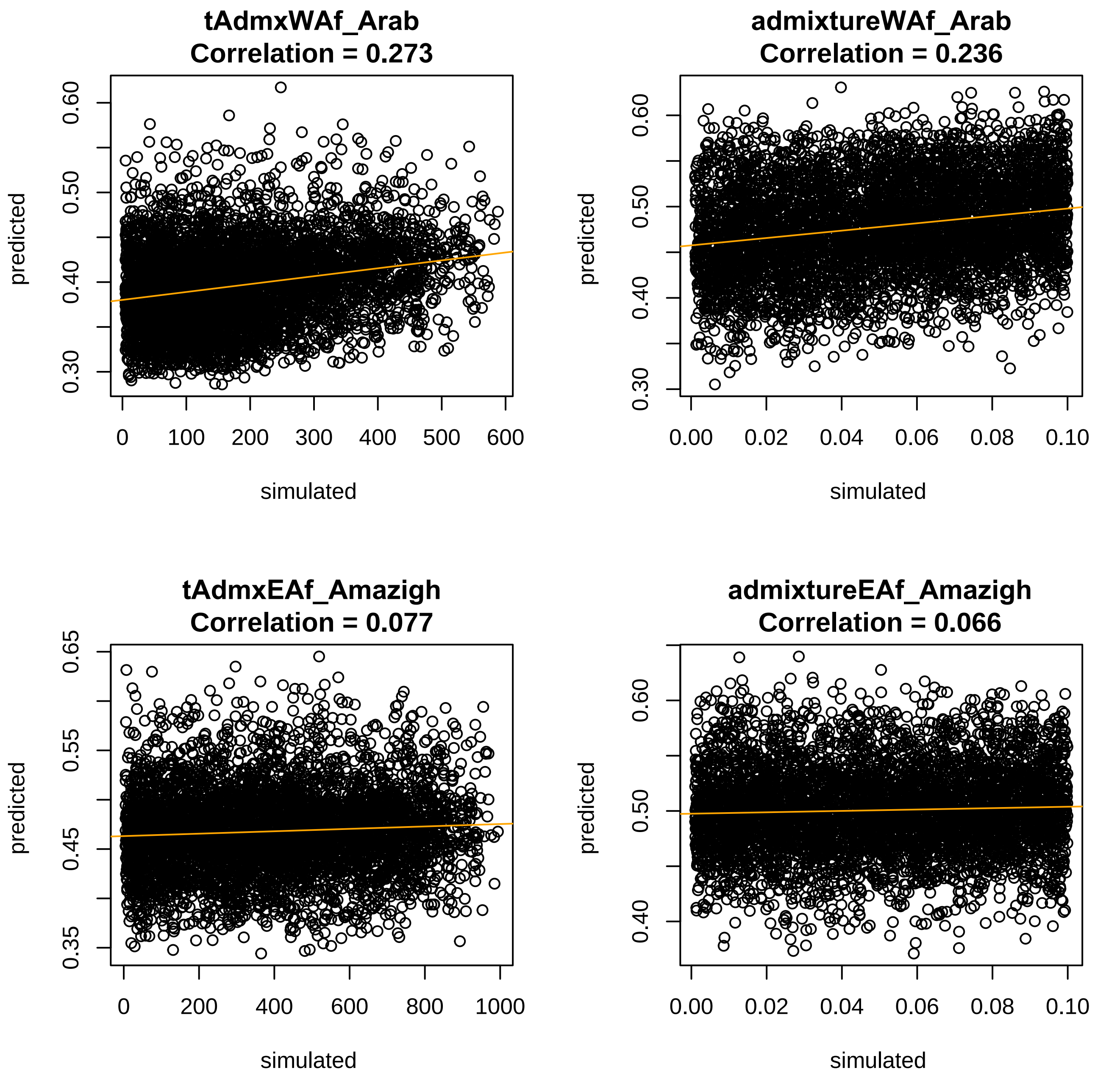
tAdmxWAf_Arab
Correlation = 0.273
admixtureWAf_Arab
Correlation = 0.236
tAdmxEAf_Amazigh
Correlation = 0.077
admixtureEAf_Amazigh
Correlation = 0.066
predicted
simulated
0.30
0.40
0.50
0.60
0.35
0.45
0.55
0.65
0
100
200
300
400
500
600
800
1000
0.00
0.02
0.04
0.06
0.08
0.10

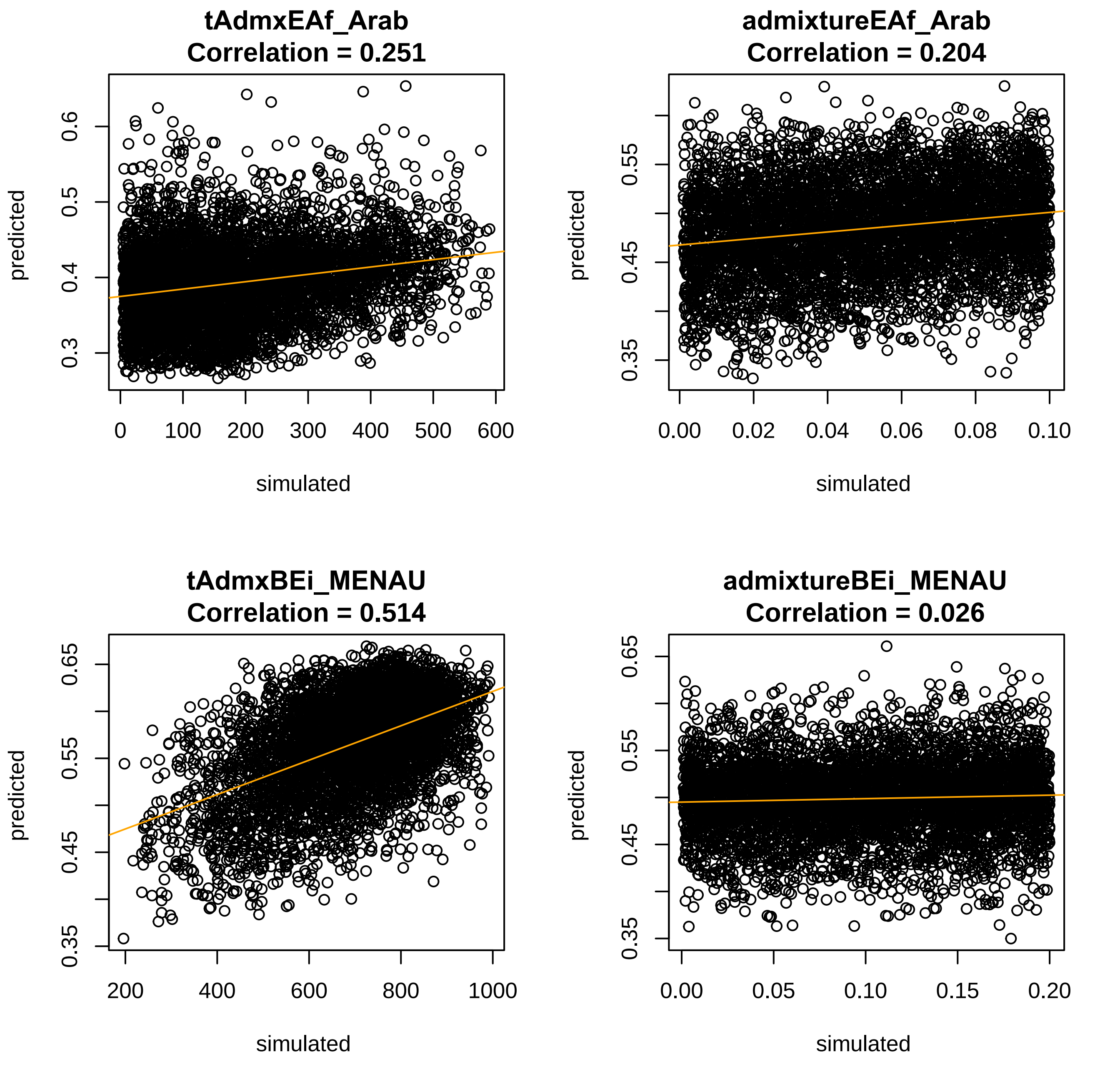
tAdmxEAf_Arab
Correlation = 0.251
admixtureEAf_Arab
Correlation = 0.204
tAdmxBEi_MENAU
Correlation = 0.514
admixtureBEi_MENAU
Correlation = 0.026
predicted
simulated
0
100
200
300
400
500
600
0.3
0.4
0.5
0.6
0.00
0.02
0.04
0.06
0.08
0.10
0.35
0.45
0.55
200
400
600
800
1000
0.65
0.05
0.15
0.20

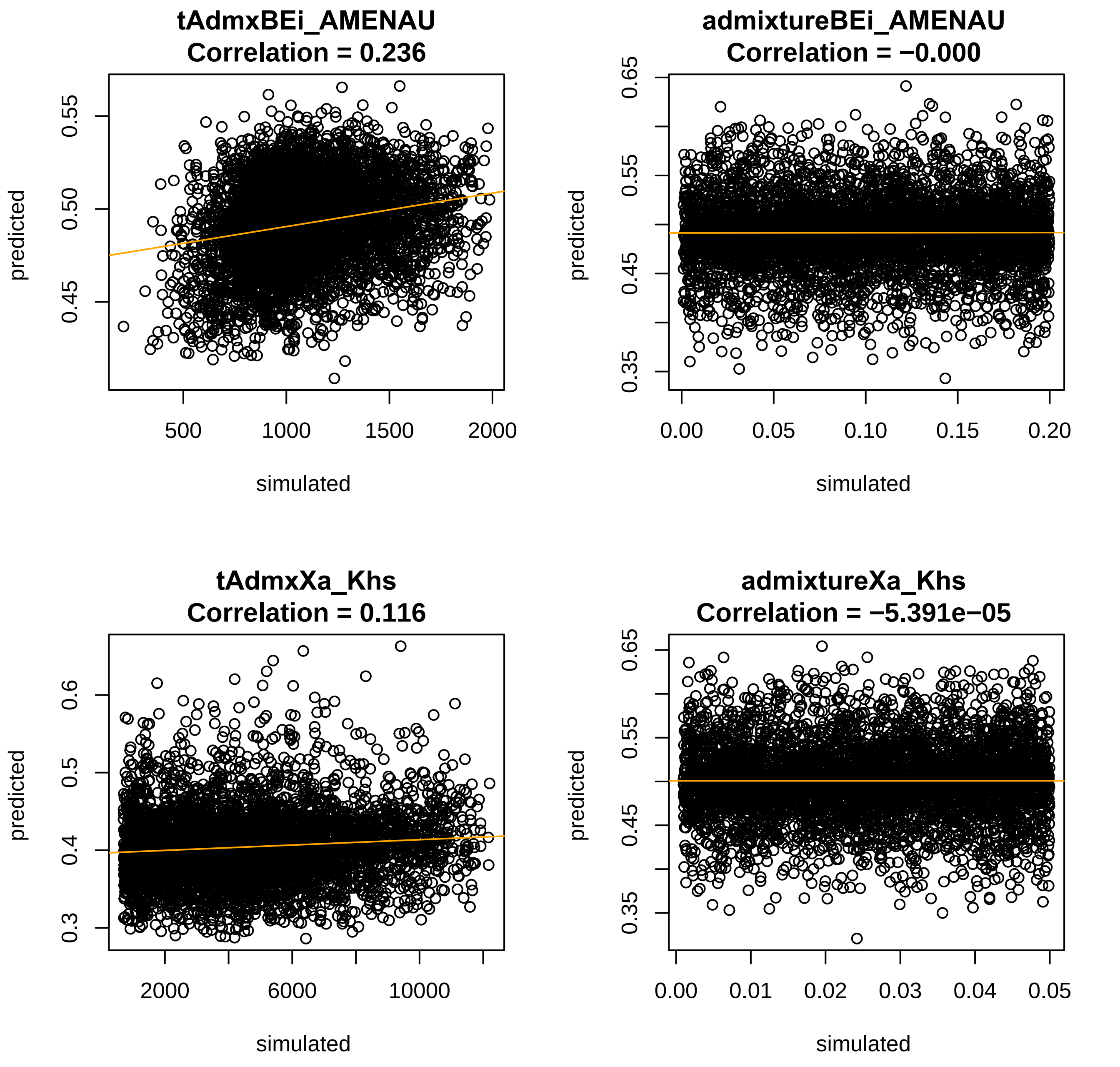

tAdmxBEi_AMENAU
Correlation = 0.236
admixtureBEi_AMENAU
Correlation = −0.000
tAdmxXa_Khs
Correlation = 0.116
admixtureXa_Khs
Correlation = −5.391e−05
predicted
simulated
0.45
0.50
0.55
500
1000
1500
2000
0.35
0.45
0.55
0.65
0.00
0.05
0.10
0.15
0.20
0.3
0.4
0.5
0.6
2000
6000
10000
0.01
0.02
0.03
0.04
0.05

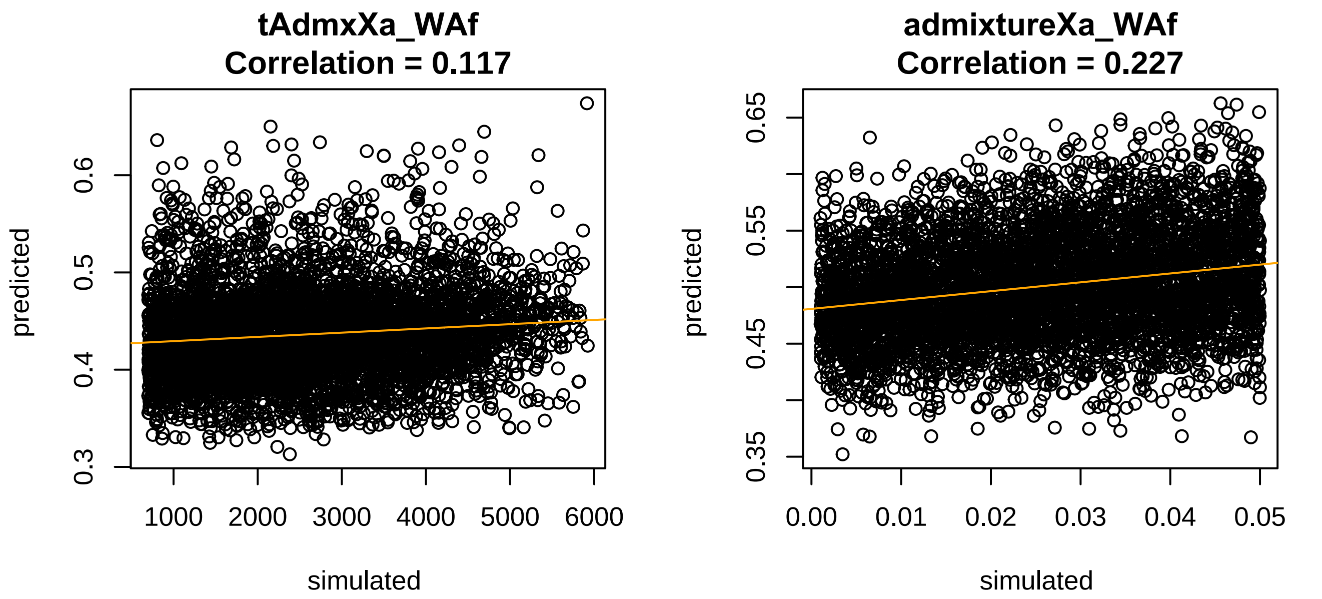

tAdmxXa_WAf
Correlation = 0.117
predicted
0.3
0.4
0.5
0.6
1000
2000
3000
4000
5000
6000
simulated
admixtureXa_WAf
Correlation = 0.227
predicted
0.35
0.45
0.55
0.65
0.00
0.01
0.02
0.03
0.04
0.05
simulated
