## Supplementary File Description for "Modelling the demographic history of human North African genomes points to soft split divergence between populations"

Additional file 1: Supplementary figures S1 to S11 and Supplementary Tables S1 to S10

Additional file 2: Xlsx file with the posterior values of the accepted models in the ABC-DL analysis (Model D in first analysis and Model D4 in second analysis). Factor 2, Kullback-Leiber, and Spearman correlation tables for the accepted models in both ABC-DL analysis.

Additional file 3: Histograms with the posterior versus prior distributions of all parameters for the best model in ABC-DL analysis (Model D4)

Additional file 4: Spearman correlation plots for all parameters in the best model in the ABC-DL analysis (Model D4)

Additional file 5: Fitness error values of all iterations of the GP4PG analysis. Parameter values for the best 10 models in the GP4PG analysis.
