## Supplementary File 1 for "Modelling the demographic history of human North African genomes points to soft split divergence between populations"

#### Figures Supplementary_ LEGENDS:

**Fig. S 1: Principal Component Analysis on genomic dataset of North Africa. Visualization of PC3 & PC4**

**Fig. S 2: CrossValidation error of ADMIXTURE analysis with K=2 to K=12. Best K is K=3**

**Fig. S 3: ADMIXTURE analysis on 364 individuals with K=2 to K=12**

**Fig. S 4: Competing topologies tested in ABC-DL analysis.** Seven different topologies included on the ABC-DL analyses considering North African Arab (NAa), North Africa Amazigh (Ama), Middle Eastern (ME), European (Eu), East Asian (EAs), East African (EAf), West African (WAf), and Ju/'hoansi (San) populations.

**Fig. S 5: Replication PCA for Model D_4 in ABC-DL analysis.** PCA for 1000 simulations of the model D_4, -the best model in the ABC-DL analysis- and the replication dataset of observed data. Observed data is an outlier in the PCA indicating that the ABC-DL model cannot properly replicate the diversity observed in the dataset.

**Fig. S 6: Box plot of the distances between each simulation in the PCA and the centroid of the PCA**. The red dot represents the observed data as an outlier of the distances.

**Fig. S 4: Competing topologies tested in GP4PG analysis.** The competing topologies for the GP4PG analysis are the same as the ones used in the ABC-DL analysis but discarding Model A, due to being the worst performing one in previous analysis and due resource consumption.

**Fig. S 5: Coordinates of the different ecodemes we are testing in the GP4PG analysis.** Each ecodeme has the exact same size and major geographical barriers such as seas and deserts has been removed for the sake of simplicity.

**Fig. S 6: Fitness of the different runs of the genetic algorithm. a.** Distribution of the fitness error of 40 independent iterations of the GP4PG algorithm with 6 competing topologies (B to G in ABC-DL) during 200 generations. Model D appears as the most selected model in a fourth of all the iterations, with D_15 as the model with the least error. **b.** PCA plot comparing the jSFS obtained from simulations of the best ABC-DL model with the 10 best GP4PG models. GP4PG simulations explain the observed data better than the ABC-DL. **c**. Same PCA plot as **b** but not including the simulations from ABC-DL result. Models C_29 and C_39 are the ones that show a more similar jSFS to the one produced by the observed data.

**Fig. S 7: Observed heterozygosity per individual compared by superpopulation.** Sub-Saharan populations present a higher heterozygosity than Eurasian populations, North African individuals have heterozygosity levels between the sub-Saharan and the Eurasians, probably due to gene flow from sub-Sharan populations to north African individuals.

**Fig. S 8: A tree-based depiction of the demographic relationships between different ecodemes and topodemes.** Each node codes for a demographic event that occurs at a given time (dashed lines).

**Fig. S 9: Fitness of the different runs of the genetic algorithm. a.** Distribution of the fitness error of 40 independent iterations of the GP4PG algorithm with 6 competing topologies (B to G in ABC-DL) during 200 generations. Model D appears as the most selected model in a fourth of all the iterations, with D_15 as the model with the least error. **b.** PCA plot comparing the jSFS obtained from simulations of the best ABC-DL model with the 10 best GP4PG models. GP4PG simulations explain the observed data better than the ABC-DL. **c**. Same PCA plot as **b** but not including the simulations from ABC-DL result. Models C_29 and C_39 are the ones that show a more similar jSFS to the one produced by the observed data.

**Fig. S 10: Observed heterozygosity per individual compared by superpopulation.** Sub-Saharan populations present a higher heterozygosity than Eurasian populations, North African individuals have heterozygosity levels between the sub-Saharan and the Eurasians, probably due to gene flow from sub-Sharan populations to north African individuals.

**Fig. S 11: A tree-based depiction of the demographic relationships between different ecodemes and topodemes.** Each node codes for a demographic event that occurs at a given time (dashed lines).

.

**TABLES:**

| | ModelA | ModelB | ModelC | ModelD | ModelE | ModelF | ModelG |
|---|---|---|---|---|---|---|---|
| **ModelA** | **1** | 0 | 0 | 0 | 0 | 0 | 0 |
| **ModelB** | 0 | **0.7941** | 0.1023 | 0.0122 | 0 | 0.0914 | 0 |
| **ModelC** | 0 | 0.0603 | **0.9395** | 0.0001 | 0 | 0.0001 | 0 |
| **ModelD** | 0 | 0.0145 | 0.002 | **0.7579** | 0 | 0.2256 | 0.0001 |
| **ModelE** | 0 | 0 | 0 | 0 | **1** | 0 | 0 |
| **ModelF** | 0 | 0.1565 | 0.002 | 0.1881 | 0 | **0.6534** | 0.0001 |
| **ModelG** | 0 | 0 | 0 | 0 | 0 | 0 | **1** |

**Table S1: Confusion matrix computed with the 7 models under evaluation. 50 randomly sampled simulations per model were used as "observed" data for the ABC-DL algorithm**. Diagonal, in bold, shows the probability of a model being correctly assigned by the A

| ModelA | ModelB | ModelC | ModelD | ModelE | ModelF | ModelG |
|---|---|---|---|---|---|---|
| 0 | 0 | 0 | 0.922 | 0 | 0.078 | 0 |

**Table S 2:Proportion of accepted simulations using postpr function for the "abc" package with tolerance = 0.0008**. Model D is present 92.2% of times in the 1000 closest simulations to the observed data.

| | ModelA | ModelB | ModelC | ModelD | ModelE | ModelF | ModelG |
|---|---|---|---|---|---|---|---|
| **ModelA** | NA | NA | NA | Inf | NA | Inf | NA |
| **ModelB** | NA | NA | NA | Inf | NA | Inf | NA |
| **ModelC** | NA | NA | NA | Inf | NA | Inf | NA |
| **ModelD** | 0 | 0 | 0 | 1 | 0 | 0.08459 | 0 |
| **ModelE** | NA | NA | NA | Inf | NA | Inf | NA |
| **ModelF** | 0 | 0 | 0 | 11.8205 | 0 | 1 | 0 |
| **ModelG** | NA | NA | NA | Inf | NA | Inf | NA |

**Table S 3: Bayes factor for the ABC-DL topology discrimination analysis**. Model D is 11.8 times better at explaining the observed data than the second-best model (Model F).

| | Model D1 | Model D2 | Model D3 | Model D4 | Model D5 |
|---|---|---|---|---|---|
| **Model D1** | **0.5888** | 0.0058 | 0.0064 | 0.0061 | 0.3933 |
| **Model D2** | 0.0019 | **0.3712** | 0.3499 | 0.2759 | 0.0011 |
| **Model D3** | 0.0003 | 0.3507 | **0.3557** | 0.2931 | 0.0001 |
| **Model D4** | 0.0003 | 0.2729 | 0.3020 | **0.4248** | 0 |
| **Model D5** | 0.3888 | 0 | 0 | 0 | **0.6112** |

**Table S 4: Confusion matrix computed with the five D models under evaluation. 50 randomly sampled simulations per model were used as “observed” data for the ABC-DL algorithm.** Diagonal, in bold, shows the probability of a model being correctly assigned by the ABC.

| Model D1 | Model D2 | Model D3 | Model D4 | Model D5 |
|---|---|---|---|---|
| 0.0167 | 0.0800 | 0.0944 | 0.7622 | 0.0468 |

**Table S 5: Proportion of accepted simulations using postpr function for the “abc” package with tolerance = 0.001**. Model D4 is present 76.22% of times in the 1000 closest simulations to the observed data.

| | Model D1 | Model D2 | Model D3 | Model D4 | Model D5 |
|---|---|---|---|---|---|
| **Model D1** | 1 | 0.2087 | 0.1768 | 0.0219 | 0.3567 |
| **Model D2** | 4.7913 | 1 | 0.8471 | 0.1049 | 1.7093 |
| **Model D3** | 5.6560 | 1.1805 | 1 | 0.1238 | 2.0178 |
| **Model D4** | 45.6714 | 9.5321 | 8.0748 | 1 | 16.2932 |
| **Model D5** | 2.8031 | 0.5850 | 0.4956 | 0.0614 | 1 |

**Table S 6: Bayes factor for the ABC-DL with different admixture patterns**. Model D4 is 8.074 times better at explaining the observed data than the second-best model (Model D3).

| SampleID | AccesionID | Population | Superpopulation | Reference |
|---|---|---|---|---|
| CEU01 | SAME123392 | Centre European Utah | European | 1000 Genomes |
| CEU02 | SAMN00801650 | Centre European Utah | European | 1000 Genomes |
| CHB01 | SAME124093 | Han Chinese | East Asia | 1000 Genomes |
| CHB02 | SAME123926 | Han Chinese | East Asia | 1000 Genomes |
| JHN01 | SAMEA3302682 | Ju\|'hoan North | South Africa San | SGDP, Mallick 2016 |

| | | | | |
|---|---|---|---|---|
| JHN02 | SAMEA3302894 | Ju\|'hoan North | South Africa San | SGDP, Mallick 2016 |
| LWK04 | SAMN00001054 | Luhya | East Africa | 1000 Genomes |
| LWK07 | SAMN00001112 | Luhya | East Africa | 1000 Genomes |
| QTR01 | SAMN03800116 | Qatar | Middle East | Fakhro 2016 |
| QTR02 | SAMN03800117 | Qatar | Middle East | Fakhro 2016 |
| TUN10 | SAMEA4969309 | Tunisia Arab | North Africa Arab | Serra-Vidal 2019 |
| TUN11 | SAMEA4969310 | Tunisia Arab | North Africa Arab | Serra-Vidal 2019 |
| TUN12 | SAMEA4969294 | Tunisia Chenini | North Africa Amazigh | Serra-Vidal 2019 |
| TUN13 | SAMEA4969295 | Tunisia Chenini | North Africa Amazigh | Serra-Vidal 2019 |
| YRI01 | SAME122984 | Yoruba | West Africa | 1000 Genomes |
| YRI02 | SAME125386 | Yoruba | West Africa | 1000 Genomes |

**Table S 7: Samples for the demographic analysis of North Africa**

| **Demographic Event** | **Prior probability of inclusion in GP4PG** |
|---|---|
| Change in Ne | 1 |
| Extintion event of a topodeme | 1 |
| Expansion event of a topodeme | 1 |
| Change in migration rate | 1 |
| Admixture event from one topodeme to another | 0.3 |

**Table S 8: Possible demographic events in GP4PG algorithm**.

| Parameter | Distribution | A | B | C | D | E | F | G |
|---|---|---|---|---|---|---|---|---|
| misclassification | U(0.0,5.0E-4) | X | X | X | X | X | X | X |
| NeSan | U(10000.0,90000.0) | X | X | X | X | X | X | X |
| NeYoruba (WAf) | U(10000.0,90000.0) | X | X | X | X | X | X | X |
| NeLuhya (EAf) | U(10000.0,60000.0) | X | X | X | X | X | X | X |
| NeTunisia_Chenini (NAfb) | U(1000.0,20000.0) | X | X | X | X | X | X | X |
| NeTunisia (NAfa) | U(1000.0,40000.0) | X | X | X | X | X | X | X |
| NeQatar (ME) | U(1000.0,40000.0) | X | X | X | X | X | X | X |
| NeCEU (EU) | U(1000.0,40000.0) | X | X | X | X | X | X | X |
| NeHan (EAs) | U(1000.0,40000.0) | X | X | X | X | X | X | X |
| tME_EU | U(200.0,800.0) | X | | X | | | | |
| NeME_EU | U(1000.0,20000.0) | X | | X | | | | |
| tNAfb_NAfa | U(10.0,150.0) | X | X | X | | | | |
| NeNAfb_NAfa | U(1000.0,20000.0) | X | X | X | | | | |
| tME_EU_EAs | U(tME_EU,3000.0) | X | | | | | | |

| | | | | | | | | |
|---|---|---|---|---|---|---|---|---|
| NeME_EU_EAs | U(1000.0,7000.0) | X | | | | | | |
| tWAf_NAfb_NAfa | U(tNAfb_NAfa,600.0) | X | | | | | | |
| NeWAf_NAfb_NAfa | U(1000.0,30000.0) | X | | | | | | |
| tEAf_ME_EU_EAs | U(tME_EU_EAs,4000.0) | X | | | | | | |
| NeEAf_ME_EU_EAs | U(1000.0,10000.0) | X | | | | | | |
| tWAf_EAf_EAs_ME_EU_NAfb_NAfa | U(tEAf_ME_EU_EAs,6000.0) | X | X | X | X | X | X | X |
| NeWAf_EAf_EAs_ME_EU_NAfb_NAfa | U(1000.0,30000.0) | X | X | X | X | X | X | X |
| tSan_WAf_EAf_EAs_ME_EU_NAfb_NAfa | U(tWAf_EAf_EAs_ME_EU_NAfb_NAfa, 12500.0) | X | X | X | X | X | X | X |
| NeSan_WAf_EAf_EAs_ME_EU_NAfb_NAfa | U(1000.0,30000.0) | X | X | X | X | X | X | X |
| tNAfb_NAfa_ME | U(tNAfb_NAfa,600.0) | | X | | | | | |
| NeNAfb_NAfa_ME | U(1000.0,15000.0) | | X | | | | | |
| tNAfb_NAfa_ME_EU | U(tNAfb_NAfa_ME,800.0) | | X | | | | | |
| NeNAfb_NAfa_ME_EU | U(1000.0,20000.0) | | X | X | X | | X | |
| tNAfb_NAfa_ME_EU_EAs | U(tNAfb_NAfa_ME_EU,3000.0) | | X | X | X | | X | |
| NeNAfb_NAfa_ME_EU_EAs | U(1000.0,7000.0) | | X | X | X | | X | |
| tNAfb_NAfa_ME_EU_EAs_EAf | U(tNAfb_NAfa_ME_EU_EAs,4000.0) | | X | X | X | | X | X |
| NeNAfb_NAfa_ME_EU_EAs_EAf | U(1000.0,30000.0) | | X | X | X | | X | X |
| tNAfb_NAfa_ME_EU | U(tME_EU,1000.0) | | | X | | | | |
| tNAfa_ME | U(150.0,600.0) | | | | X | X | X | X |
| NeNAfa_ME | U(1000.0,20000.0) | | | | X | X | X | X |
| tNAfa_ME_EU | U(tNAfa_ME,800.0) | | | | X | X | | X |
| NeNAfa_ME_EU | U(1000.0,20000.0) | | | | X | X | | X |
| tNAfb_NAfa_ME_EU | U(tNAfa_ME_EU,1000.0) | | | | X | | | |
| tNAfa_ME_EU_EAs | U(tNAfa_ME_EU,1000.0) | | | | | X | | |
| NeNAfa_ME_EU_EAs | U(1000.0,7000.0) | | | | | X | | X |
| tWAf_NAfb | U(150.0,600.0) | | | | | X | | |
| NeWAf_NAfb | U(1000.0,30000.0) | | | | | X | | |
| tNAfa_ME_EU_EAs_EAf | U(tNAfa_ME_EU_EAs,4000.0) | | | | | X | | |

| | | | | |
|---|---|---|---|---|
| NeNAfa_ME_EU_EAs_EAf | U(1000.0,30000.0) | X | | |
| tNAfb_NAfa_ME | U(tNAfb_NAfa,800.0) | | X | |
| NeNAfb_NAfa_ME | U(1000.0,20000.0) | | X | |
| tNAfb_NAfa_ME_EU | U(tNAfb_NAfa_ME,1000.0) | | X | |
| tNAfa_ME_EU_EAs | U(tNAfa_ME_EU,3000.0) | | | X |
| tEAf_NAfb | U(150.0,1000.0) | | | X |
| NeEAf_NAfb | U(1000.0,30000.0) | | | X |

**Table S 9: Parameters and prior distributions of the seven considered models in Fig. S4**

| Parameter | Distribution | D | D<br>2 | D<br>3 | D<br>4 | D<br>5 |
|---|---|---|---|---|---|---|
| misclassification | U(0.0, 5.0E-4) | X | X | X | X | X |
| NeSan | U(10000.0,90000.0) | X | X | X | X | X |
| NeYoruba | U(10000.0,90000.0) | X | X | X | X | X |
| NeLuhya | U(10000.0,60000.0) | X | X | X | X | X |
| NeTunisia_Chenini | U(1000.0,20000.0) | X | X | X | X | X |
| NeTunisia | U(1000.0,40000.0) | X | X | X | X | X |
| NeQatar | U(1000.0,40000.0) | X | X | X | X | X |
| NeCEU | U(1000.0,40000.0) | X | X | X | X | X |
| NeHan | U(1000.0,40000.0) | X | X | X | X | X |
| NeBasal_Eurasian_ghost (BEi) | U(1000.0,20000.0) | | | X | X | X |
| NeAfrican_ghost (Xa) | U(1000.0,20000.0) | | | | X | X |
| migrationNAfa_NAfb | U(0.0,5.0E-4) | | X | X | X | |
| migrationNAfb_NAfa | U(0.0,5.0E-4) | | X | X | X | |
| migrationME_NAfb | U(0.0,5.0E-4) | | X | X | X | |
| migrationNAfb_ME | U(0.0,5.0E-4) | | X | X | X | |
| migrationME_NAfa | U(0.0,5.0E-4) | | X | X | X | |
| migrationNAfa_ME | U(0.0,5.0E-4) | | X | X | X | |
| migrationEU_ME | U(0.0,5.0E-4) | | X | X | X | |
| migrationEU_NAfb | U(0.0,5.0E-4) | | X | X | X | |
| migrationNAfb_EU | U(0.0,5.0E-4) | | X | X | X | |
| migrationEU_NAfa | U(0.0,5.0E-4) | | X | X | X | |
| migrationNAfa_EU | U(0.0,5.0E-4) | | X | X | X | |
| migrationEAs_EU | U(0.0,5.0E-4) | | X | X | X | |
| migrationEU_EAs | U(0.0,5.0E-4) | | X | X | X | |
| migrationEAf_NAfb | U(0.0,5.0E-4) | | X | X | X | |
| migrationNAfb_EAf | U(0.0,5.0E-4) | | X | X | X | |
| migrationEAf_NAfa | U(0.0,5.0E-4) | | X | X | X | |
| migrationNAfa_EAf | U(0.0,5.0E-4) | | X | X | X | |
| migrationEAf_ME | U(0.0,5.0E-4) | | X | X | X | |

| | | | | | | |
|---|---|---|---|---|---|---|
| migrationME_EAf | U(0.0,5.0E-4) | | X | X | X | |
| migrationEAf_WAf | U(0.0,5.0E-4) | | X | X | X | |
| migrationWAf_EAf | U(0.0,5.0E-4) | | X | X | X | |
| migrationWAf_NAfb | U(0.0,5.0E-4) | | X | X | X | |
| migrationNAfb_WAf | U(0.0,5.0E-4) | | X | X | X | |
| migrationWAf_NAfa | U(0.0,5.0E-4) | | X | X | X | |
| migrationNAfa_WAf | U(0.0,5.0E-4) | | X | X | X | |
| migrationSan_to_WAf | U(0.0,5.0E-4) | | X | X | X | |
| migrationSan_to_EAf | U(0.0,5.0E-4) | | X | X | X | |
| tNAfa_ME | U(150.0,600.0) | X | X | X | X | X |
| NeNAfa_ME | U(1000.0,20000.0) | X | X | X | X | X |
| tNAfa_ME_EU | U(tNAfa_ME,800.0) | X | X | X | X | X |
| NeNAfa_ME_EU | U(1000.0,20000.0) | X | X | X | X | X |
| tNAfb_NAfa_ME_EU | U(tNAfa_ME_EU,1000.0) | X | X | X | X | X |
| tNAfb_NAfa_ME_EU_EAs | U(tNAfb_NAfa_ME_EU,20000.0) | X | X | X | X | X |
| NeNAfb_NAfa_ME_EU_EAs | U(1000.0,7000.0) | X | X | X | X | X |
| tNAfb_NAfa_ME_EU_EAs_EAf | U(tNAfb_NAfa_ME_EU_EAs,4000.0) | X | X | X | X | X |
| NeNAfb_NAfa_ME_EU_EAs_EAf | U(1000.0,30000.0) | X | X | X | X | X |
| tWAf_EAf_EAs_ME_EU_NAfb_NAfa | U(tEAf_ME_EU_EAs,6000.0) | X | X | X | X | X |
| NeWAf_EAf_EAs_ME_EU_NAfb_NAfa | U(1000.0,30000.0) | X | X | X | X | X |
| tSan_WAf_EAf_EAs_ME_EU_NAfb_NAfa | U(tWAf_EAf_EAs_ME_EU_NAfb_NAfa, 12500.0) | X | X | X | X | X |
| NeSan_WAf_EAf_EAs_ME_EU_NAfb_NAfa | U(1000.0,30000.0) | X | X | X | X | X |
| tSan_WAf_EAf_EAs_ME_EU_NAfb_NAfa_Xa | U(tNAfb_NAfa_ME_EU_BEi_EAs_WAf_San, 14000.0) | | | | X | X |
| NeSan_WAf_EAf_EAs_ME_EU_NAfb_NAfa_Xa | U(1000.0,30000.0) | | | | X | X |
| tNAfb_NAfa_ME_EU_BEi | U(tNAfb_NAfa_ME_EU,2000.0) | | | X | X | X |
| NeNAfb_NAfa_ME_EU_BEi | U(1000.0,20000.0) | | | X | X | X |
| tAdmxME_NA | U(60.0,tNAfa_ME) | | X | X | X | X |

| | | | | | |
|---|---|---|---|---|---|
| admixtureME_NA | U(0.001,0.20) | X | X | X | X |
| tAdmxEU_NA | U(20.0,tNAfa_ME) | | | X | X |
| admixtureEU_NA | U(0.001,0.10) | | | X | X |
| tAdmxEU_NAb | U(20.0,tNAfa_ME) | | | X | X |
| admixtureEU_NAb | U(0.001,0.10) | | | X | X |
| tAdmxME_NAb | U(60.0,tNAfa_ME) | | | X | X |
| admixtureME_NAb | U(0.001,0.20) | | | X | X |
| tAdmxMENA_Amazigh | U(tNAfa_ME,tNAfa_ME_EU) | X | X | X | X |
| admixtureMENA_Amazigh | U(0.001,0.10) | X | X | X | X |
| tAdmxWAf_Amazigh | U(10.0,tNAfb_NAfa_ME_EU) | X | X | X | X |
| admixtureWAf_Amazigh | U(0.001,0.10) | X | X | X | X |
| tAdmxWAf_Arab | U(10.0,tNAfa_ME) | X | X | X | X |
| admixtureWAf_Arab | U(0.001,0.10) | X | X | X | X |
| tAdmxEAf_Amazigh | U(10.0,tNAfb_NAfa_ME_EU) | X | X | X | X |
| admixtureEAf_Amazigh | U(0.001,0.10) | X | X | X | X |
| tAdmxEAf_Arab | U(10.0,tNAfa_ME) | X | X | X | X |
| admixtureEAf_Arab | U(0.001,0.10) | X | X | X | X |
| tAdmxBEi_MENAU | U(tNAfa_ME_EU, tNAfb_NAfa_ME_EU) | | X | X | X |
| admixtureBEi_MENAU | U(0.001,0.20) | | X | X | X |
| tAdmxBEi_AMENAU | U(tNAfb_NAfa_ME_EU, tNAfb_NAfa_ME_EU_BEi) | | X | X | X |
| admixtureBEi_AMENAU | U(0.001,0.20) | | X | X | X |
| tAdmxXa_San | U(700.0,tNAfb_NAfa_ME_EU_BEi_EAs_WAf_San) | | | X | X |
| admixtureXa_San | U(0.001,0.05) | | | X | X |
| tAdmxXa_WAf | U(700.0,tNAfb_NAfa_ME_EU_BEi_EAs_WAf) | | | X | X |
| admixtureXa_WAf | U(0.001,0.05) | | | X | X |

**Table S 10: Parameters and prior distributions of the five considered models in Fig. 2.**

Fig. S. 1

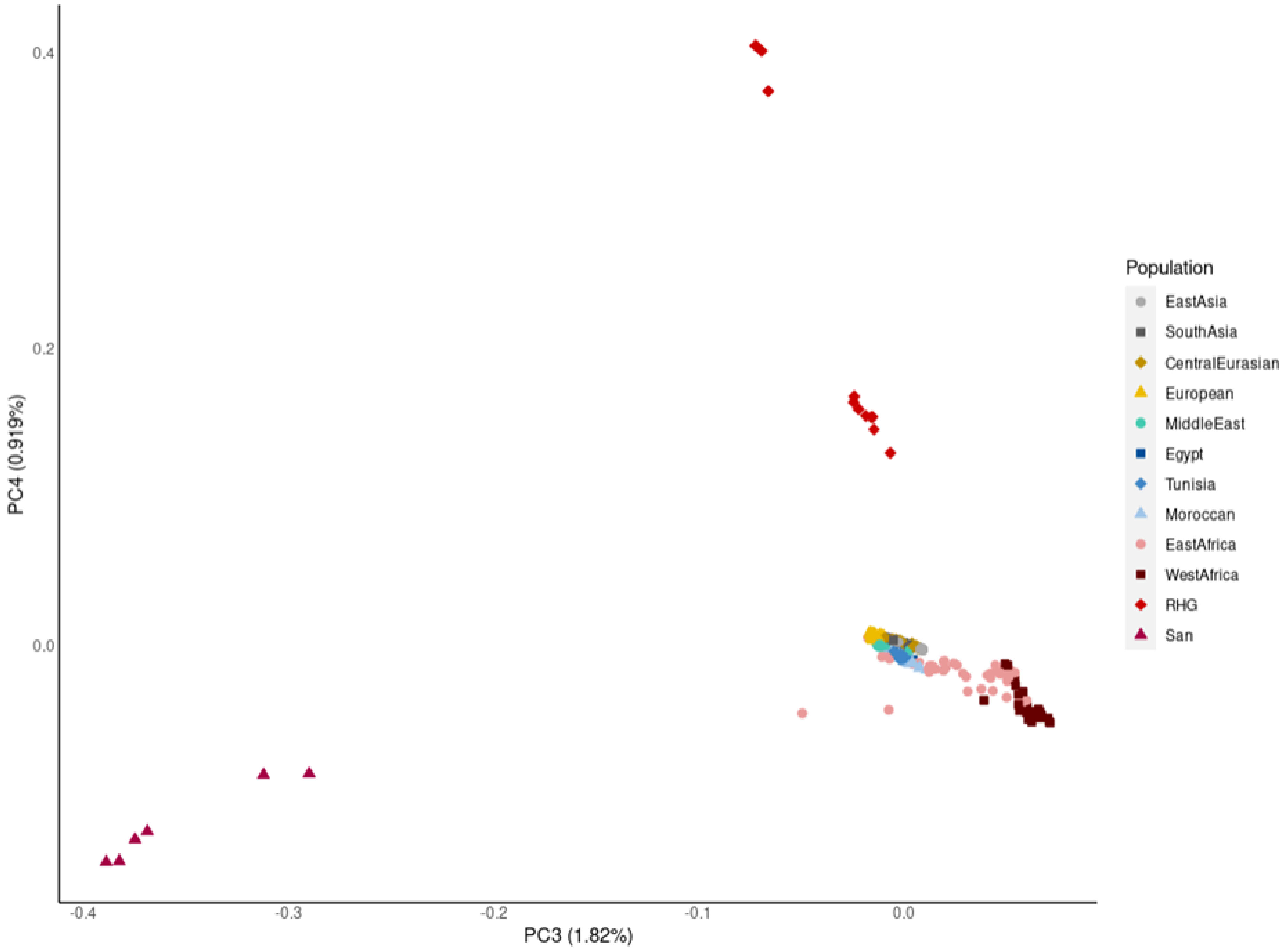

0.4
0.2
0.0
PC4 (0.919%)
-0.4
-0.3
-0.2
-0.1
0.0
PC3 (1.82%)
Population
EastAsia
SouthAsia
CentralEurasian
European
MiddleEast
Egypt
Tunisia
Moroccan
EastAfrica
WestAfrica
RHG
San

**Fig. S. 2**

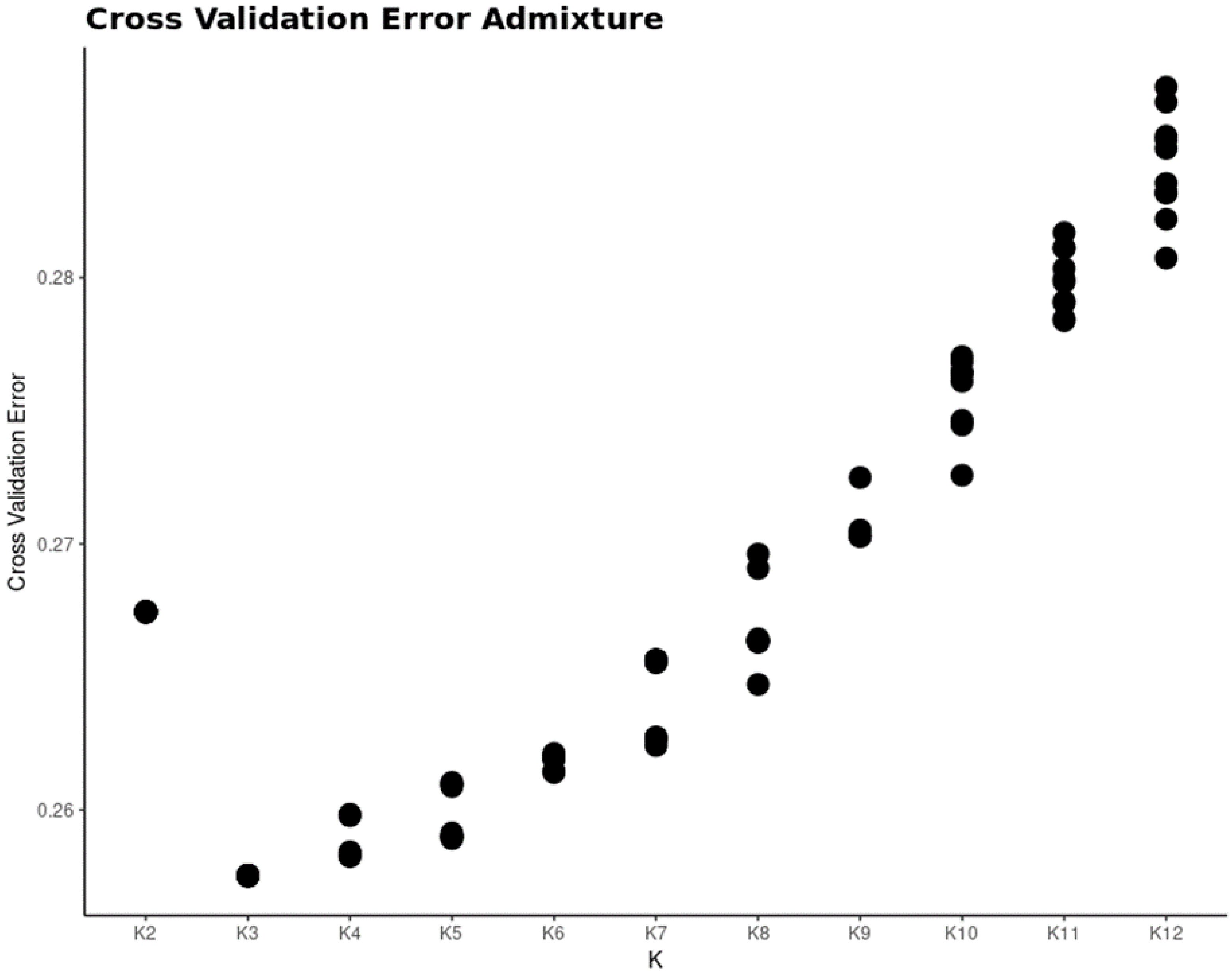

Cross Validation Error Admixture
Cross Validation Error
0.28
0.27
0.26
K2
K3
K4
K5
K6
K7
K8
K9
K10
K11
K12
K

### Fig. S. 3

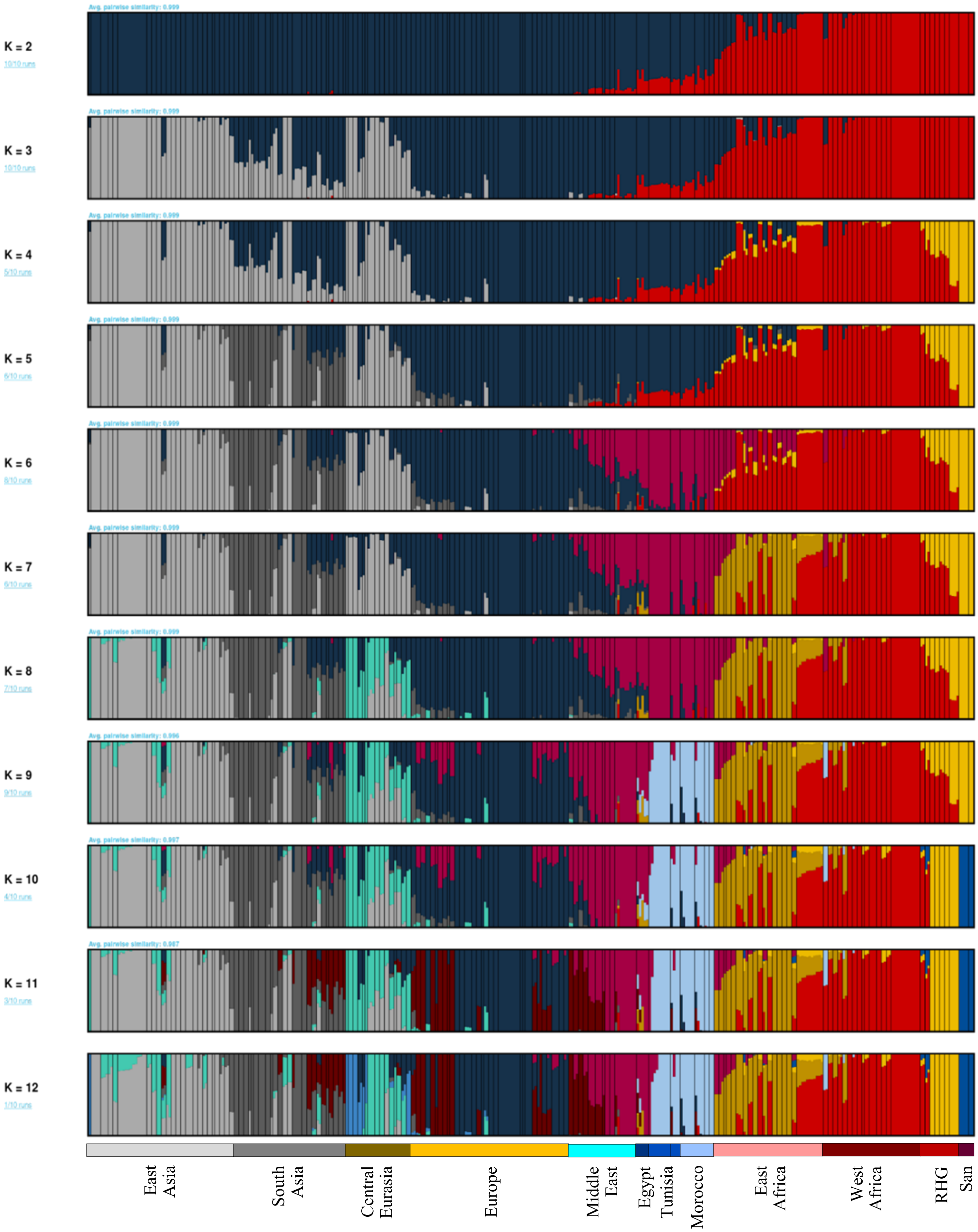
Avg. pairwise similarity: 0.999
K = 2
10/10 runs
Avg. pairwise similarity: 0.999
K = 3
10/10 runs
Avg. pairwise similarity: 0.999
K = 4
5/10 runs
Avg. pairwise similarity: 0.999
K = 5
6/10 runs
Avg. pairwise similarity: 0.999
K = 6
8/10 runs
Avg. pairwise similarity: 0.999
K = 7
6/10 runs
Avg. pairwise similarity: 0.999
K = 8
7/10 runs
Avg. pairwise similarity: 0.996
K = 9
9/10 runs
Avg. pairwise similarity: 0.997
K = 10
4/10 runs
Avg. pairwise similarity: 0.987
K = 11
3/10 runs
K = 12
1/10 runs
East Asia
South Asia
Central Eurasia
Europe
Middle East
Egypt
Tunisia
Morocco
East Africa
West Africa
RHG
San

### Fig. S. 4

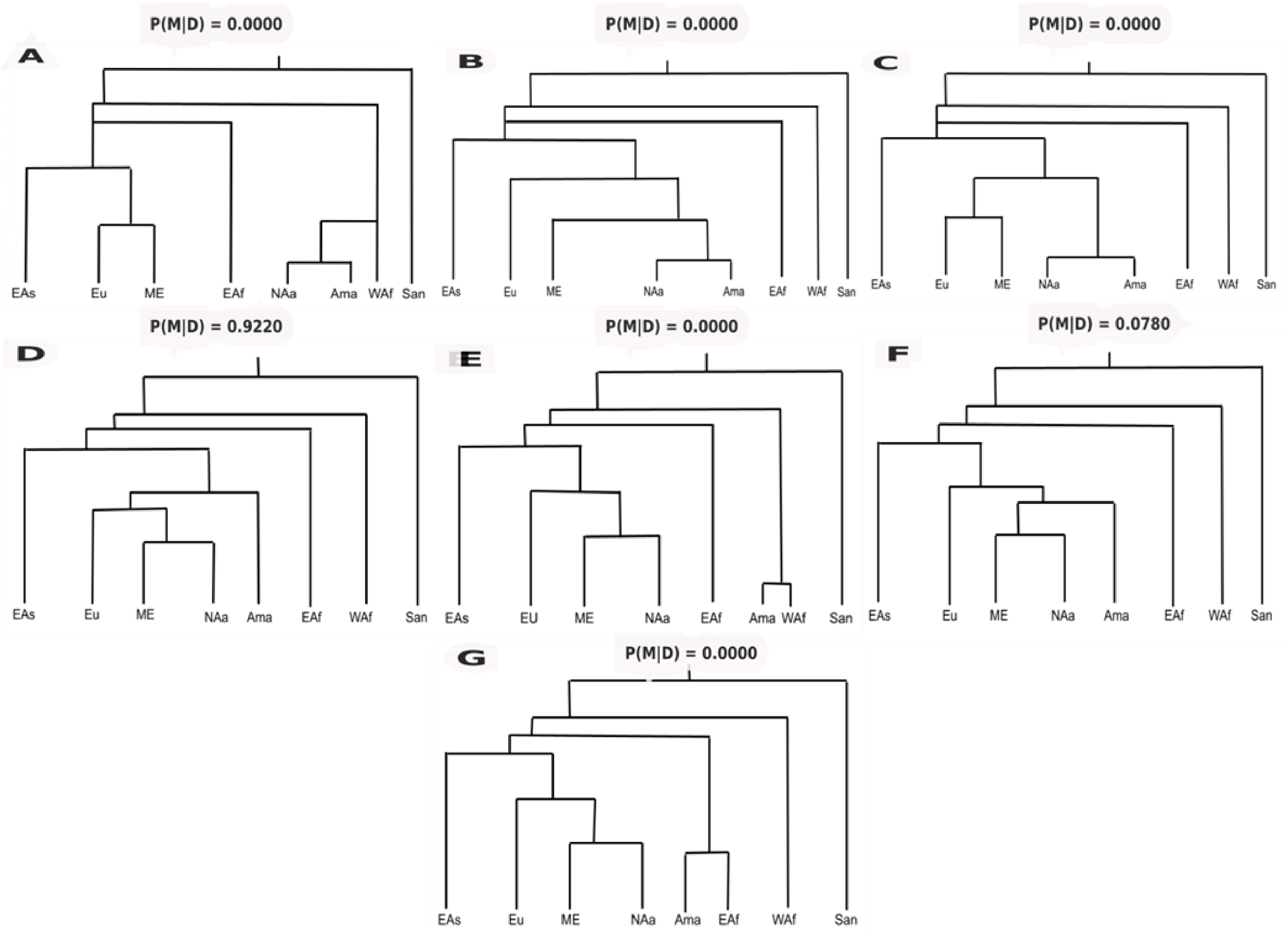
A
P(M|D) = 0.0000
EAs Eu ME EAf NAa Ama WAf San
B
P(M|D) = 0.0000
EAs Eu ME NAa Ama EAf WAf San
C
P(M|D) = 0.0000
EAs Eu ME NAa Ama EAf WAf San
D
P(M|D) = 0.9220
EAs Eu ME NAa Ama EAf WAf San
E
P(M|D) = 0.0000
EAs EU ME NAa EAf Ama WAf San
F
P(M|D) = 0.0780
EAs Eu ME NAa Ama EAf WAf San
G
P(M|D) = 0.0000
EAs Eu ME NAa Ama EAf WAf San

**Fig. S. 5**

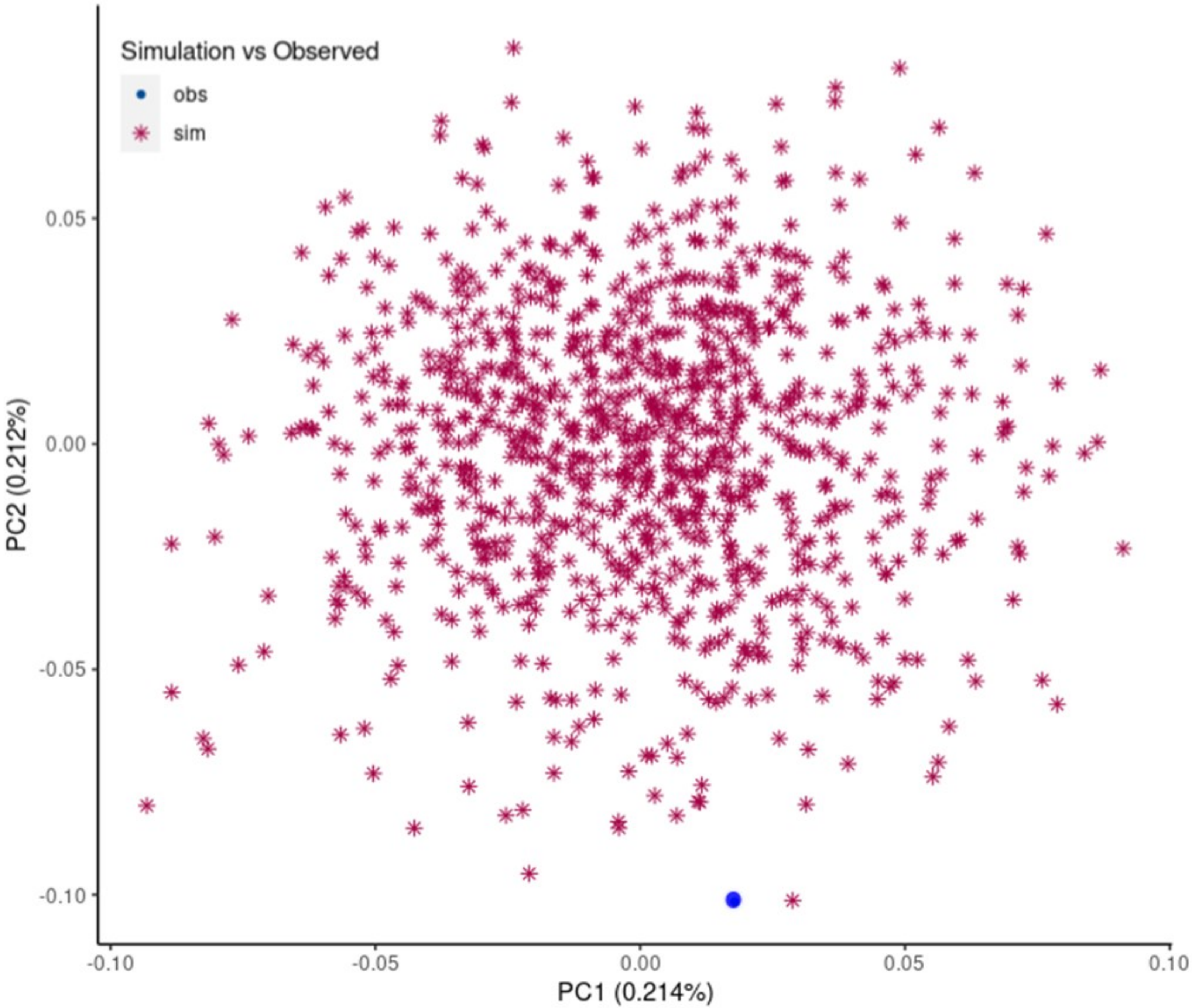

Simulation vs Observed
obs
sim
0.05
0.00
-0.05
-0.10
PC2 (0.212%)
-0.10
-0.05
0.00
0.05
0.10
PC1 (0.214%)

**Fig. S. 6**

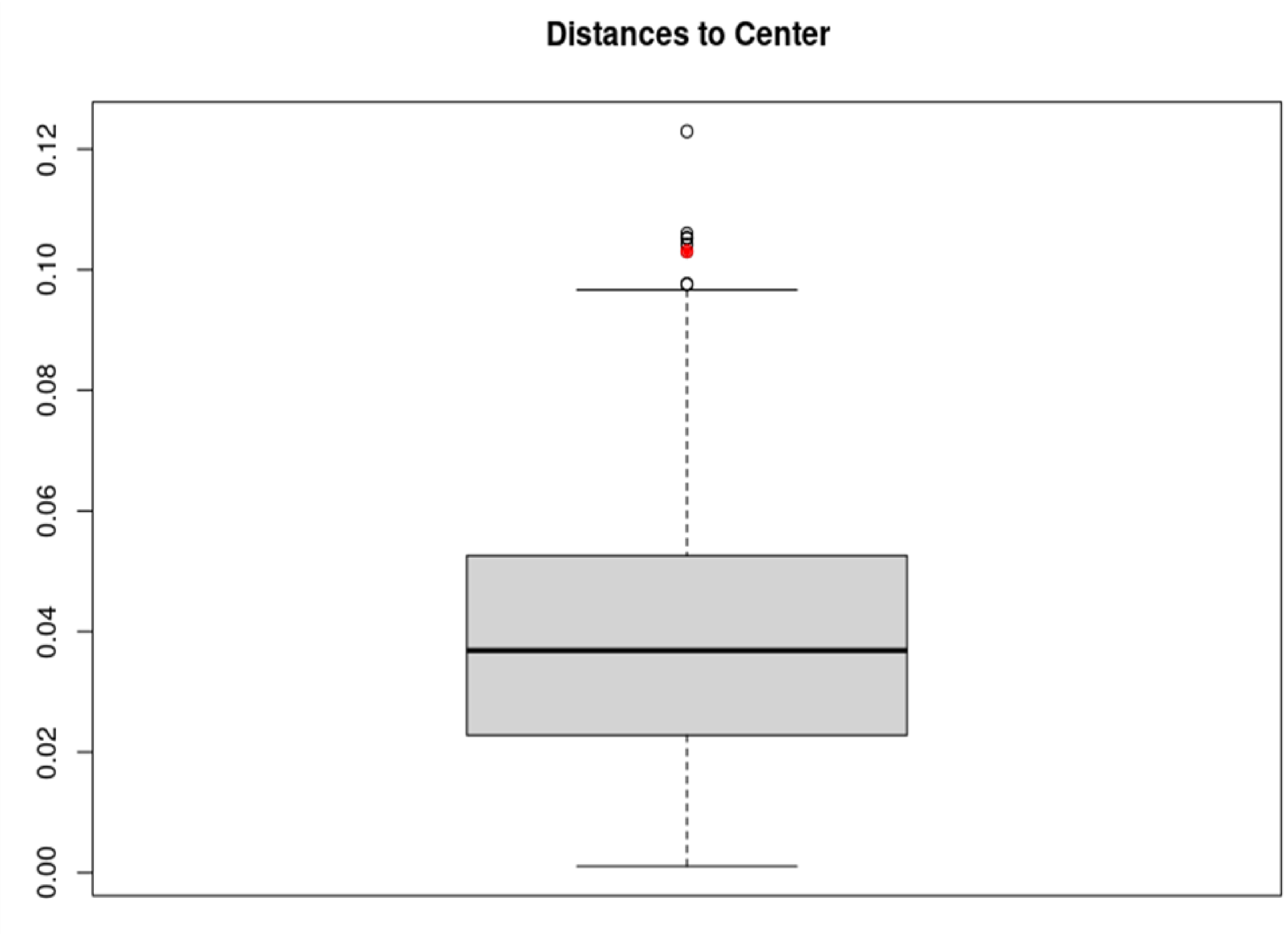

**Fig. S. 7**

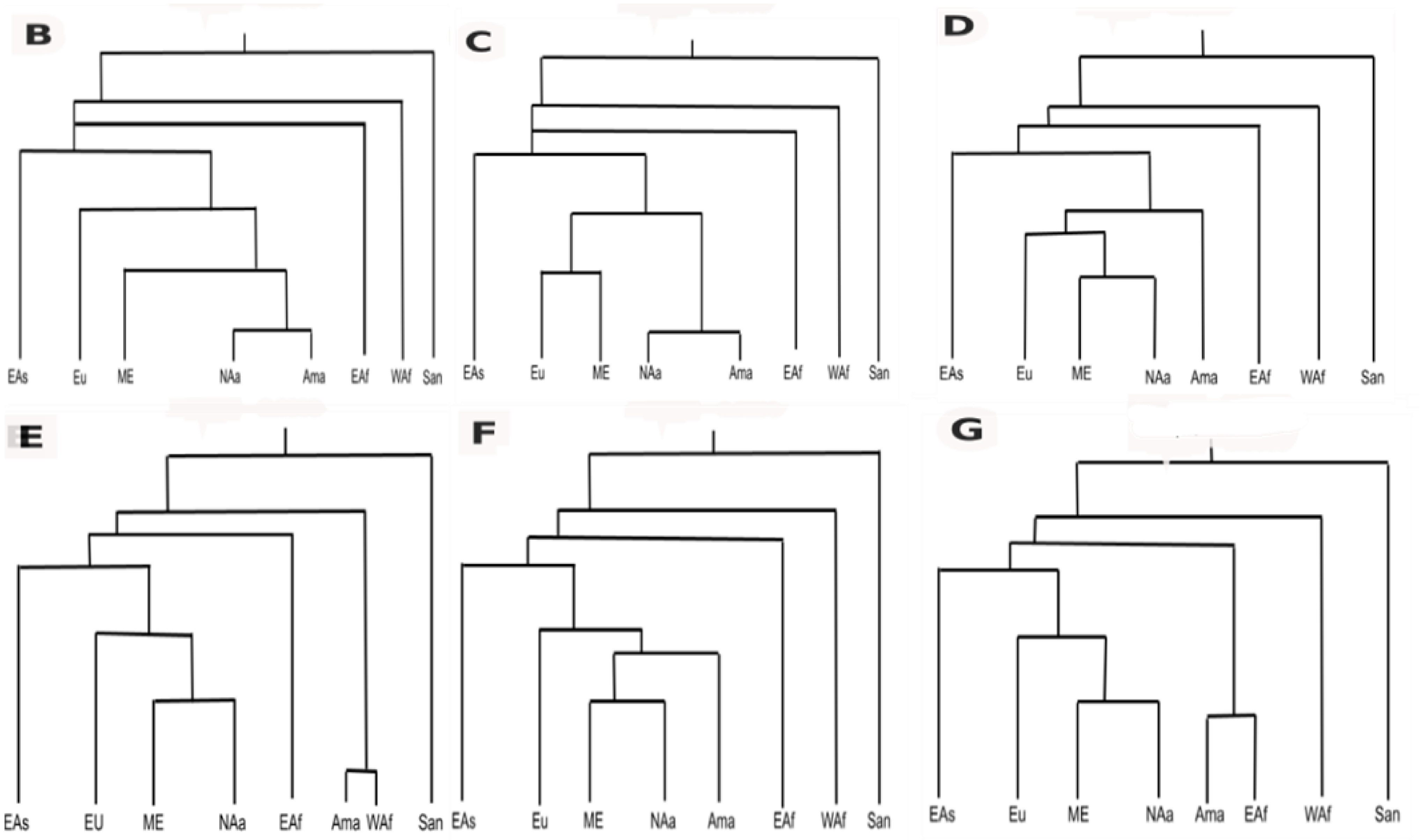
B
EAs Eu ME NAa Ama EAf WAf San
C
EAs Eu ME NAa Ama EAf WAf San
D
EAs Eu ME NAa Ama EAf WAf San
E
EAs EU ME NAa EAf Ama WAf San
F
EAs Eu ME NAa Ama EAf WAf San
G
EAs Eu ME NAa Ama EAf WAf San

**Fig. S. 8**

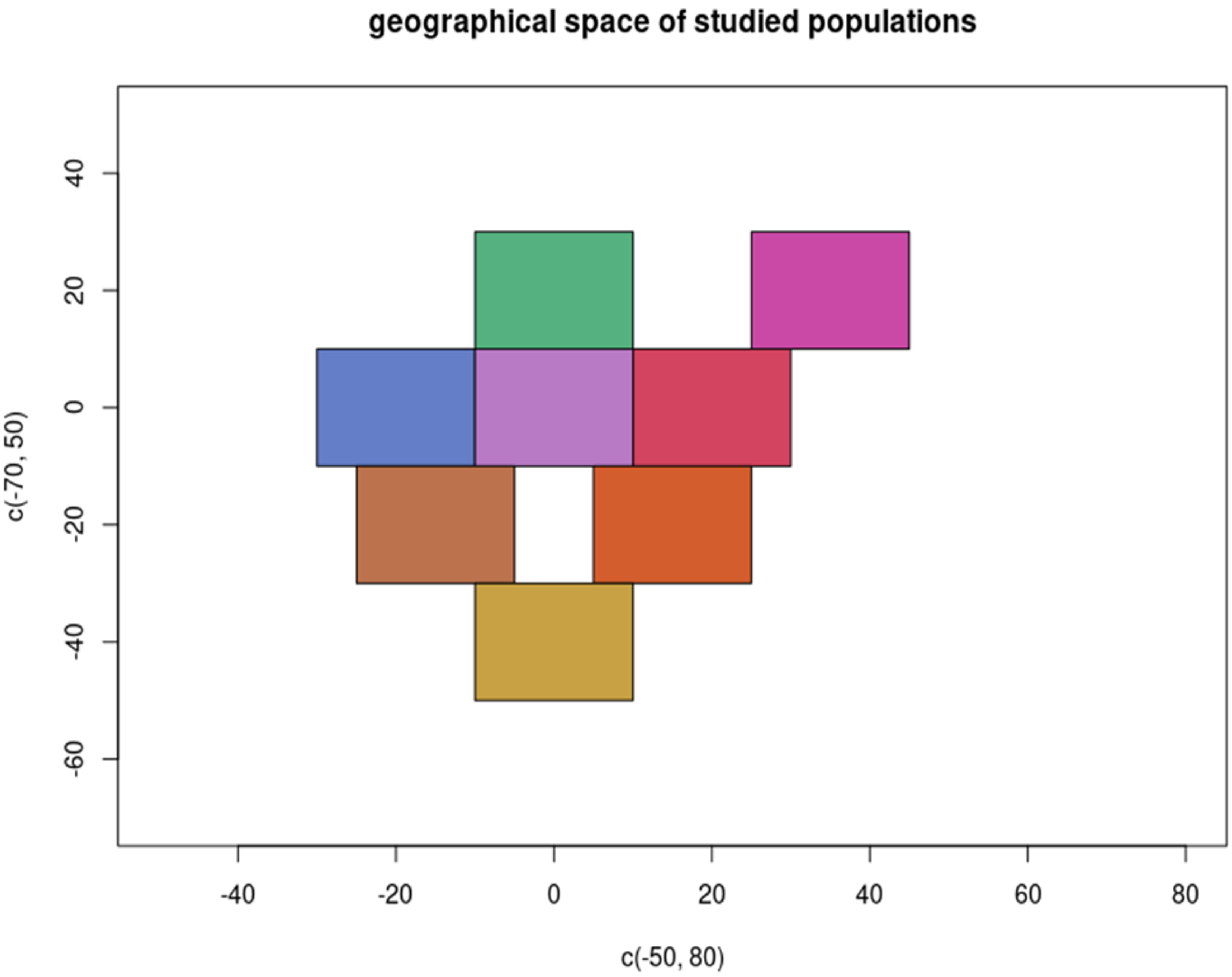
geographical space of studied populations
40
20
0
-20
-40
-60
c(-70, 50)
-40
-20
0
20
40
60
80
c(-50, 80)

**Fig. S. 9**

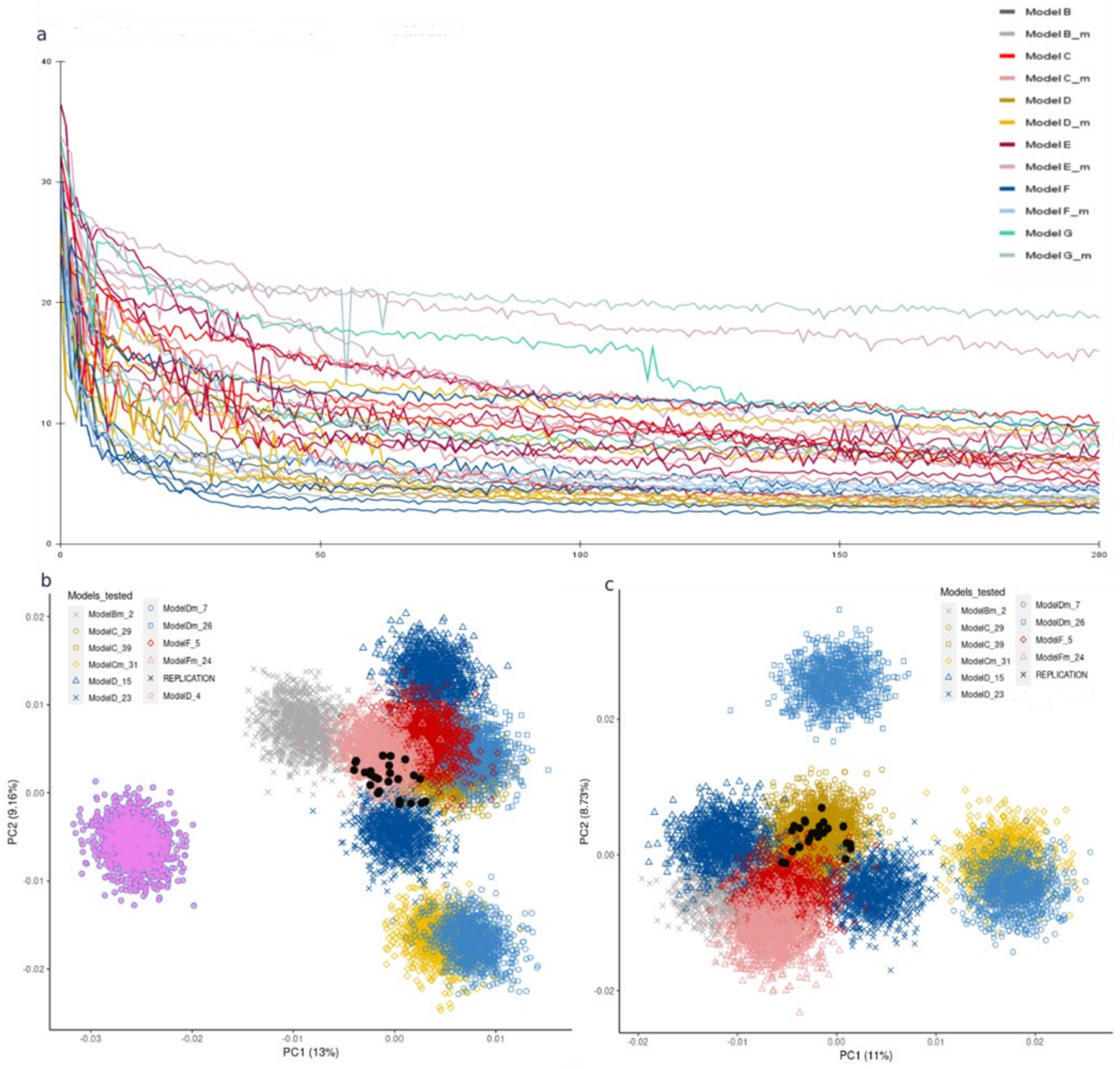
a
Model B
Model B_m
Model C
Model C_m
Model D
Model D_m
Model E
Model E_m
Model F
Model F_m
Model G
Model G_m
b
Models_tested
ModelBm_2
ModelC_29
ModelC_39
ModelCm_31
ModelD_15
ModelD_23
ModelDm_7
ModelDm_26
ModelF_5
ModelFm_24
REPLICATION
ModelD_4
PC1 (13%)
PC2 (9.16%)
c
PC1 (11%)
PC2 (8.73%)

### Fig. S. 10

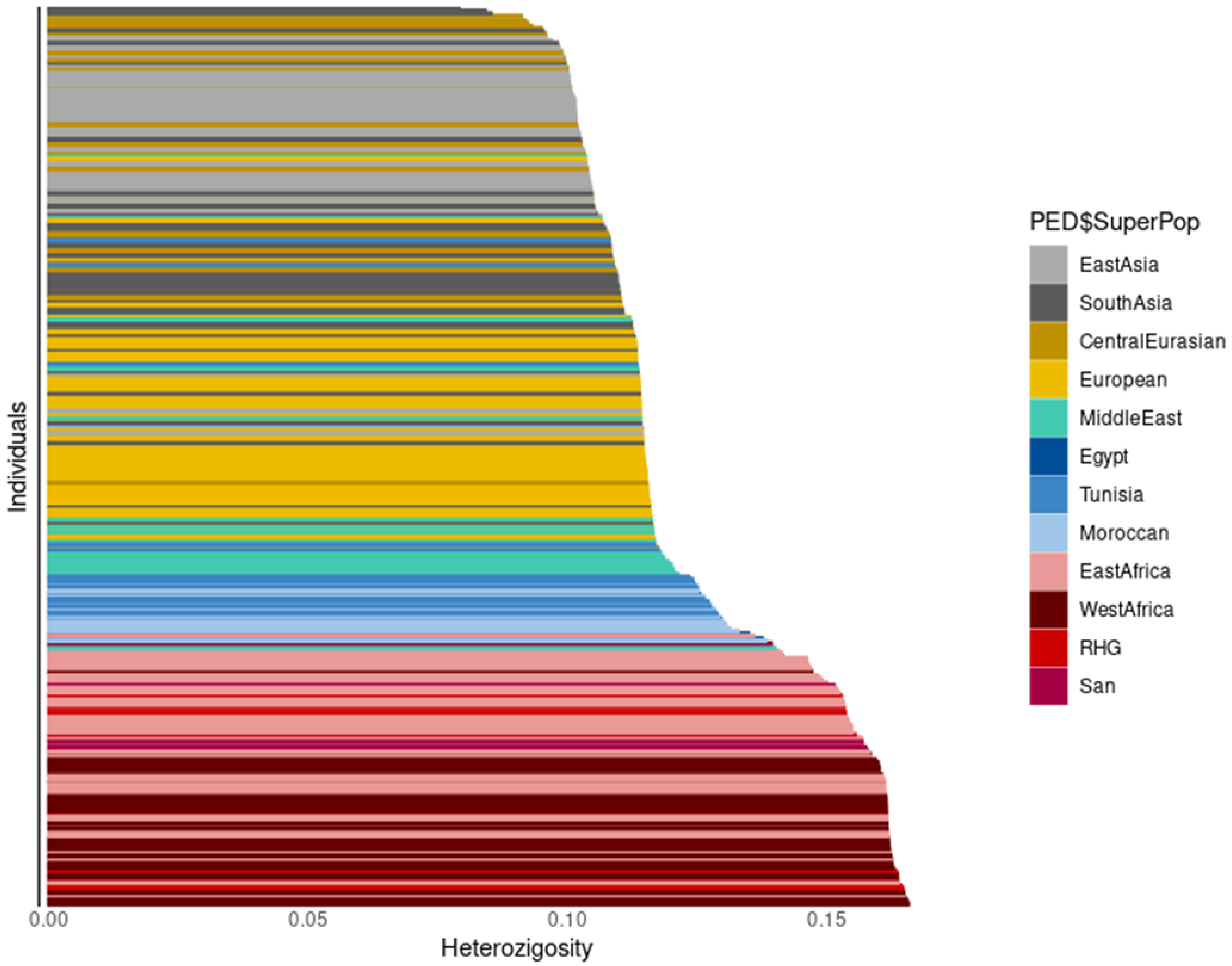

Individuals
0.00
0.05
0.10
0.15
Heterozigosity
PED$SuperPop
EastAsia
SouthAsia
CentralEurasian
European
MiddleEast
Egypt
Tunisia
Moroccan
EastAfrica
WestAfrica
RHG
San

**Fig. S. 11**

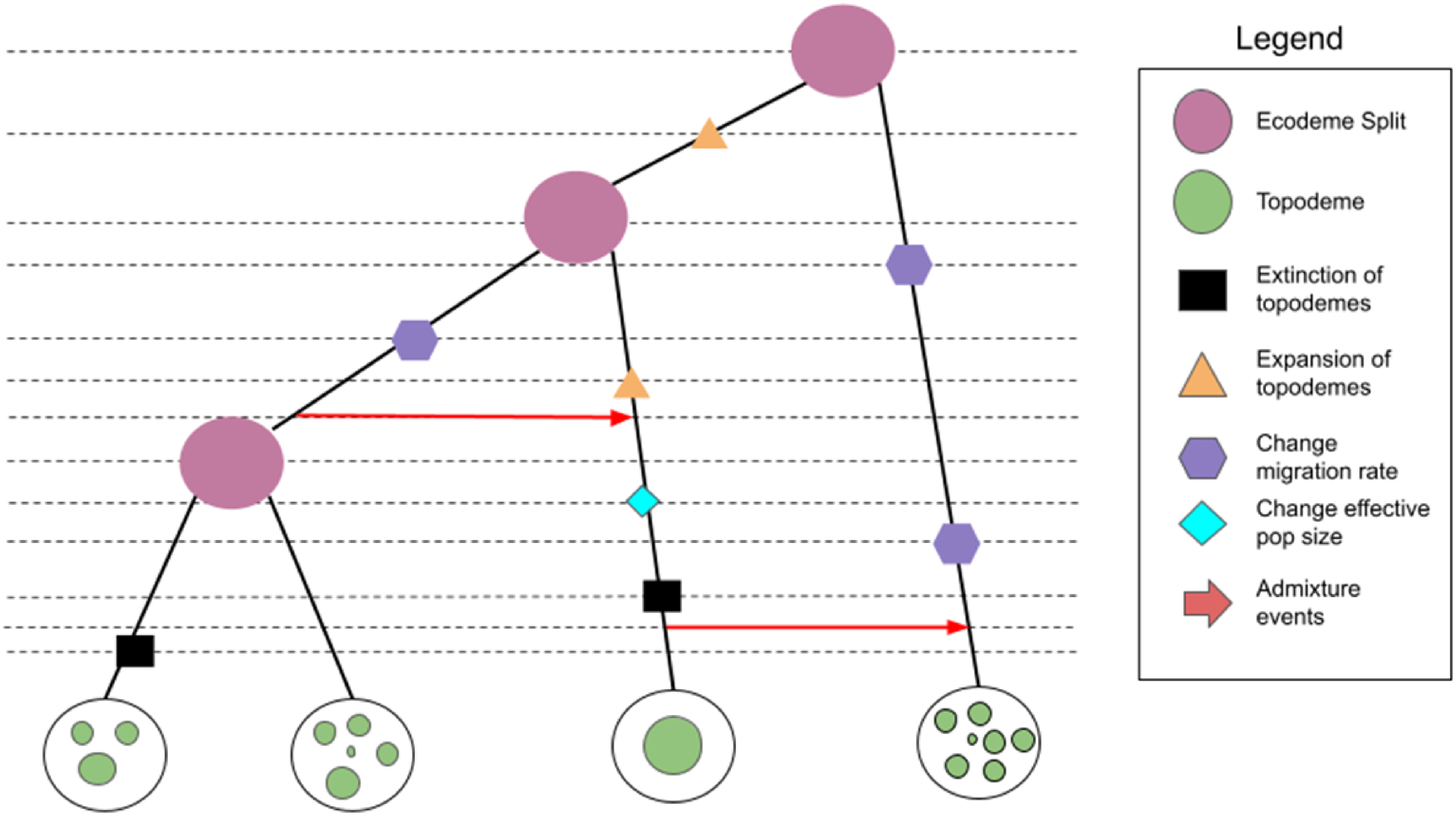
Legend
Ecodeme Split
Topodeme
Extinction of topodemes
Expansion of topodemes
Change migration rate
Change effective pop size
Admixture events
